# SIMplyBee: R package for simulating honeybee populations and breeding programs

**DOI:** 10.1101/2022.12.15.520571

**Authors:** Jana Obšteter, Laura K. Strachan, Jernej Bubnič, Janez Prešern, Gregor Gorjanc

## Abstract

**Background:** The Western honeybee is an economically important species globally, but has been experiencing colony losses that lead to economical damage and decreased genetic variability. This situation is spurring additional interest in honeybee breeding and conservation programs. Stochastic simulators are essential tools for rapid and low-cost testing of breeding programs and methods, yet no existing simulator allows for a detailed simulation of honeybee populations. Here we describe SIMplyBee, a holistic simulator of honeybee populations and breeding programs. SIMplyBee is an R package and hence freely available for installation from CRAN http://cran.r-project.org/package=SIMplyBee.

**Implementation:** SIMplyBee builds upon the stochastic simulator AlphaSimR that simulates individuals with their corresponding genomes and quantitative genetic values. To enable a honeybee specific simulation, we extended AlphaSimR by developing classes for global simulation parameters, `SimParamBee`, for a honeybee colony, `Colony`, and multiple colonies, `MultiColony`. We also developed functions to address major specificities of the honeybees: honeybee genome, haplo-diploid inheritance, social organisation, complementary sex determination, polyandry, colony events, and quantitative genetics of honeybees.

**Results:** We describe and show implementation regarding simulating a honeybee genome, creating a honeybee colony and its members, haplodiploid inheritance and complementary sex determination, colony events, creating and managing multiple colonies at once, and obtaining genomic data and honeybee quantitative genetics. Further documentation at http://SIMplyBee.info provides details on these operations and describes additional operations related to genomics, quantitative genetics, and other functionality.

**Discussion:** SIMplyBee is a holistic simulator of honeybee populations and breeding programs that simulates individual honeybees with their genomes, colonies with colony events, and individual- and colony-level quantitative values. SIMplyBee provides a research platform for testing breeding and conservation strategies and their effect on future genetic gain and variability. Future development of SIMplyBee will focus on improving the simulation of honeybee genomes, optimizing the performance of the simulator, and including spatial awareness to crossing functions and phenotype simulation. We welcome the honeybee genetics and breeding community to join us in the future development of SIMplyBee.

## Background

The Western honeybee (*Apis mellifera*) is an economically important species globally, playing a major role in pollination and food production. The value of insect pollinators is estimated at 150 billion euros per year worldwide, which is approximately 10 percent of the global agriculture production [1, 2, 3]. In recent decades, both wild and managed populations have been experiencing increased colony losses due to numerous biotic and abiotic factors [4, 5, 6]. Besides the economic loss, high colony mortality and human-mediated hybridisation have also driven the loss of within species diversity during the last century and put native subspecies at risk [7, 5, 8, 9]. Although honeybees are a diverse species that is differentiated into 7 evolutionary lineages and 33 subspecies [10, 11], two subspecies, *A. m. ligustica* and *A. m. carnica*, dominate the vast majority of commercial beekeeping operations [12]. The loss of genetic variability can decrease the fitness of the populations and further increases the susceptibility of populations to ecological and anthropogenic factors [5, 9].

Due to increased colony losses and decline in genetic diversity there has been an increased interest in honeybee management programs, either for breeding, conservation, or both. Breeding programs aim to improve honeybee production, behaviour, and resistance to pathogens, and manage genetic diversity that enables long-term response to selection. Conservation programs aim to preserve populations of endangered or native species by managing genetic diversity, reducing inbreeding depression, maintaining locally adaptive traits, and reducing the prevalence of pathogens.

The increased interest in honeybee breeding has spurred additional research in quantitative genetics of honeybees. Stochastic simulators are an essential tool for *in-silico* development and testing of quantitative genetic and statistical methods, and breeding strategies [13, 14, 15, 16]. While simulations rest on a number of assumptions, they enable cost effective and rapid testing of hypotheses before practical deployment. There are a number of simulators available for the most commercially interesting mammalian or plant species [13, 15, 16]. Due to the differences in biology and social organisation, these simulators cannot simulate honeybee populations. Although there are existing honeybee simulators, they are either too simplistic, do not simulate genomes and genetic and phenotypic values of individual honeybees, lack flexibility to simulate the honeybee colony life cycle or the entire breeding program, or are not available as open source [14, 17]. One of such honeybee simulators is BeeSim [14] that accounts for the quantitative genetics of the honeybees, but simulates quantitative values on the colony level, does not account for the colony events, and is also not publicly available. Another honeybee simulator, BEEHAVE [17], simulates colony and population dynamics as well as environmental variation to explore causes of colony failures and colony performance, but does not include genetics.

The aim of this work was to develop a holistic simulator of honeybee population management programs, SIMplyBee. SIMplyBee simulates i) genomes and quantitative values of individual honeybees as well as whole colonies, ii) major biological, reproductive, and organisational specificities of the honeybees, and iii) colony events. SIMplyBee is freely available for installation from CRAN (http://cran.r-project.org/package=SIMplyBee) with extensive help pages, examples, and vignettes. See also http://www.SIMplyBee.info. We welcome contributions from the community at https://www.github.com/HighlanderLab/SIMplyBee. In the following, we describe the theory and technical implementation in the SIMplyBee, demonstrate the use of SIMplyBee, and discuss the potential use of SIMplyBee and our plans for its future development.

## Implementation

SIMplyBee builds upon an established simulator, AlphaSimR [18, 15], and shares its core simulation principles and functionality. AlphaSimR is a stochastic simulator that simulates individuals with their corresponding genomes and quantitative genetic and phenotypic values. The most important classes in AlphaSimR are the `SimParam` class for global simulation parameters and the `Pop` class for objects that hold a group of individuals with their individual identification, parent identifications, as well as genomes and trait values.

To enable a honeybee specific simulation, we extended AlphaSimR by developing dedicated classes: `SimParamBee` for global simulation parameters, `Colony` for a honeybee colony, and `MultiColony` for multiple honeybee colonies. We also developed functions to simulate honeybee populations and its events and facilitate an inspection or analysis of the results. The functions address major specificities of the honeybees: honeybee genome, honeybe biology, including haplodiploid inheritance, complementary sex determination, social organisation, and polyandry, and colony events. Functionally, we organised these functions in five functional groups related to genome and genomic information, caste operations, colony and multicolony operations, quantitative genetics, and auxiliary operations. From the operational standpoint of the SIMplyBee R package, we separated these functions into four operational levels with respect to their simplest return objects: level 0 being auxiliary functions returning standard R class objects such as vectors, matrices, and lists; level 1 returning an AlphaSimR `Pop` class object; level 2 returning a SIMplyBee `Colony` class object; and level 3 returning a SIMplyBee `MultiColony` class object. The total codebase spans over 16,000 lines of R code, documentation, and unit tests.

## Results

Here, we present the SIMplyBee functionality. We briefly describe the biological mechanisms behind the functionality and demonstrate its use. We describe: i) simulating honeybee genome; ii) creating a honeybee colony and its members; iii) haplodiploid inheritance and the complementary sex determination locus *CSD*; iv) colony events; v) working with multiple colonies; and vi) honeybee genomics and quantitative genetics.

### Honeybee genome and initiating a honeybee simulation

To initiate the simulation we first need to simulate honeybee genomes and set simulation parameters (Figure 1). The honeybee genome is small in its physical length, only 250 million bp, but large in its genetic length, 23 Morgans, due to a very high recombination rate of 2.3 × 10^−7^ per bp [19]. In SIMplyBee, we generate honeybee genome sequences with the approximate (Markovian) coalescent simulator MaCS [20] according to the most complete honeybee demographic model [21]. We are currently simulating three subspecies: *A. m. ligustica*, *A. m. carnica*, and *A. m. mellifera*.

**Figure 1.**
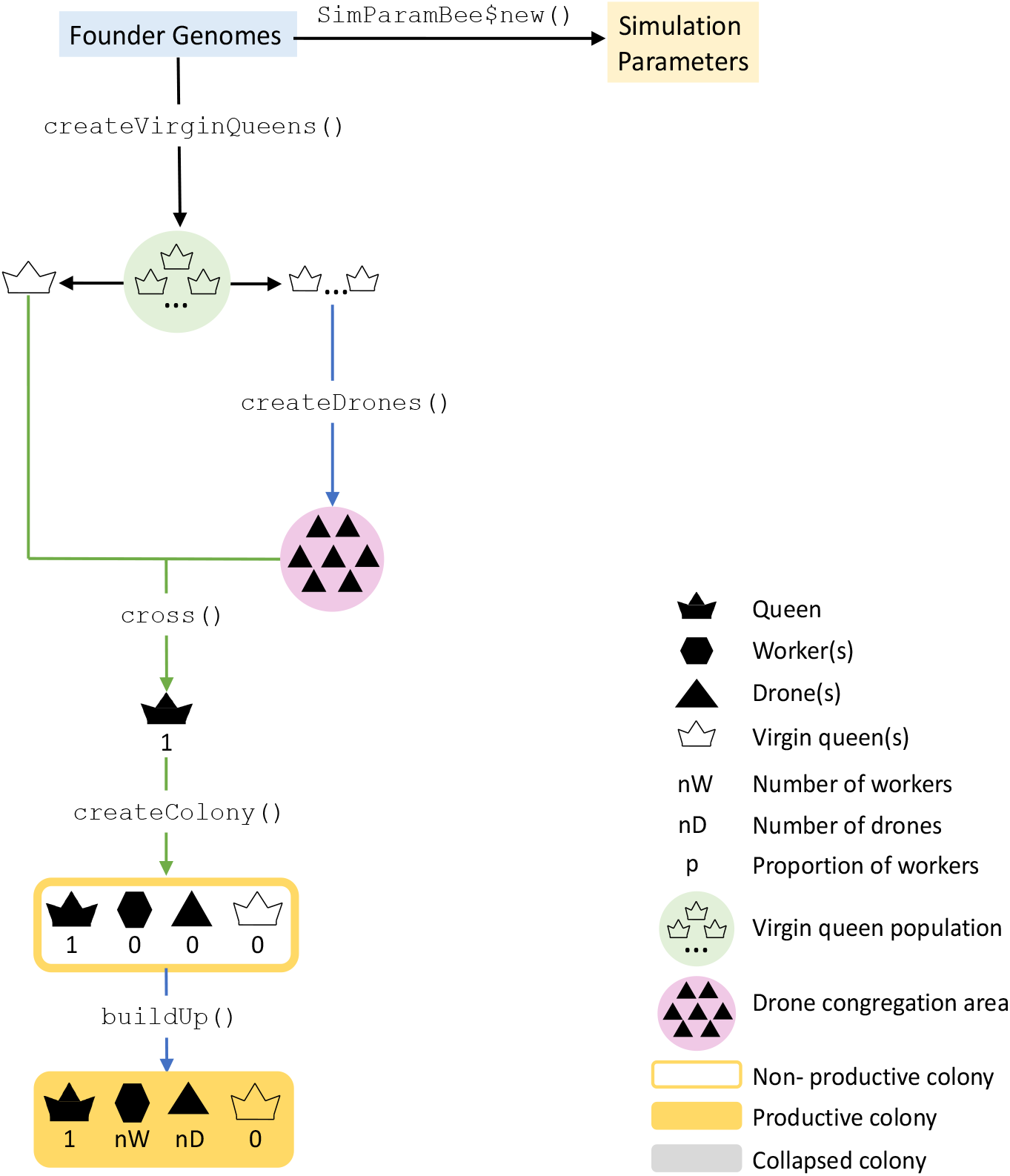
A flow chart of initialising a honeybee simulation in SIMplyBee. The first step is simulating a desired number of founder genomes and specifying the global simulation parameters in a new `SimParamBee` object. Next, we create the base virgin queens from the founder genomes. We can simulate any number (nInd) of virgin queens with the maximum being the number of simulated founder genomes. We choose one virgin queen as the future queen of the colony (left). On the other side (right), we select one virgin queen to provide drones for the DCA. We could select more virgin queens as future queens to create more colonies, and more virgin queen to contribute to the DCA. We next cross the virgin queen to a sample of drones from the DCA and use it to create a colony. We next build-up a colony, which adds in a desired number of workers and drones. The build-up also results in a productive colony.

To start the simulation we initially install the SIMplyBee package and load it.

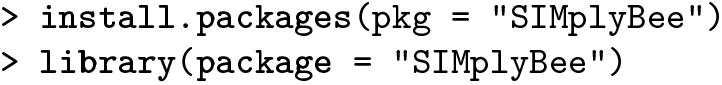

First, we generate founder honeybee genomes using `simulateHoneybeeGenome()`. Alternatively, we can import chromosome haplotypes, say from drones, phased queen or worker genotypes. Here, we simulate 10 *A. m. carnica* honeybees with only three chromosomes, each with only 100 segregating sites. These numbers are not realistic, but enable a fast demonstration. You can read more about initiating a simulation in the Additional File 1.

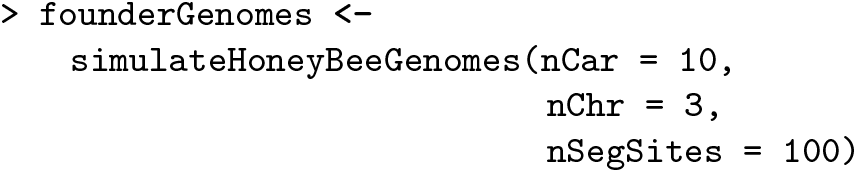

Second, we set global simulation parameters with `SimParamBee`. `SimParamBee` builds upon the AlphaSimR class `SimParam`, which contains global user-defined simulation parameters that apply to all individuals and populations, including genome and trait parameters, but also global pedigree and recombination events. `SimParamBee` additionally holds honeybee-specific information: default numbers of workers (`nWorkers`), drones (`nDrones`), and virgin queens (`nVirginQueens`) in a full-size colony, default number of drones that a queen mates with (`nFathers`), default proportions of workers that leave in a colony swarm (`swarmP`) or are removed in a colony split (`splitP`), and a default percentage of workers removed during colony downsize (`downsizeP`). These default numbers can be changed according to the needs of a simulation. The default numbers can even be replaced by providing functions to sample numbers. For example, to sample variable number of fathers, workers, drones, and virgin queens from Poisson or truncated Poisson distributions (Additional File 7). Most SIMplyBee functions that take the number of individuals as an argument can accept these sampling functions as input, meaning that the output of such function calls will be stochastic. `SimParamBee` also holds information about the *CSD* locus: the chromosome (`csdChr`), the physical position on the chromosome (`csdPos`), and the number of alleles (`nCsdAlleles`). `SimParamBee` also holds the caste of each individual in the simulation. The caste can be a queen, father (drones that successfully mated and died), worker, drone, or virgin queen. The caste can change during the life of a honeybee. For example, after successful mating, a virgin queen becomes a queen and drones become fathers.

Here, we show how to set the `SimParamBee` with the default number of workers in a colony being 100, default number of drones in a colony being 10, and the *CSD* locus to have 32 alleles. We save the output to the `SP` object, which enables its direct use for other SIMplyBee functions without explicitly passing it as an argument.

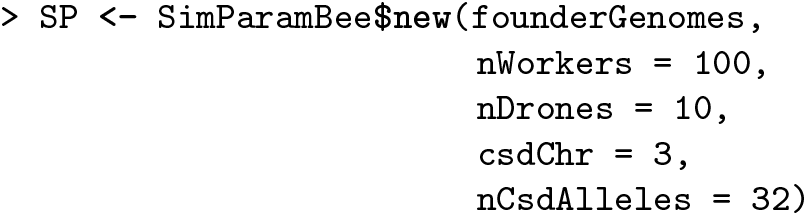

### Colony as an operational unit

Honeybees live in colonies organized into two castes, the queens and the workers, and the drones. The queen is a single reproductive diploid female, the workers are non-reproductive diploid females and perform various colony maintenance tasks (collect food, nurse larvae, clean cells, etc.), and the drones are reproductive haploid males. A single colony can contain up to 65,000 workers ([22]. Drones can represent up to 20% of a honeybee colony [23]. In SIMplyBee, we accounted for this social organisation by creating a class `Colony` that holds all the above mentioned individuals as AlphaSimR populations. For the ease of use, we refer to all these groups as “castes”. We also consider the drones that the queen mated with (fathers) and the virgin queens (queen-cells or emerged virgin queens) as “castes”, although the term is not biologically correct. The `Colony` further contains technical information about the colony, its identification (`id`) and location (`location`) coded as (`latitude, longitude`) coordinates. Further, it contains logical information about the past colony events: `split`, `swarm`, `supersedure`, or `collapse`. It also contains `production` status, which indicates whether we can collect a production phenotype from the colony. The latter is possible when the colony is built-up to its full size and has not swarmed. The production is turned off when a colony downsizes, collapses, swarms, or is a split from a colony.

Here we show how to create a colony in SIMplyBee (Figure 1). From the founder genomes, we create a base population of virgin queens (an AlphaSimR’s `Pop` class object). We check whether they are really virgin queens with the function `isVirginQueen()`. SIMplyBee contains a similar `is*()` function for checking the caste of each honeybee, where `*` is the inquired caste. These functions return `TRUE` or `FALSE` when an individual belongs or does not belong to the caste.

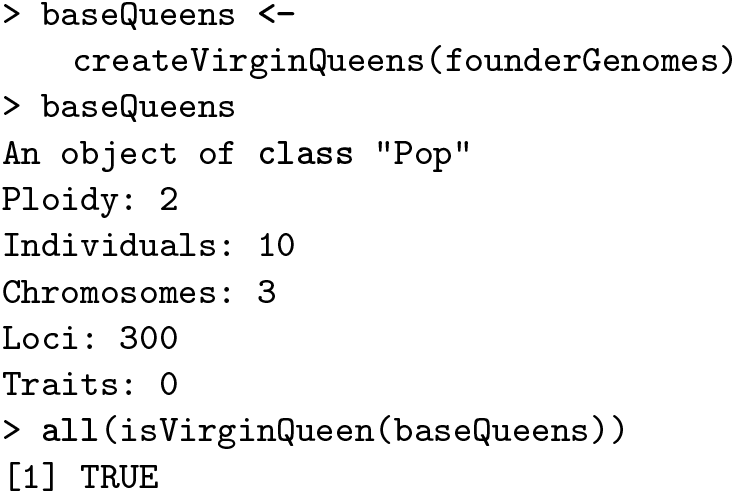

From the first virgin queen, we create a `Colony` object. We can also use `n` simulated queens to simulate `n` colonies. Printout of a `Colony` object returns its basic information: the `id` (1), location (not set, hence `NA`), queen (not yet available, hence `NA`), the number of fathers, workers, drones, and virgin queens, as well as colony event statuses.

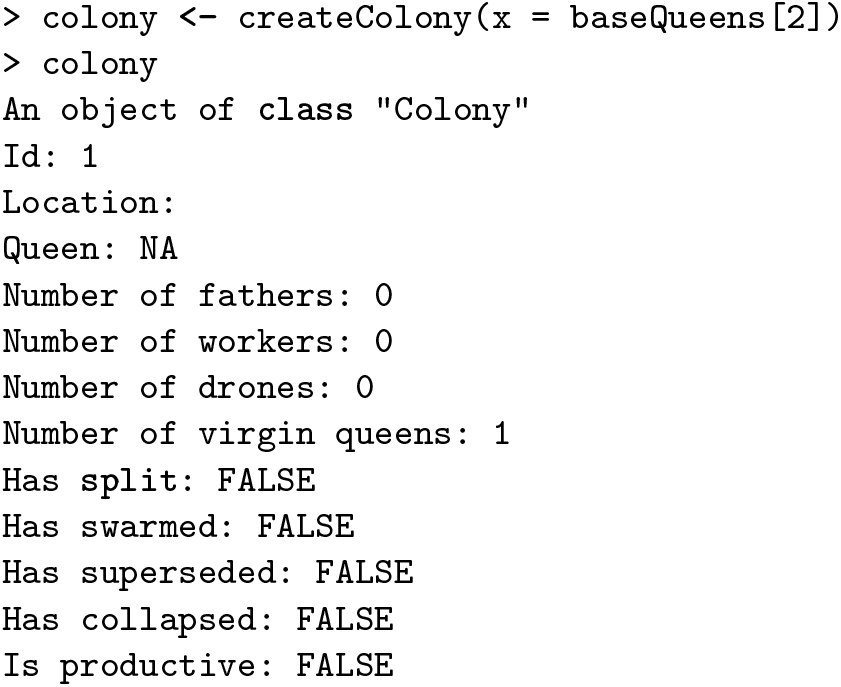

From the second virgin queen, we create drones for mating, the drone congregation area (DCA) (Figure 1). You can use more than one virgin queen to create a DCA. Technically, virgin queens do not create drones, only queens with colonies do that. However, to kick-start the simulation, we need drones. The function `createDrones()` can therefore work with a virgin queen or a mated queen in colony as the input to create drones. SIMplyBee contains a `create*()` function for each of the castes.

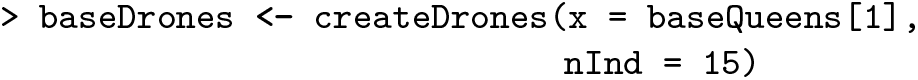

In honeybees, the virgin queens mate with multiple drones, a phenomena termed polyandry. A honeybee virgin queen will undergo several mating flights to a drone congregation area (DCA) that consists of thousands of drones from up to 240 colonies [24]. There, she will mate with 6 to 24 drones [25] and store the sperm in spermatheca for life.

Here, we now show how to mate our virgin queen in the colony with the function `cross()` to promote her to a queen so she can lay eggs for workers and drones. After the crossing, we inspect the colony printout and see that the identification of the queen is `“2”`, that there are 15 fathers, and that there are no more virgin queens in the colony.

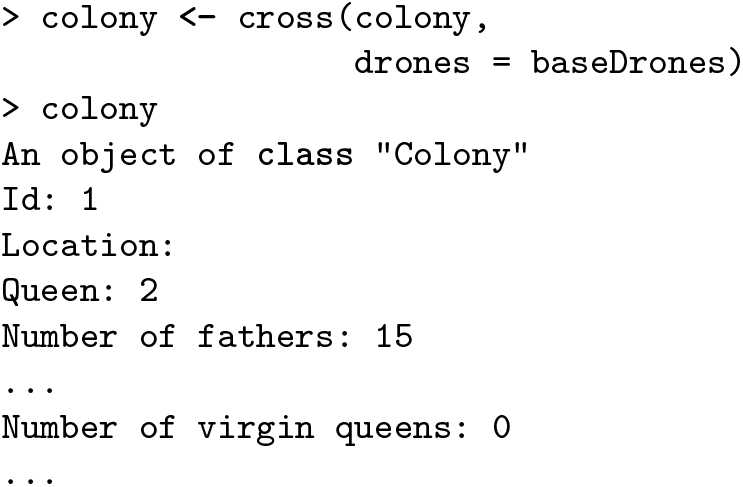

In SIMplyBee, we developed additional functionality regarding open or controlled mating that we describe in detail in the Crossing vignette in Additional File 4. There is a function to i) create a DCA for open mating, `createDCA()`, or a DCA with drones from sister queens, as commonly found on honeybee mating stations, `createMatingStationDCA()`; ii) sample a desired number of drones from a DCA, `pullDroneGroupsFromDCA()`; iii) create a cross plan, which includes information about which drones will mate with each virgin queen, `createRandomCross-Plan()`; iv) cross a virgin queen to a selected population of drones or according to a user defined cross plan, `cross()`.

Next, we show to to build-up the colony with workers and drones with the `buildUpColony()` function. We can specify the number of workers and drones. Without specifying these numbers, the function uses default numbers in the `SimParamBee` object. Building up the colony always switches the production status to `TRUE`.

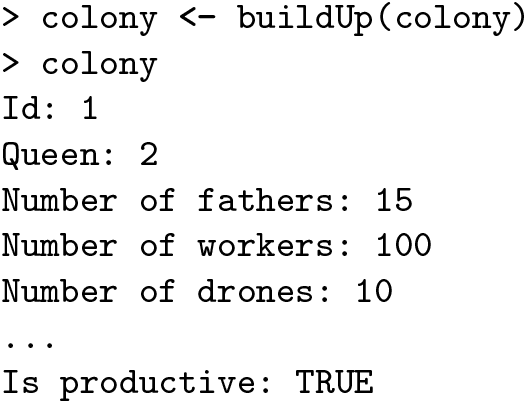

SIMplyBee also contains `n*()` functions to count individuals in each caste.

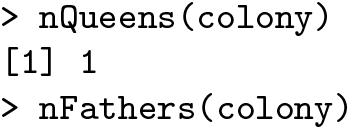

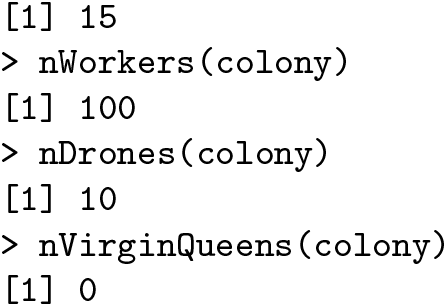

We can access individuals using `get*()` functions. Note that these functions copy individuals and hence leave individuals in the colony – check `pull*()` functions for “pulling” individuals out of the colony.

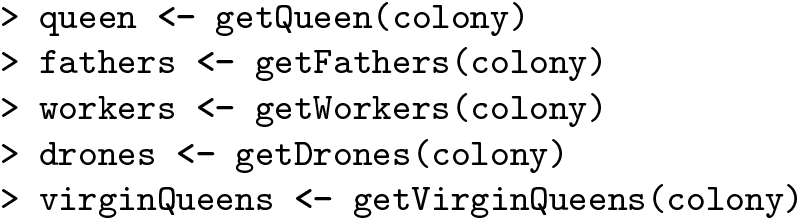

We can access the caste information of every individual with the *getCaste()* function.

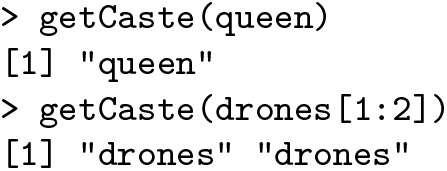

The `getCaste()` can be very useful when you have a group of honeybees and you do not know their source.

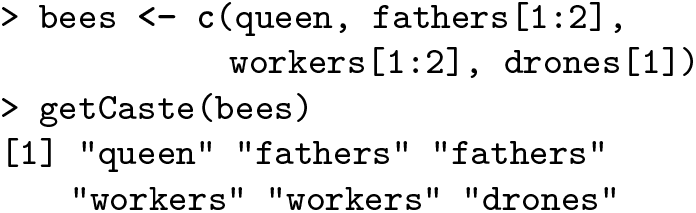

Additional functions for caste operations include obtaining the identifications of caste members, and set or get the year of birth and age of the queen. To work with castes within a `Colony` object, use `addCastePop()`, `replaceCastePop()`, or `removeCastePop()`, which all return a modified `Colony` object.

### Haplodiploidy and *CSD*

Honeybees belong to the insect order Hymenoptera. The order is characterised by haplo-diploid inheritance [26, 27, 28]. In SIMplyBee, we accounted for the haplo-diploidy by simulating queens and workers as proper diploids and males as doubled haploids. Doubled haploids are fully homozygous individuals. However, we only use one (haploid) genome set in all drone operations inside the functions.

Second, besides haplo-diploidy, sex in honeybees is determined by the complementary sex determination (*CSD*) locus. Fertilised eggs that are heterozygous at the *CSD* locus develop into diploid females, while homozygotes develop into non-viable diploid drones [29]. In SIMplyBee, we assign a specific region of the genome to represent the *CSD* gene. This region corresponds to the position of the *CSD* on chromosome three [30]. We simulate the *CSD* region as a sequence of non-recombining biallelic SNPs which determine a *CSD* allele. To account for balancing selection [31] at the *CSD* locus, we edit the initial founder genomes to achieve the desired number and frequency of *CSD* alleles in a population. The user can control the number of possible *CSD* alleles (2^length^) by controlling the length of the locus (in number of SNPs) in the simulated population.

We retrieve *CSD* alleles with `getCsdAlleles()`, which for diploids reports two non-recombining haplo-types as strings of 0s and 1s representing respectively ancestral and mutation alleles along the *CSD* locus. Here we show the queen’s *CSD* alleles. The first row of the output shows locus identifications (`chromosome_-locus`) and the first column shows haplotype identifications (`individual_haplotype`). We see that the two sequences are different, meaning that she is heterozygous, as expected – otherwise her egg would have developed into a diploid drone that would have been killed by workers.

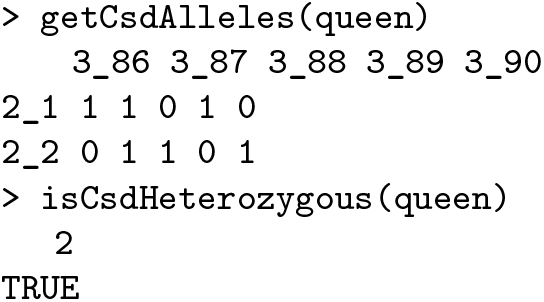

The *CSD* heterozygosity of honeybees is critical. Comparing *CSD* alleles of this queen and the drones she mated with (compare `getCsdAlleles(queen)` and `getCsdAlleles(fathers)` – not shown) shows no allele matches, which means we do not expect any homozygous brood in this colony. We obtain the theoretical brood homozygosity of a queen with `pHom-Brood()` and realized number of homozygous offspring with `nHomBrood()`.

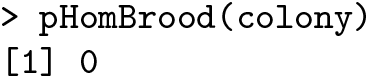

Let’s create an inbred colony, by mating a virgin queen from our colony with her brothers, and inspect the expected brood homozygosity.

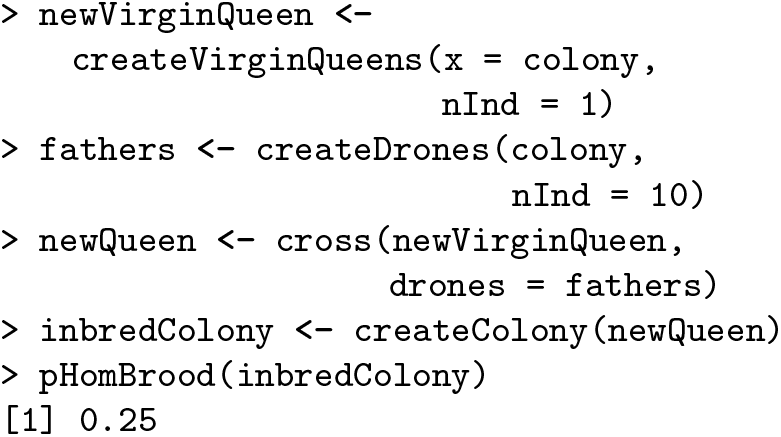

We see that the expected brood homozygosity is 0.25, hence we expect that about 25% of diploid brood will be homozygous. We now add workers to the colony to observe how many of them are homozygous. Inheritance is a random process, so a realised number of homozygotes will deviate from the expected proportion.

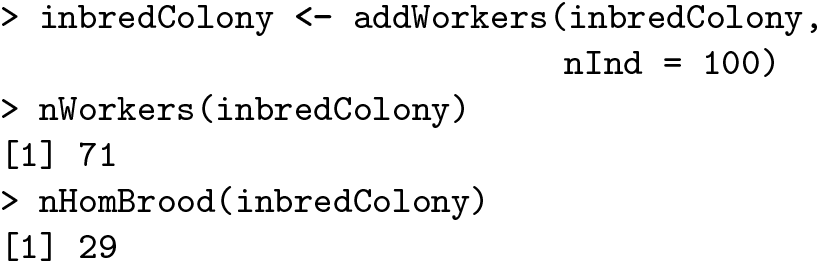

We tried adding 100 workers, but we only got 71. The difference of 29 is due to non-viable `CSD` homozygous brood. The information about the number of homozygous brood is stored in the queen’s miscellaneous slot and is updated every time we create offspring from her.

### Colony events

A honeybee colony can experience a series of events during its life: swarming, superseeding, splitting, and collapsing. We present the details of colony events and their simulation in the Additional File 3.

In swarming a proportion of workers leave the hive with the queen, while the rest of the workers and drones stay in the hive. New virgin queens emerge and compete in the colony. The winner undergoes mating flights as described above. In SIMplyBee, we created a function `swarm()` to swarm a colony (Figure 2). An input parameter to the function is also the percentage of workers that leaves with the swarm, `p`. This can be either a fixed number or a function that samples the `p` from either a uniform distribution or in some cases also from a beta distribution that accounts for the number of individuals in a colony (colony strength). You can read more about the sampling functions in the Additional File 7. The `swarm` function returns an R list with two colonies: `swarm` that contains the old queen and a proportion `p` of workers, and `remnant` that contains the rest of workers, all the drones, and virgin queens that are daughters of the queen that swarmed. The function also changes the `swarm` status to `TRUE` and `production` status to `FALSE`

**Figure 2.**
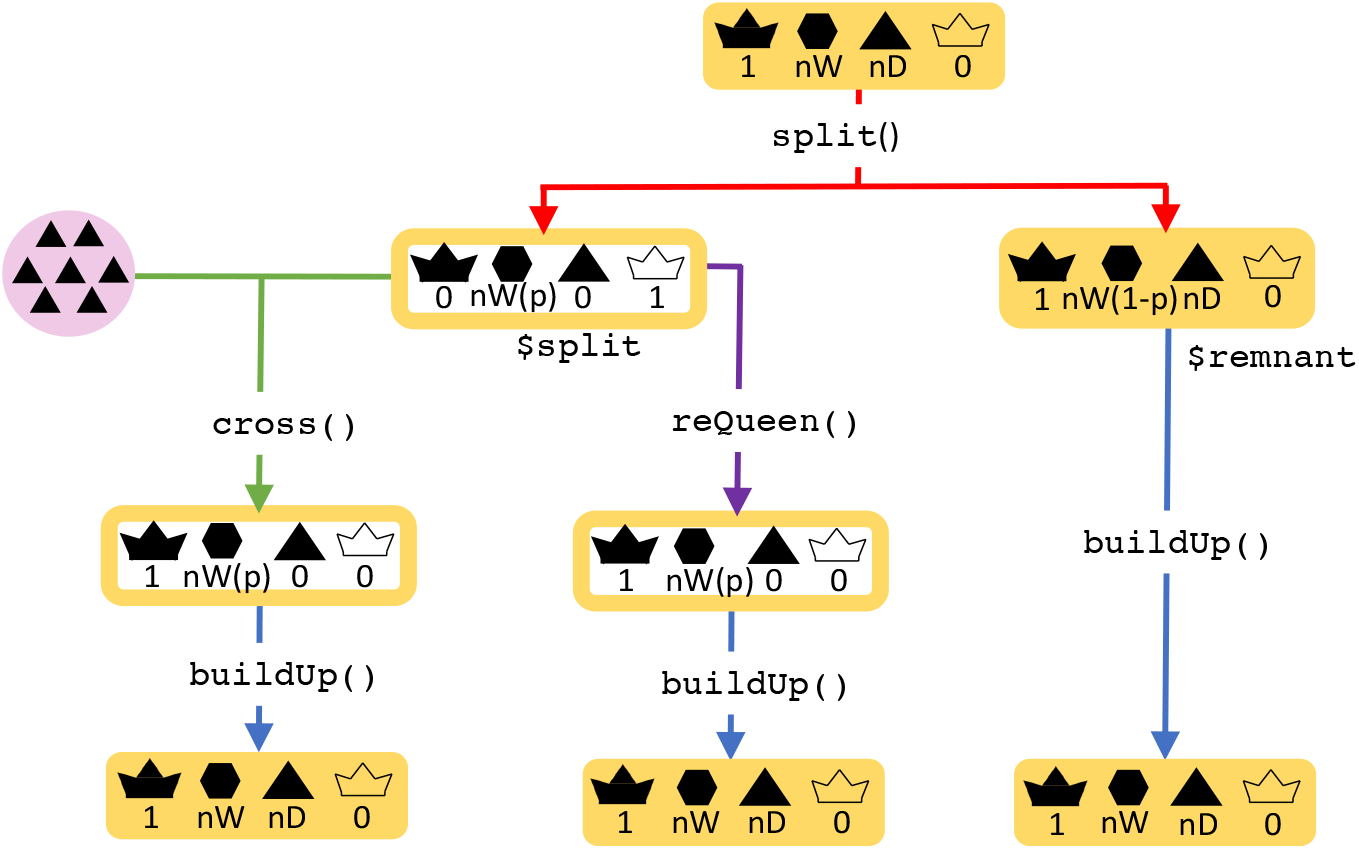
A flow chart of the colony swarming event in SIMplyBee. The `swarm()` function returns an R list with two colonies, `swarm` and `remnant`, both of which are non-productive. Parameter `p` represent the proportion of workers that leave with the swarm. After the swarm, the user can cross() the virgin queen of the remnant colony, or use an already mated queen from another source using reQueen(), which mimics the beekeepers’ options. Refer to the key in Figure 1.

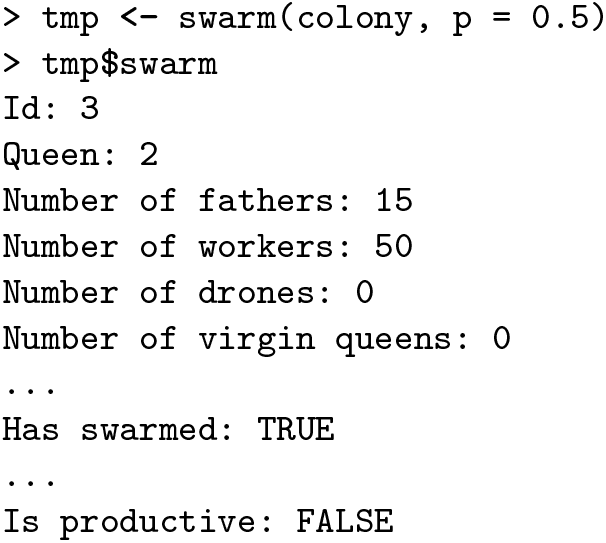

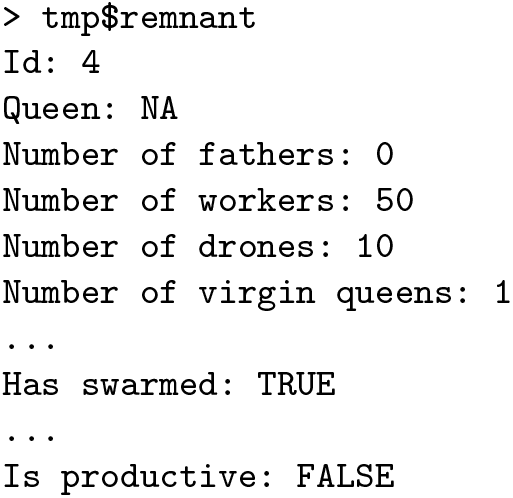

To prevent swarming of strong colonies, beekeepers often split them by taking away a proportion of workers and starting a new colony with a new queen. The rest of the workers stay in the hive with the old queen. In SIMplyBee, we created a function split() that takes a colony and proportion of workers that we remove with the split, `p`. The `split` function returns an R list with two colonies: `split` that contains proportion `p` of workers taken from the main hive and virgin queens, and `remnant` that contains the queen, the remaining workers and drones (Figure 3). After the split, the remnant colony is still productive, while the split is not.

**Figure 3.**
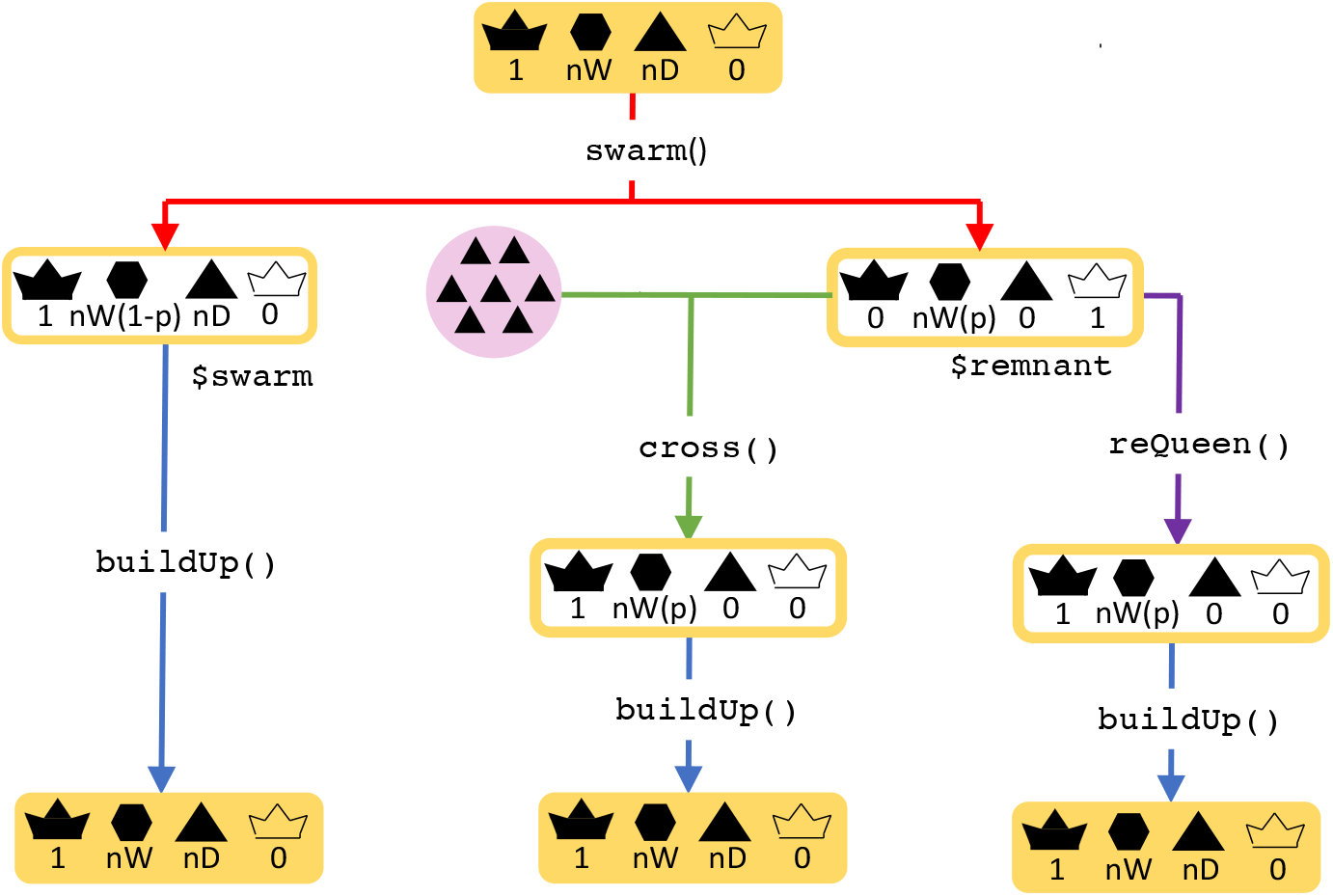
A flow chart of the colony splitting event in SIMplyBee. The `split()` function returns an R list with two colonies, `split` and `remnant`, where the *split* is non-productive and the *remnant* is productive. Parameter p represents the proportion of workers that are removed in a split. After the split, the user can cross() the virgin queen of the remnant colony, or use an already mated queen from another source using reQueen(), which mimics the beekeepers’ options. Refer to the key in Figure 1.

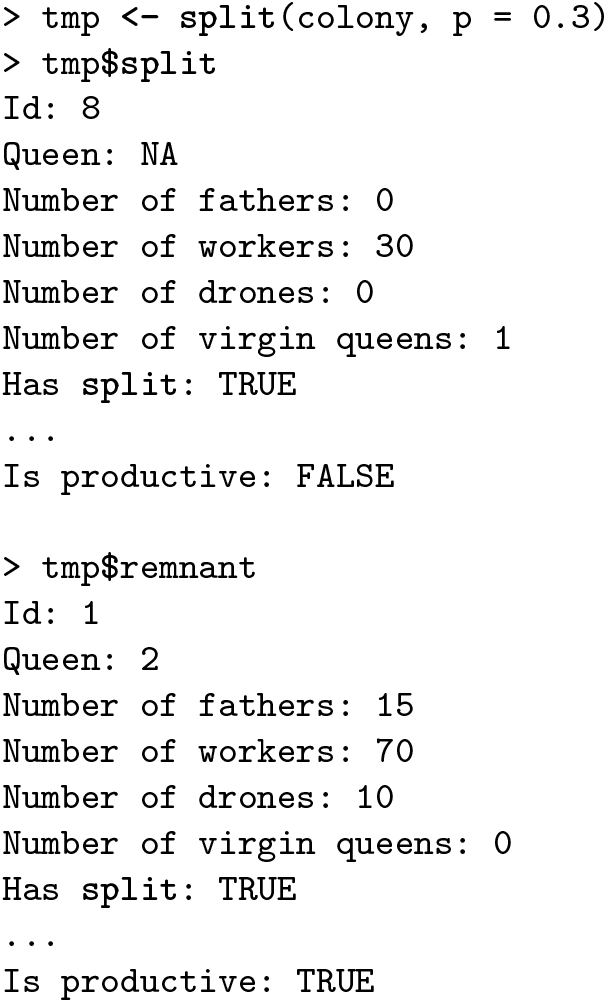

In supersedure, the queen dies or is killed and its workers raise new virgin queens. In SIMplyBee, we created a function `supersede()` that removes the queen and produces new virgin queens from the brood (Figure 4). After a supersedure, the colony is still productive because the workers are still present and working within the colony.

**Figure 4.**
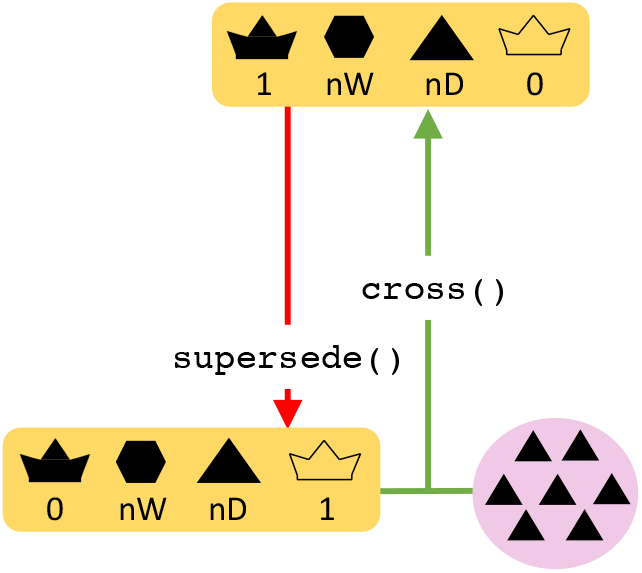
A flow chart of the colony supersedure event in SIMplyBee. The `supersede()` function returns a queen-less colony with a virgin queen. After a supersedure, a colony remains productive since the colony is still at its full size but a cross() is required for a new queen. Refer to the key in Figure 1.

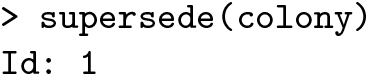

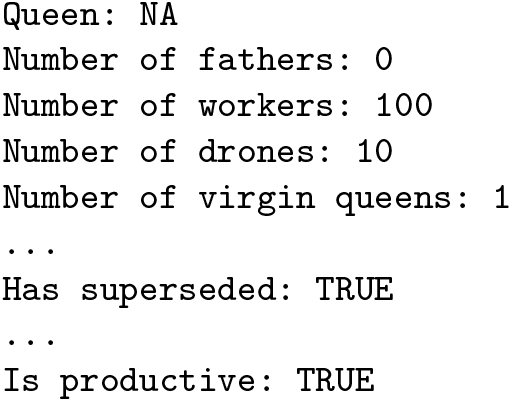

Finally, some colonies can collapse due to the death of all its members. In SIMplyBee, we created a function `collapse()` that collapses a colony by changing the `collapse` status to `TRUE` (Figure 5). The function keeps the individuals in the colony to enable study of genetic and environmental causes contributing to the collapse. In reality, dead honeybees would also be present in a collapsed colony.

**Figure 5.**
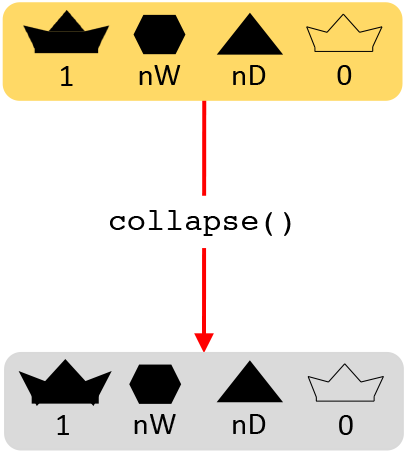
A flow chart of the colony collapse event in SIMplyBee. The collapse function keeps all the individuals in a colony, but turns on the collapse parameter, hence marking the colony as collapsed and all the individual within it as dead. Further simulation with a collapsed colony is not allowed in SIMplyBee. Refer to the key in Figure 1

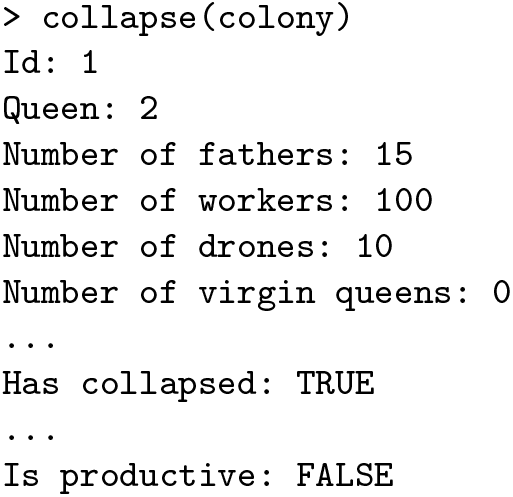

SIMplyBee also includes functions to build-up a colony, as shown above, a function to downsize a colony named `downsize()`, and a function to combine a strong and a weak colony named `combine()`. You can read more about simulating events in SIMplyBee in Additional File 3.

### Working with multiple colonies

Beekeepers regularly work with a collection of colonies at once. To this end, we created a `MultiColony` class that collects a list of colonies to represent an apiary, a region, an age group, etc. You can read more about working with multiple colonies in the Additional File 2.

In SIMplyBee, we use a function `createMulti-Colony()` to create a `MultiColony` object. Here, we take three of the remaining base population virgin queens to create an apiary with three virgin colonies. The printout of the object returns basic information including the number of all, empty, and NULL colonies, and information about the colony events for the colonies within.

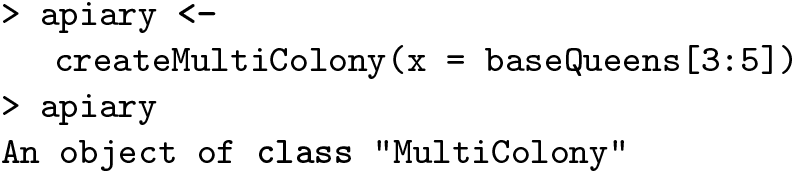

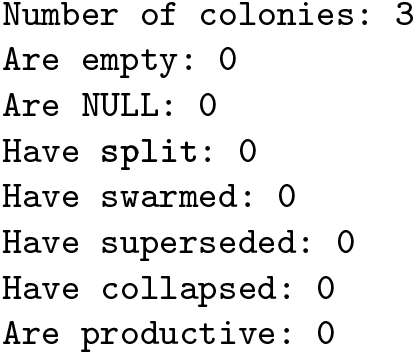

All the functions for managing a `Colony` object can also be applied to a `MultiColony` object, which streamlines simulation scripts. These functions include functions for crossing, adding or removing individuals, simulating Colony events, etc. Here, we demonstrate a short script to cross the `apiary`, build-up its colonies, and swarm some of them. We start by creating a DCA from the remaining base population virgin queens and sampling three groups of drones to mate the three virgin queens. As already mentioned, when we sample individuals in SIMplyBee, we can use either fixed numbers or use sampling functions (Additional File 7).

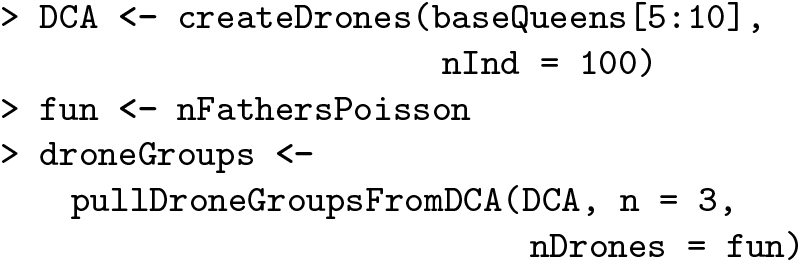

Next, we cross the virgin queens in the apiary with the provided drones groups. We test for the presence of queens before and after mating to show that mating was successful. For this we can use `is*Present()` functions, where `*` is the caste, that check the presence of a caste in a colony. We then build-up all the colonies.

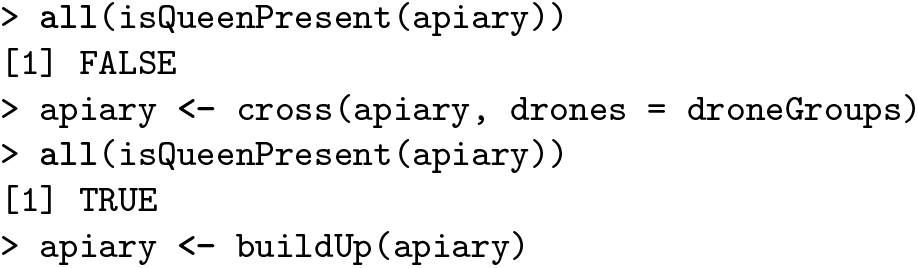

Now, we use the function `pullColonies()` to sample one colony that will swarm. This returns an R list with two `MultiColony` objects: `pulled` with the sampled colonies, and `remnant` with the remaining ones. We save the latter back in the apiary.

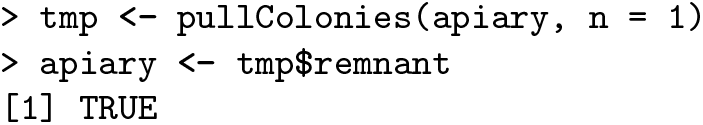

Now, we swarm the pulled colonies. When applied to `MultiColony`, all the colonies are swarmed with the same `p`, unless specified otherwise. You can read more about simulating colony events for `MultiColony` in the Additional File 3. The swarm function returns an R list with two `MultiColony` objects, `remnant` and `swarm`.

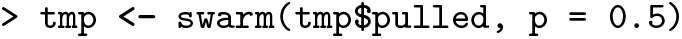

Assuming that we did not catch the swarm(s), we combine the colonies that did not swarm with the swarm remnant(s) into an updated apiary.

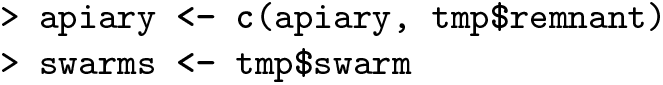

### Genomics and Quantitative genetics

Similar to extracting the *CSD* genomic sequence, SIMplyBee provide functions to extract whole-genome information for any set of individuals using `get*Haplo()` or `get*Geno()`. Here, `*` can be `SegSite` to extract all segregating/polymorphic loci tracked in the simulation, `Snp` to extract marker loci, or `Qtl` to extract Quantitative Trait Loci. There is also `getIbdHaplo()` to extract Identity By Descent information, with IBD alleles defined as those originating from the base population genomes. These functions leverage AlphaSimR functionality, but work with SIMplyBee `Colony` or `MultiColony` objects and in addition take the caste argument to extract information only for a specific `caste`. For example, to extract genotypes of the first five workers in the colony at the first five tracked segregating sites we call the code below. See further details in the Additional File 5.

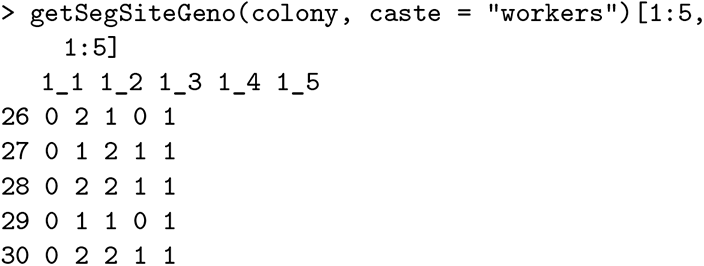

Honeybee phenotypes are characterized with two important phenomena. First, in honeybee keeping and breeding, phenotypes are mostly collected on a colony level as opposed to on an individual level. Second, phenotypes in honeybees are a complex interaction between queen and worker effects that are often negatively correlated [32, 33]. For most traits, the queen indirectly contributes to the colony phenotype by laying eggs [34, 35] and stimulating the workers through pheromones [36, 37], while workers contribute directly by doing the actual work.

In SIMplyBee, we simulate genetic and phenotypic values for each individual honeybee and enable also calculating colony-level values. Quantitative genetic simulation is initiated in the `SimParamBee` by specifying the assumptions about the genetic architecture of traits in `SimParamBee`, including the number of quantitative trait loci, distribution of their effects, as well as genetic and environmental variances and covariances. Let’s initiate another simulation and specify two negatively correlated traits that will represent the queen and worker effect for honey yield. You can read a more extensive explanation of this simulation in the Additional file 6.

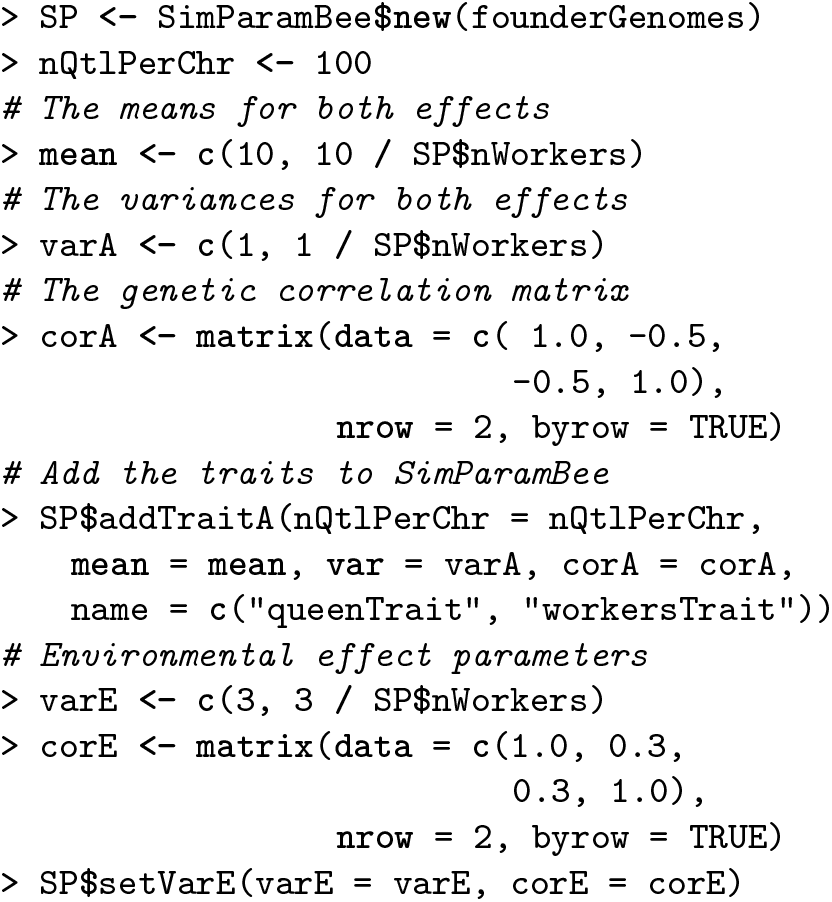

This initiation triggers calculation of individual-level genetic and phenotypic values. Using an AlphaSimR `Pop` class object, genetic and phenotypic values are stored respectively in `gv` and `pheno` slots. We can access the genetic and phenotypic values of population or colony members with functions `getGv()` and `get-Pheno()`, both of which have the `caste` argument and work on `Colony` and `MultiColony`.

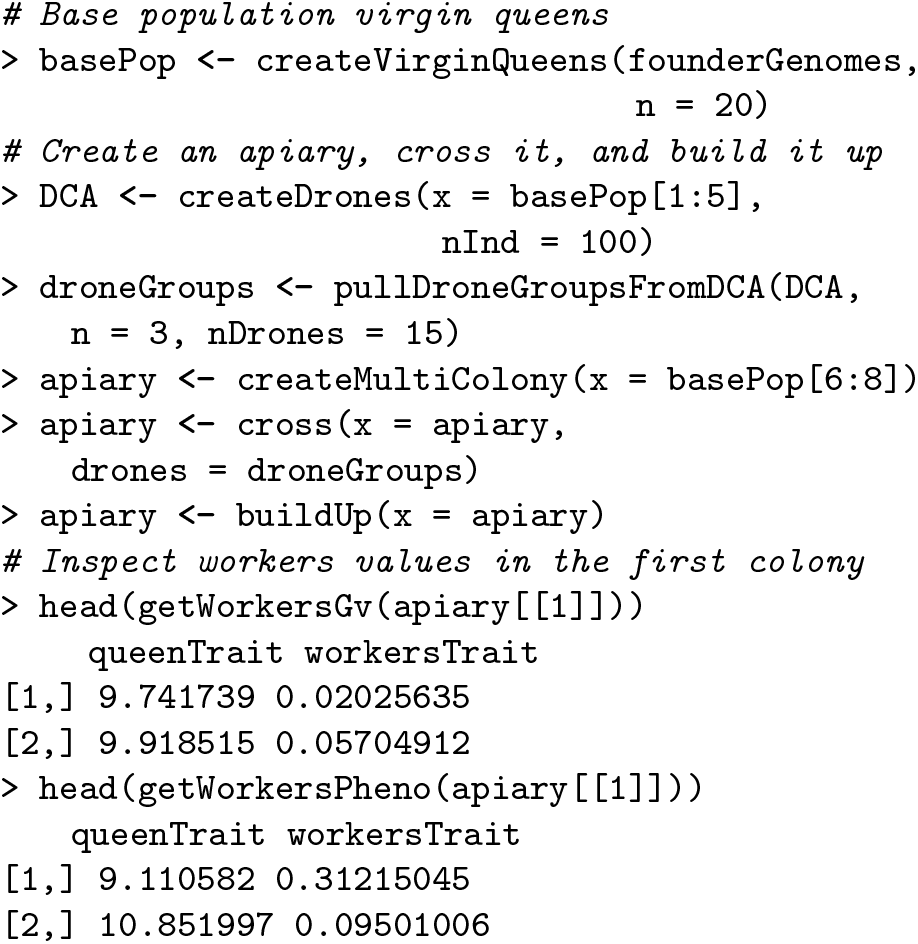

To obtain colony-level values we provide `calc-ColonyValue()` function that maps individual values to colony values. While we provide an established mapping function from the literature [38, 14, 39], users can provide their own mapping function. Examples of such quantitative genetic simulations of one or multiple correlated traits are shown in the Additional File 6.

Here, we compute the colony-level genetic and phenotypic values for the colonies in our apiary.

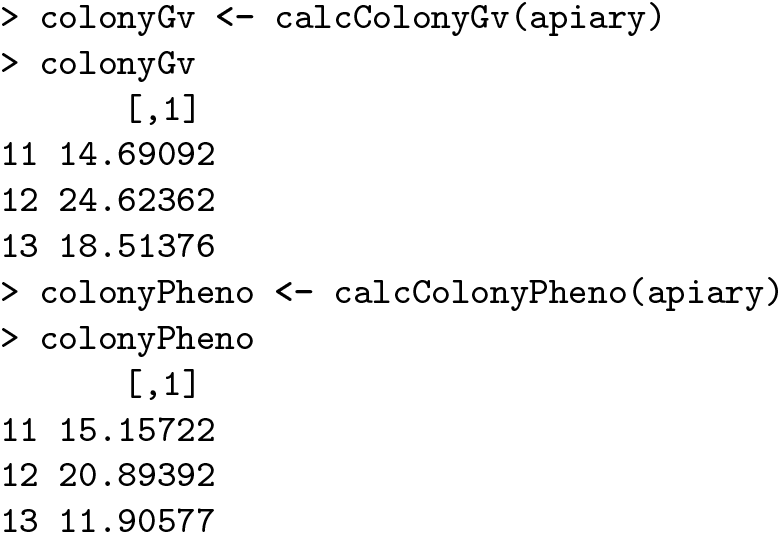

We see that the best colony according to the genetic as well as the phenotypic value is colony with ID “12”, hence we would select it for further reproduction. We can pour the values into the `use` parameter of the `selectColonies()`.

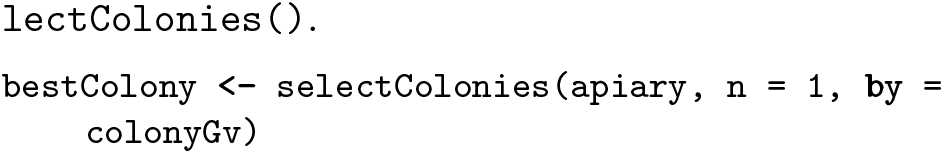

## Discussion

SIMplyBee is an R package for holistic simulation of honeybee breeding and conservation programs. In comparison to previously developed general genetics and breeding simulators [15, 16], it enables the simulation of honeybee-specific genomes, social organisation, and behaviours. SIMplyBee differs from previously developed honeybee-specific simulators [14, 17] by enabling the simulation of individual honeybees, individual-level and colony-level quantitative values, and colony events that can affect genetic and phenotypic variation in a population.

SIMplyBee provides a valuable research platform for testing different population-management decisions and answering various questions regarding design of breeding schemes. SIMplyBee can be used to test the effect of various decisions in a breeding program on genetic gain, genetic diversity and inbreeding; or to test the accuracy of inferences with competing quantitative genetic models. For example, users can test the effect of different phenotyping schemes by varying the frequency of phenotyping or the measuring scale. Furthermore, SIMplyBee can be used to test different mating control designs and the effect of varying the number of sires or drone producing queens on a mating station. The users can also test different selection strategies by varying the time of selection, the number of selected queens, or the sources of information (pedigree, genomic, and phenotypic data). The list of such potential studies is long. SIMplyBee is also a valuable platform for answering questions regarding the conservation of honeybees. Users might be interested in the effect of mating and management decisions on the genetic diversity in a population along the whole genome or only at the *CSD* locus. Users might also be interested comparing how different migration or import practices and associated policies affect genetic diversity, or how to design a conservation program to preserve the existing genetic diversity.

While developing SIMplyBee, we observed that there is very limited knowledge of relationship between individual-level and colony-level phenotypes. This knowledge includes the contribution of individual worker’s honey yield to the total colony honey yield, the effect of the number of workers on the total colony honey yield, the effect of colony events on colony honey yield, etc. While this knowledge gap is understandable, given the sheer number of honeybees in a colony, future advances in sensor and beekeeping technologies and data science (machine learning) will provide evermore fine-grained data. Such data could further contribute to explaining the relationship between individual-level and colony-level phenotypes. SIMplyBee can serve as a research platform to model such relationships. Less ambitiously, SIMplyBee can serve as a research platform to test assumptions of the current quantitative genetic models [39, 14, 38]. Another challenge that we faced in developing SIMplyBee was in providing functionality to calculate statistical genetic values, that is, breeding values, dominance deviations, and epistasis deviations. Since SIMplyBee leverages AlphaSimR [15], all these values can be calculated using `bv()`, `dd()`, and `aa()` functions. However, caution is required because these values are computed “relative” to the population of individuals at hand, which means that for a honeybee simulation we would either have to report these values relative to each colony population, which would make the output useless, or we would have to create a large “meta” population object of all currently living honeybees. Also, these functions and the underlying theory assume Hardy-Weinberg equilibrium [40]. Further development is required to address this aspect in SIMplyBee.

Future development of the SIMplyBee package will focus on additional features and improving the functionality and efficiency of existing features. Our immediate focus is on the following three features. First, we are working on a new honeybee demographic model to include more subspecies and improve estimation of the model parameters ([41]). While we are currently using MaCS [20]) to simulate the genome, we are also contributing honeybee species and demography to the stdpopsim consortium [42, 43], which uses the msprime (backward in time) [44, 45] and SLiM (forward in time) [46] simulators. Second, we will further optimize the speed and memory performance of SIMplyBee. A simulation of an entire real-size honeybee colony or a breeding program with such colonies can be computationally demanding because a single colony can hold up to several tens of thousands of workers. We are profiling memory and compute bottlenecks in SIMplyBee and will optimize functions by leveraging C++ via the Rcpp package [47] or by adopting alternative approaches for achieving the same target. For example, by working with the expectation and variance of genetic values in progeny [48, 49] instead of simulating tens of thousands of workers. Third, we will add spatial awareness. Colony location plays a major role in honeybee mating and colony performance. The current implementation enables setting the location of every `Colony` and `MultiColony` object. We will develop functionality to create a DCA or sample the drones for a virgin queen mating according to the location of colonies, for example, in a certain radius, since virgin queens are more likely to mate with drones from nearby colonies. Furthermore, we will add spatially-aware simulation of environmental effects. Honeybee colony performance depends heavily on the environment due to food provision, weather, pests, etc. Such environmental conditions usually change continuously through space, hence colonies closer together usually experience more similar environmental conditions than colonies further apart. The framework for such spatially aware simulation and modelling has already been developed and tested in a livestock setting [50].

We welcome the honeybee genetics and breeding community to join us in the future development of SIMplyBee. The development is hosted on GitHub at https://github.com/HighlanderLab/SIMplyBee. We welcome users and developers to fork this git repository and provide “pull request (PR)” contributions. Each pull request is reviewed by one of the developers within the core team. Based on the review, pull requests are updated before being merged into the development branch. The development branch is periodically merged into the main branch for publication on CRAN and user installation. For each function we request documentation with examples and unit tests to ensure future changes will not break the functionality.

In this work, we described the usage of SIMplyBee for simulating honeybee populations. However, other bee species share a similar organisation and behaviour as the honeybee. Hence, SIMplyBee could also be used to simulate other *Apis* species. For example, *Apis florea*, the dwarf honeybee, and *Apis cerana*. *Apis florea* importantly contributes to pollination in some countries of the Middle East and Asia. Its range is predicted to increase due to climate change [51] and SIMplyBee could be used to model a breeding program in this bee species as well.

## Conclusions

We developed a stochastic simulator, SIMplyBee, for holistic simulation of honeybee populations and population management programs. SIMplyBee builds upon its predecessors by simulating genomes of individual honeybees and the corresponding individual-level quantitative values. SIMplyBee stores individual honeybees as caste population within a colony object, which enables the simulation of colony events and calculation of colony-level values. Colonies can be further organised into multi-colony objects for ease of use. SIMplyBee provides a valuable research platform for honeybee genetics, breeding, and conservation. Possible uses include testing the effects of breeding or conservation decisions on genetic gain and genetic variability in honeybee populations, testing the performance of existing and novel statistical methods, etc. Future directions include improvements to the simulation of honeybee chromosomes through new demographic models, addition of spatial awareness in mating and phenotype simulation, reducing computational bottlenecks, and encouraging community engagement. We welcome the honeybee genetics and breeding community to collaborate with us in improving SIMplyBee.

## Availability and requirements

**Project name**: SIMplyBee

**Home page**: http://SIMplyBee.info

**Installation**: http://cran.r-project.org/package=SIMplyBee

**Development**: https://github.com/HighlanderLab/SIMplyBee

**Operating systems**: Windows, Linux, and MacOS

**Programming language**: R

**License**: MIT + file

## Ethics approval and consent to participate

Not applicable.

## Consent for publication

Not applicable.

## Availability of data and materials

The data and material for this study are available in the SIMplyBee GitHub repository https://github.com/HighlanderLab/SIMplyBee and https://SIMplyBee.info.

## Competing interests

Not applicable.

## Funding

JO acknowledges support from the ARRS Research program P4-0133. JO, LS, JB, JP, and GG acknowledge support from the ARRS research project L4-2624. JB acknowledges support from the ARRS PhD studentship 1000-20-0401. JB and JP acknowledge the support from the ARRS Research program P4-0431. LS and GG acknowledge support from the BBSRC DTP (EASTBio) CASE PhD studentship with AbacusBio and the BBSRC ISP grant BBS/E/D/30002275 to The Roslin Institute. For the purpose of open access, the authors have applied a Creative Commons Attribution (CC BY) license to any Author Accepted Manuscript version arising from this submission.

## Authors’ contributions

JO and GG initiated the project, planned the SIMplyBee implementation, and led the SIMplyBee development. LS, JB, and JP contributed to SIMplyBee development, documentation, and testing. JO wrote the first draft of this manuscript. All authors have contributed to the final version of the manuscript.

## Acknowledgements

The authors would like to thank R. Chris Gaynor for suggestions on how to leverage AlphaSimR functionality to implement honeybee specificities in SIMplyBee, and Philip Greenspoon for suggestions on improving the manuscript.

## Additional Files

Additional file 1 — Honey biology vignette

This vignette introduces SIMplyBee package by describing and demonstrating how SIMplyBee implements honeybee biology. Specifically, it describes how to initiate simulation with founder genomes and simulation parameters, how to create and build-up a colony, the colony structure, and complementary sex determining (*CSD*) locus. This vignette can also be found on https://www.github.com/HighlanderLab/SIMplyBee and https:\\SIMplyBee.info.

Additional file 2 — Multiple colonies vignette

This vignette introduces working with multiple colonies by demonstrating how to create and work with `MultiColony` objects in SIMplyBee. This vignette can also be found on https://www.github.com/HighlanderLab/SIMplyBee and https:\\SIMplyBee.info.

Additional file 3 — Colony events vignette

This vignette introduces the colony events and how to simulate them in SIMplyBee. It shows how to simulate swarming, splitting, superseding, and collapsing either a `Colony` or `MultiColony` objects. This vignette can also be found on https://www.github.com/HighlanderLab/SIMplyBee and https:\\SIMplyBee.info.

Additional file 4 — Crossing vignette

This vignette demonstrated how to cross virgin queens in SIMplyBee. It demonstrates how to cross a single or multiple virgin queens, cross either with pre-selected population/group of drones or according to a cross plan, and cross queens on an open DCA or mating station. This vignette can also be found on https://www.github.com/HighlanderLab/SIMplyBee and https:\\SIMplyBee.info.

Additional file 5 — Genomics vignette

This vignette demonstrates how to obtain genomic information of simulated honeybees. It also demonstrates, how to compute honeybee genomic relationship matrices in SIMplyBee. This vignette can also be found on https://www.github.com/HighlanderLab/SIMplyBee and https:\\SIMplyBee.info.

Additional file 6 — Quantitative genetics vignette

This vignette describes and demonstrates how SIMplyBee implements quantitative genetics principles for honeybees. Specifically, it describes three different examples where we simulate a single colony trait, two colony traits, and two colony traits where one trait impacts the other one via the number of workers. This vignette can also be found on https://www.github.com/HighlanderLab/SIMplyBee and https:\\SIMplyBee.info.

Additional file 7 — Sampling functions vignette

This vignette introduces sampling functions that sample either the number of caste individuals or proportion of workers that stay or are removed in colony events. This vignette can also be found on https://www.github.com/HighlanderLab/SIMplyBee and https:\\SIMplyBee.info.

## Additional file 1 – Honeybee biology vignette

### Introduction

This vignette describes and demonstrates how SIMplyBee implements honeybee biology. Specifically, it describes:

1. initiating simulation with founder genomes and simulation parameters,
2. creating and building up a colony,
3. colony structure, and
4. complementary sex determining (*CSD*) locus.

First, you need to install the package with `install.packages(pkg = “SIMplyBee”)`.

Now load the package and dive in! You load the package by running:

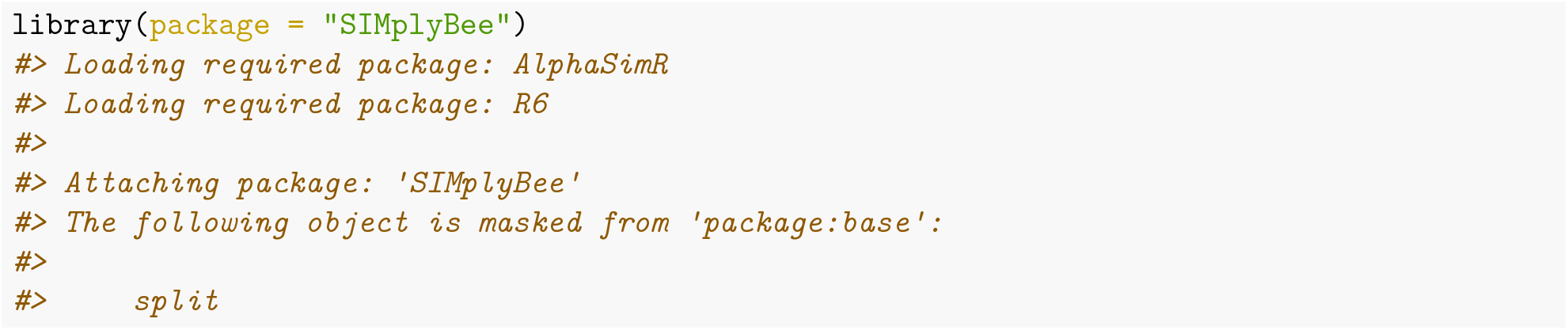

### Initiating simulation with founder genomes and global parameters

Figure 1 visualizes the initiation of the simulation. First, we simulate some honeybee genomes that represent the founder population. You can quickly generate random genomes using AlphaSimR’s `quickHaplo()`. These founder genomes are rapidly simulated by sampling 0s and 1s, and do not include any species-specific demographic history. This is equivalent to all loci having allele frequency 0.5 and being in linkage equilibrium. We use this approach only for demonstrations and testing.

**Figure 1:**
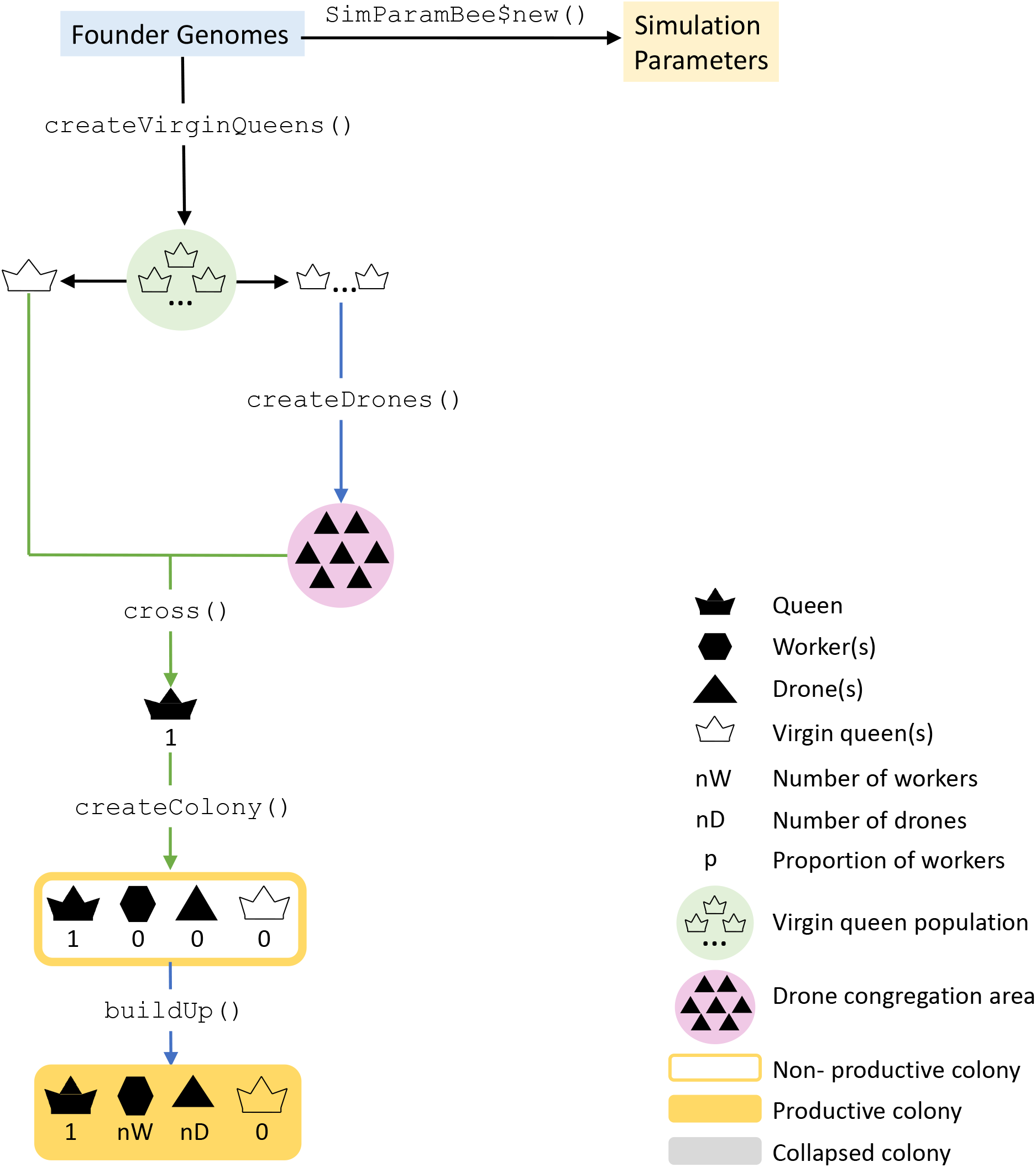
Simulation initiation.

Alternatively, you can more accurately simulate honeybee genomes with SIMplyBee’s `simulateHoneybeeGenom()`. This function simulates the honeybee genome using MaCS (Chen et al., 2009) for three subspecies: *A. m. ligustica*, *A. m. carnica*, and *A. m. mellifera* according to the demographic model described by Wallberg et al. (2014).

As a first demonstration, we will use `quickHaplo()` and simulate genomes of two founding individuals. In this example, the genomes will be represented by only three chromosomes and 1,000 segregating sites per chromosome. Honeybees have 16 chromosomes and far more segregating sites per chromosome, but we want a quick simulation here.

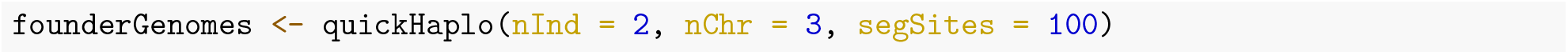

Alternatively, we use `simulateHoneybeeGenomes()` to sample genomes of a founder population including 4 *A. m. mellifera* (North) individuals and 2 *A. m. carnica* individuals. The genomes will be represented by only three chromosomes and 5 segregating sites per chromosome. These numbers are of course extremely low because we want a quick examample for demonstrative reasons. This chunk of code should take a few minutes to run.

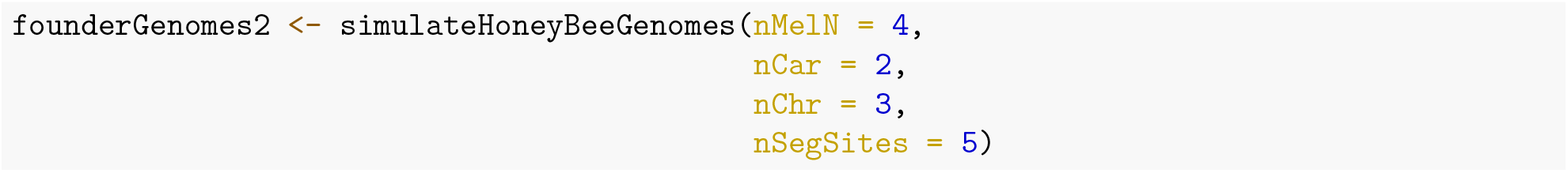

Unfortunately, due to the complexity of this function, even using such small numbers takes a while to run. Simulating a group of founder genomes with more realistic numbers will therefore require a lot of time to run. We suggest running this part to an external server and save the outcome as an RData file, which we can load in our environment and work with it.

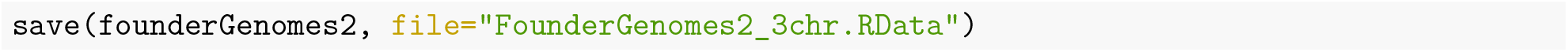

Besides specifying the number of individuals, chromosomes, and segregating sites, `simulateHoneybeeGenomes()`, also takes a number of genomic parameters: effective population size, ploidy, length of chromosomes in base pairs, genetic length of a chromosome in Morgans, mutation rate, recombination rate, and time of population splits. The default values for these numbers follow published references (Wallberg et al., 2014). While you can change these parameters, we don’t advise doing this because such demographic models, and their parameters, are estimated jointly, so we should not be changing them independently. You can read more about these parameters in the help page:

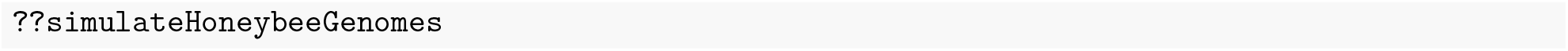

Now we are ready to setup global simulation parameters using `SimParamBee`. `SimParamBee` builds upon AlphaSimR’s `SimParam`, which includes genome and trait parameters, but also global pedigree and recombination events. We usually save the output of `SimParamBee` as the `SP` object (we will assume this in all vignettes). Namely, all SIMplyBee functions will use this object if you don’t directly specify `SimParamBee` argument. `SimParamBee` additionally holds honeybee specific simulation information (Figure 1):

- default number of workers (`SP$nWorkers`) and drones (`SP$nDrones`) in a full-sized colony; these numbers are used by functions such as `createWorkers/Drones()`, `addWorkers/Drones()` and `buildUp()`;
- default number of drones that a virgin queen mates with (`SP$nFathers`)
- the *CSD* information: the chromosome of the *CSD* (`SP$csdChr`), the position (`SP$csdPos`), and the desired number of *CSD* alleles in a population (`SP$nCsdAlleles`). The number of *CSD* alleles determines the length of the *CSD* locus (`SP$nCsdSites`): `nCsdAlleles = nCsdSites**2`. By default, the *CSD* is placed on its real genomic position on chromosome 3. However, if the user simulates less than three chromosomes, the *CSD* is placed on chromosome 1;
- pedigree for each individual created in the simulation (`SP$pedigree`) if requested by `SP$setTrackPed(TRUE)`; and
- caste information for each individual created in the simulation (`SP$caste`).

You can read more about the `SimParam` and `SimParamBee` in their help pages (`help(SimParam)` and `help(SimParamBee)`).

Below we use set the number of *CSD* alleles and default number of workers and drones in a colony:

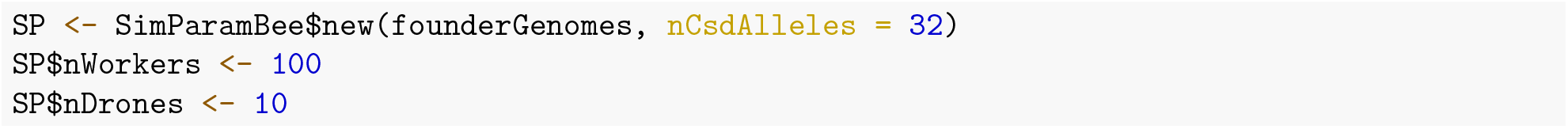

After creating the `SimParamBee` object, you can inspect it! This returns a lot of output and we suggest you return back to this point once you are comfortable with the basic functionality!

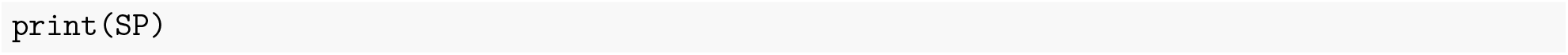

From the simulated founder genomes, we can create virgin queens (Figure 1). These will serve as our our first honeybee individuals (the so called base or founder population). In AlphaSimR and SIMplyBee, individuals are stored in `Pop` class objects, that hold a group of individuals with their individual identification, parent identifications, as well as genomes and trait values. So, the `basePop` is a population (`Pop` class object) of two individuals, our two virgin queens. If we print out `basePop`, we see some basic information about the population: the ploidy, number of individuals, chromosome, loci, and traits. We next check whether our individuals are of certain caste with `is*()` functions, where `*` can be either `queen`, `worker`, `drone`, `virginQueen`, or `father`. These functions return `TRUE` if the individual is a member of the caste in question and `FALSE` is it is not. These functions check the caste information in the `SP$caste`. Here, we use `isVirginQueen()` to check whether our base population individuals are virgin queens.

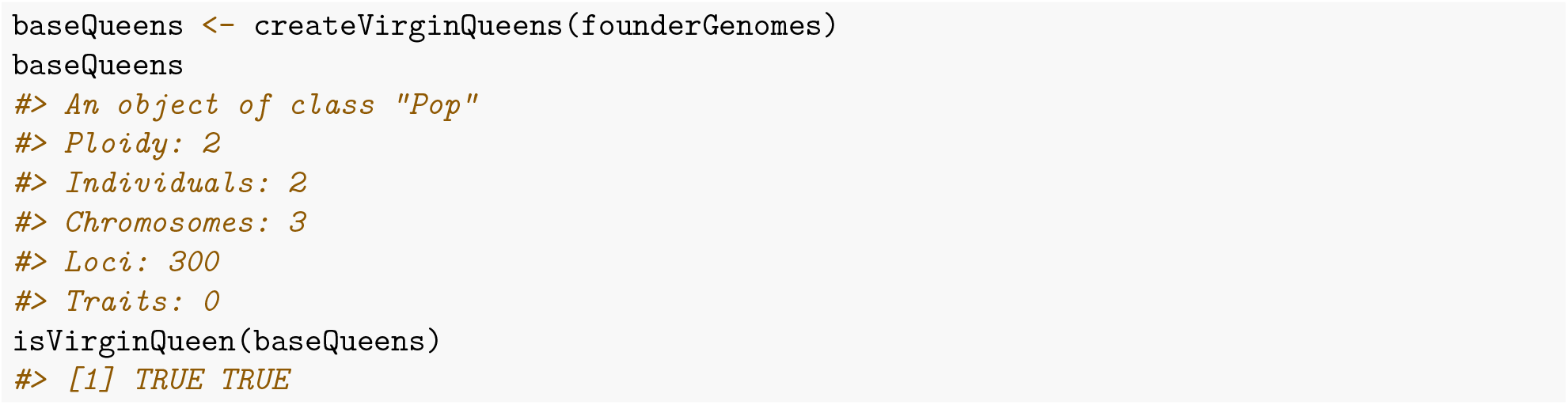

Similarly, you can use the function `getCaste()` to get the caste of each individual.

We will use the first virgin queen to create five drones for future mating. Note that virgin queens do not create drones. Only queens with colonies create drones. However, to get the simulation up and running, we need drones and the function `createDrones()` can work both with virgin queens or colonies (we will present colonies in the next section). You can use more than one virgin queen to create the drones or even an entire drone congregation area (DCA) with as many drones per virgin queen as you want (`nInd`).

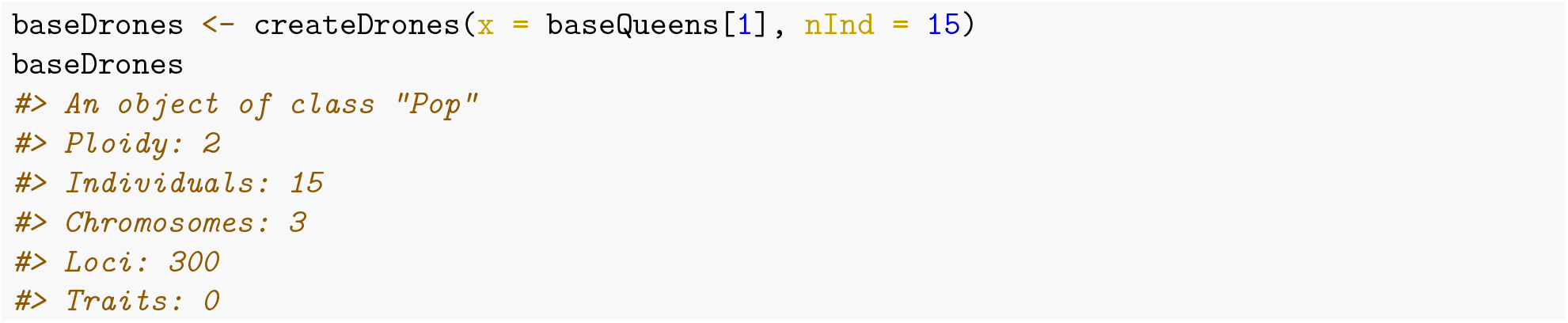

### Creating and building up a colony

We will use the other virgin queen to create a colony. You can use more than one virgin queen to create more than one colony. In SIMplyBee, a honeybee colony is stored in an object of `Colony` class. You can create a new colony with the function `createColony()`. You can create a completely empty colony or a colony with either a virgin or a mated queen. The `Colony` class organises all its members in five castes: `queen`, `fathers`, `workers`, `drones`, and `virginQueens`. We describe the castes in next section. The `Colony` further contains technical information about the colony, its identification `id` and `location` coordinates coded as (`latitude`, `longitude`). Further, it contains logical information about the past colony events: `split`, `swarm`, `supersedure`, or `collapse`. It also contains `production` status, which indicates whether we can collect a production phenotype from the colony. The latter is possible when the colony is built-up to its full size and has not swarmed. The production is turned off when a colony downsizes, collapses, or swarms, and for the split of a split colony. You will learn about these colony events in the Colony events vignette.

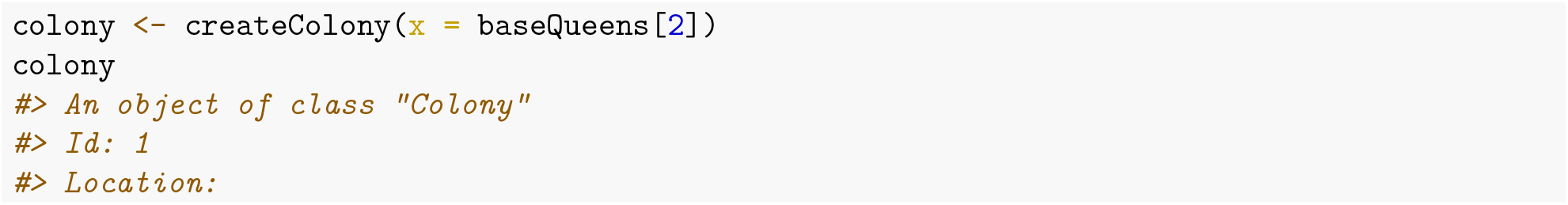

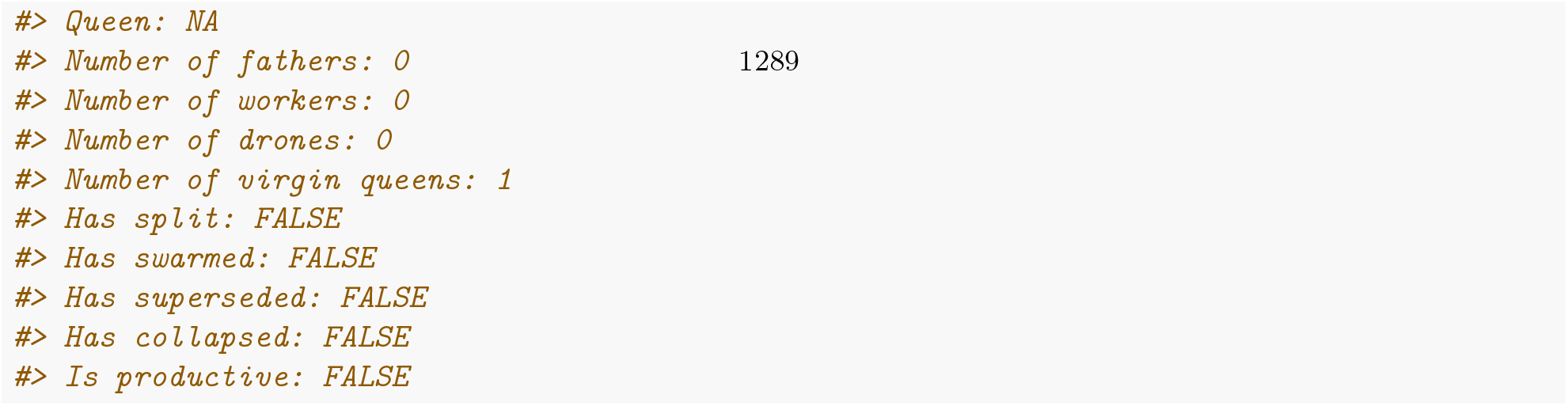

We see all the above mentioned information in the printout of the `Colony` object. For this specific colony, we see that the ID of the colony is “8”, the location is not set, and there is no queen (hence `NA`). There are consequently no fathers in the colony, nor any workers, drones or virgin queens. All the events are set to `FALSE` (you will learn more about events in the Colony events vignette) and the colony is not productive, since it does not include any individuals.

Let’s now mate our virgin queen, so that she is promoted to a queen and can start laying eggs of her own workers and drones.

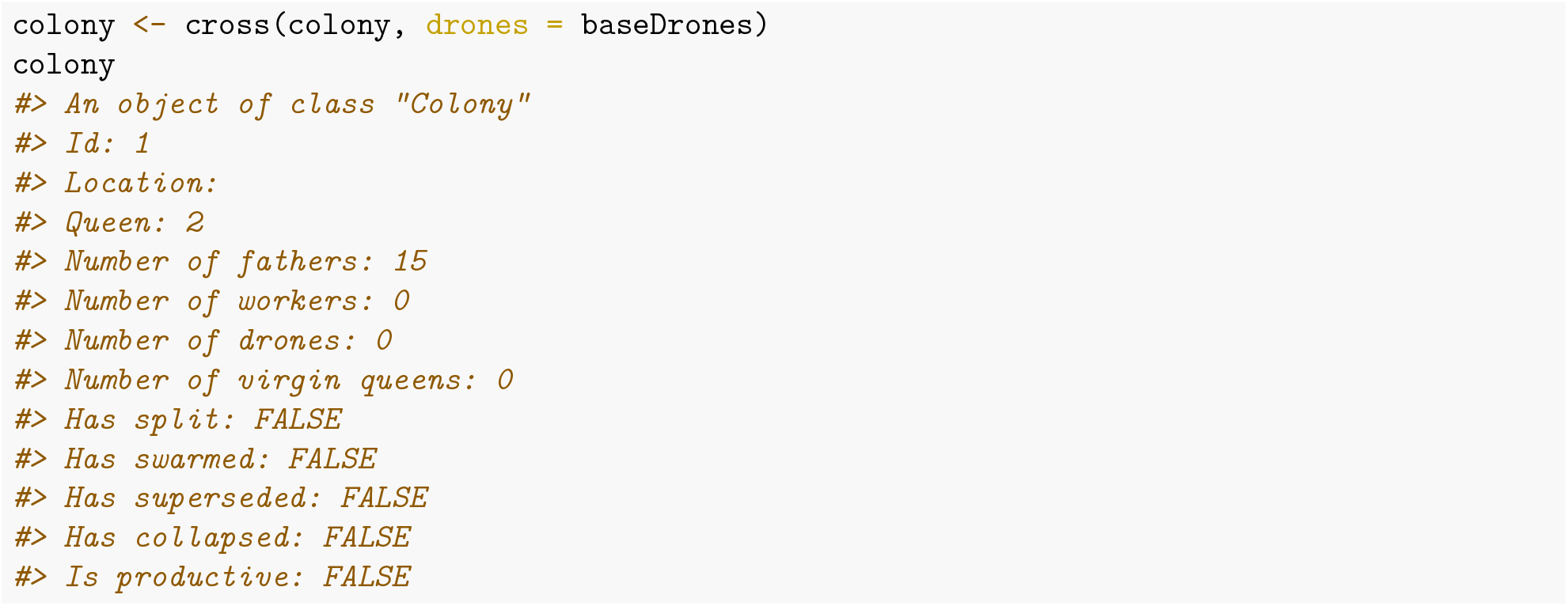

We see that the virgin queen is now a queen – hence we have a queen with the ID “2” and no virgin queens in our colony.

Next, let’s build up our colony using the function `buildUp()` that adds in workers and drones. This function takes parameters `nWorkers` and `nDrones`, where we specify how many workers and drones to add. However, if these numbers are not specified in the function’s call, the function uses the default numbers from the `SimParamBee` object (`SP$nWorkers` and `SP$nDrones`). This function also always turns the `production` status to `TRUE`, since it assumes we are building the colony up to its full-size.

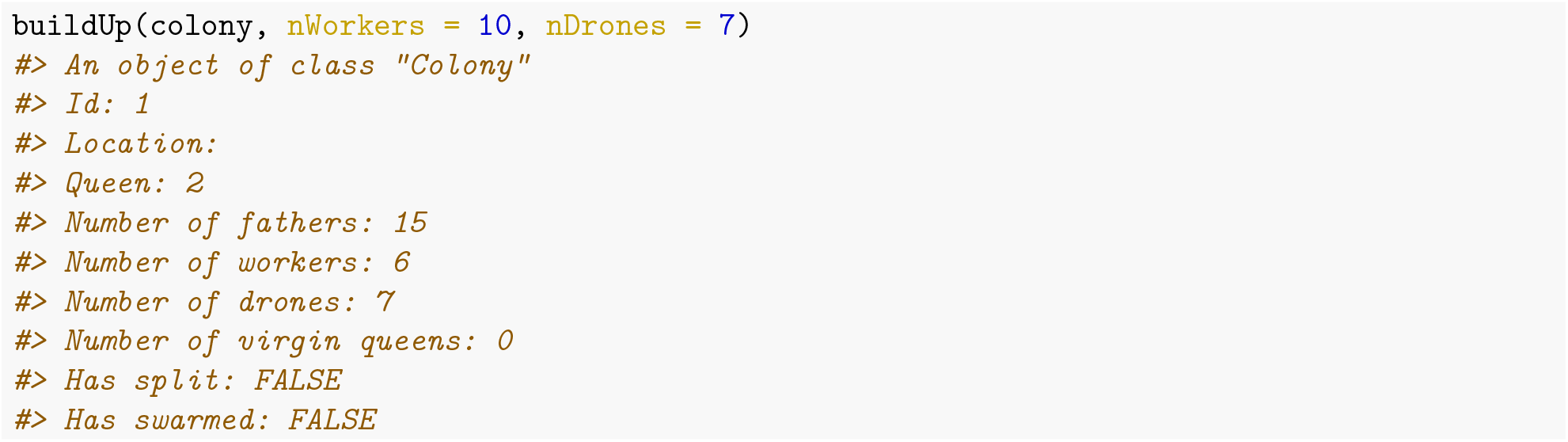

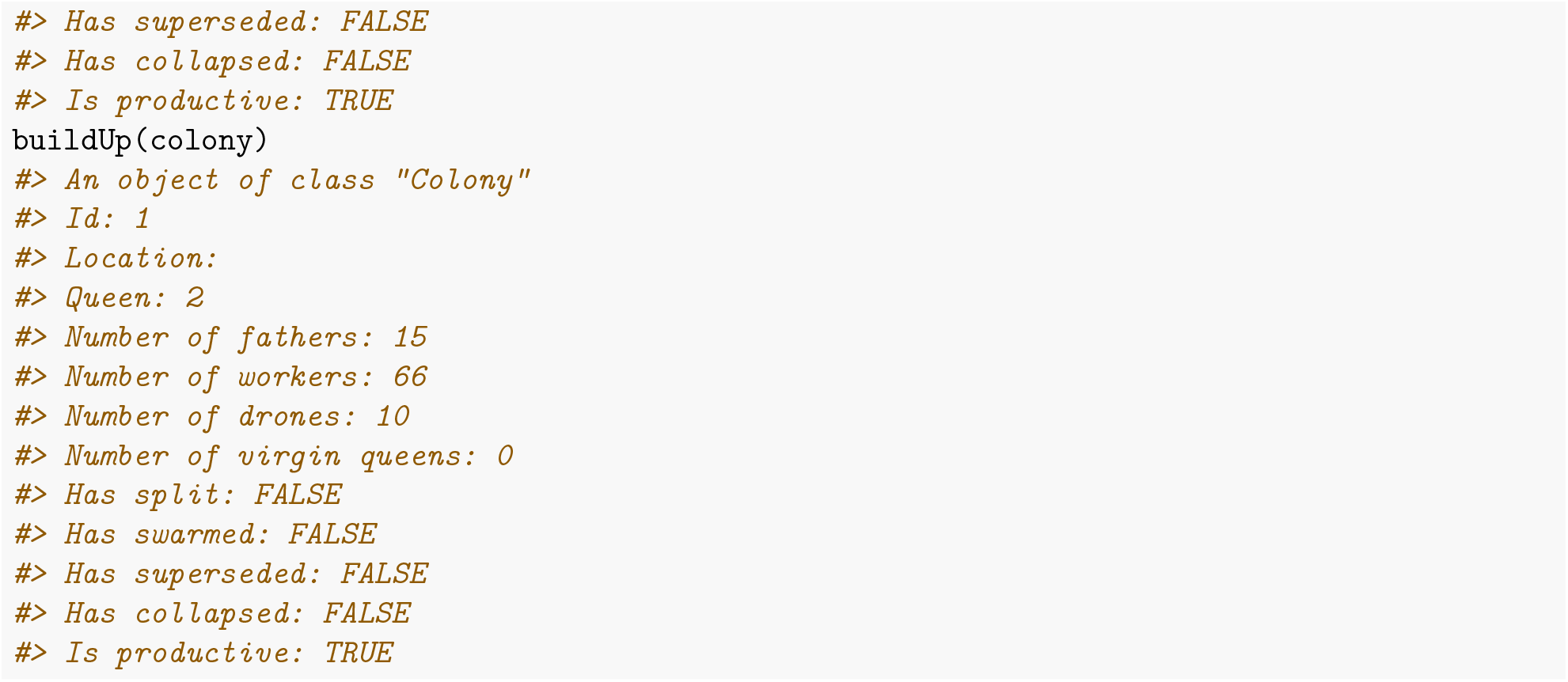

All the functions in SIMplyBee return objects, hence we need to save them as an object, otherwise they are lost.

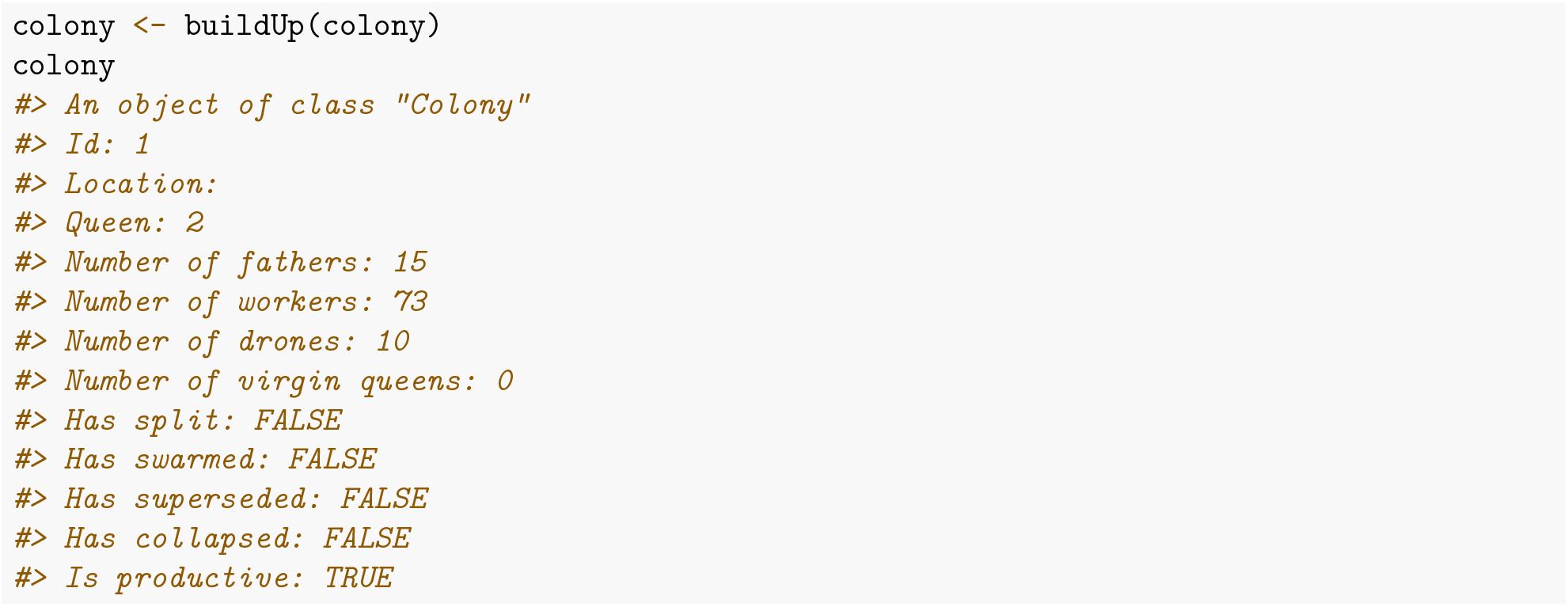

### Colony structure

Lets explore our colony. In every colony we have different groups of individuals (castes). These include: queen, fathers, workers, drones, and virgin queens. The queen controls the colony, workers do all the hard work, drones disseminate queen’s genes, and one of the virgin queens will eventually replace the queen. We also store fathers, which represent drones that the queen mated with. The fathers caste is effectively the drone sperm stored in queen’s spermatheca. Storing fathers enables us to generate colony members on demand. SIMplyBee contains `n*()` functions to count the number of individuals in each caste, where `*` is `queen`, `fathers`, `workers`, `drones`, and `virginQueens`. Let’s count how many individuals we have for each caste in our colony.

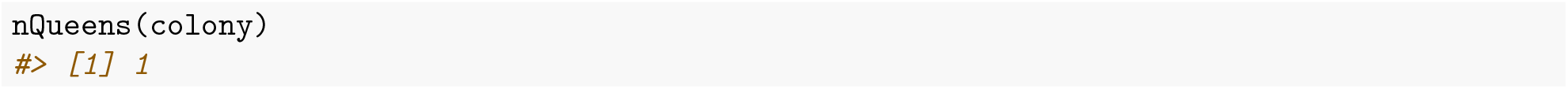

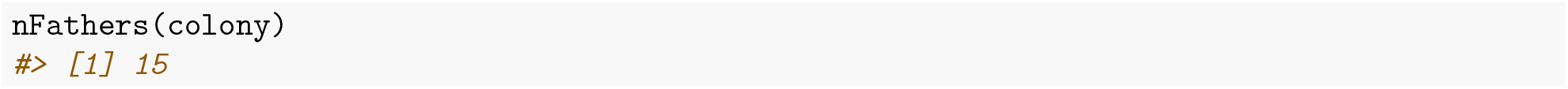

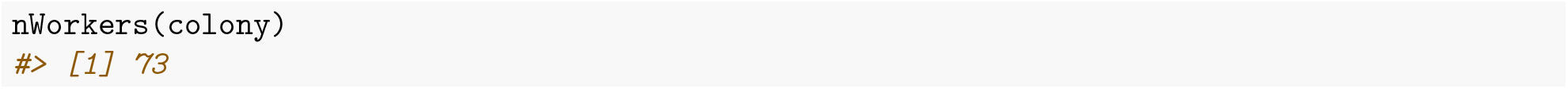

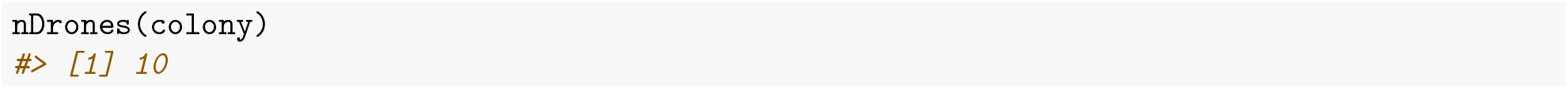

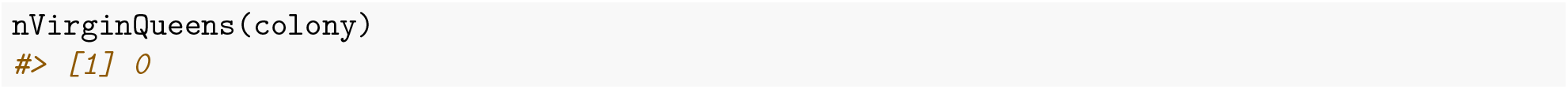

Next, we can access the individuals of each caste with `get*()` functions. These functions leave the colony and its members intact (they do not change the colony) by copying the individuals.

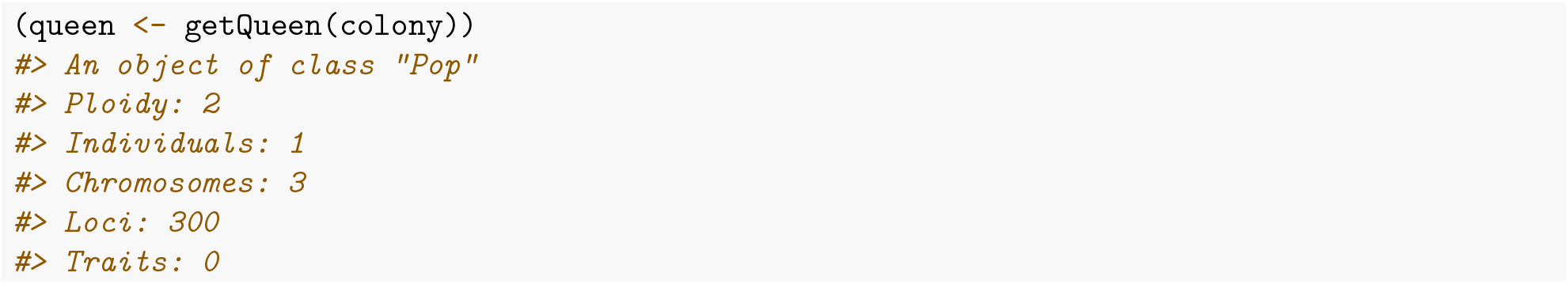

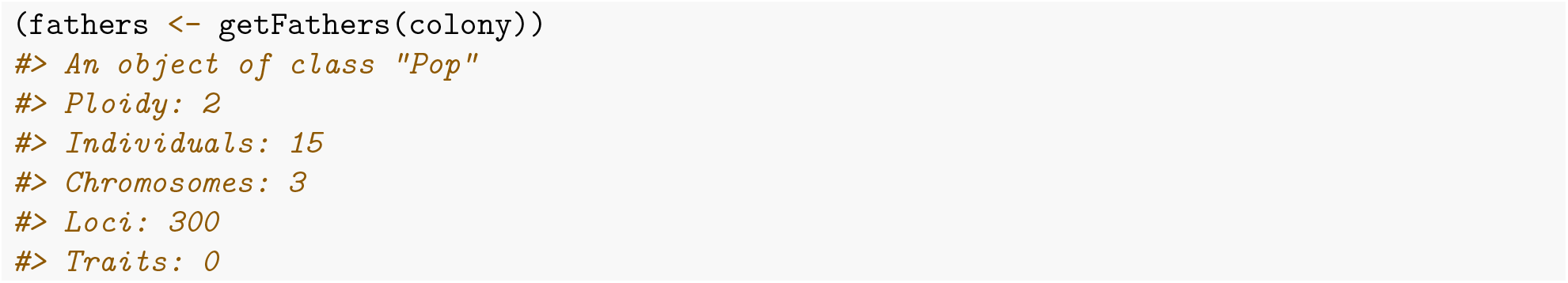

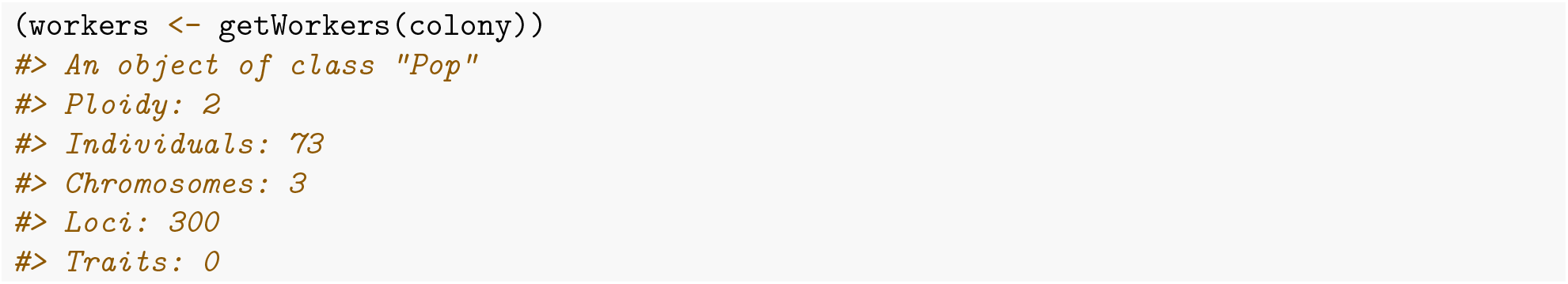

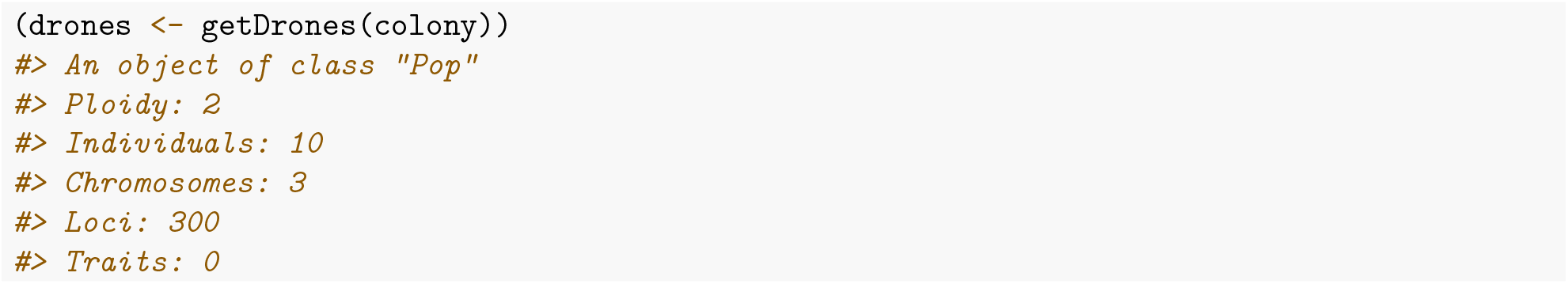

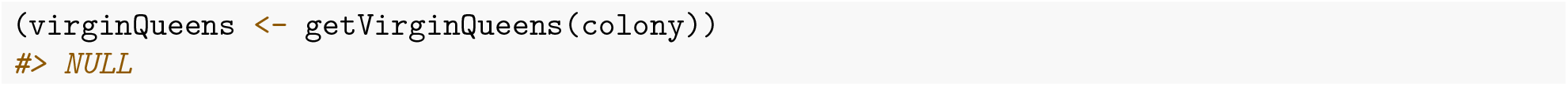

As you see above, there are no virgin queens present in the colony at this moment, since the queen is active. Future colony events might change this.

Should you want to pull out, that is, remove castes or their members, have a look at `pull*()` functions. These functions return a list of objects: pulled being the `pulled` individuals (`Pop` object), and `remnant` being the remaining colony without the pulled individuals.

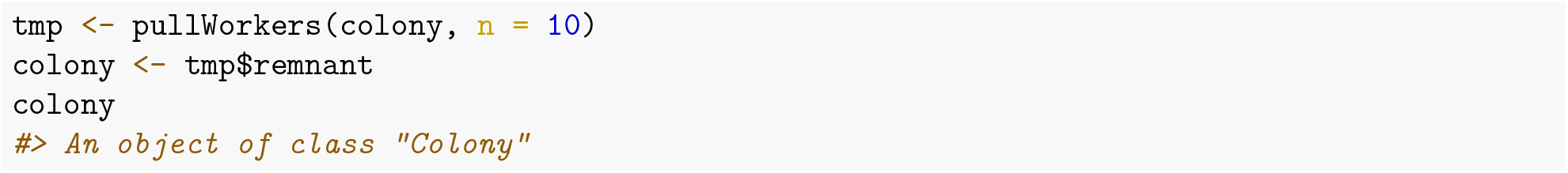

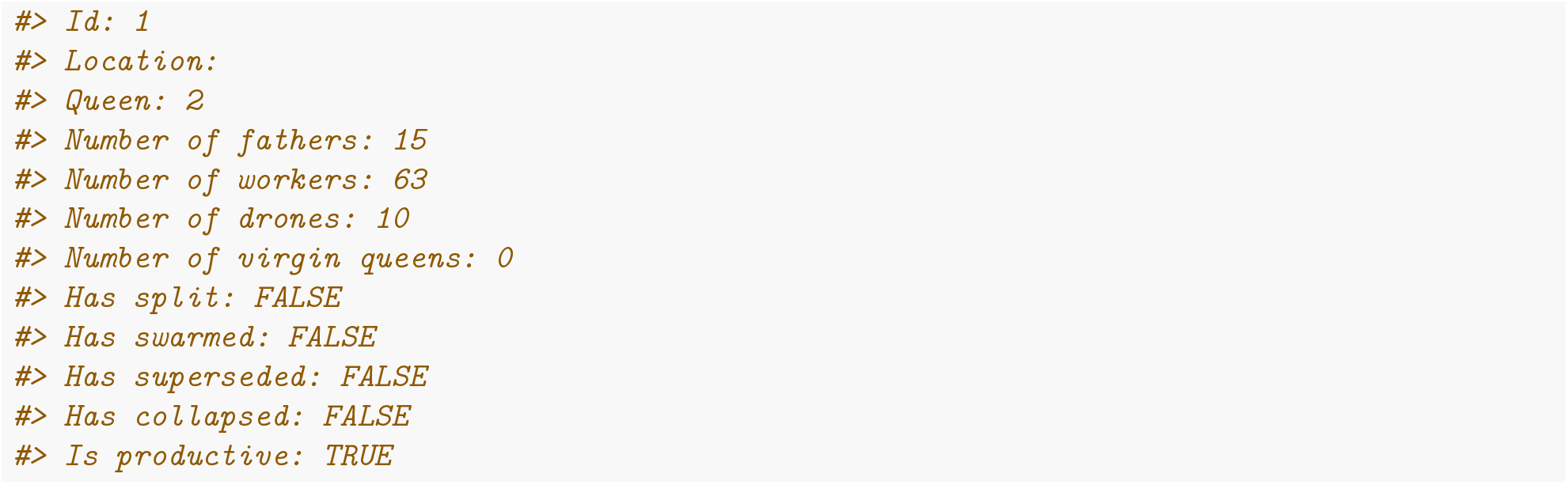

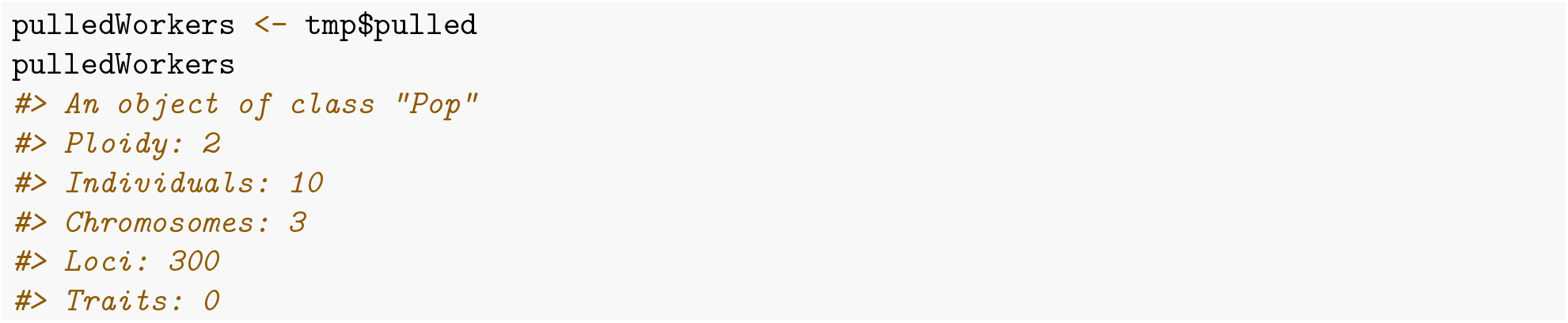

Next, you can obtain the caste of each individual with the `getCaste()` function. As already mentioned above, a similar group of functions are the `is*()` functions that check whether an individual is of specific caste. Let’s now obtain the caste of colony members:

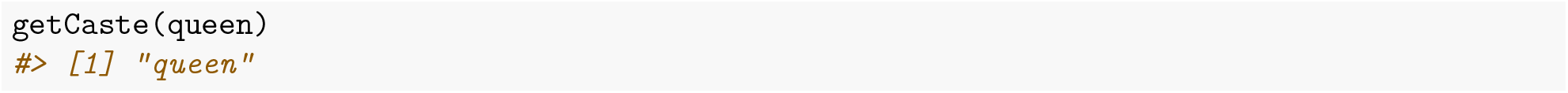

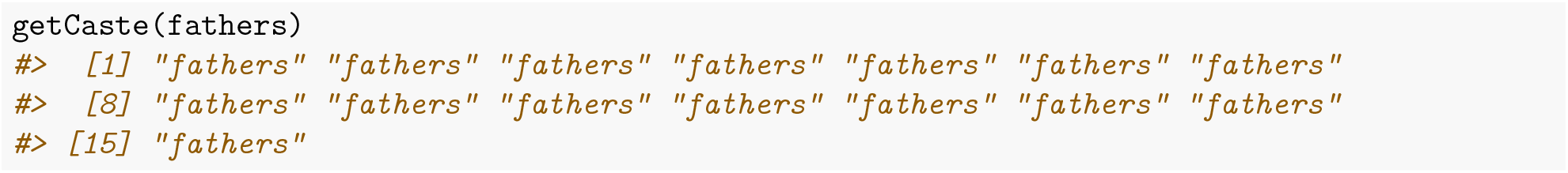

and so on. When you have a collection of bees at hand and you might not know their source, the `getCaste()` can be very useful:

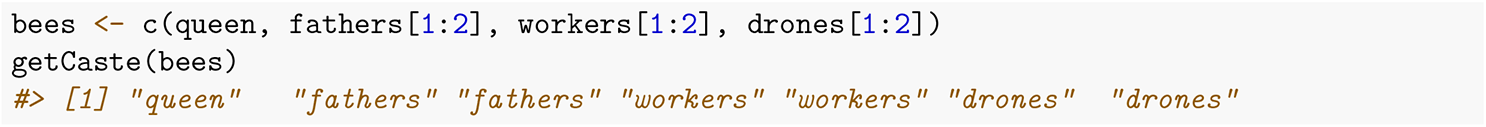

### Complementary sex determining locus

The complementary sex determiner (*CSD*) locus, well, complements sex determination. Fertilised eggs that are heterozygous at the *CSD* locus develop into workers. On the other hand, homozygous eggs develop into unviable drones. These drones are usually discarded by workers. SIMplyBee does not store these unviable drones, but it does store their number in the queen’s miscellaneous slot (queen@misc). Here, you can find the total number of workers and drones produced by the queen (`nWorkers` and `nDrones`) and how many of the diploid offspring were homozygous at the *CSD* (`nHomBrood`). There is also a `pHomBrood` slot, that represents the theoretical (expected) proportion of offspring that are expected to be homozygous based on queen’s and father’s *CSD* alleles. You can obtain `pHomBrood` and nHomBrood values with the corresponding `pHomBrood()` and `nHombrood()` functions that can be applied either on the queen (`Pop` class) or colony (`Colony` class) directly. You can obtain the entire `misc` slot with the `getMisc()` function.

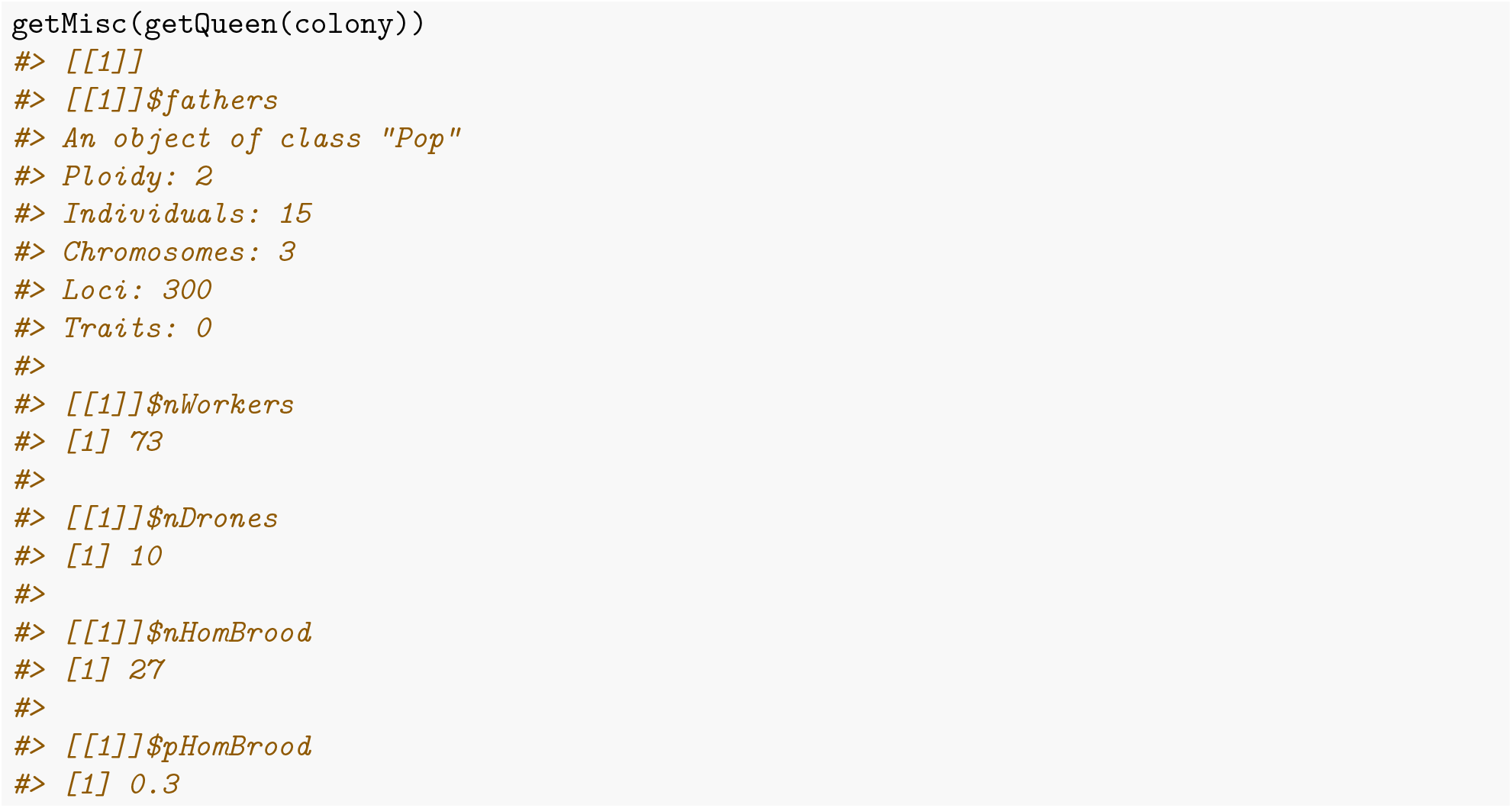

Technically, in SIMplyBee we represent the *CSD* locus as a series of bi-allelic single nucleotide polymorphisms that don’t recombine. So, the *CSD* locus is represented as a non-recombining haplotype and different haplotypes represent different *CSD* alleles. By varying the number of sites within the *CSD* locus we can control the number of distinct alleles (see `help(SimParamBee)`).

We can retrieve information about *CSD* alleles with `getCsdAlleles()`. For details on where the *CSD* locus is and the number of distinct alleles, see `help(SimParamBee)`. Looking at the below output, the first row shows marker identifications (chromosome_locus) and the first column shows haplotype identifications (individual_haplotype). The alleles are represented with a sequence of 0’s and 1’s. You can see that the two sequences are different, meaning that the queen is heterozygous, as expected.

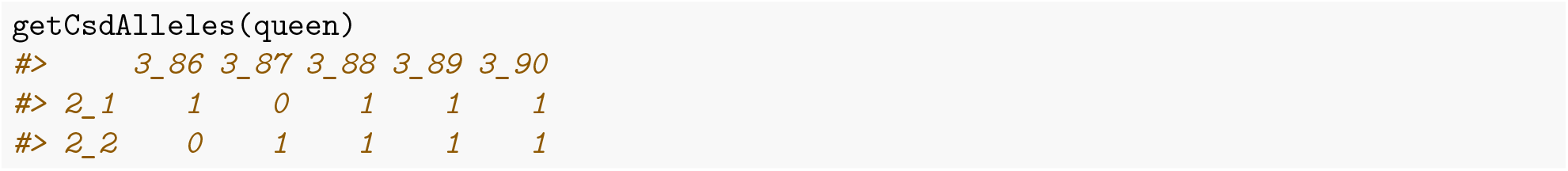

A keen geneticist would immediately inspect *CSD* alleles of fathers to check for any similarity with the queen’s *CSD* alleles. Let’s boost a chance of such an event by creating an inbreed colony. We will create a virgin queen from the current colony and mate her with her brothers. Oh, dear.

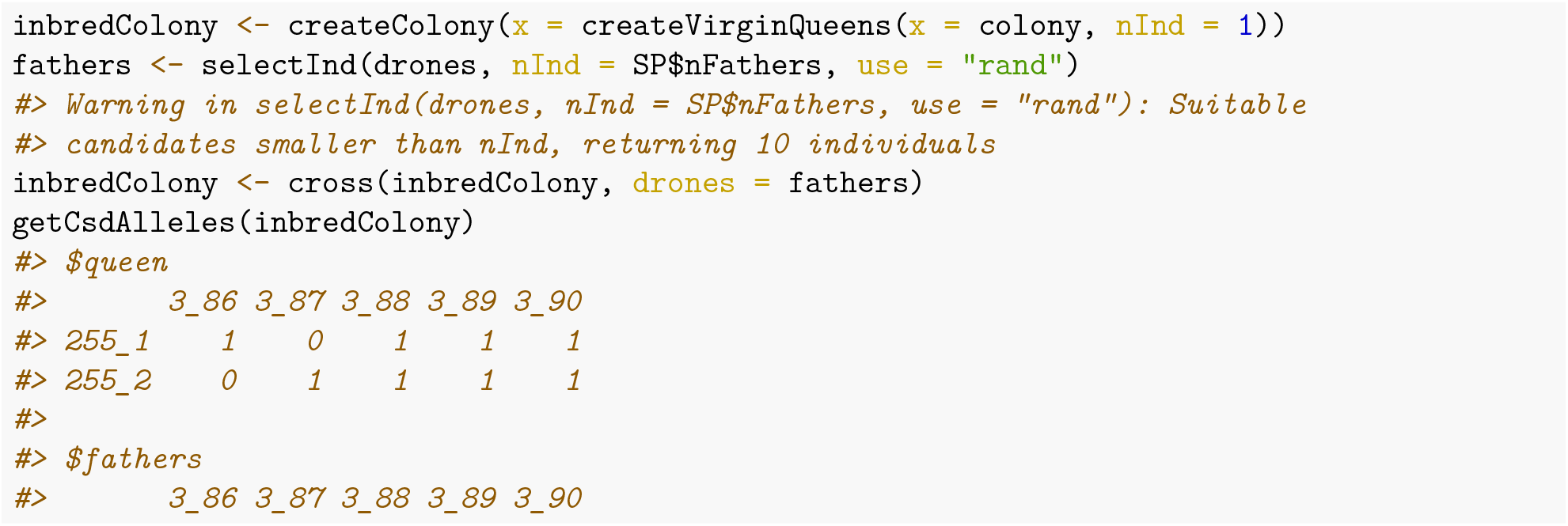

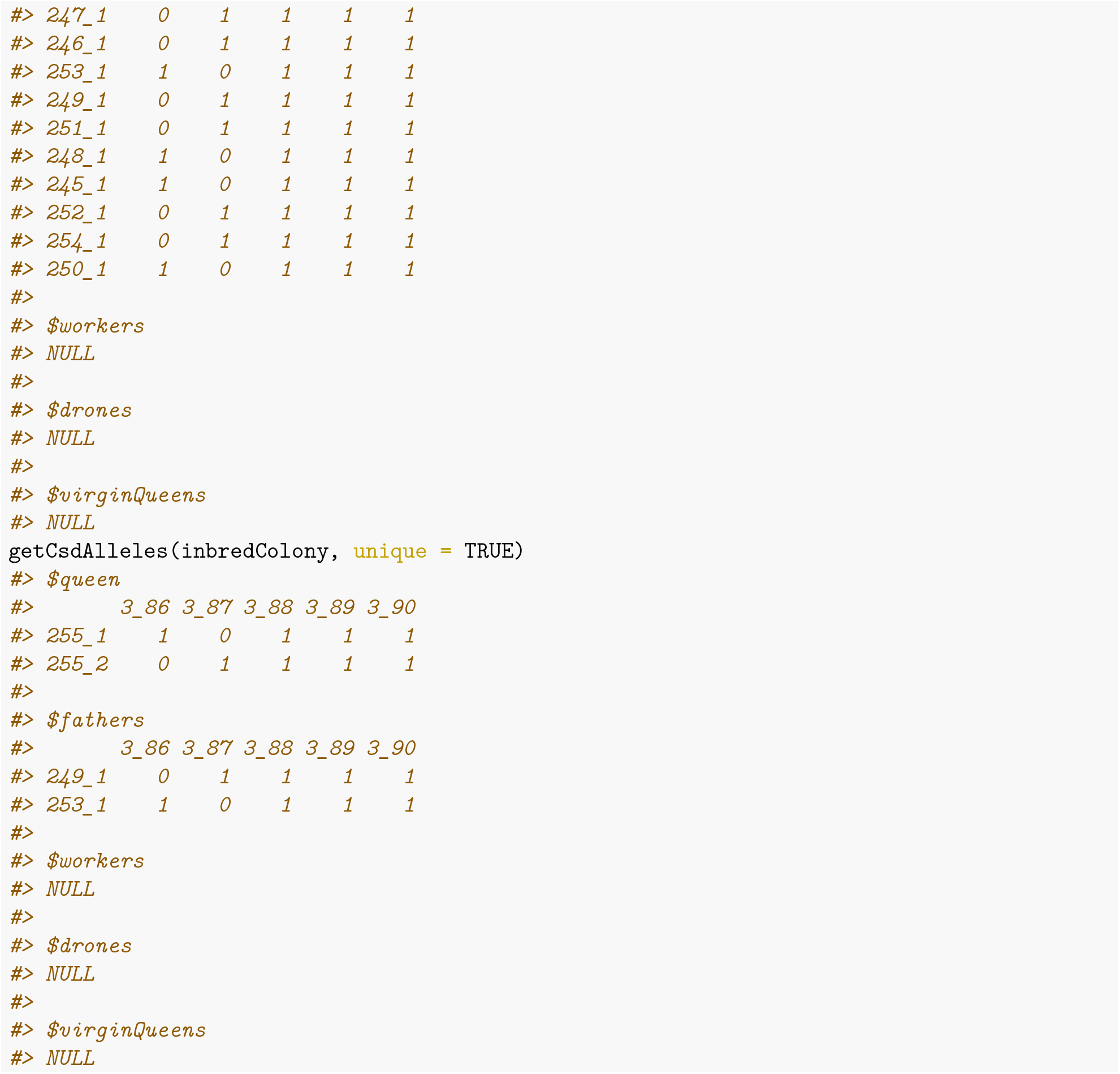

Can you spot any matches? Let’s calculate the expected proportion of homozygous brood from this mating.

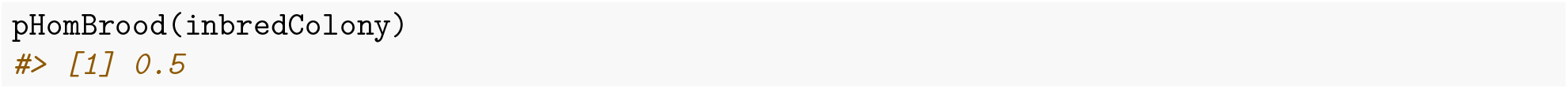

Let’s see how many homozygotes will we observe. Note that inheritance is a random process, so a realised number of homozygotes will deviate from the expected proportion.

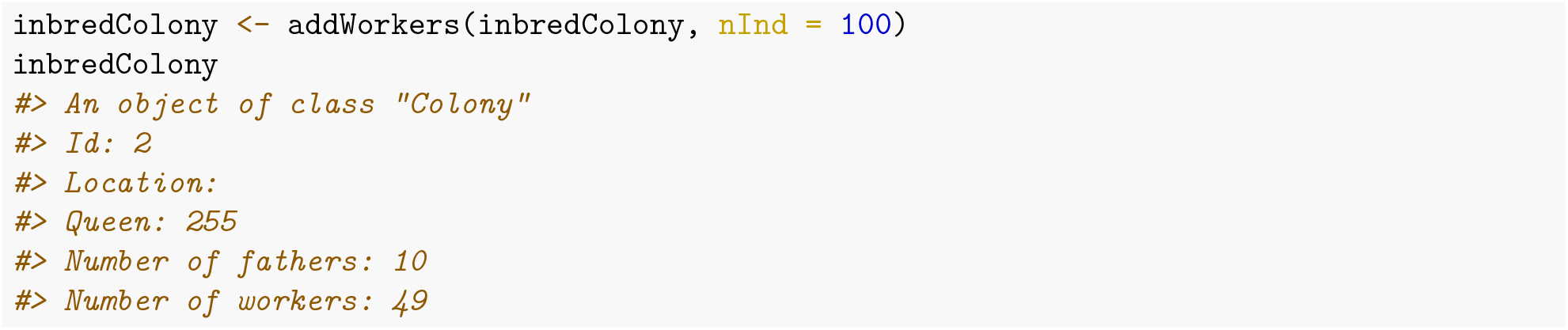

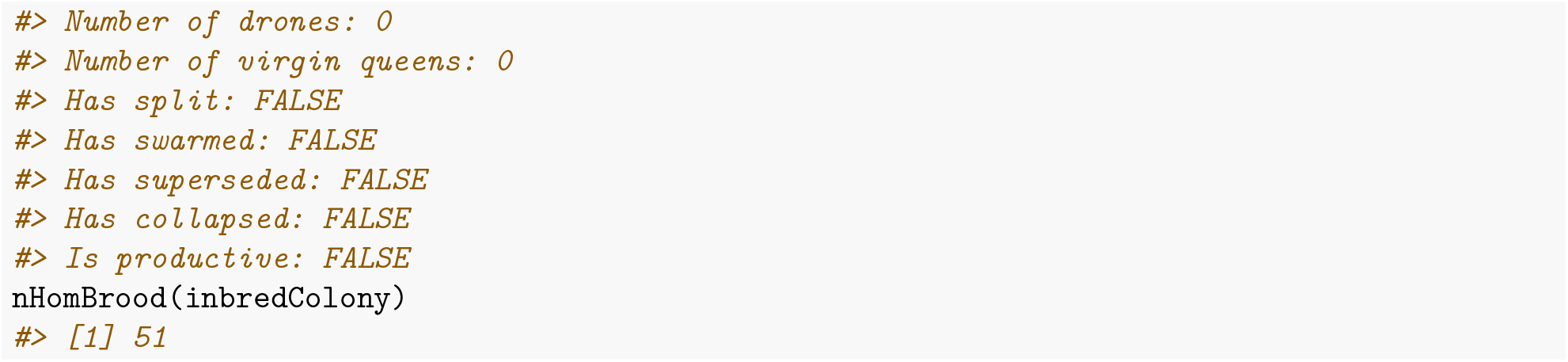

We tried adding 100 workers, but we only got 49. The difference of 51 is due to *CSD* homozygous brood. Let’s add another set of workers to show variation in the realised numbers and accumulation of information.

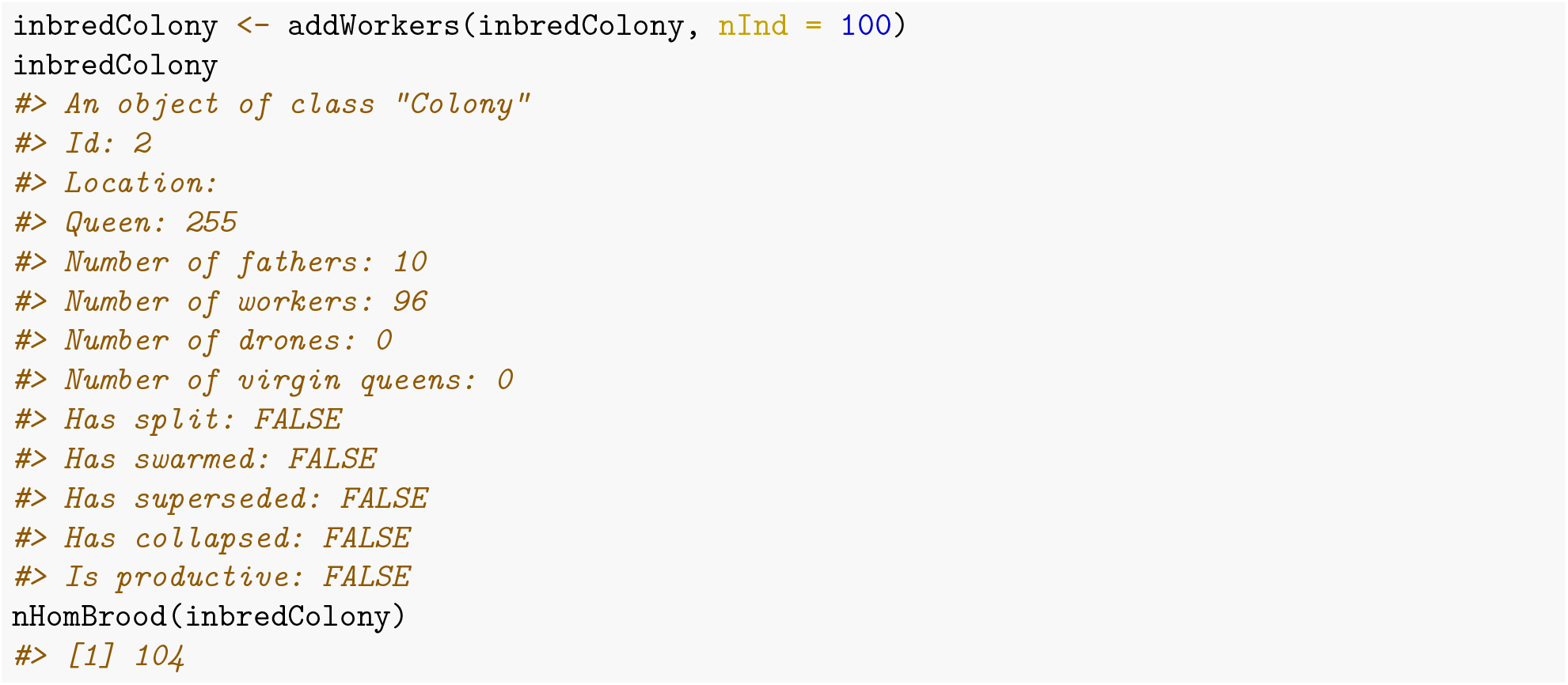

In total we tried adding 200 workers. We got 96 workers and 104 homozygous brood. To see all this information, we can inspect the miscellaneous slot of the queen that contains the fathers population as well as the cumulative number of workers, drones, homozygous brood, and the expected proportion of homozygous brood.

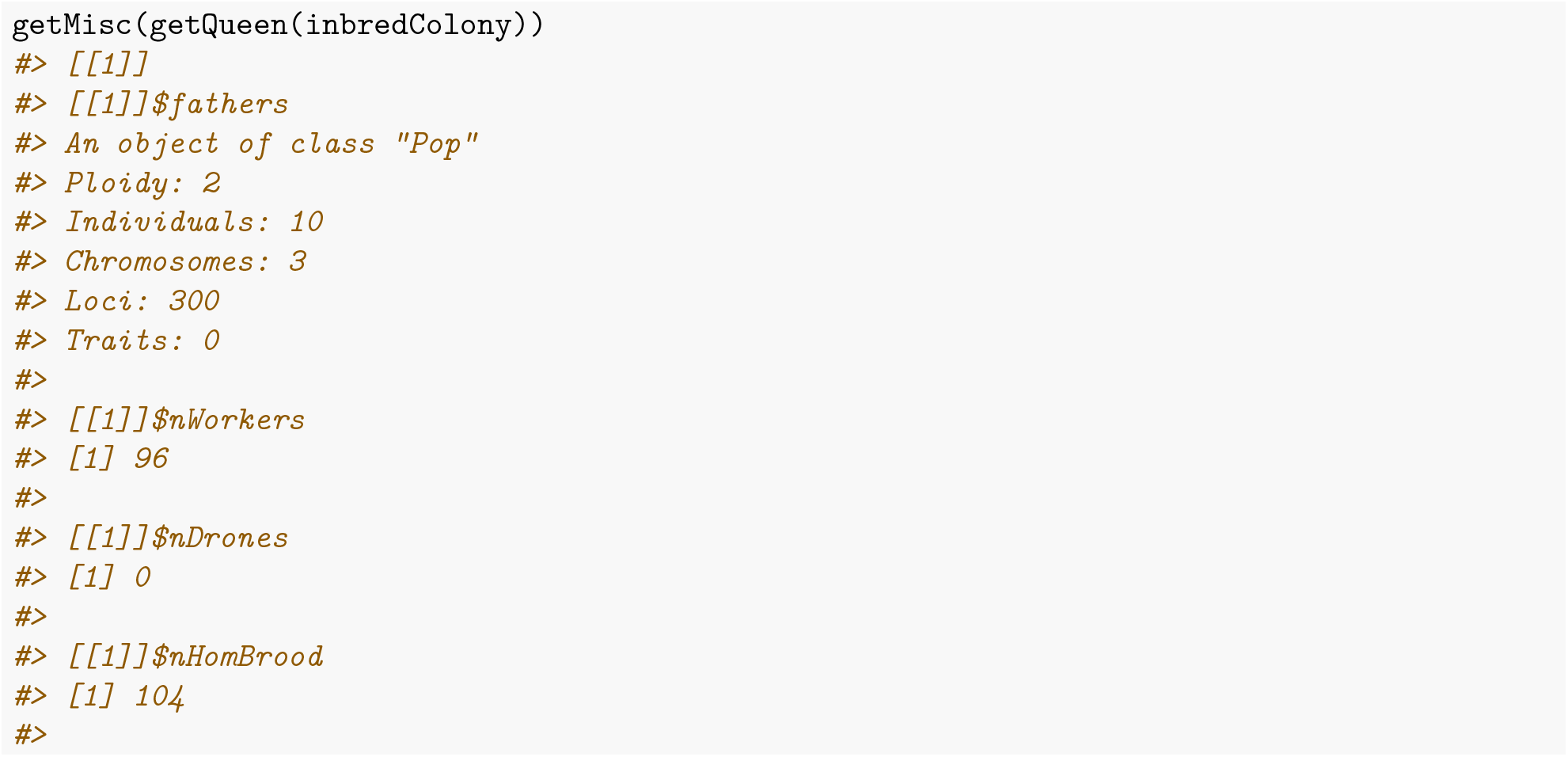

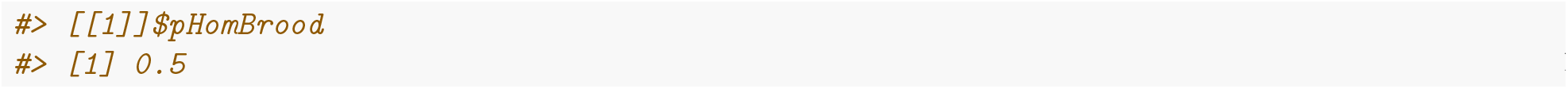

## Additional file 2 – Multiple colonies vignette

### Introduction

We have already introduced the Colony class that holds colony-specific information and caste individuals. However, when working with honeybees, we usually do not work with a single colony, but with apiaries or even whole populations of colonies. To cater for this, SIMplyBee provides a `MultiColony` class. It behaves as a list of `Colony` objects but with additional functionality – you can apply function directly to the `MultiColony` objects. A `MultiColony` can represent different apiaries or sub-populations in terms of either age of the queens or geographical location of the apiaries etc. This vignette demonstrates creating and working with `MultiColony` objects. First, we again load the package.

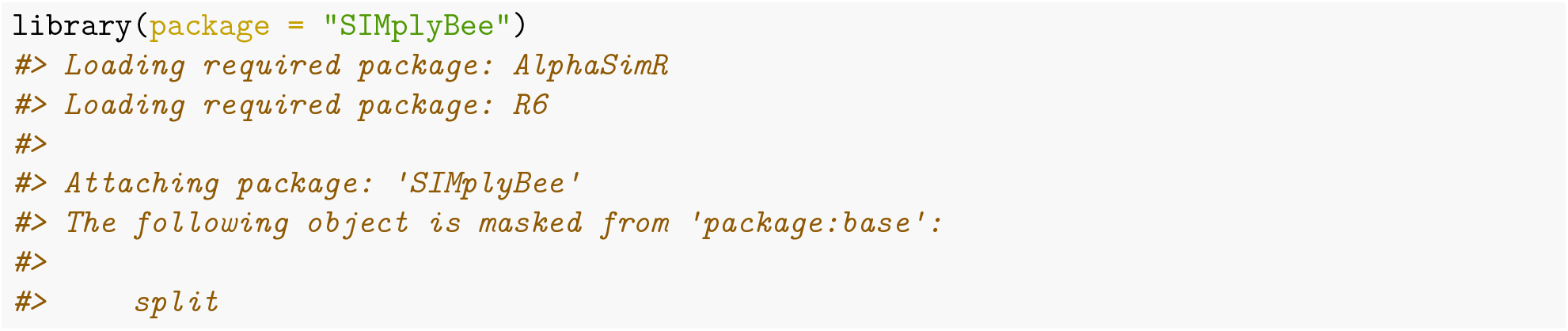

### Creating a MultiColony object

We create a `MultiColony` object with `createMultiColony()` function. Let’s say you want to create a `MultiColony` object that represents a single apiary. The first option is to initialise an empty `MultiColony` object that represents an empty apiary without any colonies and individuals within them.

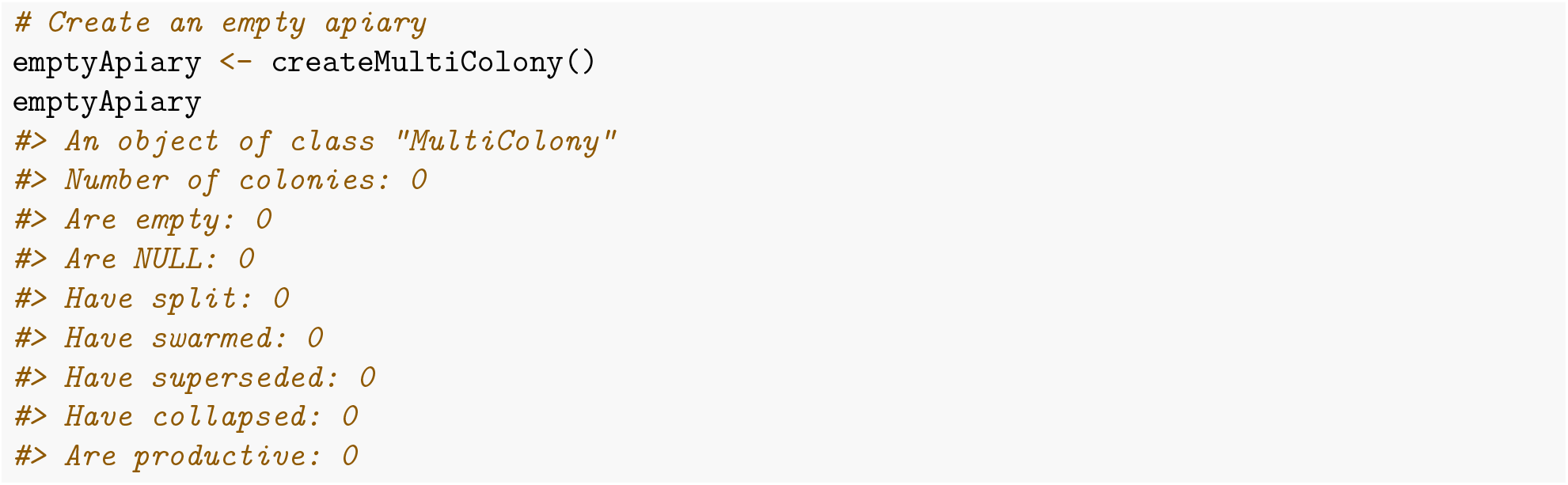

Let’s inspect the printout of the `MultiColony` object. This tells how many colonies are within, how many of them are `empty` and contain no individuals, how many are `NULL` objects, how many have experienced a split, swarm, supersedure, or a collapse (you can read more about these events in the Colony events vignette), and how many of them are productive, meaning that we can collect a production phenotype from them such as honey yield.

The second option is again to create an empty `MultiColony` object that represents an empty apiary without any individuals within, but with a defined number of colony slots.

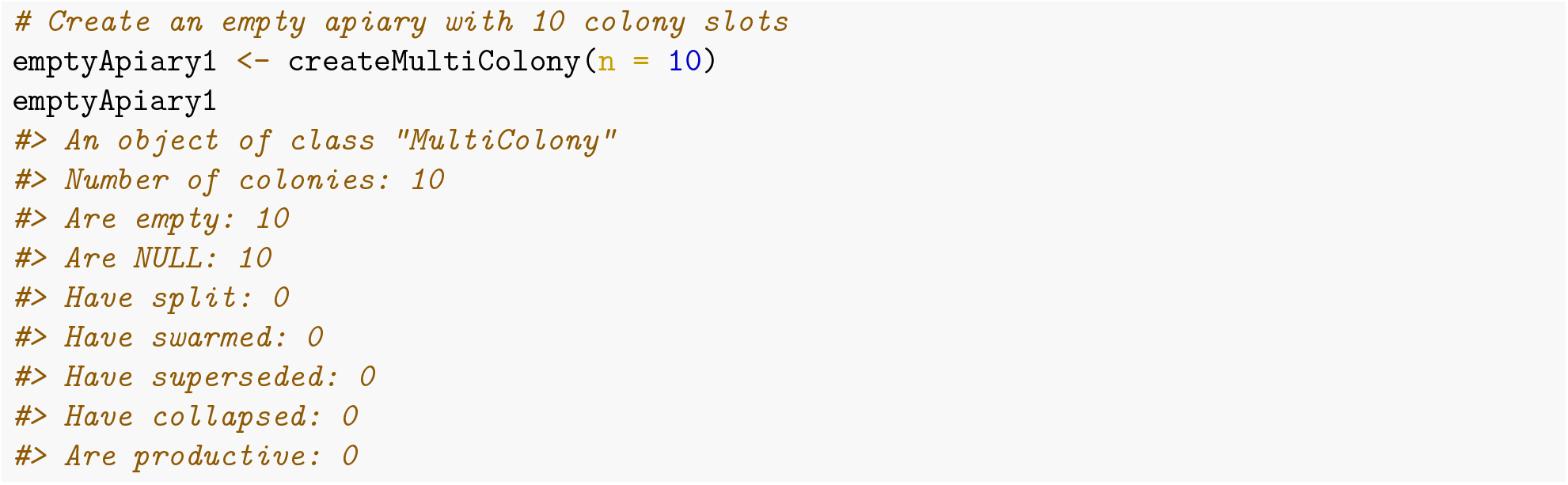

The third option is to create a `MultiColony` object with a population of either virgin or mated queens. For this, we first have to initialise the simulation with founder genomes and creating a base population of virgin queens. We will use 10 virgin queens to produce drones and create a DCA.

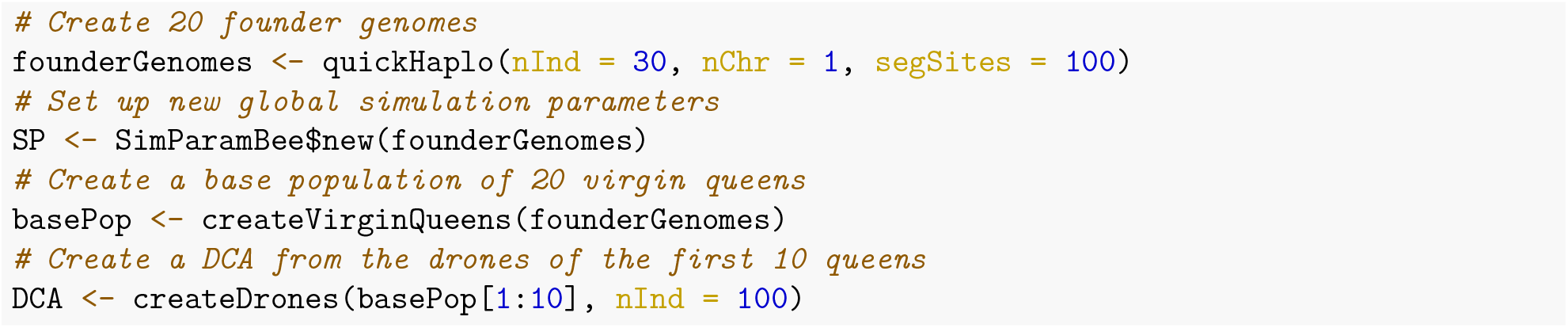

We will now create an apiary with 10 virgin colonies with the `createMultiColony()` function by providing the second set of 10 virgin queens as the input parameter. Let’s call this apiary `apiary1` and say that it is positioned at the location (`1,1`).

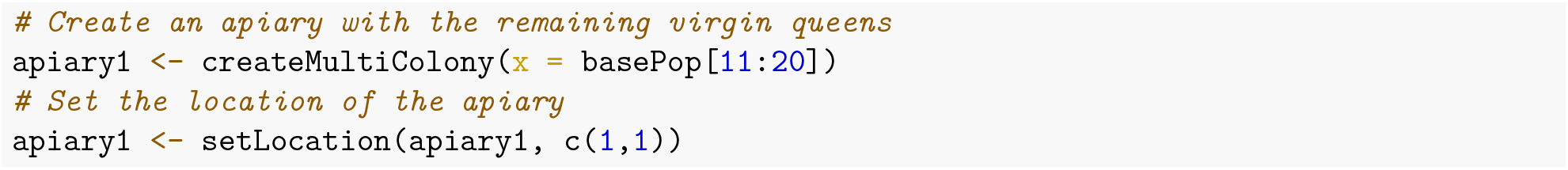

Let’s now use functions `isQueenPresent()` and `isVirginQueensPresent()` to confirm all the colonies are virgin.

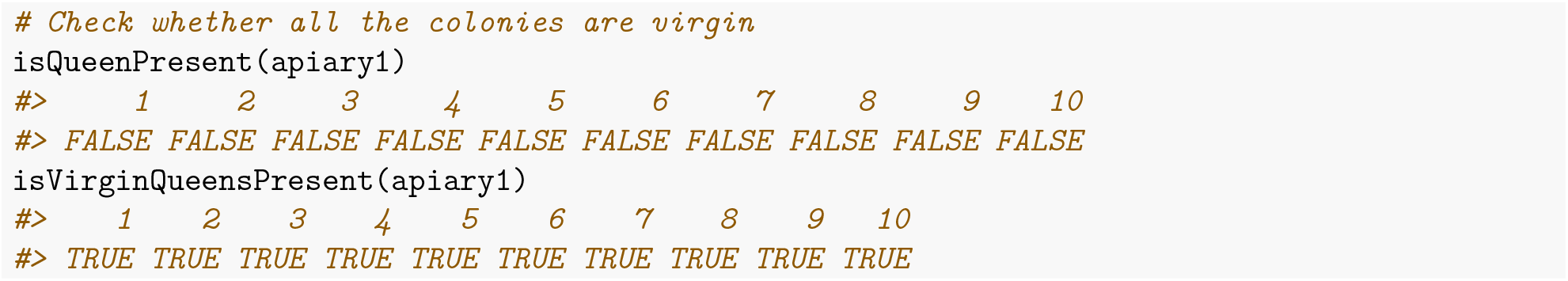

### MultiColony operations

Once we have a non-empty `MultiColony` object, we can do basic operations on it. First, we can select some colonies by either specifying their IDs, desired number or percentage of randomly selected colonies.

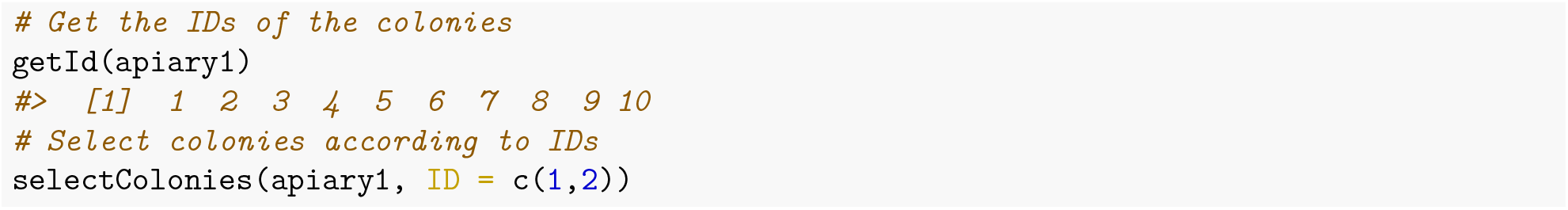

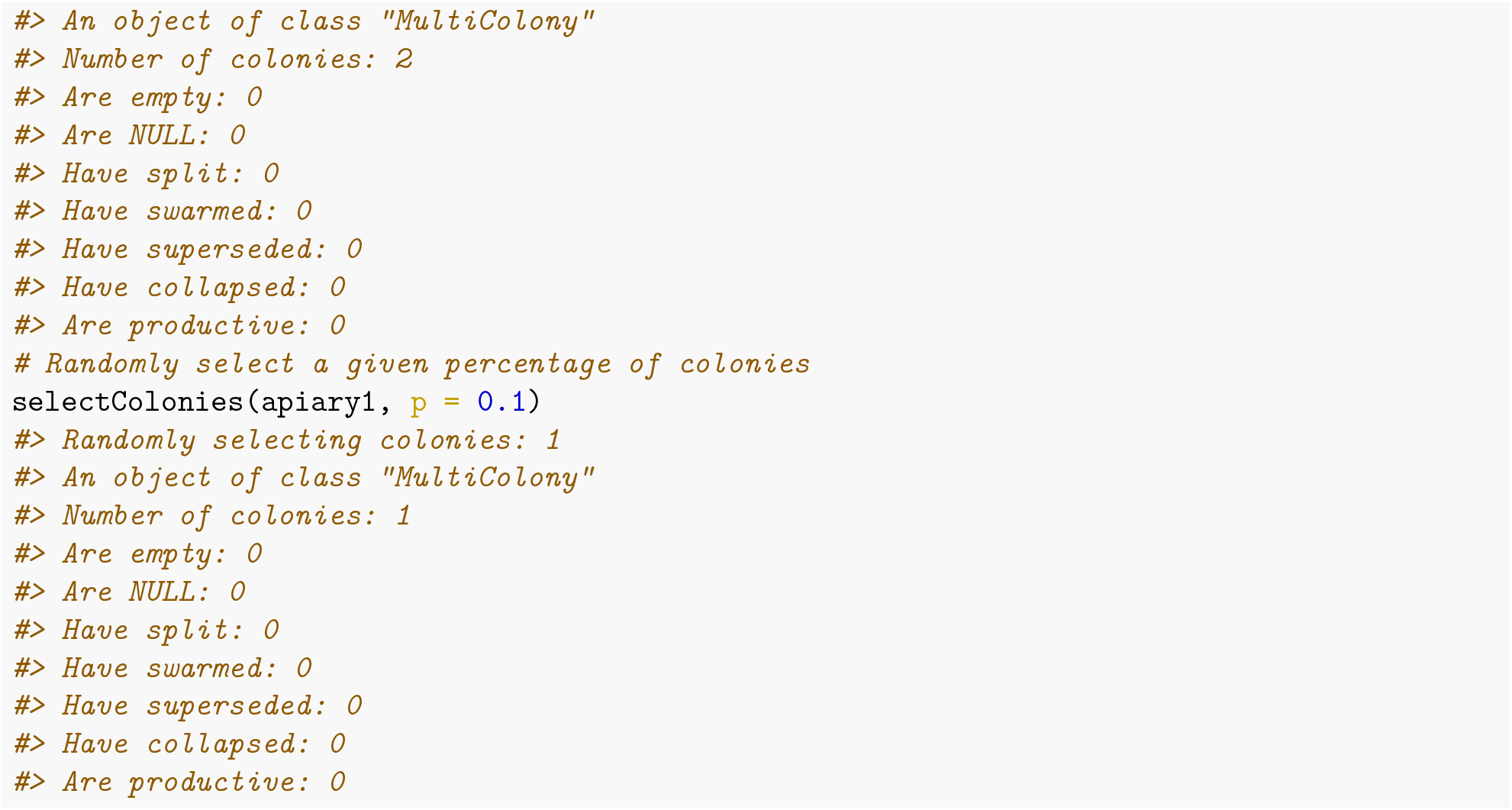

Second, we can pull some colonies from the `MultiColony` object. This means, that the pulled colonies are removed from the original object. The function `pullColonies()` therefore returns two object – the pulled colonies and the remnant colonies.

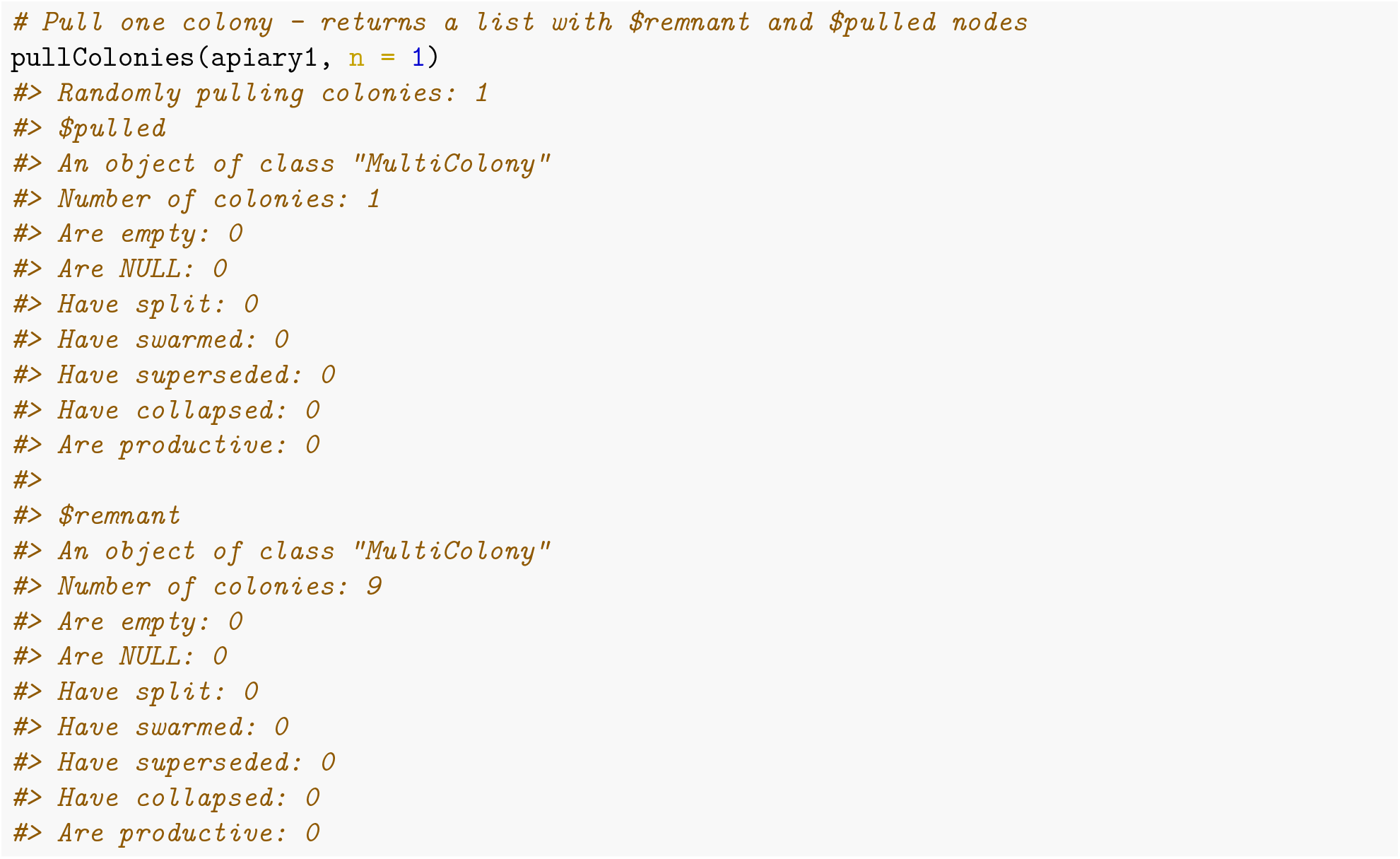

Third, we can also remove some colonies from the `MultiColony` object with `removeColonies()` function.

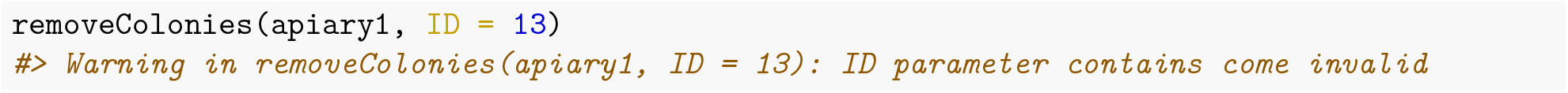

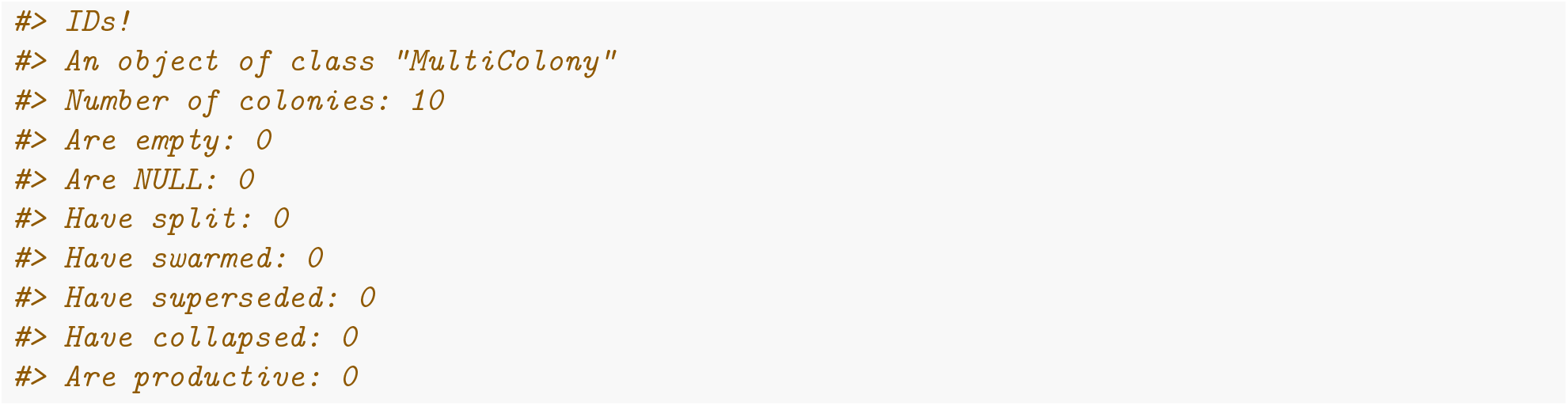

These three functions can also select, pull, and remove colonies based on some values (phenotypes, genetic values…). You can read more about that in the Quantitative genetics vignette.

### Crossing a MultiColony

Next, we will cross all the virgin queens in the apiary with the `cross()` function to groups of drones that we collected from the DCA with the `pullDroneGroupsFromDCA()` function. We have to collect at least as many groups of drones as we have colonies in our `MultiColony`.

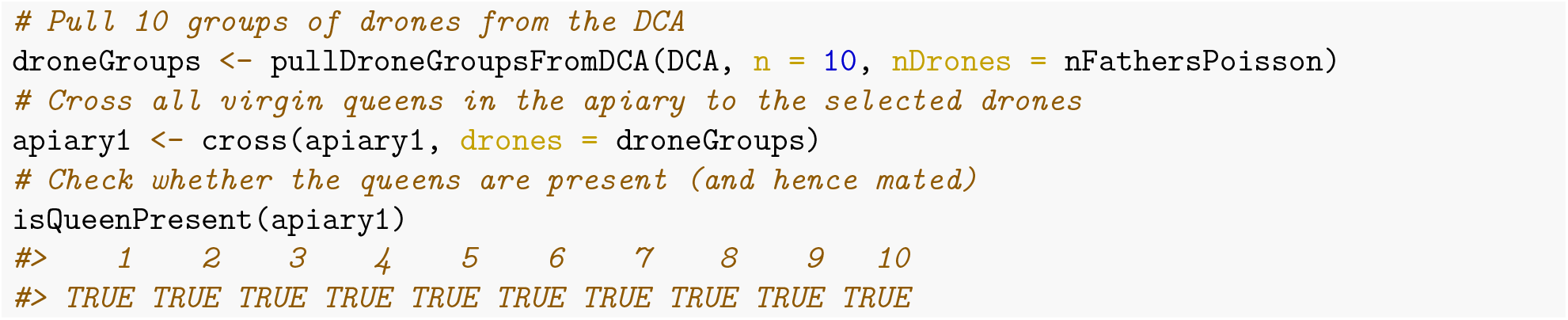

Once we have mated queens in the apiary, we can apply all the event functions directly to the `MultiColony` object: `buildUp()`, `downsize()`, `swarm()`, `split()`, `supersede()`, `collapse()` but also all the functions that either add, replace, or remove individuals from the castes. Let’s say we want to build-up all the colonies in our apiary.

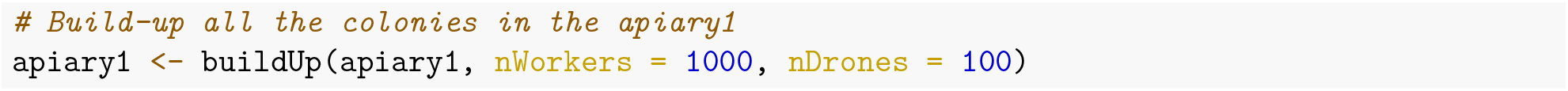

Furthermore, we can use the `pullColonies()` or `selectColonies()` to subset the colonies that will for example swarm, collapse, or supersede (presented in the Colony events vignette), or the ones that we decided to split (check out the Colony events vignette).

### Working with multiple MultiColony objects

Let’s now initiate another `MultiColony` named as `apiary2` that is placed at location (`2,2`). Here, we define different `MultiColony` object according to the location of the apiary, but the objects could also be defined according to the age of the queens (such as `age0`, `age1`…). `apiary2` contains only virgin queens and we want to mate them to a DCA made of drones from `apiary1`.

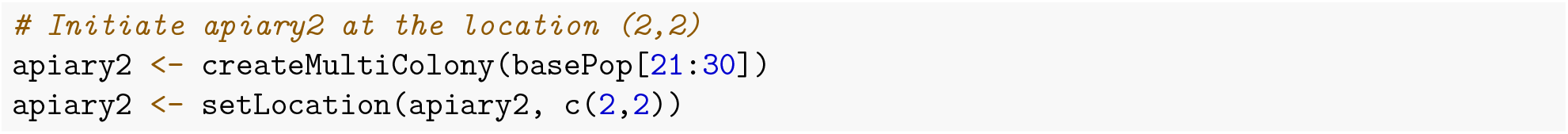

Since some time has passed, we want to first replace the drones in `apiary1` with new drones. We can do that with `replaceDrones()` function.

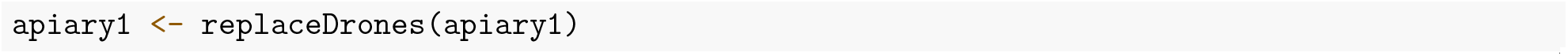

Now that we have a new set of drones, we can create a DCA with the function `createDCA()` and mate virgin queens in apiary2 to the DCA.

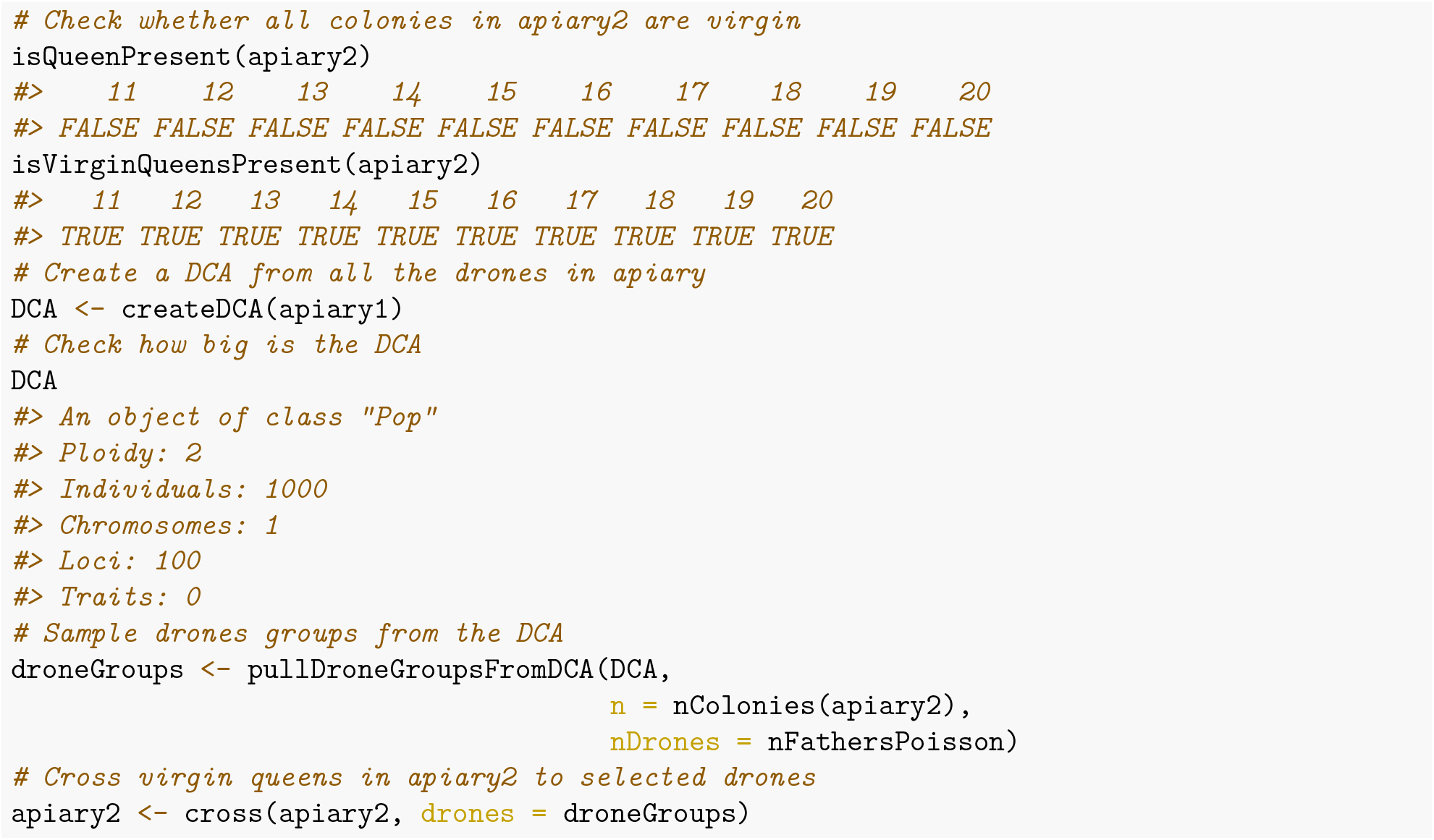

To learn more about the `nFathersPoisson()` function and other similar functions, read the Sampliong functions vignette.

## Additional file 3 – Colony events vignette

### Introduction

This vignette will introduce you to honeybee colony events. SIMplyBee implements the most important natural events like swarming, supersedure, and collapse of the colony, and a beekeeping management practice called splitting. All functions that implement colony events work both on `Colony` and `MultiColony` objects.

First, we start by loading the package.

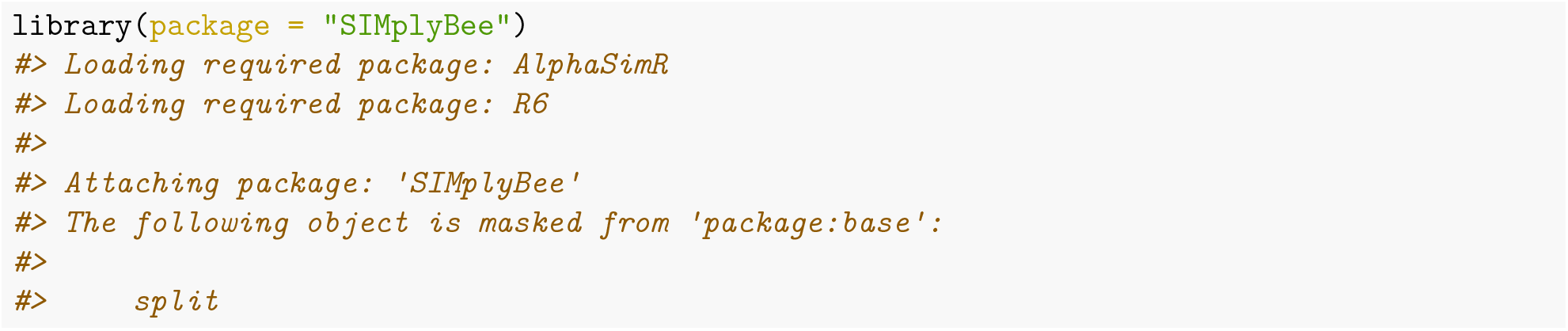

The code below generates 6 colonies so we can demonstrate various colony events.

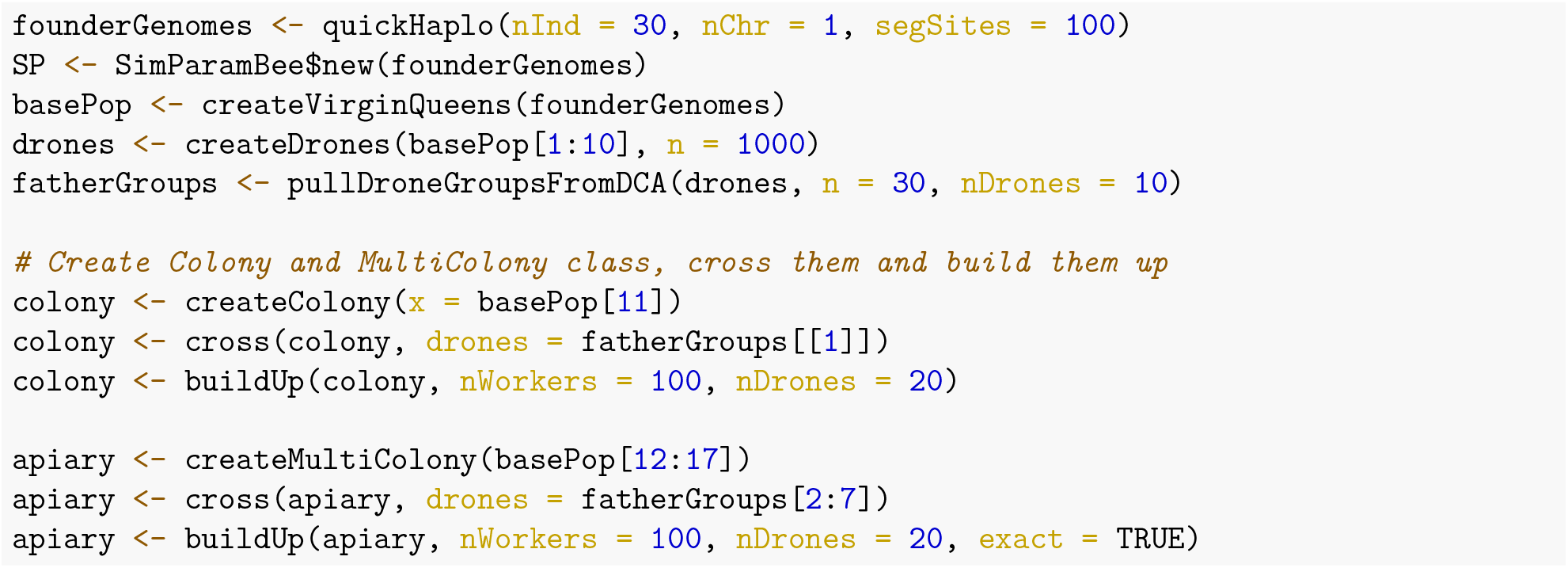

### Swarming

Swarming is the process through which honeybee colonies produce new colonies. When a honeybee colony outgrows its hive, becomes too congested, or too populated for the queen’s pheromones to spread among workers, then the swarming begins. The workers start building swarm cells for new virgin queens. When the queen is ready, she leaves the hive and is followed by about 70% of the workers in a massive cloud of flying bees, the swarm (Rangel and Seeley, 2012). The swarm will cluster on a nearby tree or bush and remain there until they find a suitable new home.

The virgin queens developing in the old hive are daughters of the queen that swarmed and are attended by the remnant workers that did not leave with the swarm. After few days, the new virgin queens begin to emerge. Typically, the first queen to emerge will kill the rest of virgin queens to assume the role as the new queen for the colony. She will then go on a mating flight to find drones to mate with to begin laying eggs and rebuilding the workforce in the colony (Clemson Cooperative Extension, 2021; Rangel and Seeley, 2012).

In SIMplyBee, function `swarm()` simulates swarming (Figure 1). The function takes a `Colony` class object and a percentage `p` of workers that leave with the swarm. The function returns a list with two `Colony` class objects, `swarm` and `remnant`. The `swarm` contains the old queen and `p` percentage of workers that left the hive. The `remnant` contains the rest of workers (`1-p`), all the drones, and virgin queens that are daughters of the old queen that swarmed.

**Figure 1:**
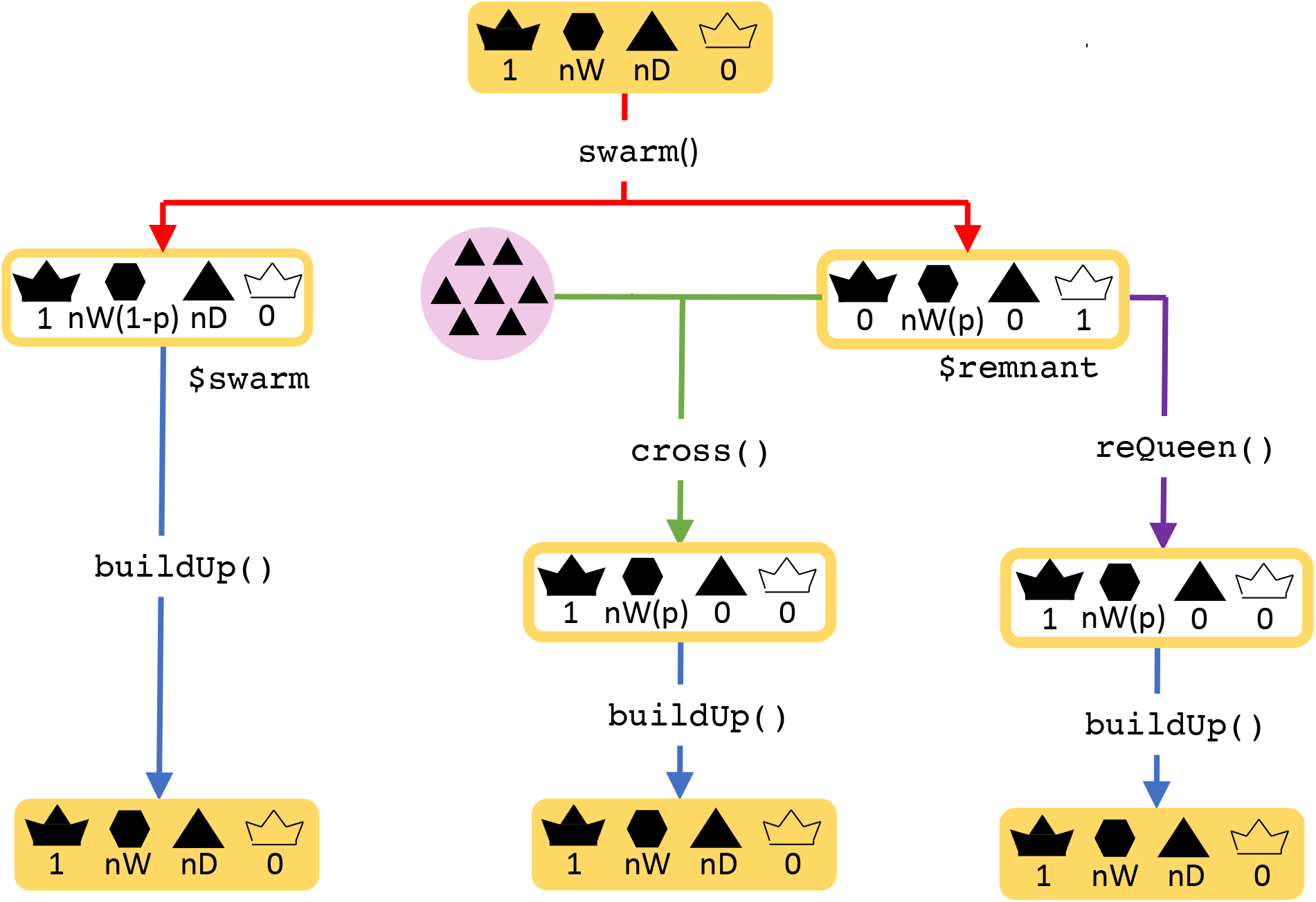
Swarm function.

#### Swarming a Colony

Let’s swarm our `colony`:

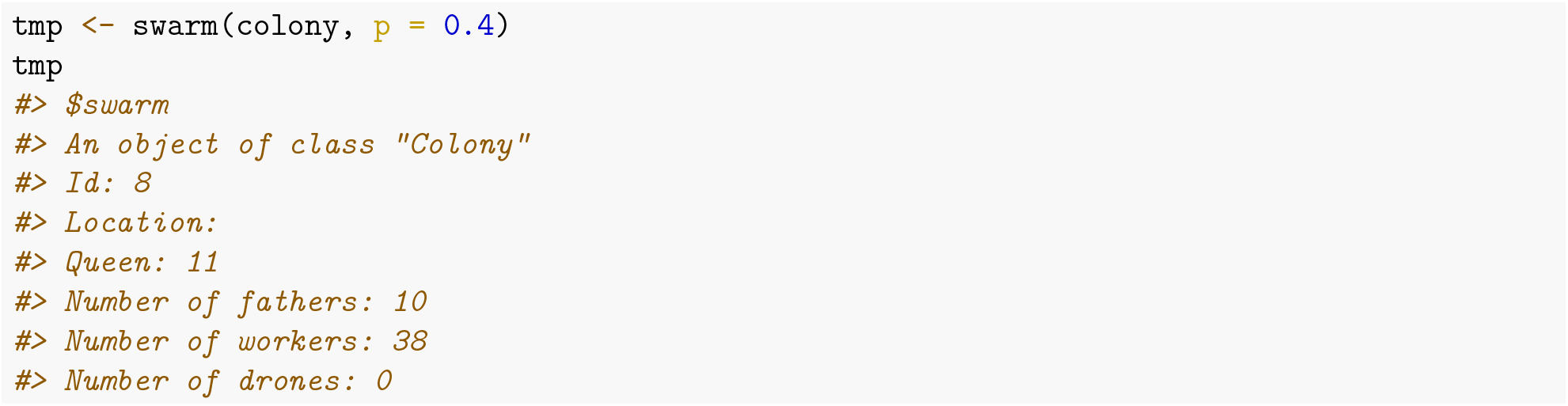

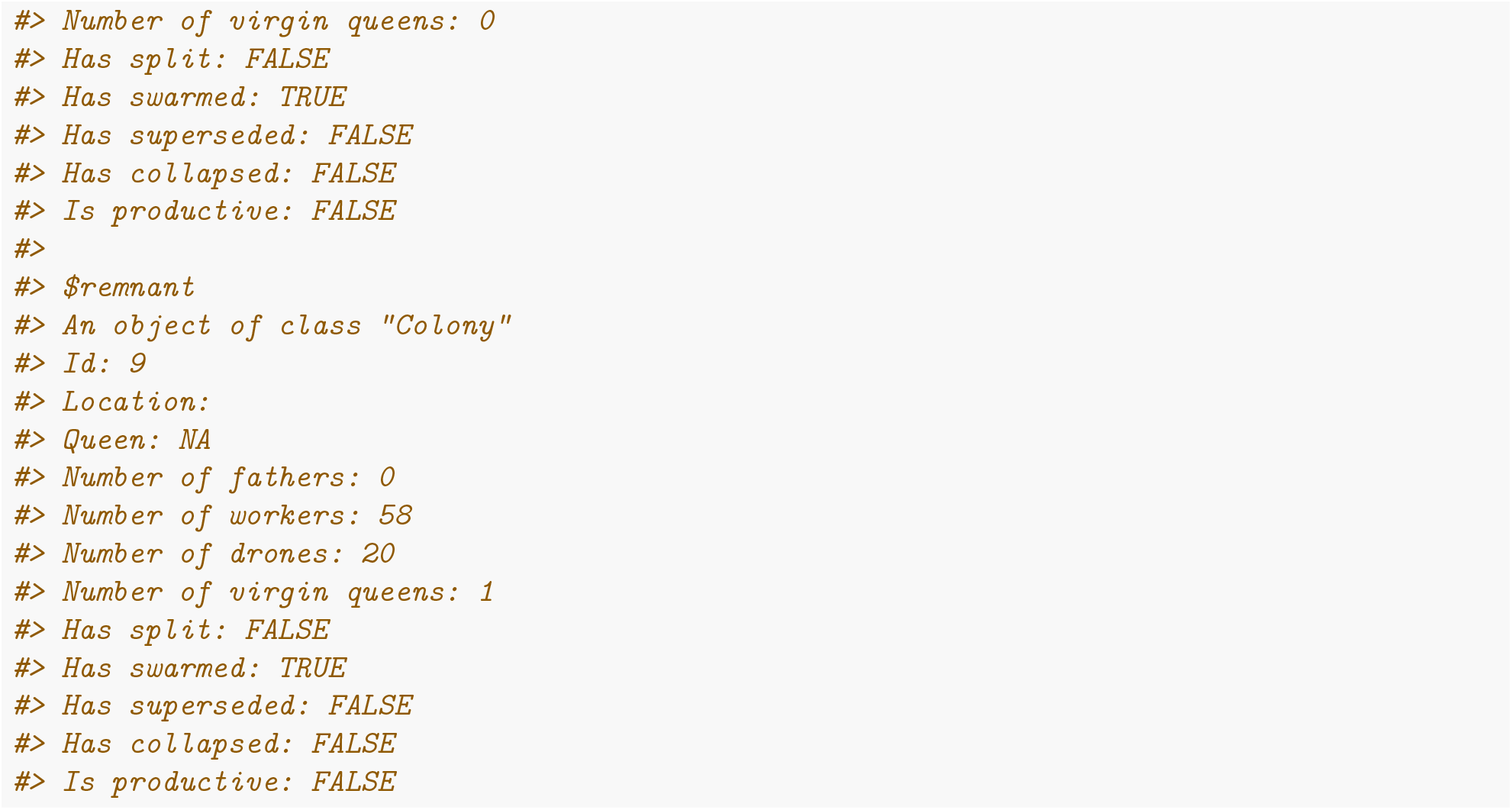

We see that the remnant colony does not have a queen but has one virgin queen that will have to be mated. It also has 60 workers since we set `p` argument to 0.4, meaning that 40% of workers left with the swarm. All the drones remained in the remnant. Note that the `swarm` status is turned to `TRUE`.

The swarm contains the old queen, no virgin queens, and 40 workers, since we set the proportion p to 0.4. Same as in the remnant, the `swarm` status is turned to `TRUE` in the swarm.

Swarming also turns the `production` slot to `FALSE` for both, the swarm and the remnant colony, since we would not be able to collect honey from such colonies (or other products).

The swarm stays genetically identical to the “old” colony, although downsized. Assuming that we have caught the swarm, we assign it back to the original colony object. The remnant has a new queen and is hence genetically different from the original colony. Thus, we assigned it to a new colony object.

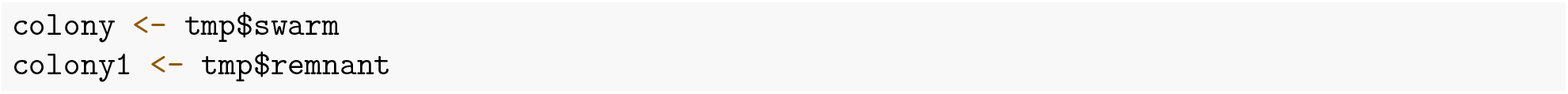

After swarming, the colony would usually build-up back to the full size and the virgin queens would mate. Building-up a colony turns the production status back to `TRUE`.

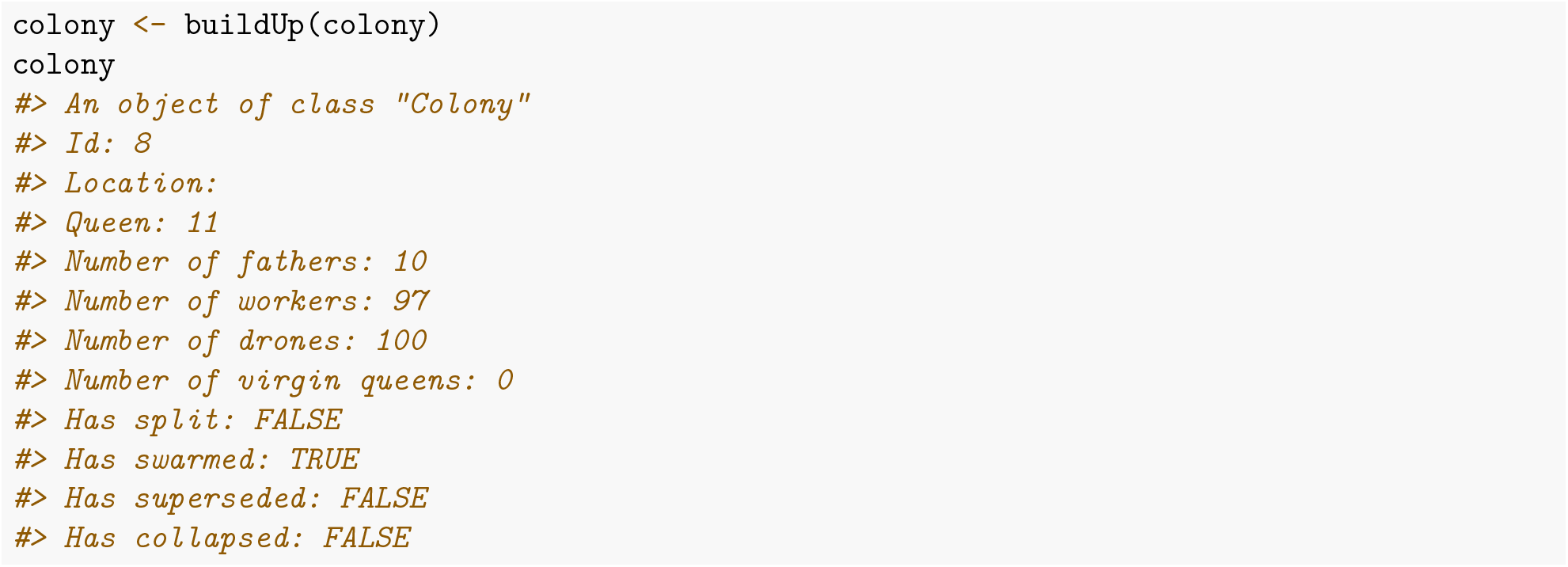

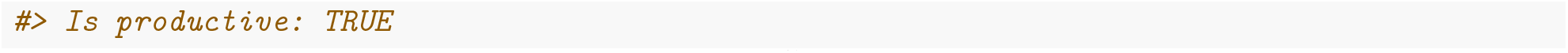

Instead of setting the `p` every time we call the `swarm()` function, we can save the `swarmP` argument in the `SimParamBee` object. The `swarm()` function will then use this percentage if `p` is not set.

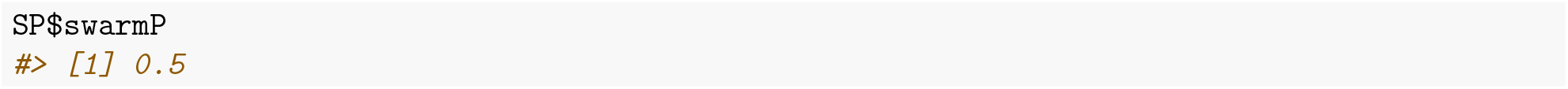

The default value is 0.5, but we can set any value we want.

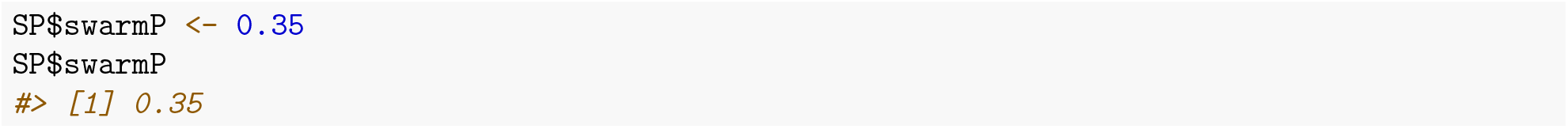

You can also use a non-fixed `p` parameter by using the function `swarmPUnif` that samples the `p` from a uniform distribution between values 0.4 and 0.6 irrespective of the colony strength. You can read more about this in the Sampling functions vignette.

#### Swarming a MultiColony

We swarm a `MultiColony` object in the same was we swarm a single `Colony` – with the `swarm()` function. The `swarm()` function is here applied to each colony in the `MultiColony` object with the same parameters. The function now returns a list with two `MultiColony` objects – one containing the swarms and the other containing the remnants.

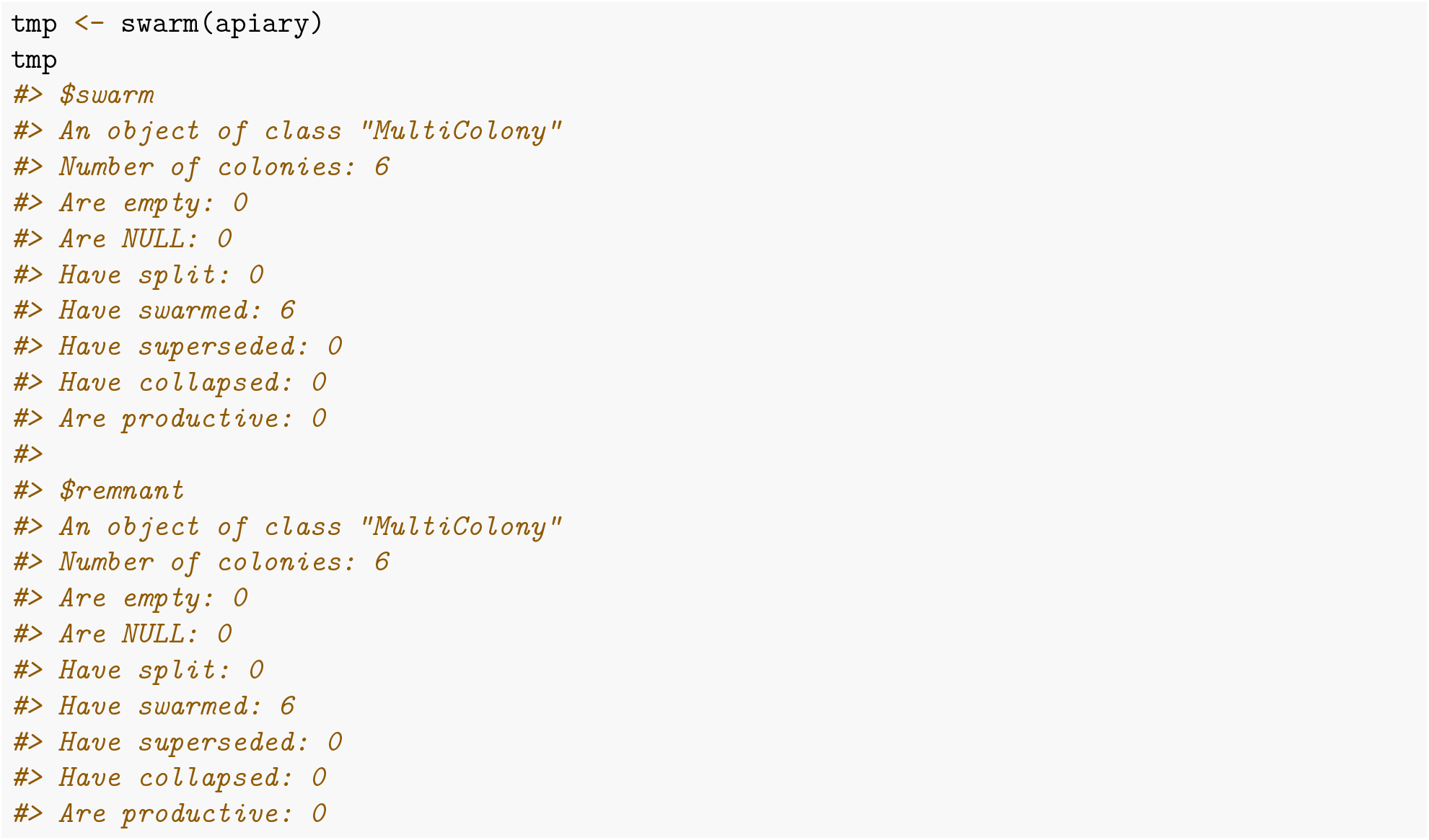

We see that we get six swarms and six remnants from the apiary with six colonies. We can inspect individuals colonies to ensure they swarmed according to the parameters. Let’s inspect the swarm and remnant of the third colony.

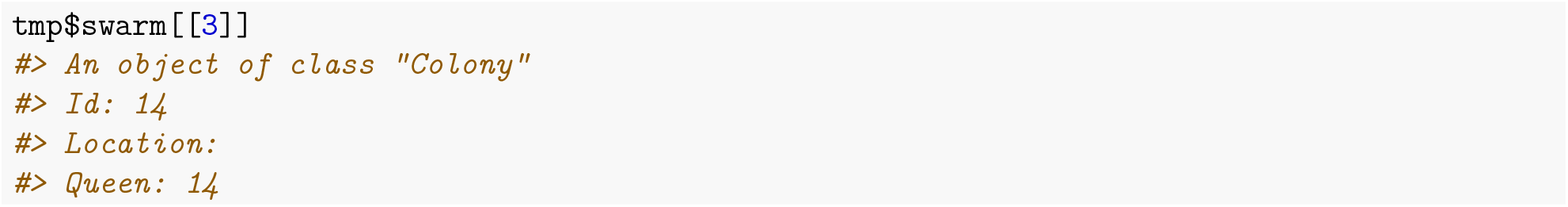

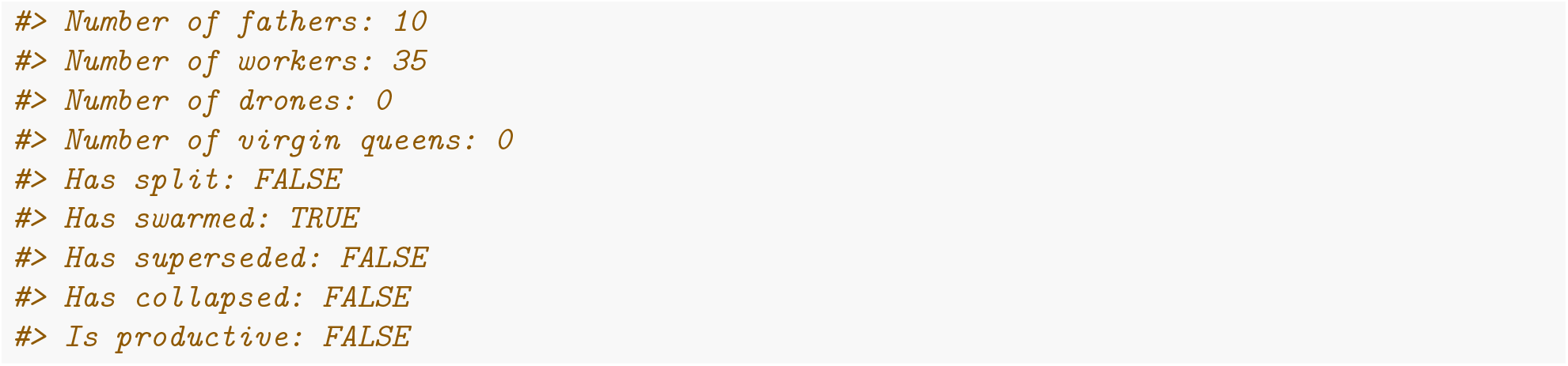

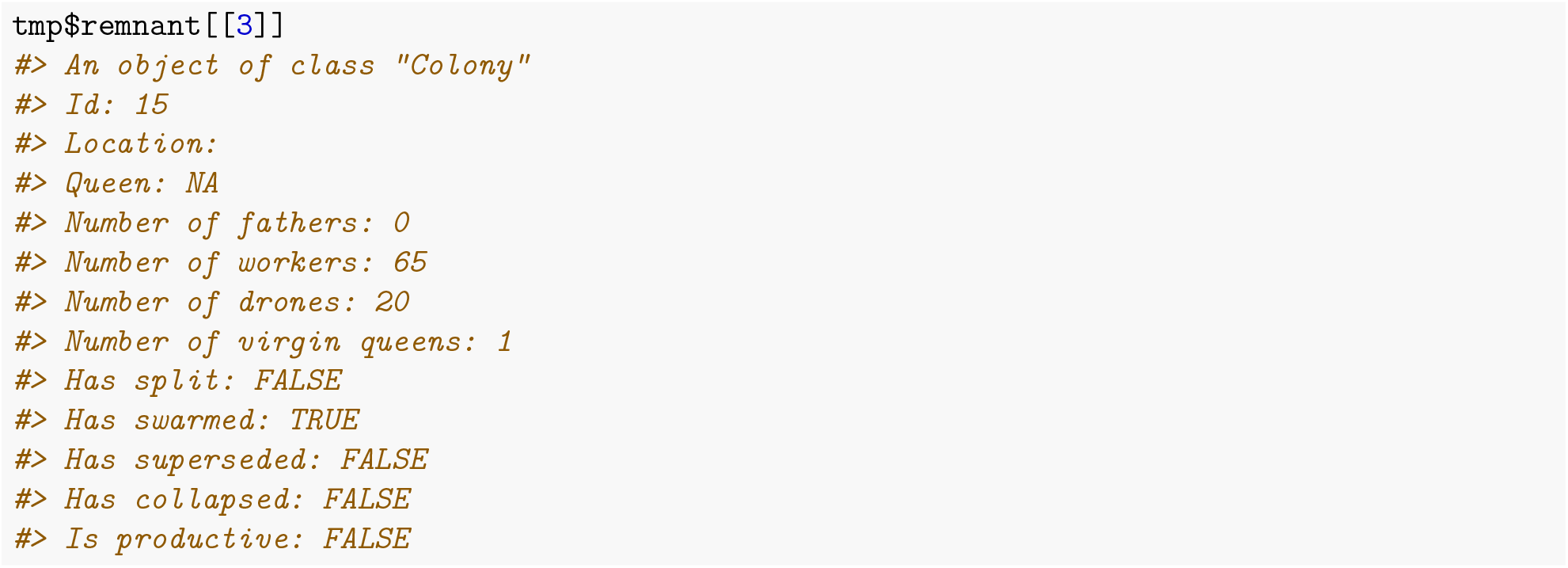

We see that the the third colony was swarmed with `p` of 35% as specified in the `SimParamBee`, hence the swarm left with 35 workers and the old queen, and the remnant stayed behind with a new virgin queen and 65 workers.

Above, all the colonies in a `MultiColony` are swarmed with the same percentage. However, we can also specify a different `p` for each colony.

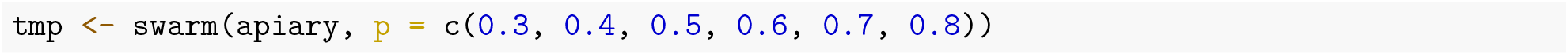

If we now inspect the first and the second swarm, we see that each colony has a different percentage of workers that stayed or left.

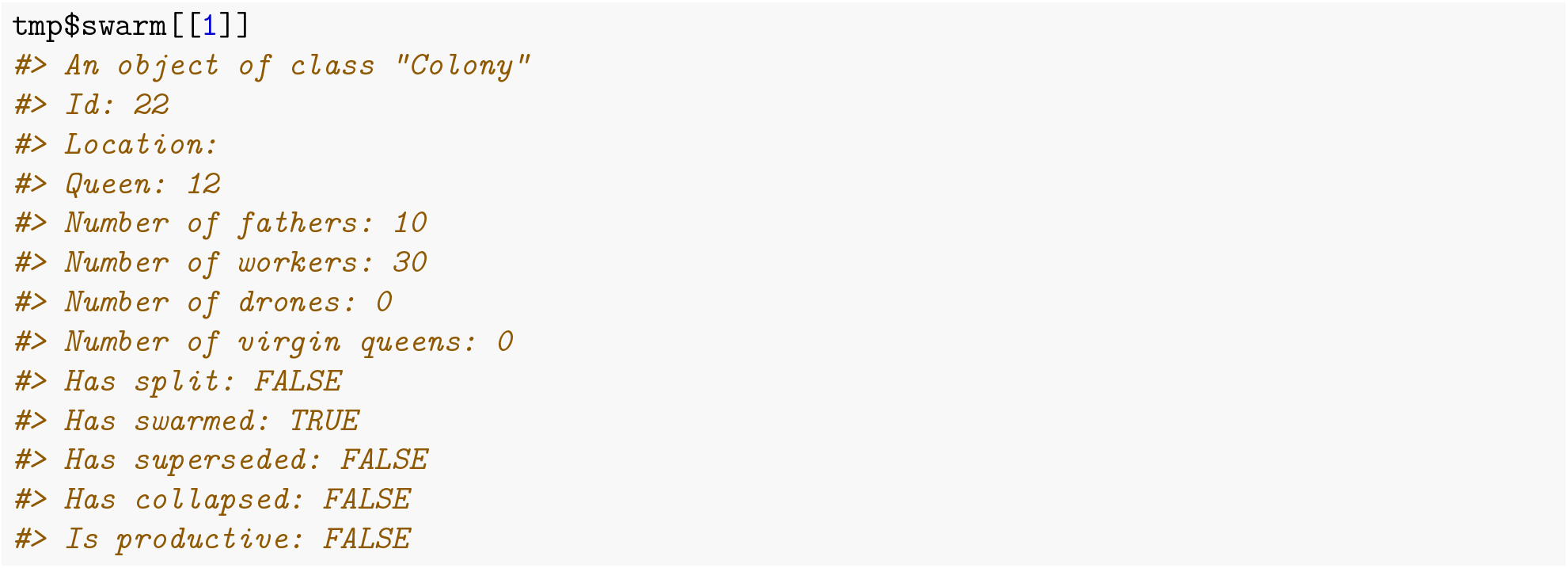

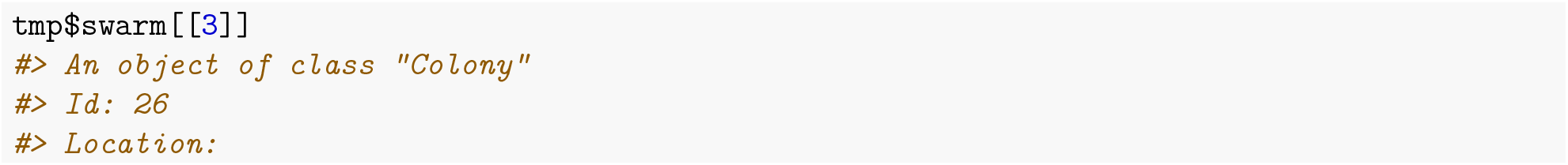

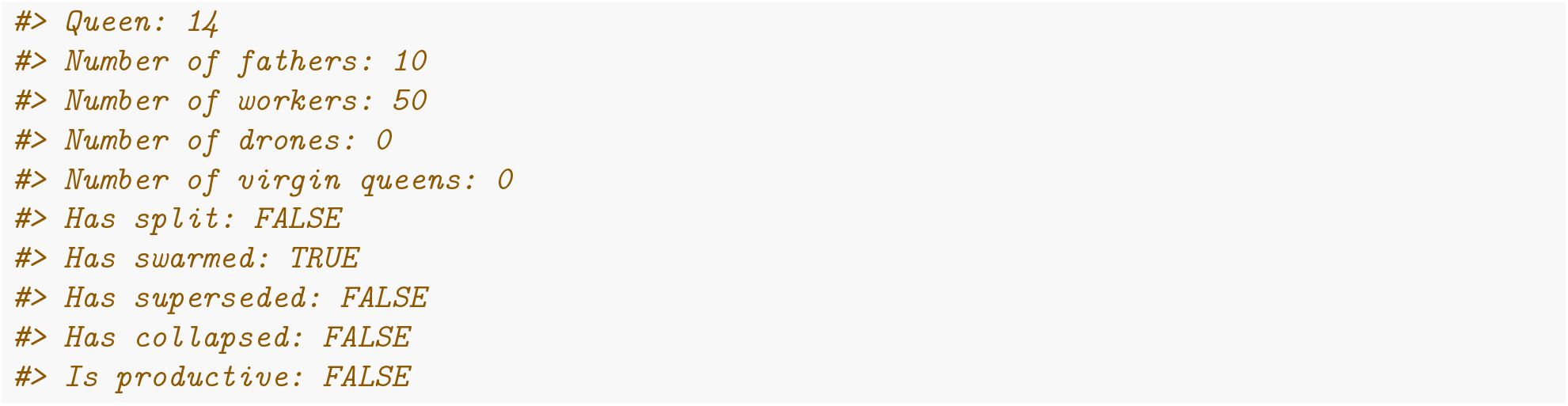

If you want to track the genetics, you might want to assign the swarms back to the original apiary and create a new apiary from the remnants. However, if you want to track the position, the remnant actually stay in the original position and would hence be assigned back to the same apiary, while the swarm would be assigned to a new apiary or even be lost. Here, we will track the genetics and assign the swarm back to the original apiary and remnants to a new apiary.

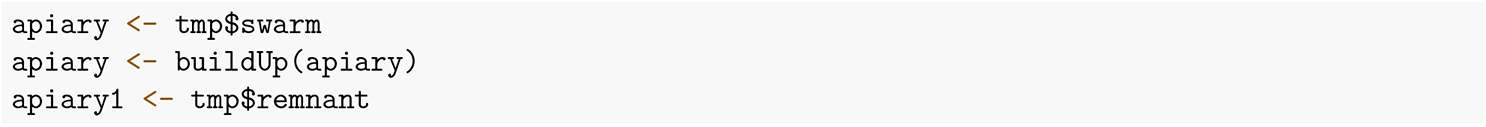

### Split

Colony splitting is a common beekeeping technique to limit swarming. A percentage of workers, brood, and food stores are split away to create a new colony or combine two split. Old queen normally stays in the original (remnant) colony. We created a function `split()` that works on `Colony` and `MultiColony` objects (Figure 2). It accepts the `p` argument as a proportion of workers that will be split away for a new colony. The output of the function is a list of two `Colony` or `MultiColony` objects: remnant that contains the old queen, drones, and (`1-p`) workers; and split that doesn’t contain a queen, but contains virgin queens and `p` workers. The `split()` function follows the same principles as the `swarm()`, hence we will limit explaining the outputs.

**Figure 2:**
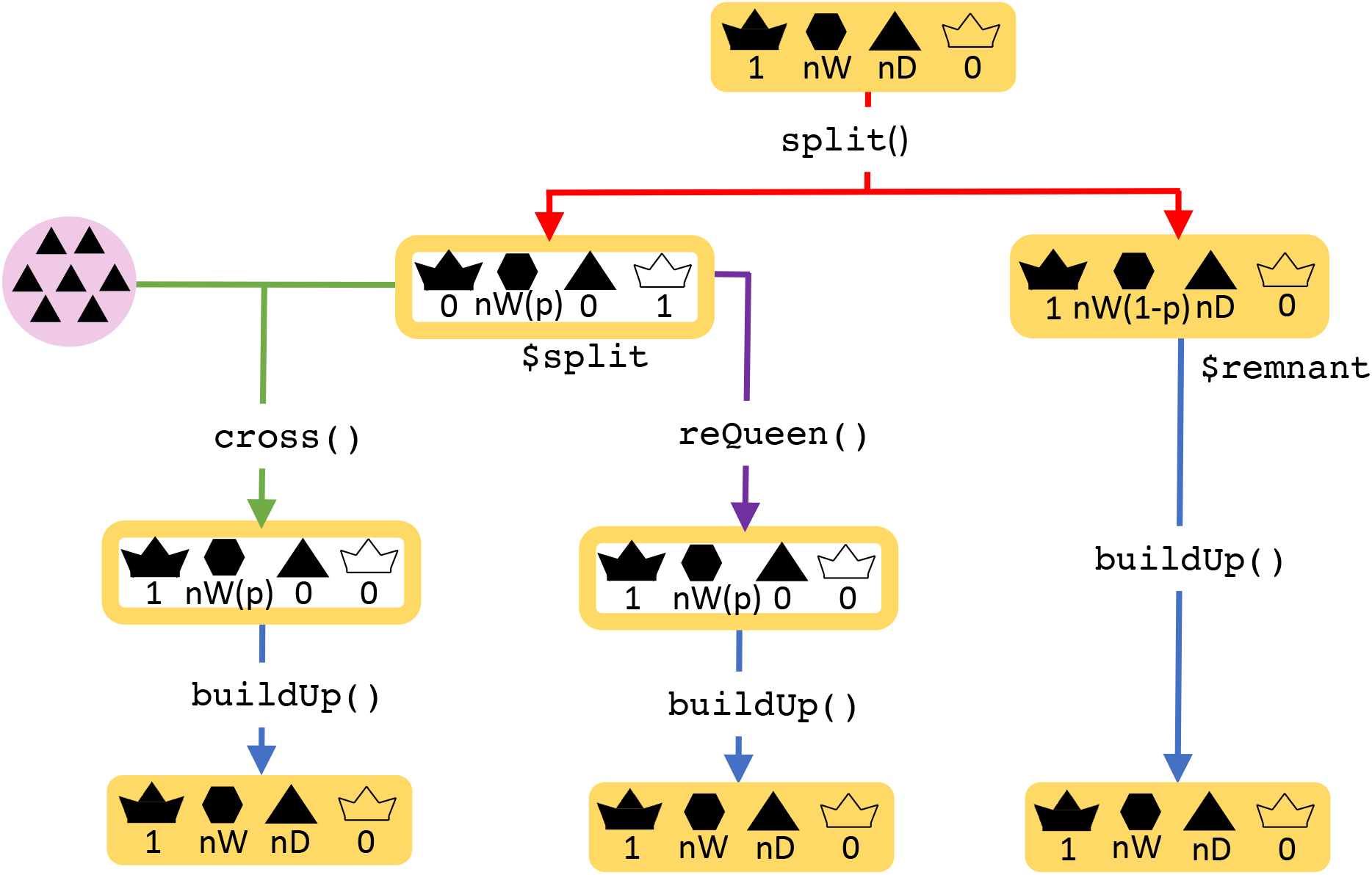
Split function

#### Splitting a Colony

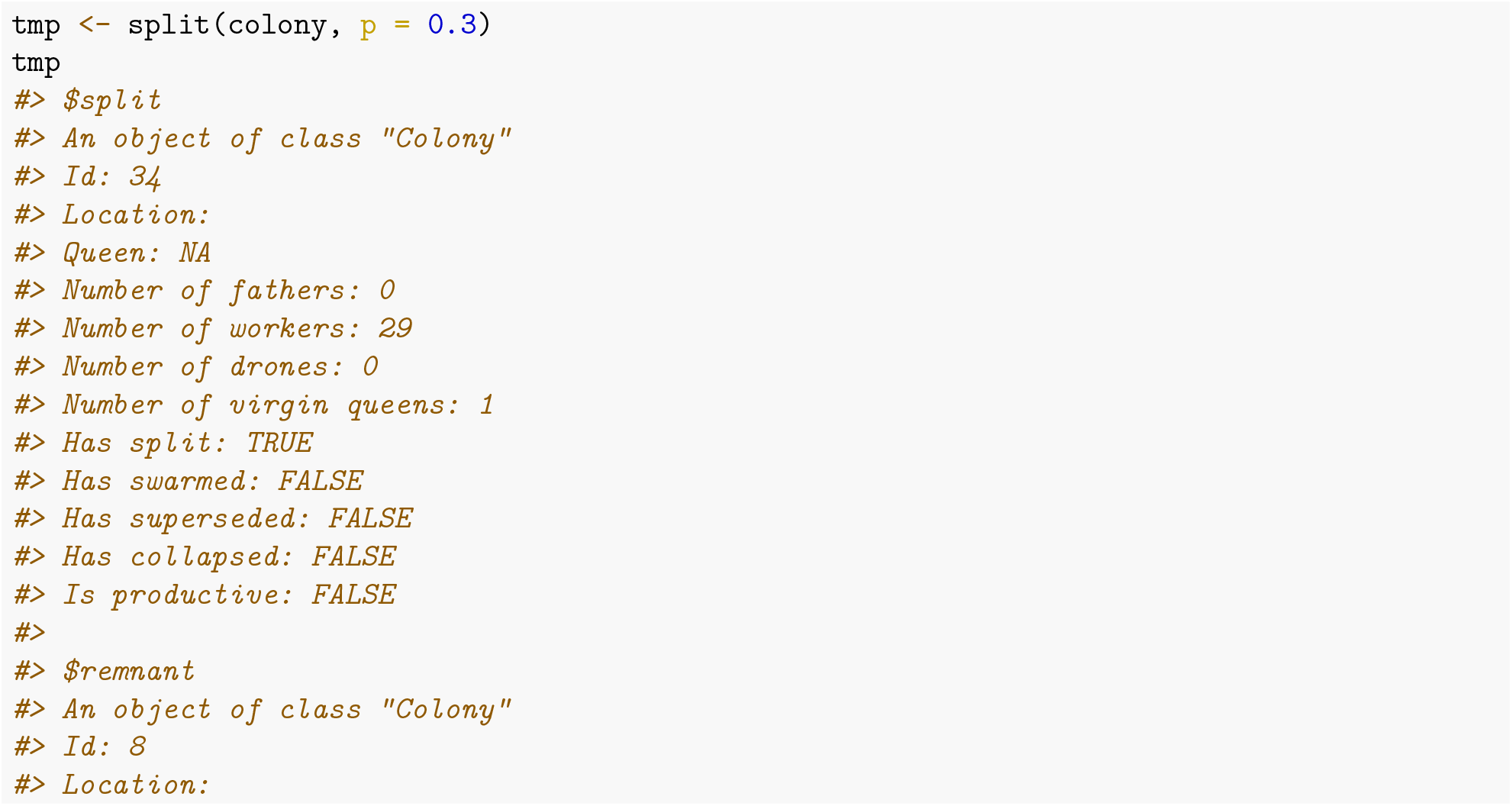

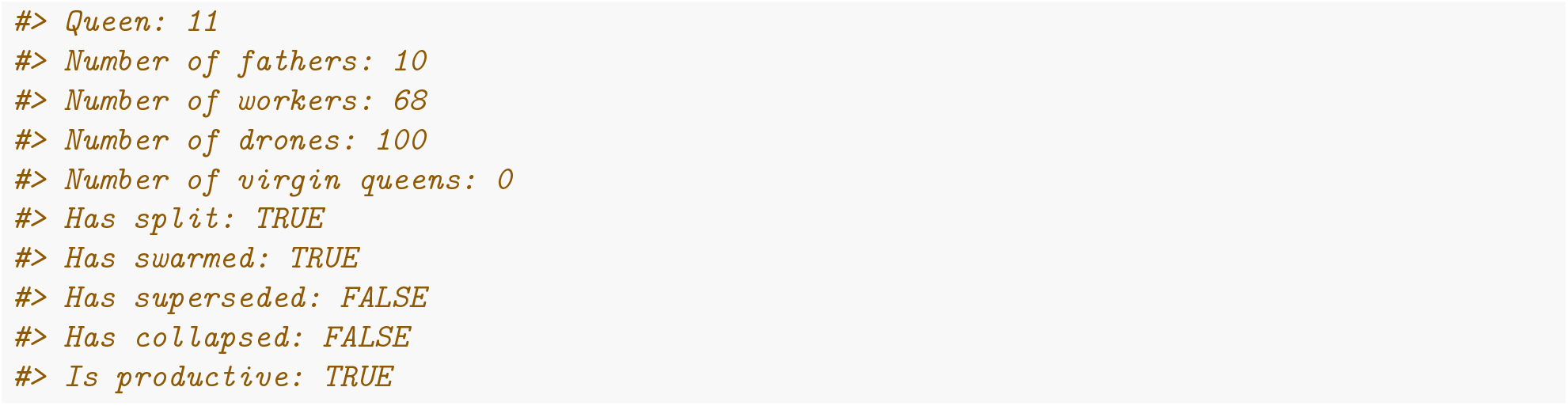

We see that remnant contains the old queen and 70% of workers since we set `p` argument to 0.3, meaning that 30% of workers are removed in the split. The `split` status of the remnant colony is turned to `TRUE`. The `production` status of the remnant stays `TRUE`, because in reality we always split a colony in a way that does not threaten the production of the remnant colony.

If we inspect the split, we see that it contains 30% of the workers, the `split` status is turned to `TRUE`, and the `production` status is turned to `FALSE`, since in reality, these small colonies would not be productive.

We would usually consider the remnant as the original colony, since it tracks its genetics and location.

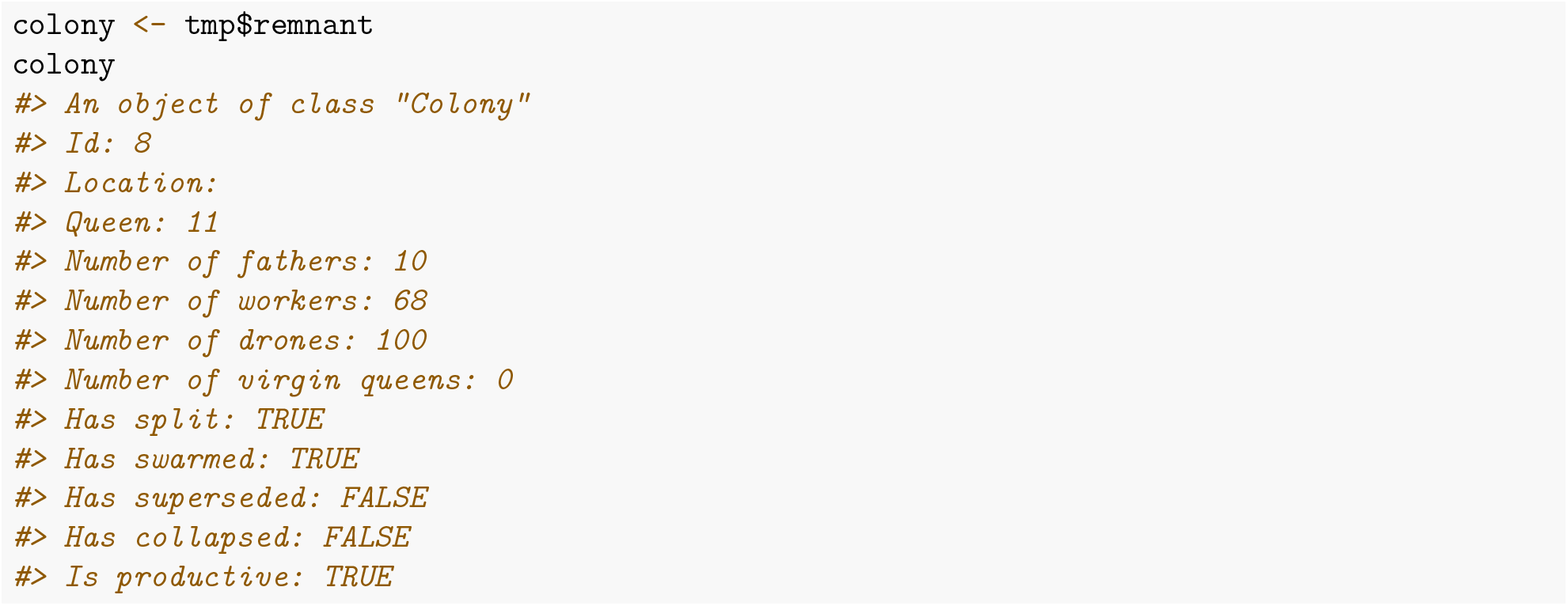

After a split, the colony would build-up back to their full-size.

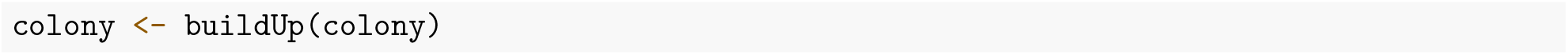

The `p` argument for splitting can be also saved in `SP` object, so we do not specify it each time we call the function – see `SimParamBee$splitP`.

#### Splitting a MultiColony

We split a `MultiColony` object in the same way we split a `Colony` (as shown in swarm).

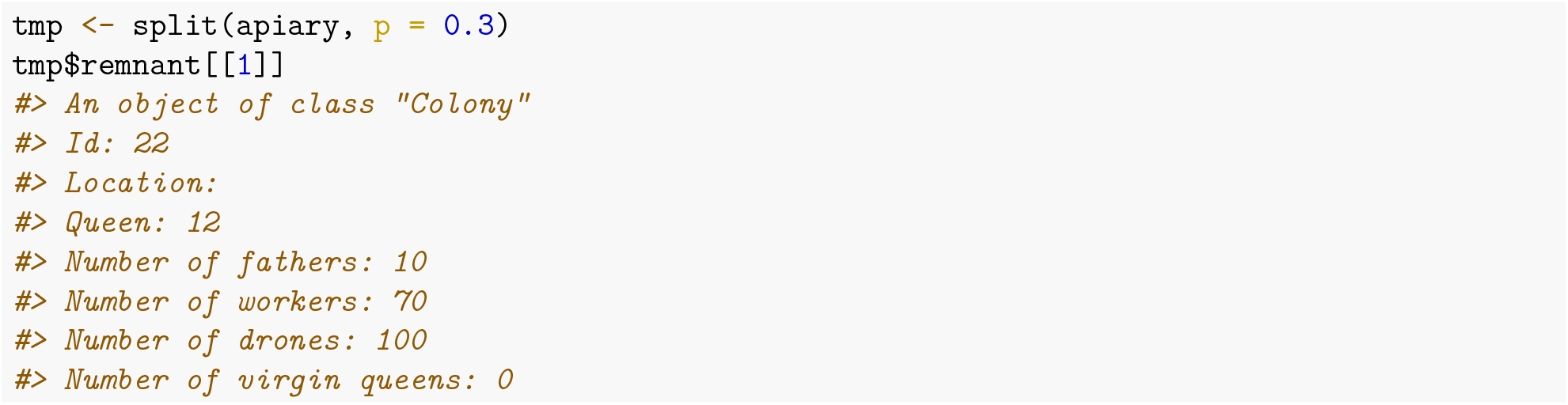

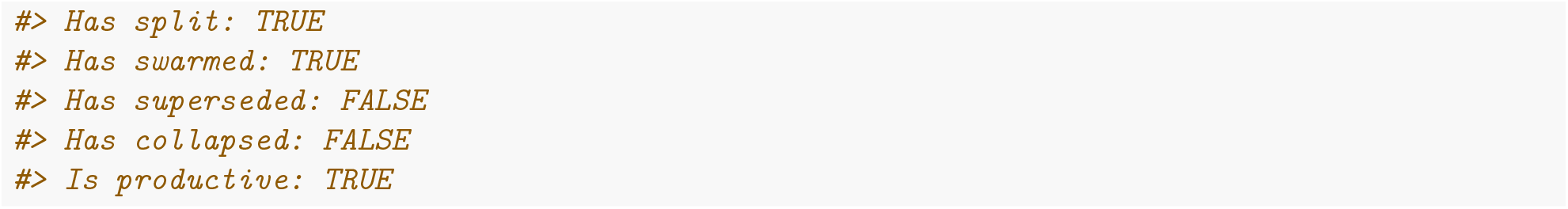

We again see that in remnant we have the queen and 70 workers:

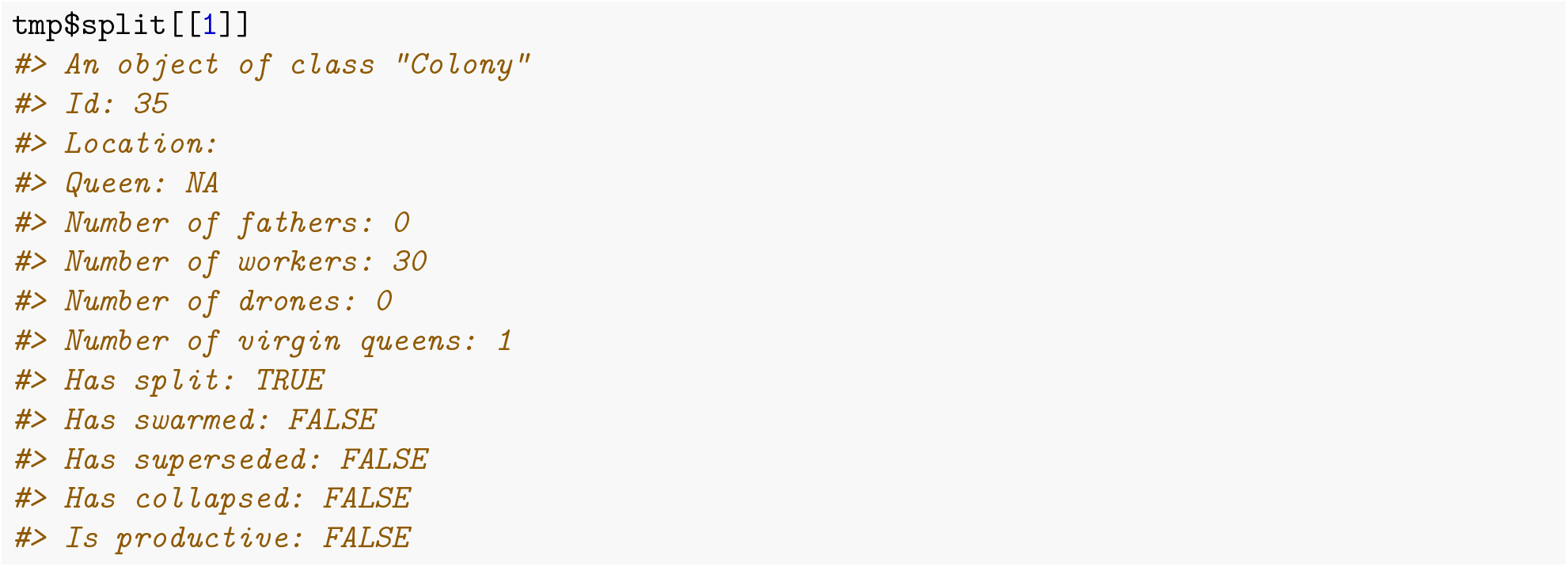

and a virgin queen with 30 workers in split. We can use the vector of `p`, different for each colony same as shown for the `swarm()` function above.

After the split, we would assign the remnant colonies back to the apiary and build them up.

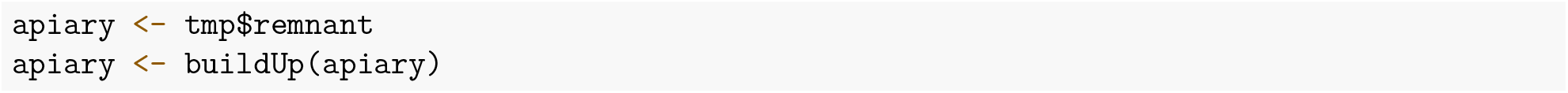

### Supersedure

Supersedure is a replacement of the queen by one of her daughters without interference of the beekeeper. Supersedure is a natural way of re-queening a colony without swarming. There are many reasons for supersedure: poor physical condition of a queen, old age, diseases, depleted spermatheca, poorly bread queen, reduced pheromone output and many others (Hamdan, 2010).

Function `supersede()` removes the old queen and triggers the creation of new virgin queens from the brood (Figure 3). The function returns a single `Colony` or `MultiColony` object (not a list of two).

**Figure 3:**
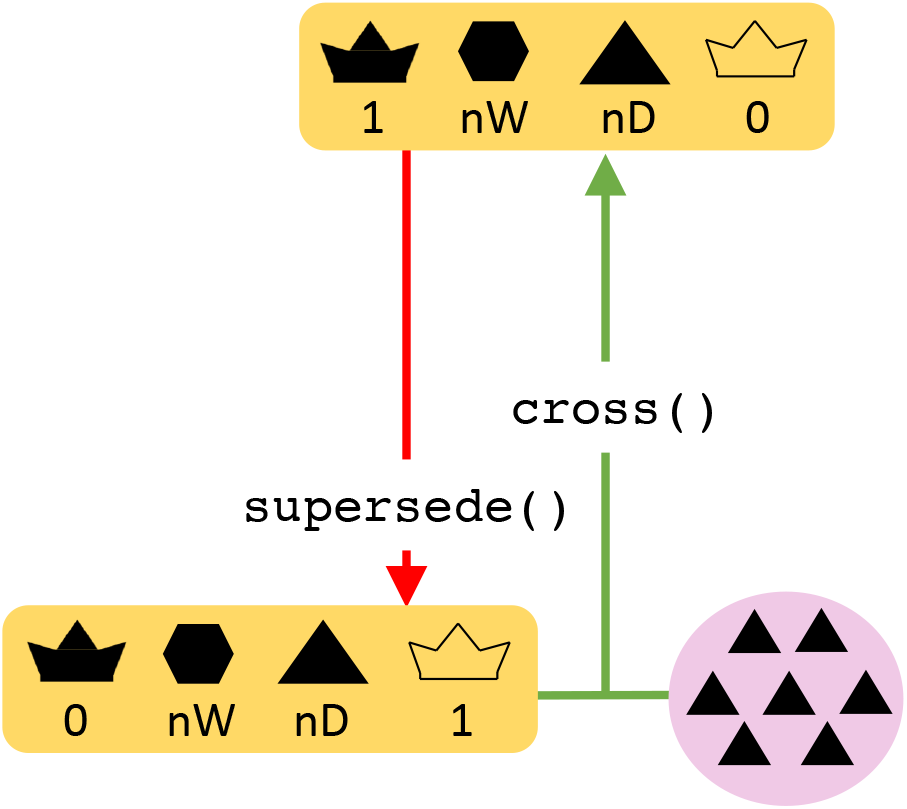
Supersede function

#### Superseding a Colony

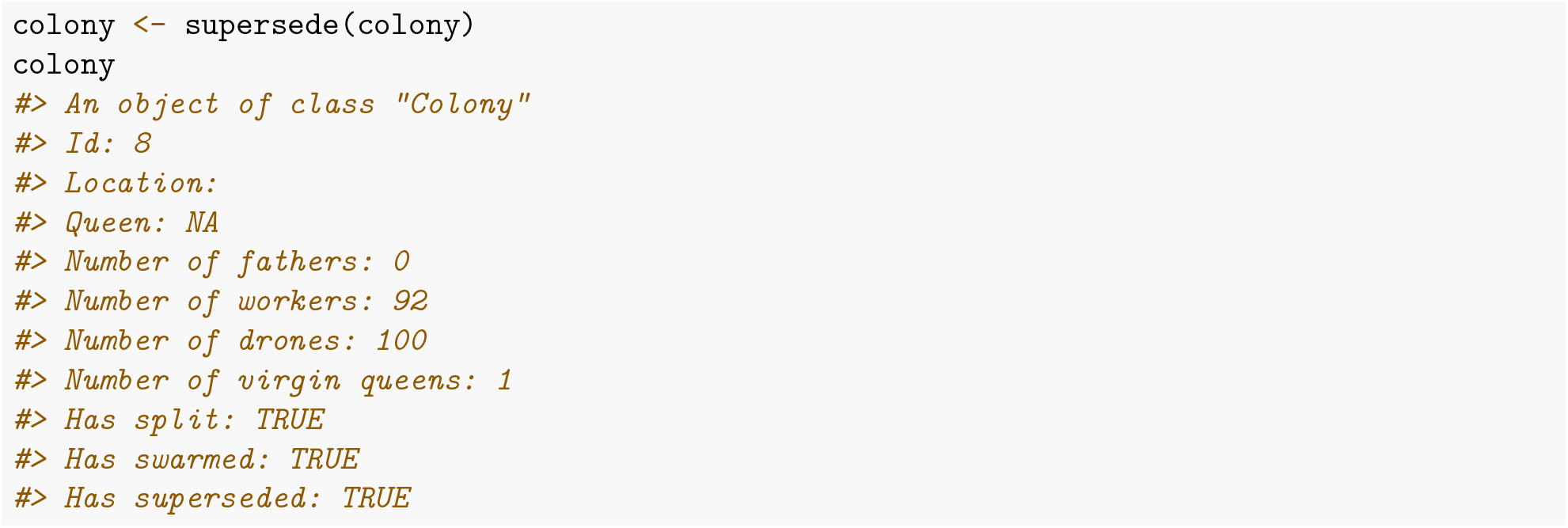

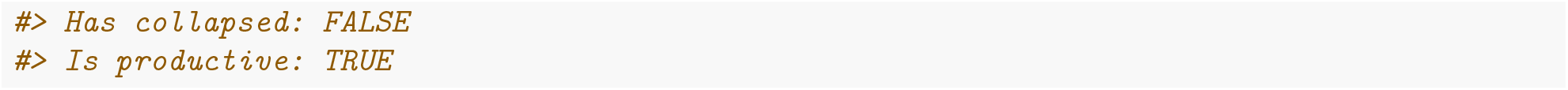

We see that after a supersedure, the old queen is removed and a new virgin queen is ready to mate. Hence, the next step in the simulation would be to cross this virgin queen. We also see that the `supersedure` status is set to `TRUE`. The `production` status stays set to `TRUE`, since the colony did not loose any individuals (it stayed at it’s full-size).

#### Superseding a MultiColony

The function `supersede()` works both on `Colony` and `MultiColony` classes. We supersede a `MultiColony` in a same way as a `Colony`.

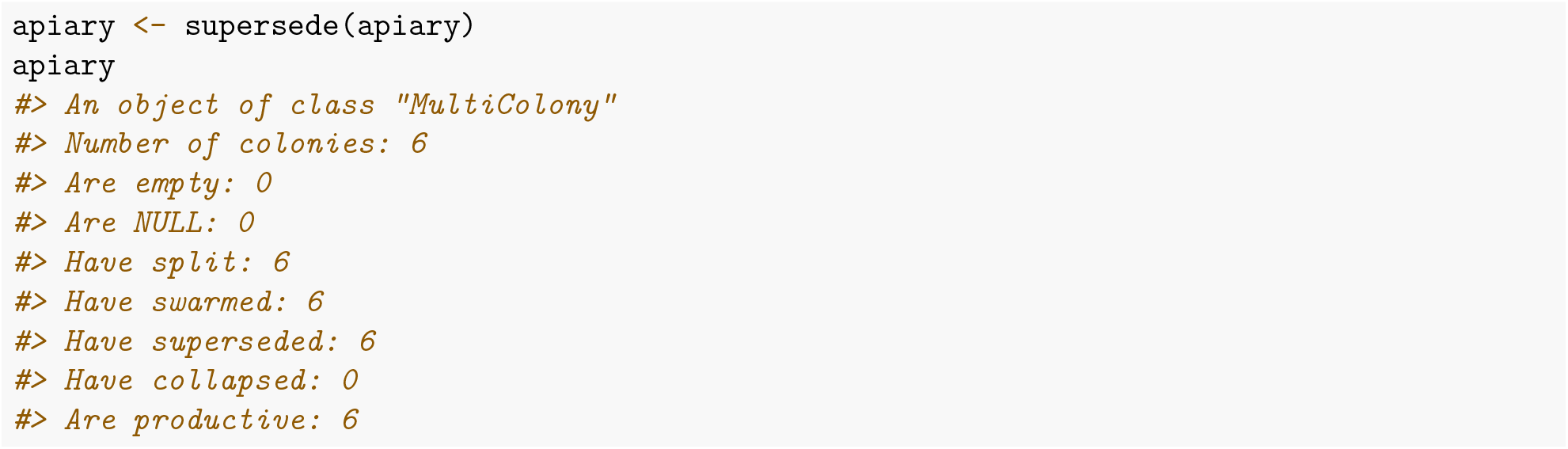

### Collapse

Collapse of the colony is a term that describes the death of a colony – when all individuals within a colony die. There are many reasons for the collapse of a honey bee colony: diseases, starvation, queen problems, contamination with pesticides, etc. Colony losses can be high, possibly up to 60% of colonies per year. High colony losses can significantly influence genetic structure of a population and hence genetic diversity and genetic gain.

Function `collapse()` simulates the collapse of a colony (Figure 4). This function does not remove individuals from the colony, but sets the `collapse` status of the colony to `TRUE` and `production` status to `FALSE`. This is to mimic reality, where bees would still be present in the colony, although being dead. This allows us to extract any genetic material from the colony even after collapse, say to study genetic causes related to the collapse. Future operations in terms of reproduction or simulation of events are not allowed with a collapsed colony.

**Figure 4:**
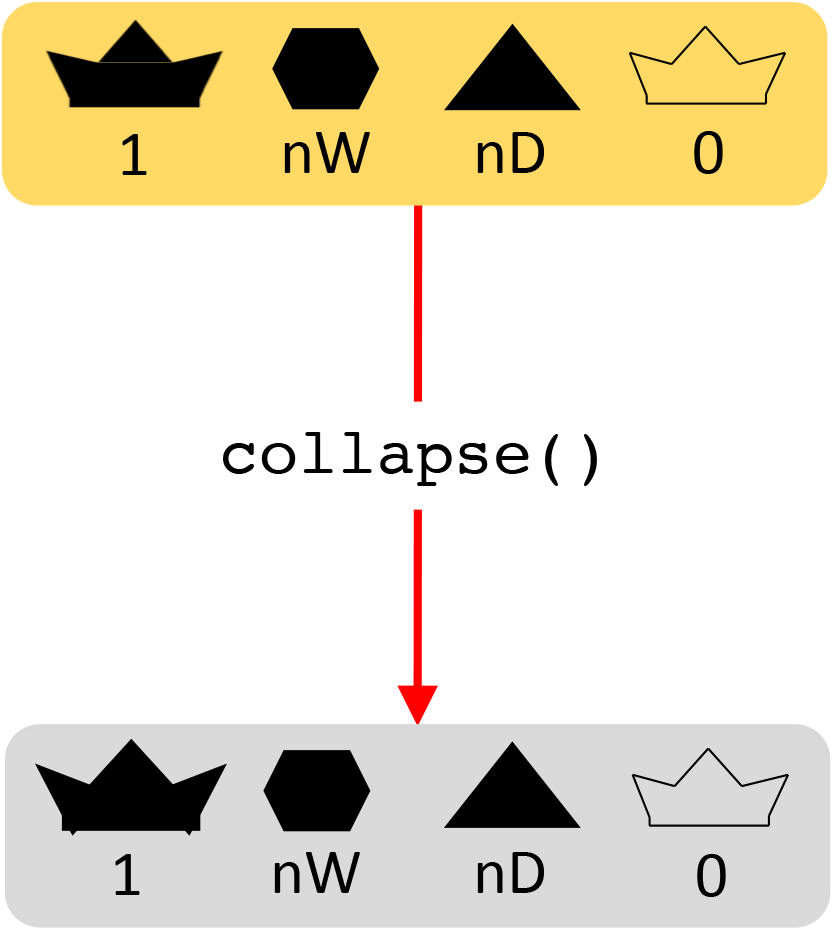
Collapse function

#### Collapsing a Colony

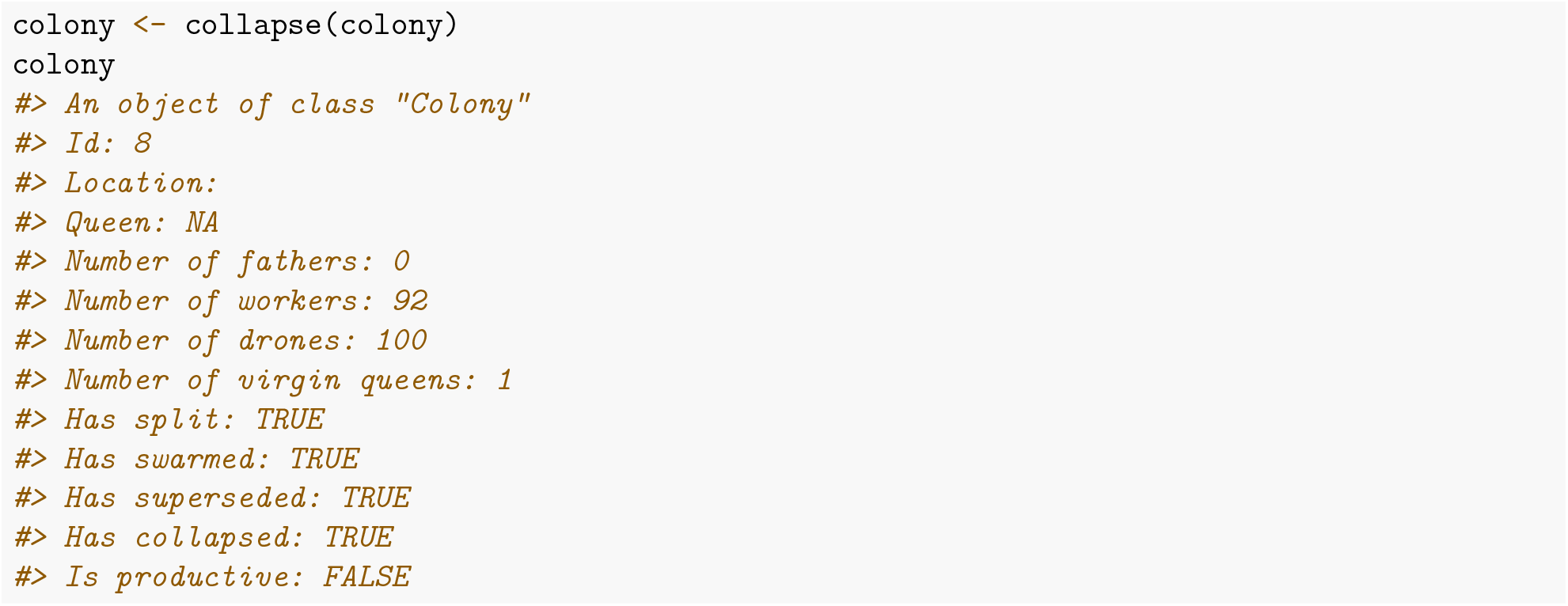

#### Collapsing a MultiColony

`collapse()` function is used when you want to keep collapsed colonies for subsequent analyses. If you don’t need the collapsed colony, you can also simply select the surviving colonies with the `selectColonies()` function.

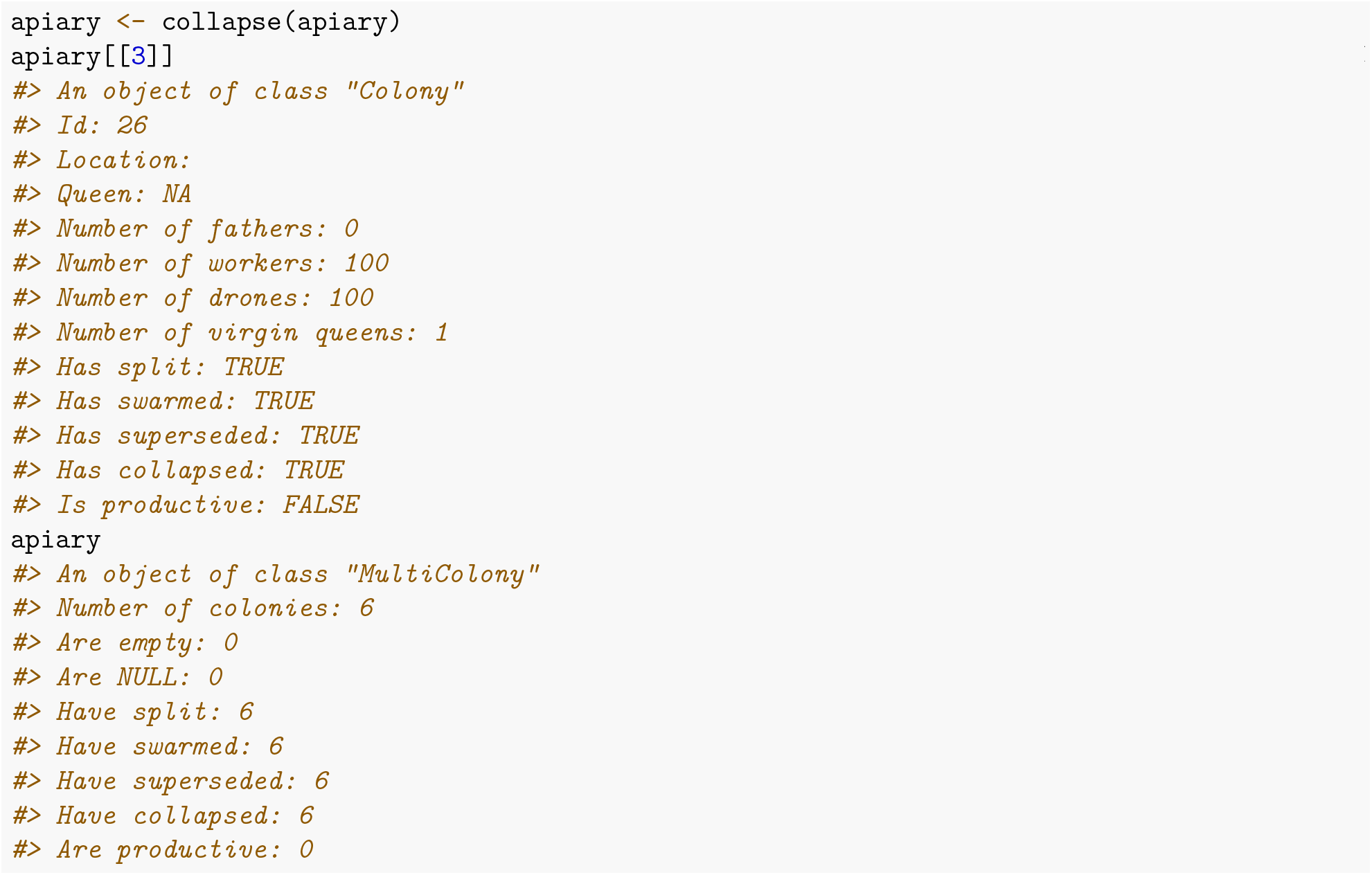

## Additional file 4 – Crossing vignette

### Introduction

This vignette shows how you can cross virgin queens in SIMplyBee. Here, we present how you can cross:

- single or multiple virgin queens (class `Pop`), virgin queen in a colony (class `Colony`), or all the virgin queens in an apiary or population (class `MultiColony`);
- cross either with pre-selected population of drones or according to a cross plan, and
- cross queens at an open drone congregation area (DCA) or at a mating station.

Start by loading the package:

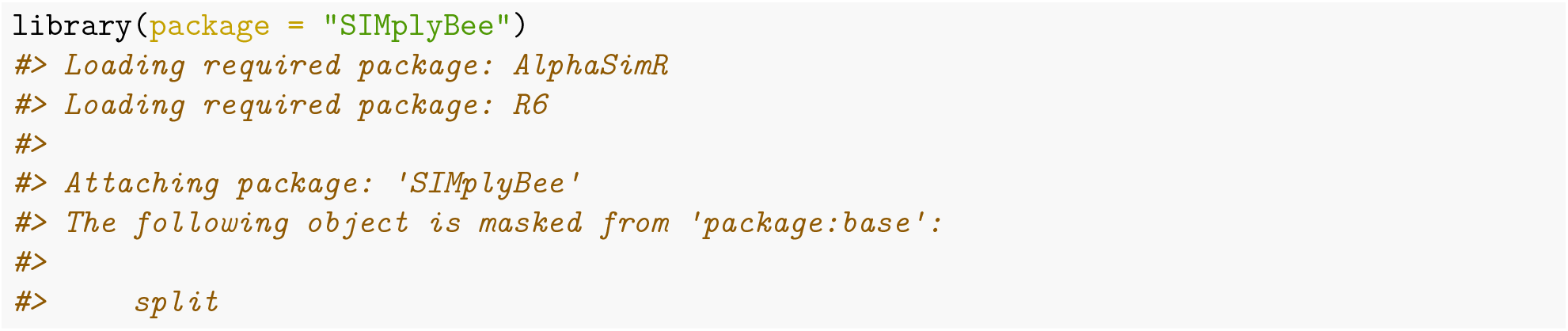

First, we create a founder population and some virgin queen, virgin colonies, and virgin apiaries that we will later cross.

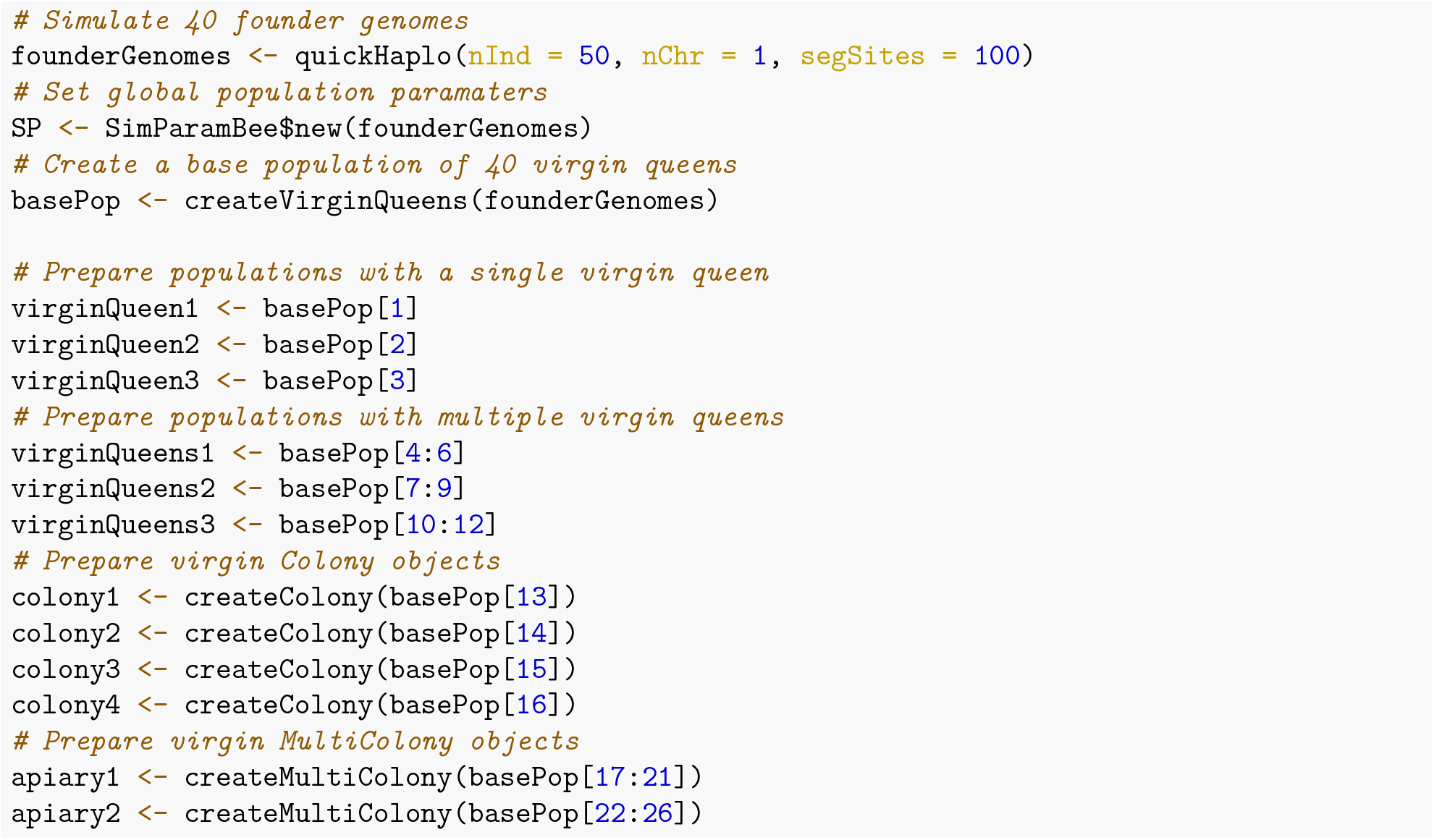

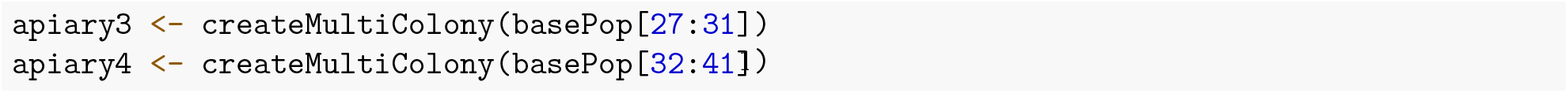

We will now create a groups of drones from the remaining queens with 1,000 drones per queen that will represent a drone congregation area (DCA).

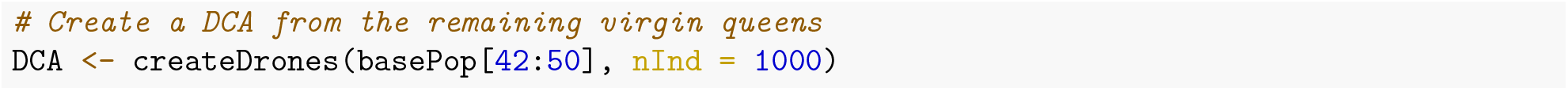

### Cross virgin queens at an open DCA

#### Cross by pre-selecting drone populations

We start by crossing our populations and colonies to pre-selected populations of drones. We pre-select the groups by pulling a desired number of drone packages from a DCA with the function `pullDroneGroupsFromDCA()`. This function requires you to specify a group of drones (`DCA`), how many groups you want to pull from the DCA (`n`), and how many drones per group you want (`nDrones`). For `nDrones`, you can either specify an integer or a sampling function, which results in a different number of drones in each of the pulled groups (you can read more about this in the Sampling functions vignette). These sampling functions are particularly useful in crossing simulations:

- `nFathersPoisson()`: samples the number of drones from a Poisson distribution with a default mean of 15 (the user can specify a different mean) – the output can contain the value 0 and
- `nFathersTruncPoisson()`: samples the number of drones from a zero truncated Poisson distribution with a default mean of 15 (the user can specif a different mean) – the output does not contain the value 0.

If these functions do not satisfy your needs, you can specify your own sampling function(s).

We can pull the drone groups out separately for each crossing or pull them out all at once.

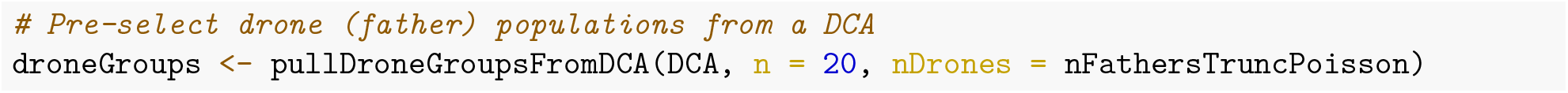

Once we inspect these drone groups, we see there is a different number of drones in each group because we used a sampling function:

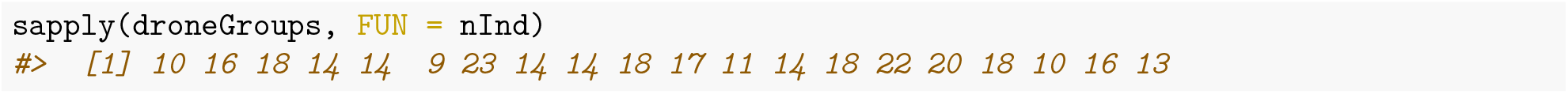

Now, we can cross our virgin queens to drone groups.

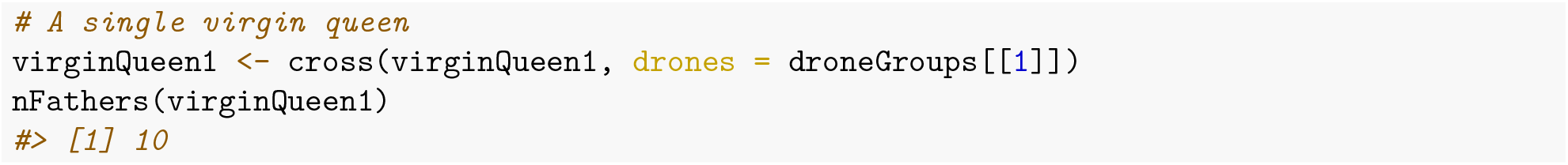

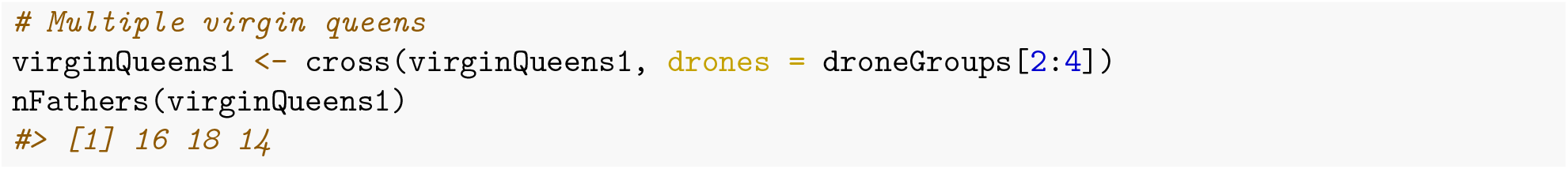

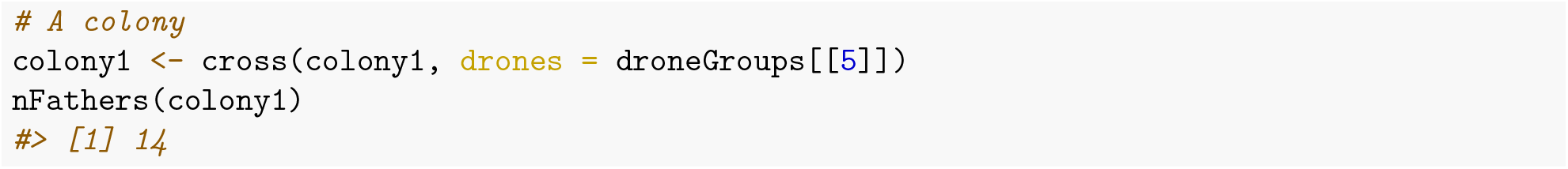

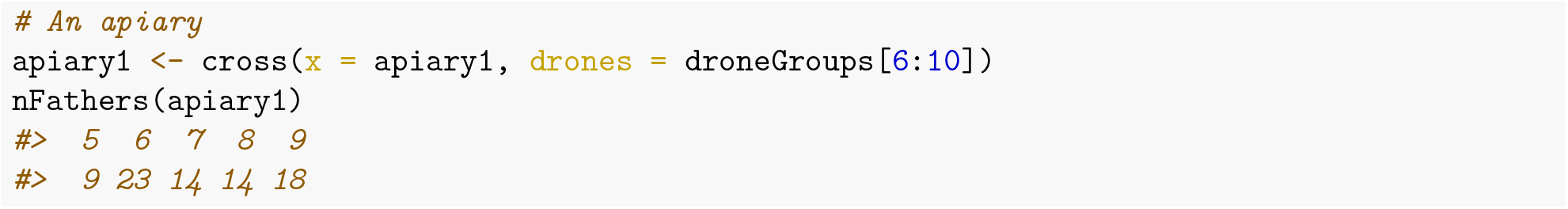

### Cross according to a cross plan

Another option is to provide a cross plan with IDs of the virgin queens/colonies and drones, and a single drone population with all the drones listed in the cross plan. You can create a cross plan with the function `createRandomCrossPlan()`. This function creates a cross plan by randomly sampling a desired number of drone IDs from a DCA and assigning them to either virgin queen ID or colony ID. When crossing a virgin queen from a colony, you have to provide the colony ID, since there could be multiple virgin queens within the colony. In that case, the random selection of one virgin queen occurs within the `cross()` function. To create a cross plan you therefore have to provide the IDs of either the virgin queens or the colonies you want to cross (but not both in the same cross plan!!!), the drone population, and the number of drones you want to cross to a particular virgin queen. This can again be a fixed number or a sampling function. We can create a separate cross plan for each mating or create one combined cross plan for multiple matings (but can not have virgin queen and colony IDs in the cross plan at the same time!!!). Here, we again mate a single virgin queen, a population of virgin queens, virgin queens from a colony, and from an apiary.

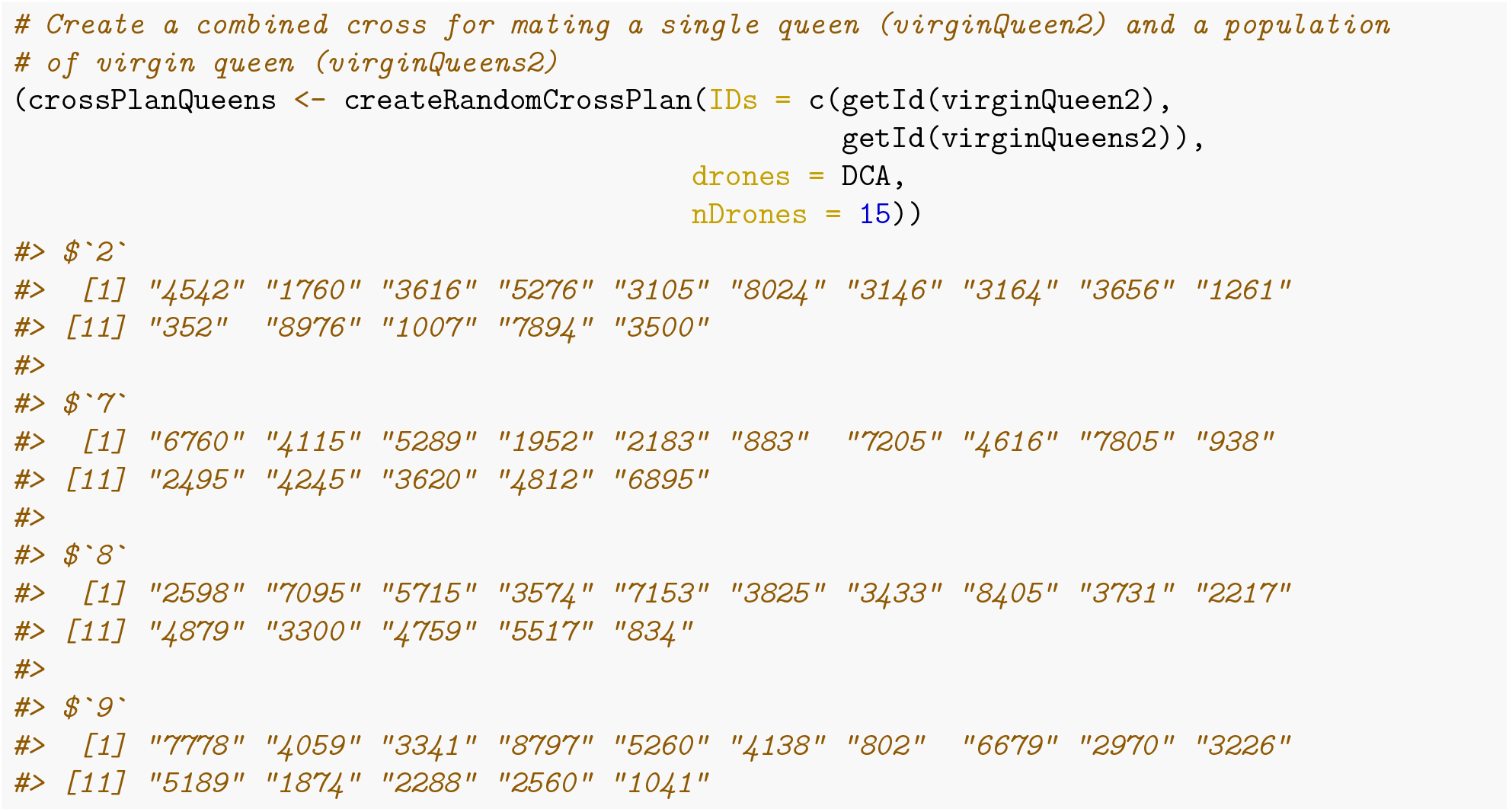

We see that the created cross plan is a list with IDs corresponding to the virgin queen’s IDs and the elements of each list being the IDs of the drones the virgin queen will mate with. Now we can cross these virgin queens by providing the `crossPlanQueens` to the `cross()` functions.

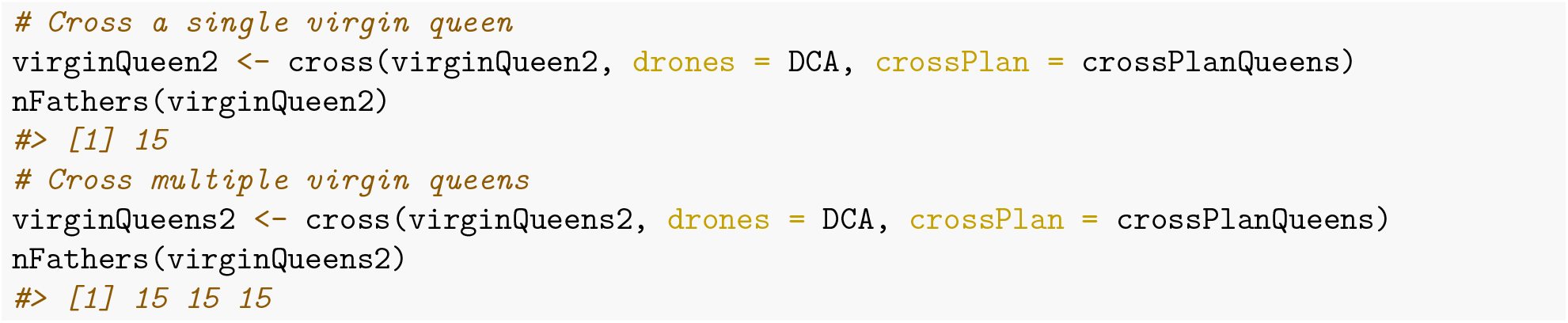

As already mentioned, we need to create a separate plan for crossing virgin queens already in colonies, since here we need to provide colony IDs.

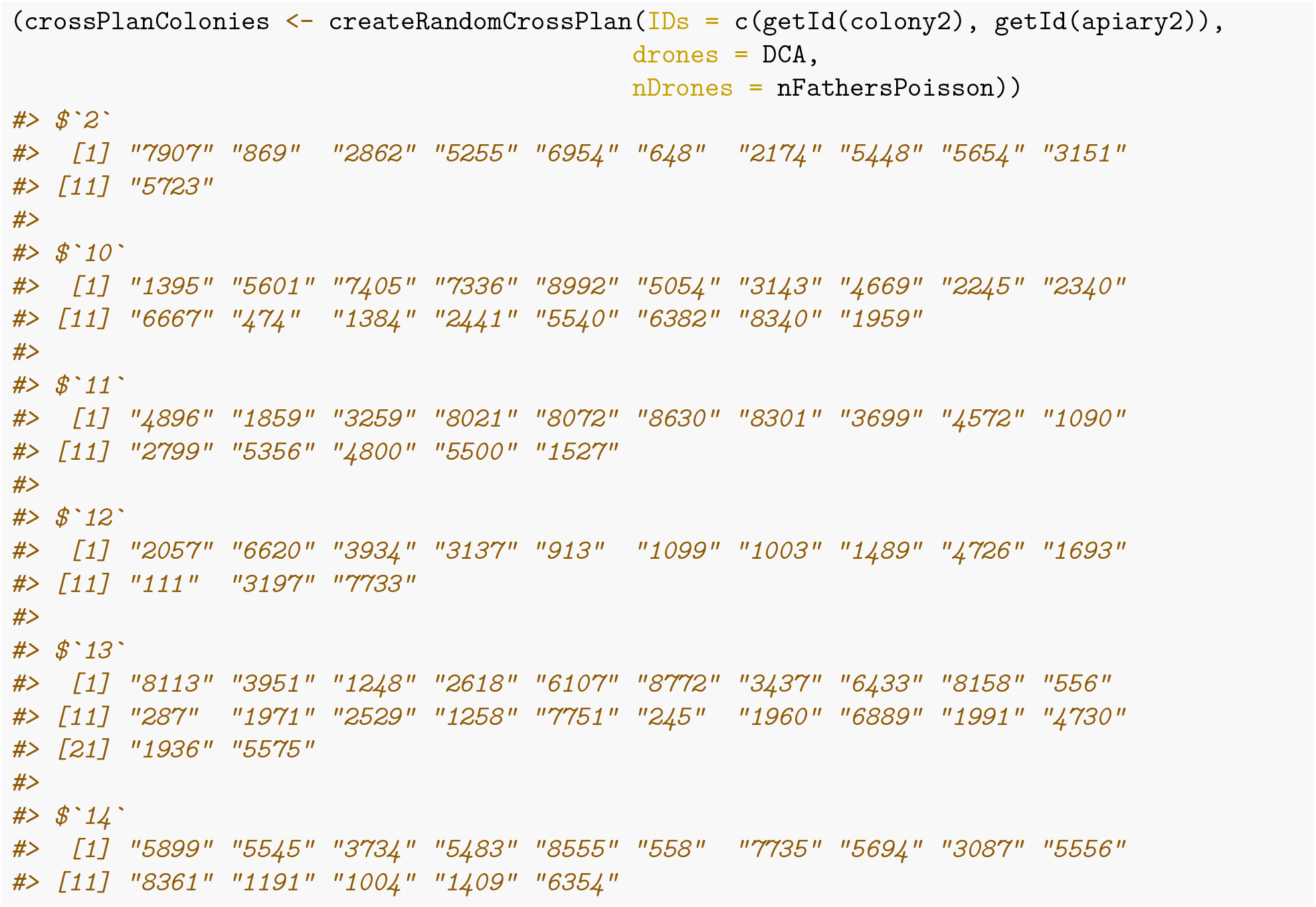

We again see that the cross plan is a list, now with colony IDs and elements of the list being the ID of drones the virgin queen within each colony will mate with. Now, we can cross our “colonies” by providing the `crossPlanColonies` to the `cross()` function.

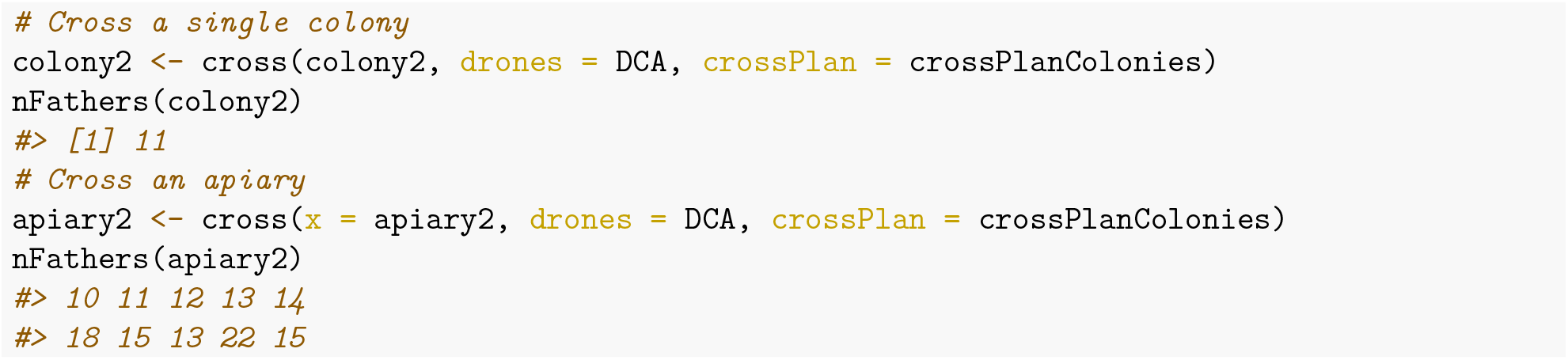

### Cross virgin queens at a mating station

Mating virgin queens at a mating station is no different than mating them at an open DCA – the difference is in the DCA itself. In the case of open mating, the DCA consists of drones from multiple queens, all of which are usually unknown. In the case of a mating station, the DCA consists of drones coming from a sister group of drone producing queens (DPQ), the queen of which is known. This allows us to track the pedigree on the paternal side.

To simulate this situation, we first create a mating station DCA using `createMatingStationDCA()` function, which takes a single “sire” colony (queen of the DPQs). From the “sire” colony, the function first produces a desired number of sister DPQs, and next produces a desired number of drones per DPQ. The produced drones represent the mating station’s DCA.

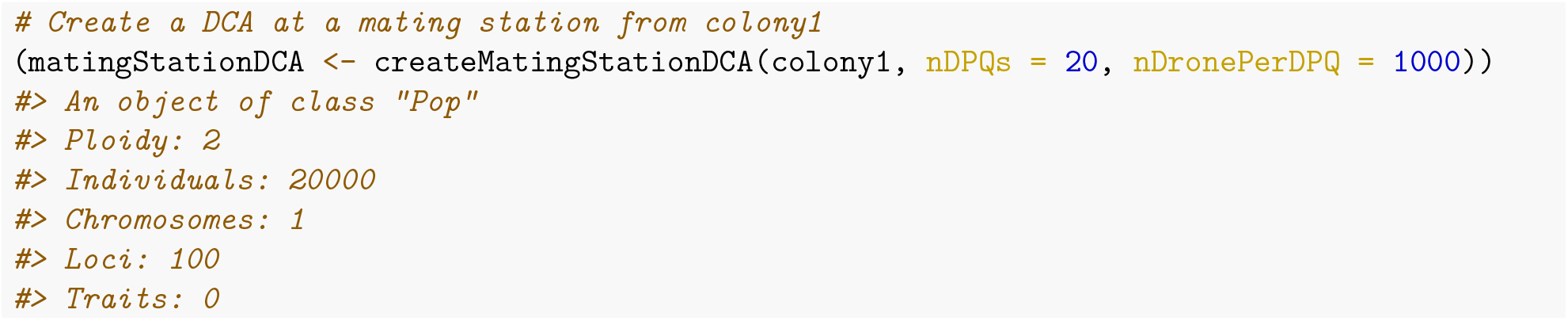

We see that the output of the function is a single population of 20,000 drones that represents the DCA. Once you have the DCA, you can cross virgin queens either by pulling out populations of drones or creating a mating plan as described above.

Here, we will mate a single colony (`colony3`) and a group of colonies (`apiary3`) at a mating station according to a cross plan.

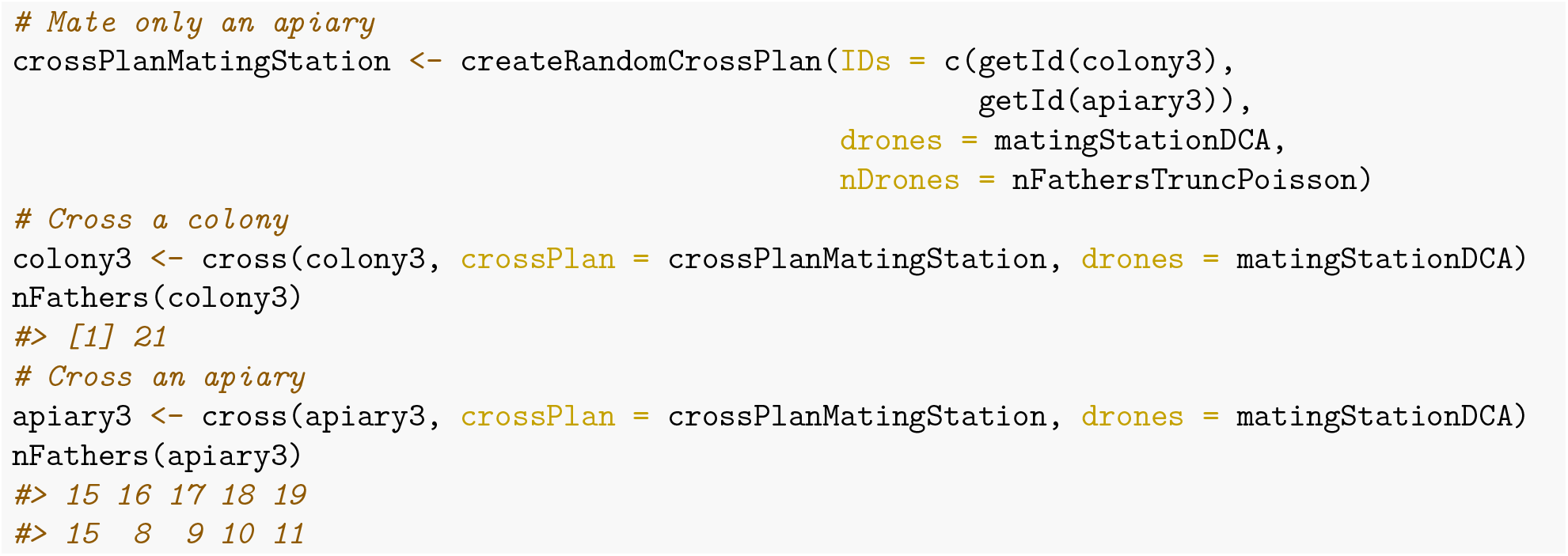

### Cross virgin queens with different methods

It could happen that you have some virgin colonies in an apiary and you want to inseminate one of the virgin queens artificially with a single drone, take three of them to a mating station, and mate the rest of them openly at a local DCA. Since the cross plan is a named list, you can concatenate multiple cross plans into one. Let’s mate the multicolony `apiary4` in such a manner.

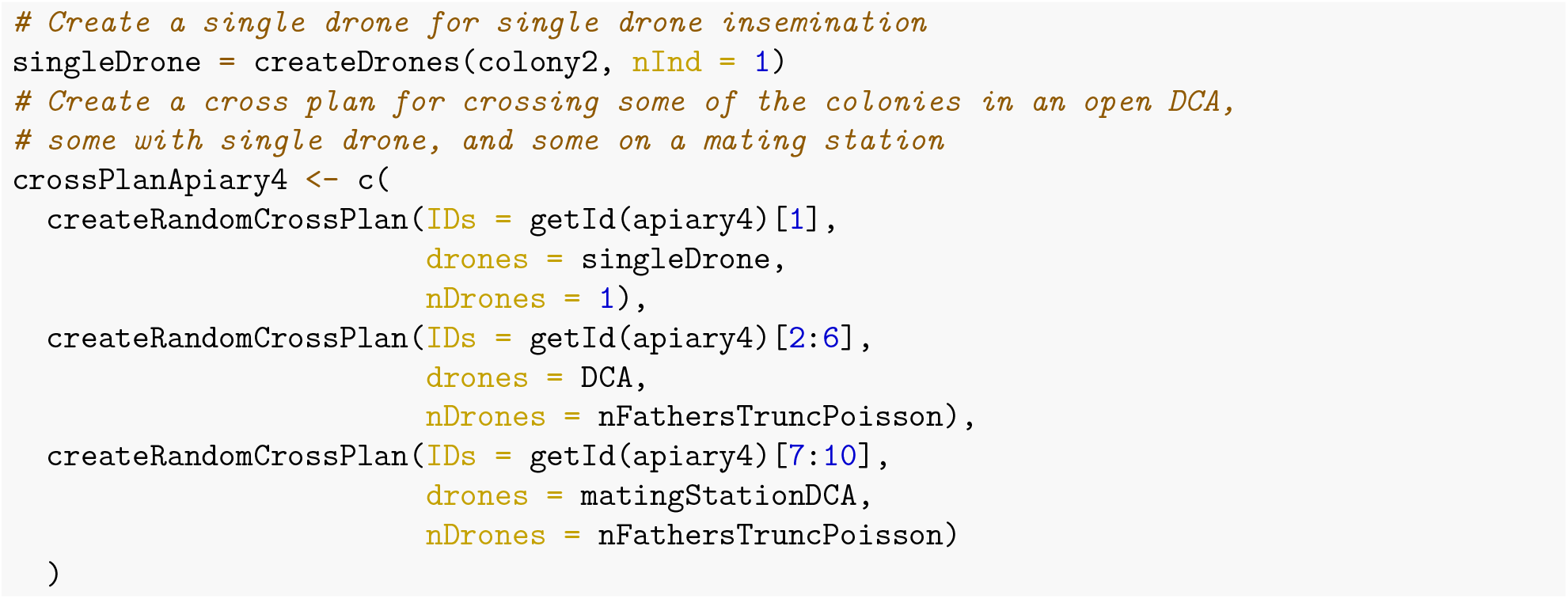

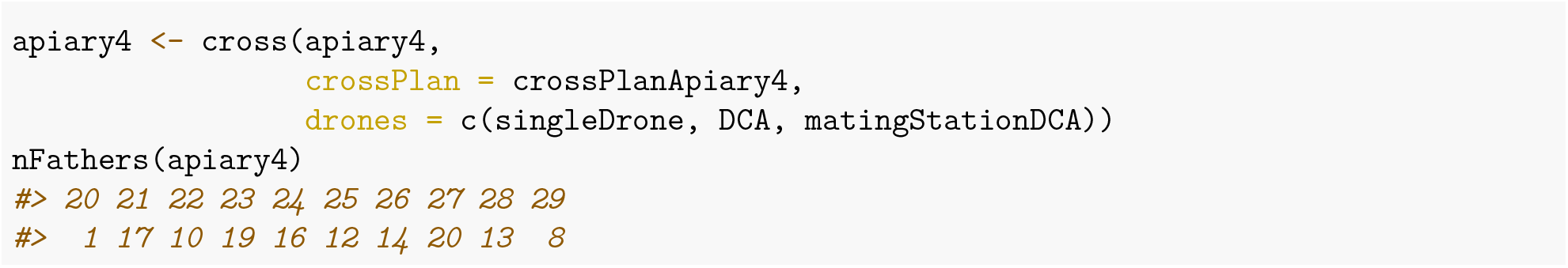

## Additional file 5 – Genomics vignette

### Introduction

This vignette demonstrates how SIMplyBee manages and manipulates the honey bee’s genomic information. Specifically, it describes:

- how to obtain the genomic information,
- how to pool genotypes, and
- how to compute genomic relationship matrices.

Let’s first create a colony.

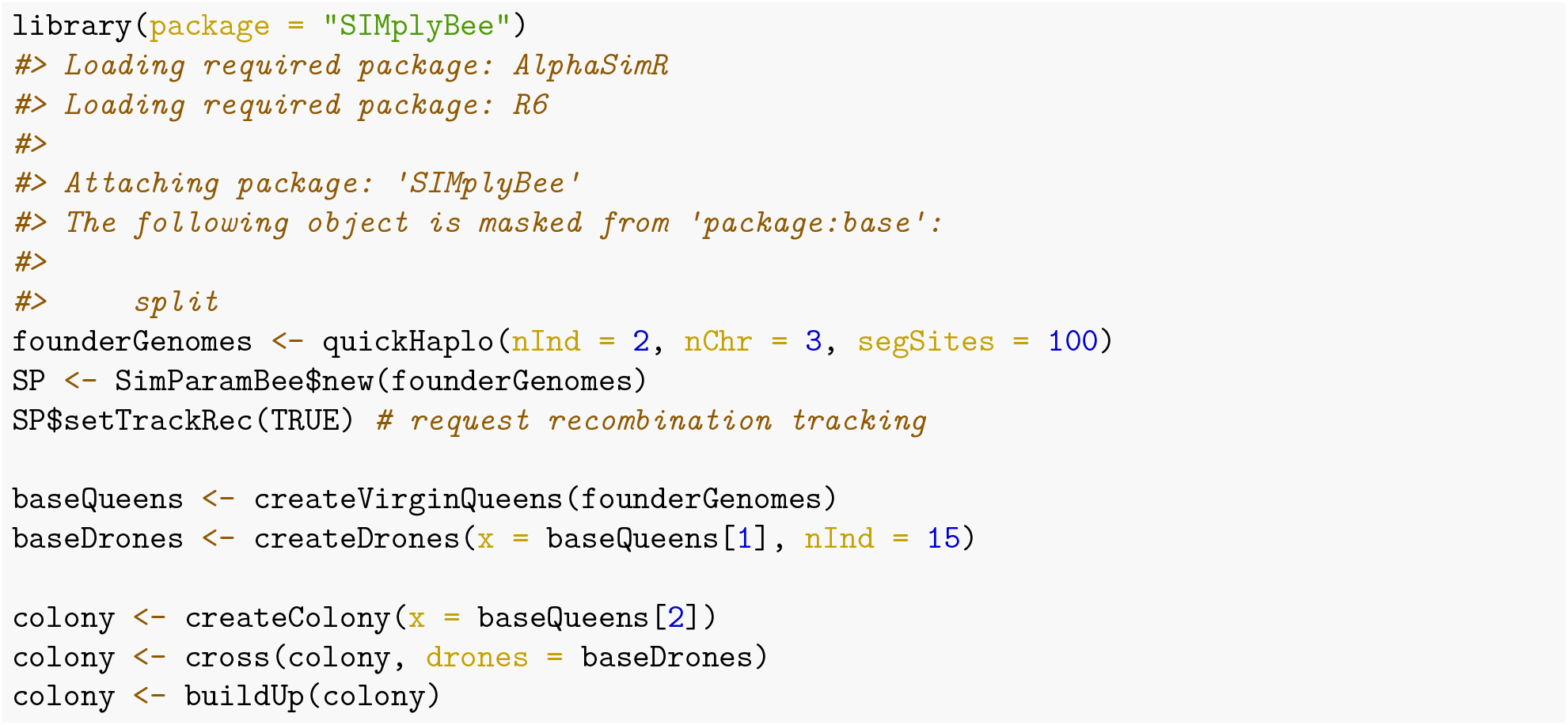

### Obtaining genomic information

Honeybees have a haplo-diploid inheritance system where queens and workers are diploid and drones are haploid. In SIMplyBee, we simulate drones as doubled-haploids, that is, as fully homozygous diploid individuals. This means that they have two identical sets of chromosomes. When they produce sperm, their gametes all have the same one set of chromosomes. Despite them being diploid, we generally return a haploid set of chromosomes from drones, unless specifically requested that you want the doubled-haploid genotype.

Following AlphaSimR, SIMplybee has a group of genome retrieval functions `get*Haplo/Geno()` which extract haplotypes and genotypes for all segregating sites (`SegSites`), quantitative trait loci (`QTL`), markers (`SNP`), and the identical by descent (`IBD`) haplotypes. Here, site, locus and marker are all synonyms for a position in the genome. These functions leverage AlphaSimR functionality, but work with SIMplyBee’s `Colony` or `MultiColony` objects and in addition take the `caste` argument to extract information for a specific caste. Another argument you can use with this function is `collapse = TRUE/FALSE`. If `collapse = TRUE` then all of the information is collapsed together and a single matrix is returned, if `collapse = FALSE` we return a list by caste or by colony.

We recommend that you study the index of available `get*()` functions in SIMplyBee and read this vignette for a short demonstration.

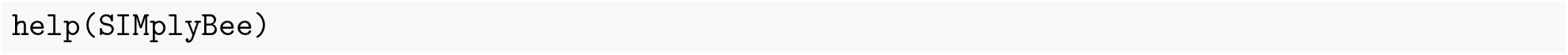

To show all this functionality, let’s get haplotypes and genotypes across the segregating sites for the different castes using `getSegSitesGeno()` or `getSegSitesHaplo()`. The first row of the output shows marker identifications (chromosome_locus) and the first column shows haplotype identifications (individual_haplotype). The alleles are represented with a sequence of 0’s and 1’s. Let’s first obtain the information at the segregating sites for the queen (we limit the output to the first 10 sites):

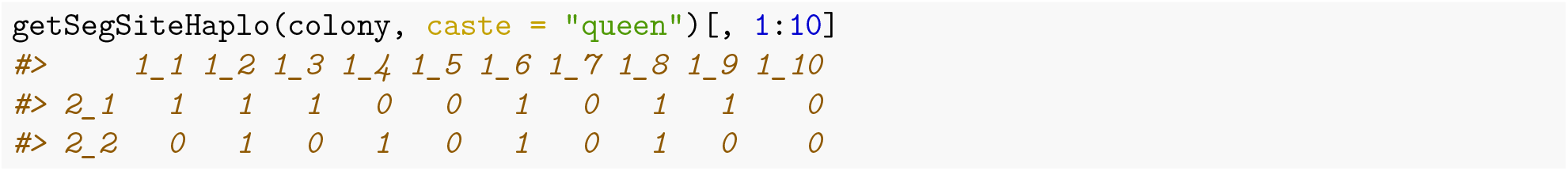

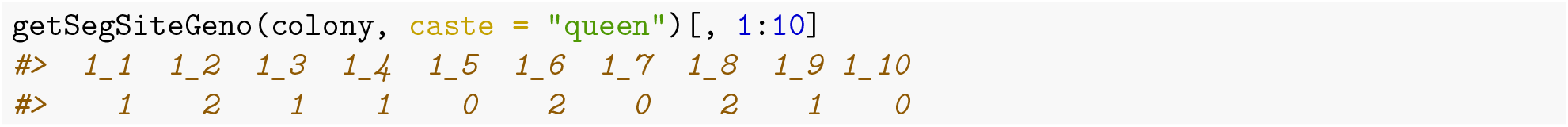

Now for the fathers:

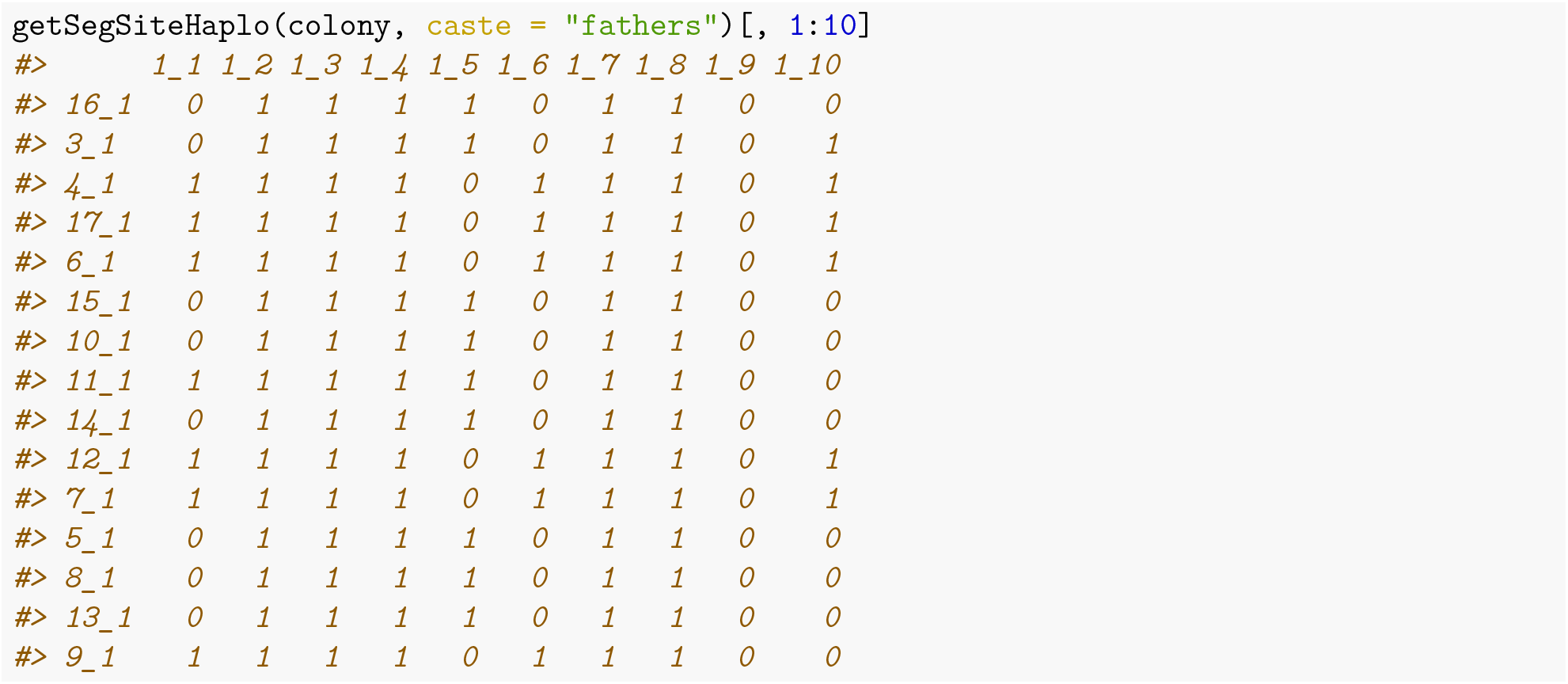

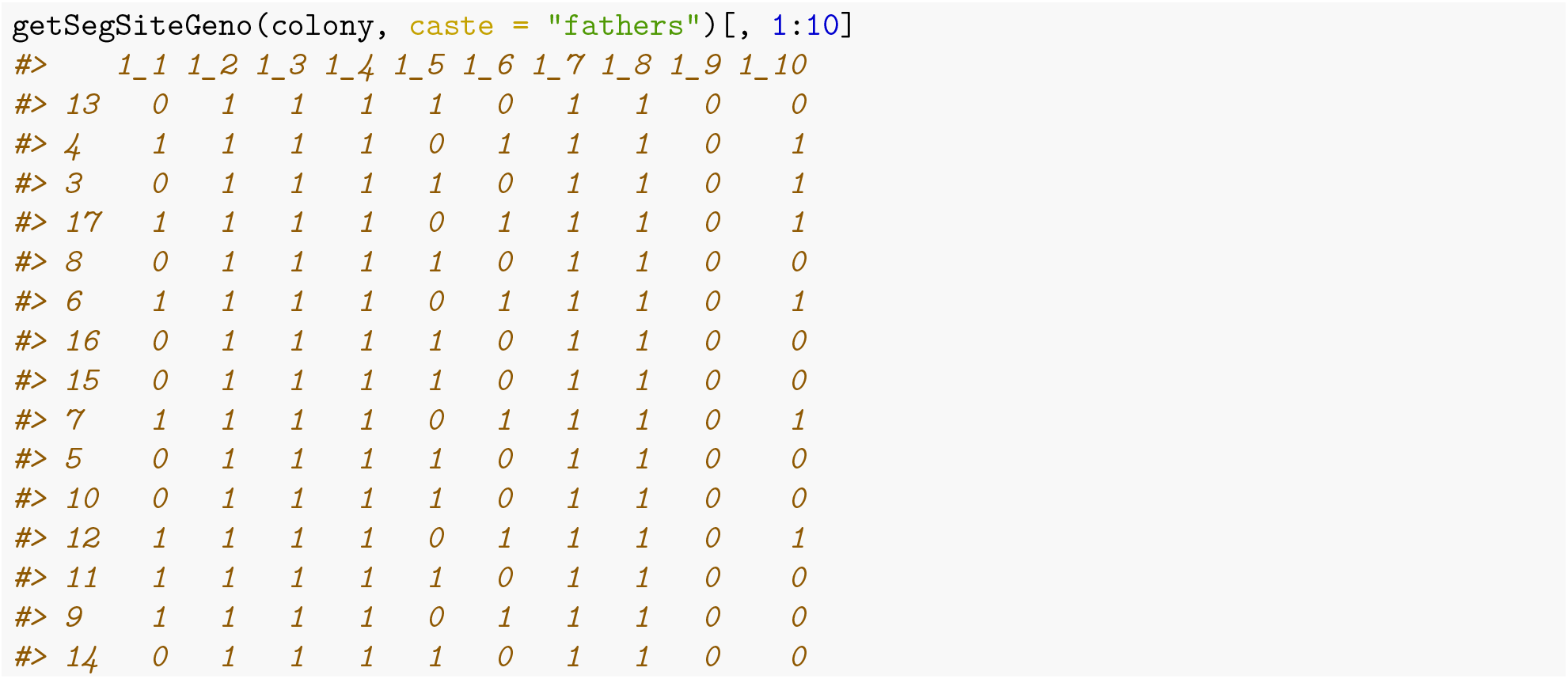

Since father are drones, and these are haploid, we get one row per father. We can retrieve the doublet-haploid (diploid implementation) state, if this is desired (showing just one father to show this clearly):

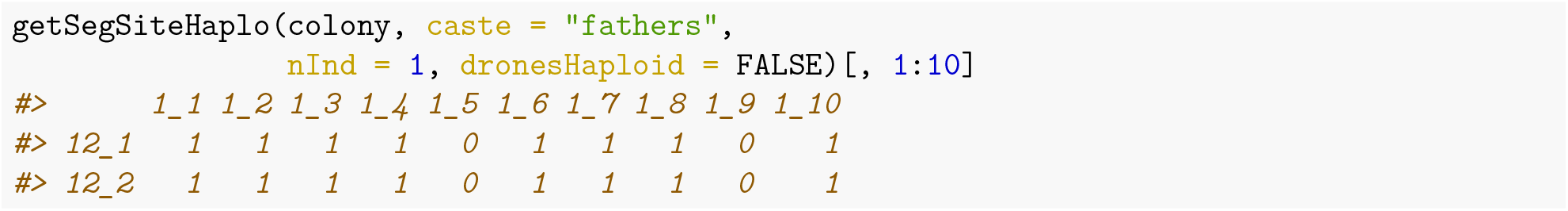

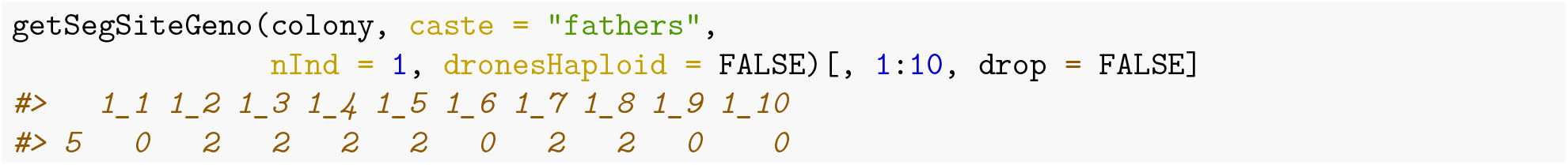

Now two workers:

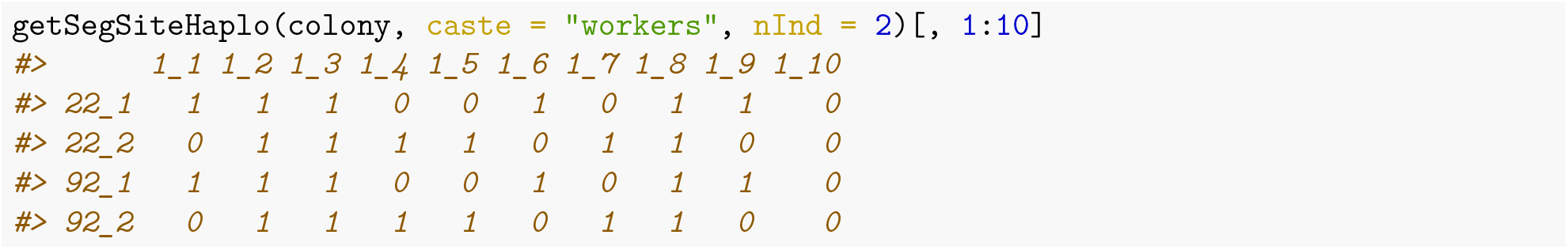

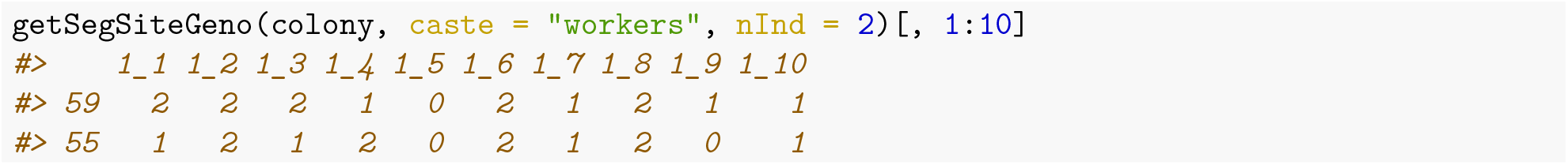

And finally four drones:

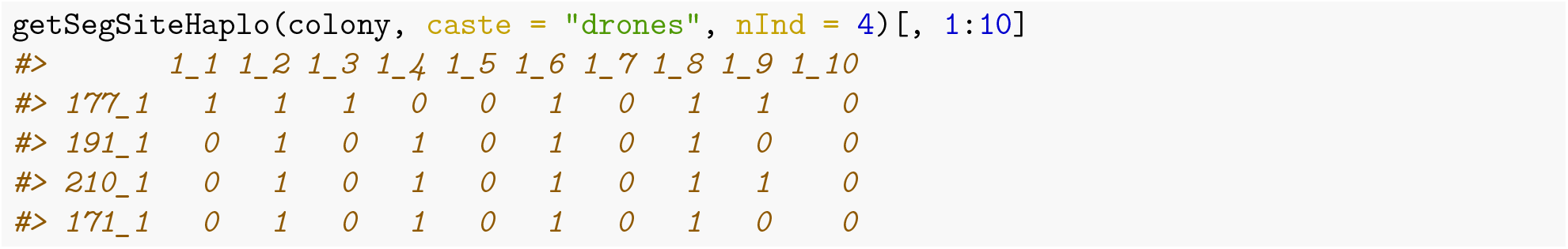

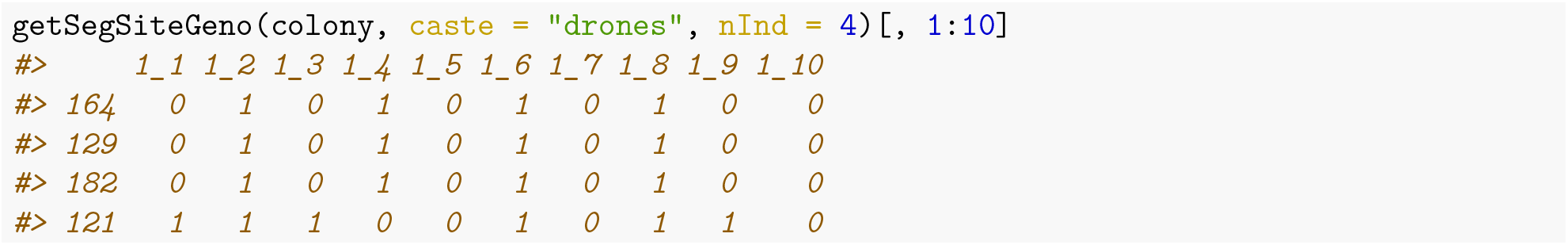

You can also use `caste = “all”` to get the haplotypes and phenotypes from every individual in the colony. If the argument `collapse` is set to `FALSE`, then the function returns a list with haplotypes for each caste. Let’s explore the structure of the output:

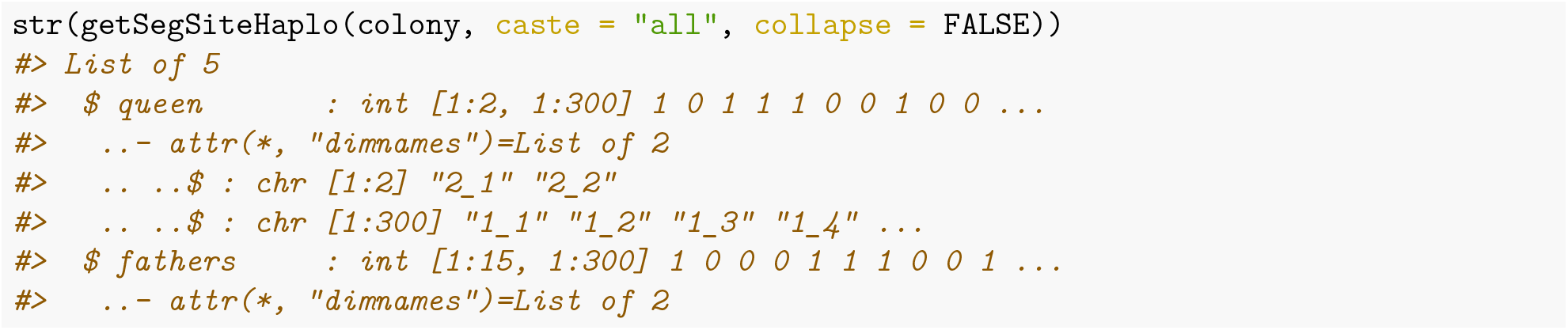

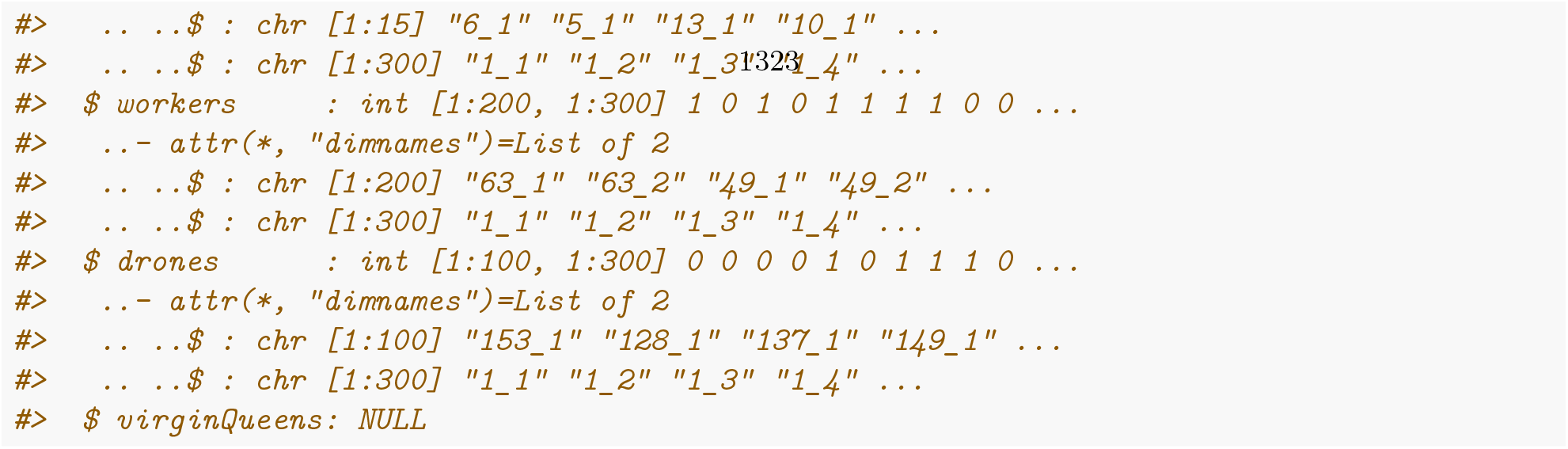

If the argument `collapse` is set to `TRUE`, the function returns a single matrix with haplotypes of all the individuals. The same behaviour is implemented for all the functions that extract genomic information

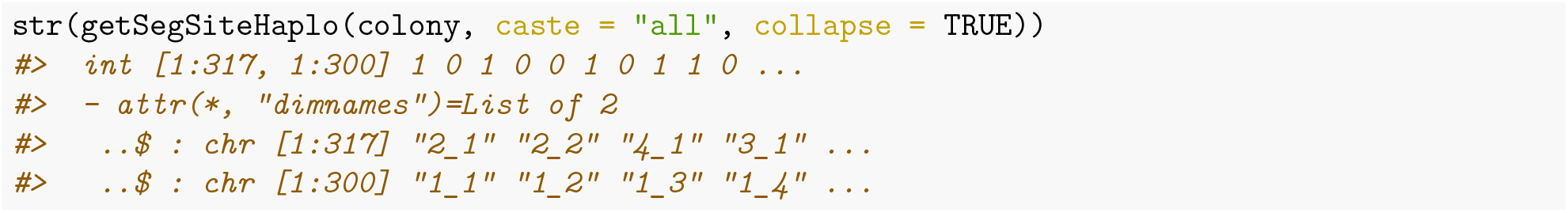

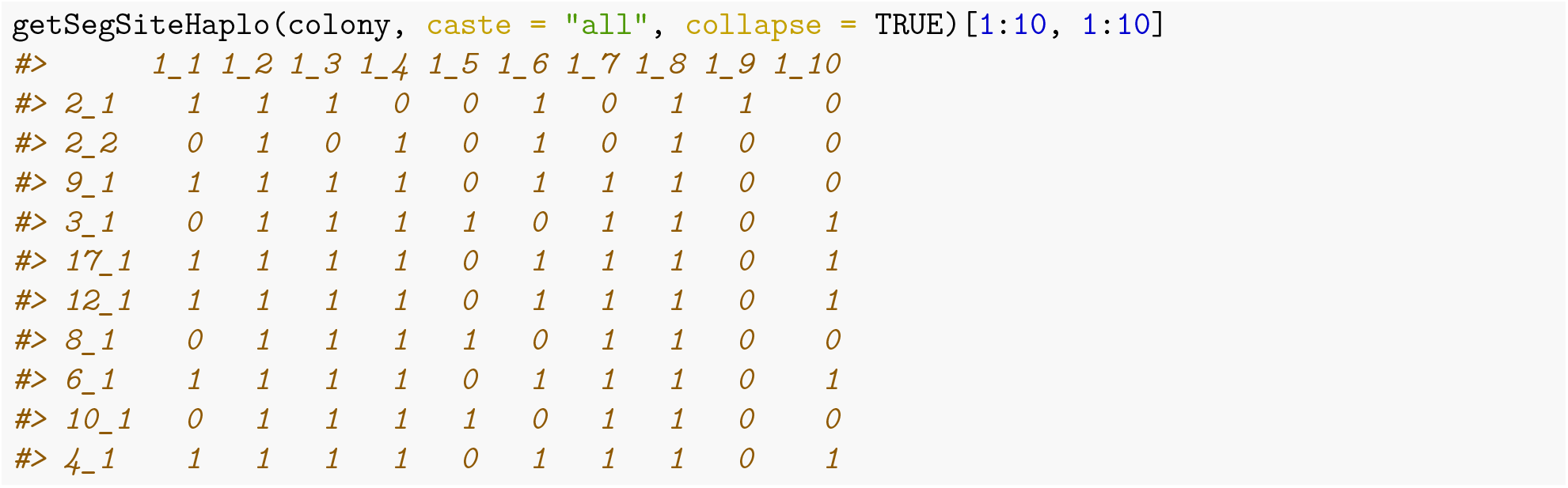

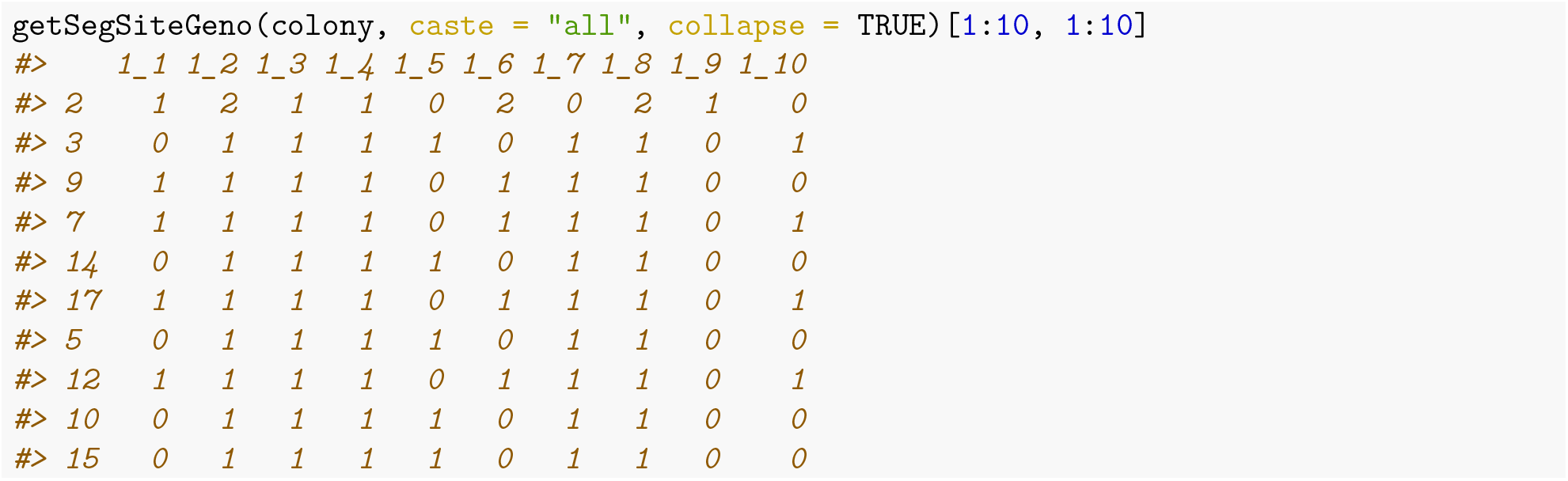

SIMplyBee also has shortcuts for these haplotype and genotype functions to make life a bit easier for the user:

- `getQueenSegSitesHaplo()`
- `getQueenSegSitesGeno()`
- `getFathersSegSitesHaplo()`
- `getFathersSegSitesGeno()`
- `getWorkersSegSitesHaplo()`
- `getWorkersSegSitesGeno()`
- `getDronesSegSitesHaplo()`
- `getDronesSegSitesGeno()`
- `getVirginQueensSegSitesHaplo()`
- `getVriginQueensSegSitesGeno()`

Similar aliases exist also for extracting information about quantitative trait loci (`QTL`), markers (`SNP`), and the identical by descent (`IBD`) haplotypes.

### Pooling genotypic information

Unfortunately, in real life it’s challenging to get the genotype of every individual honeybee and so SIMplyBee provides the function `getPooledGeno()` to imitate real life data. `getPooledGeno()` returns a pooled genotype from individual genotypes to mimic the genotyping of a pool of colony members. A comparison of pooled and individual genotypes also allows the user to compare the two and see the impact of pooled samples on results.

Firstly let’s obtain the genotypes of the workers and of the queen so that they’re easier to work with:

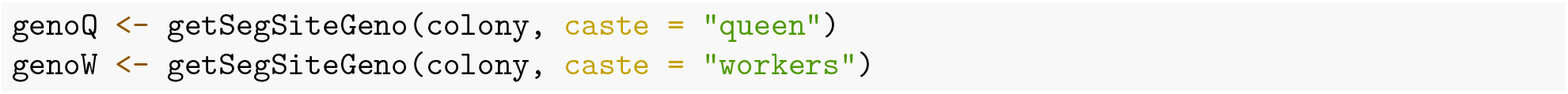

The function `getPooledGeno()` required also the sex of individuals whose genotype are getting pooled (`F` for females and `M` for males).

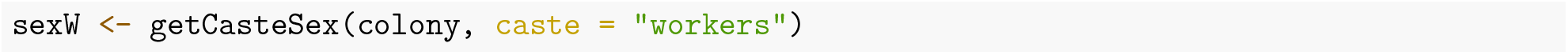

You have two options when choosing what kind of pooled genotypes you would like, using the `type =` argument. You can use `type = “mean”` for the average genotypes and `type = “count”` for the counts of reference and alternative alleles.

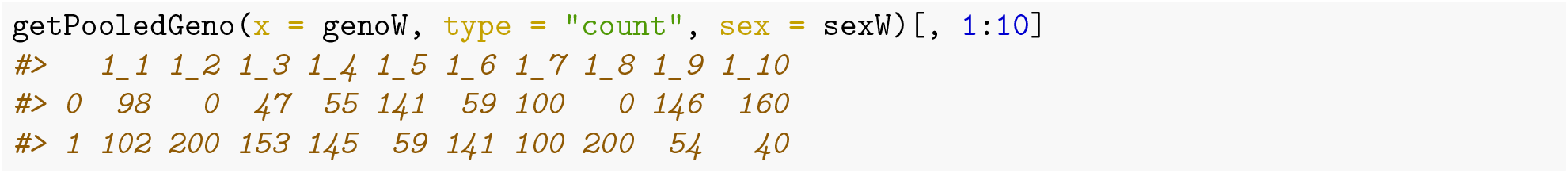

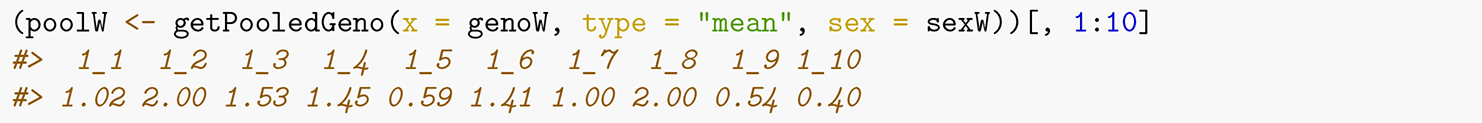

Now lets plot and compare the pooled workers to the queen’s genotype (note the use of jitter for queen’s genotype on the x-axis so we can spread out the dots in the plot!).

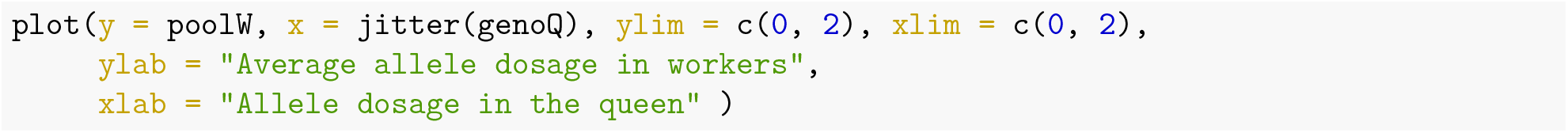

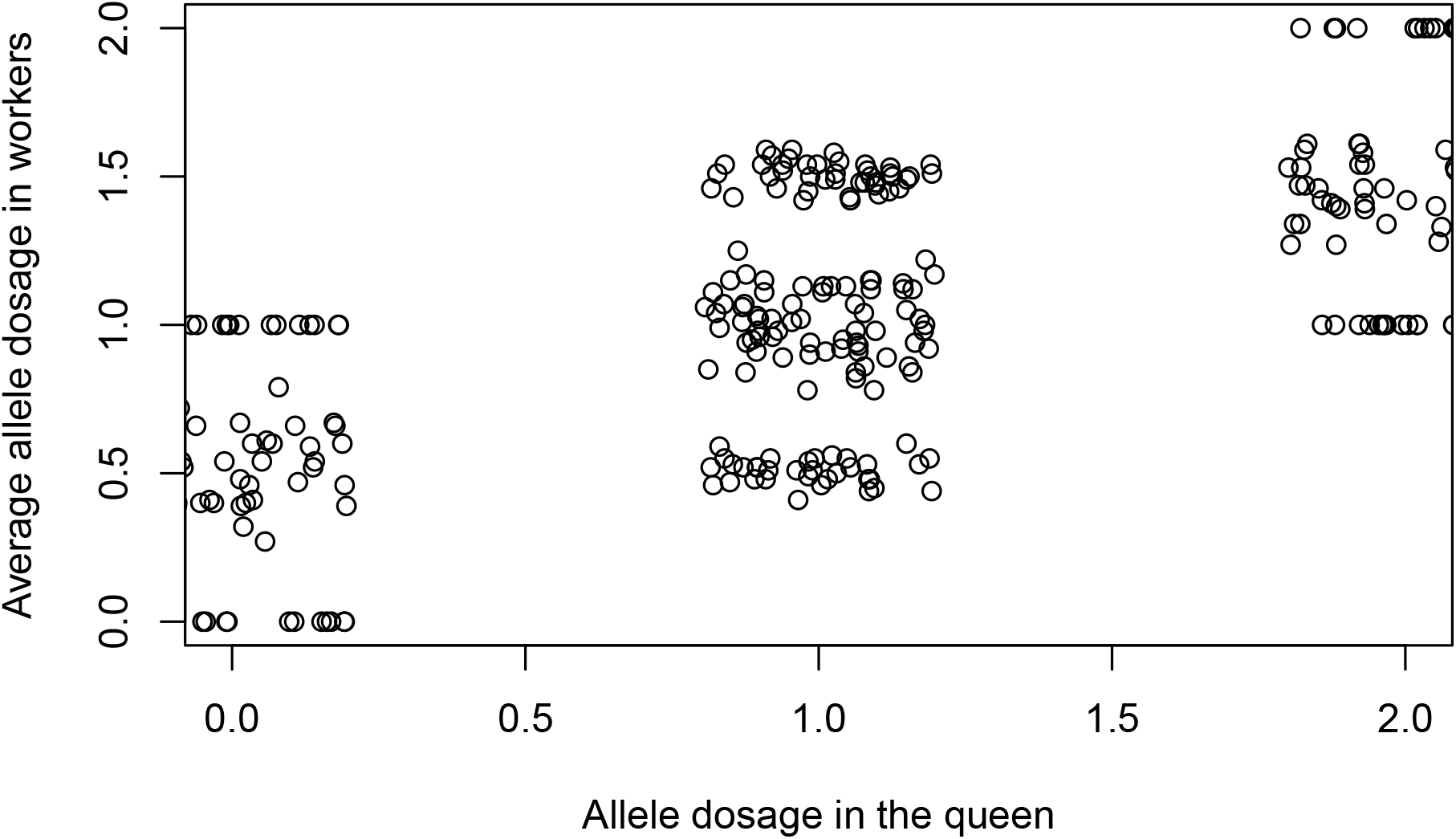

### Computing Genomic Relationship Matrices

This section introduces the calculations of IBD and IBS genomic relationship matrices, so let’s have a quick reminder of what these mean. Identity-by-state (IBS) is a term used when two alleles, two segments or sequences of the genome are identical. Identity-by-descent (IBD) is when a segment of matching (IBS) DNA shared by two or more individuals has been inherited from a common ancestor.

Using IBD and IBS can allow a user to look into the relationships based on the genomic data. We’ll demonstrate this by calculating some Genomic Relationship Matrices (GRM) using SIMplyBee’s `calcBeeGRMIbs()` and `calcBeeGRMIbd()`.

Let’s look at the `calcBeeGRMIbs()` first. This function returns a Genomic Relatedness Matrix (GRM) for honeybees from IBS genomic data (bi-allelic SNP represented as allele dosages) following the method for the sex X chromosome (Druet and Legarra, 2020).

To see this, let’s obtain the genotypes and sex information of all individuals in the colony.

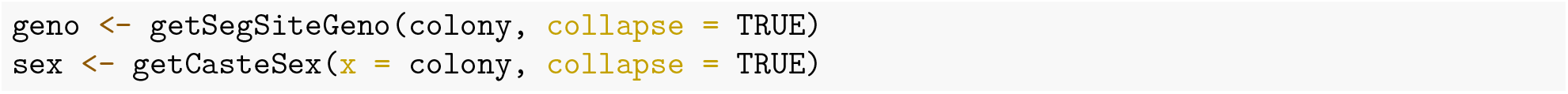

Now let’s calculate the IBS GRM, we will use the genotypes to calculate this:

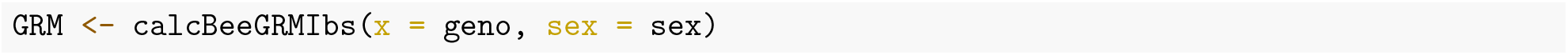

This produces a matrix that we can plot and summarise – its useful to summarise diagonal and off-diagonal values separately.

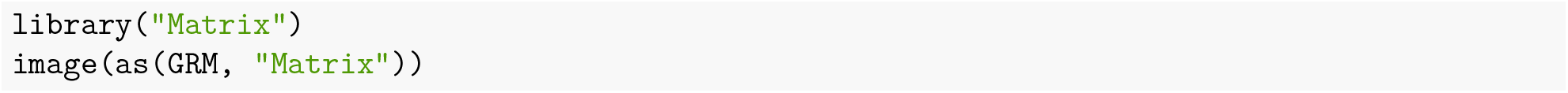

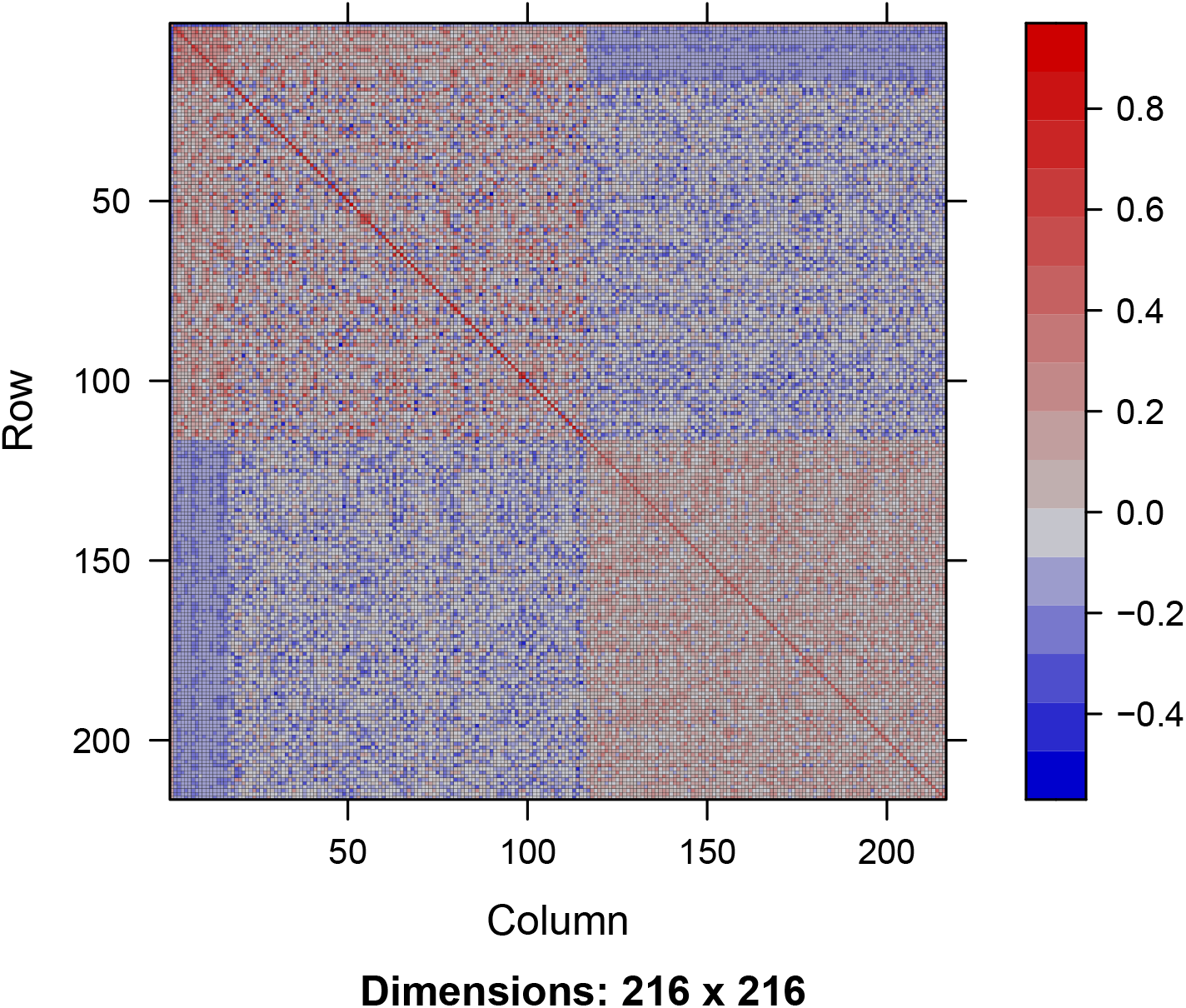

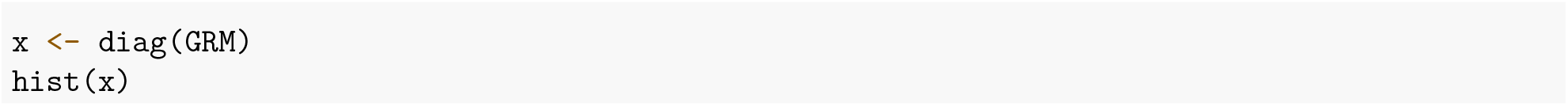

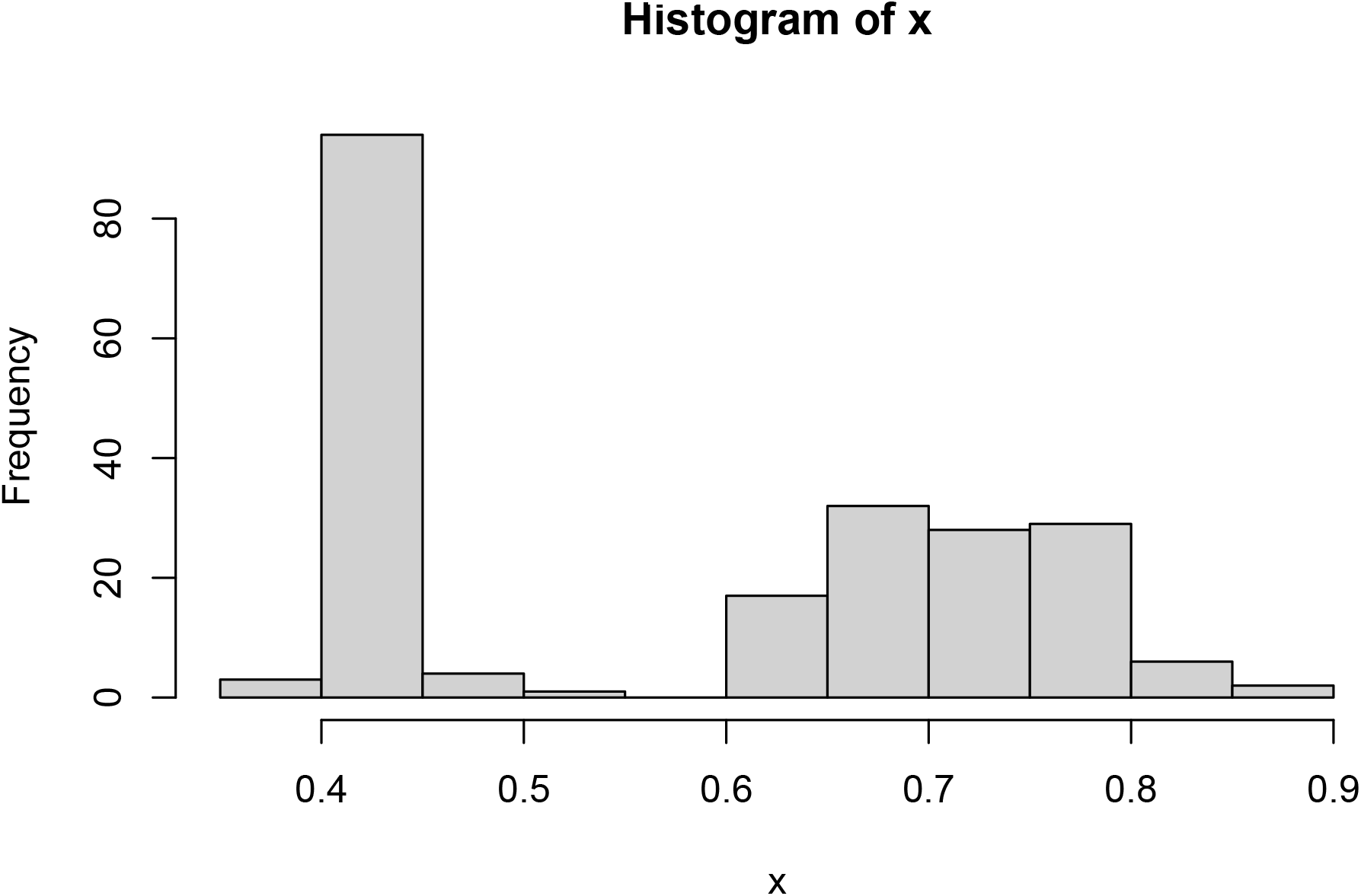

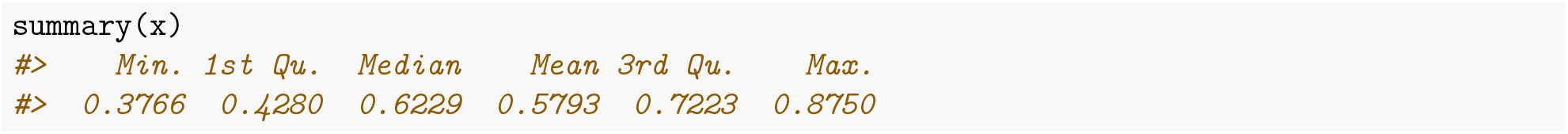

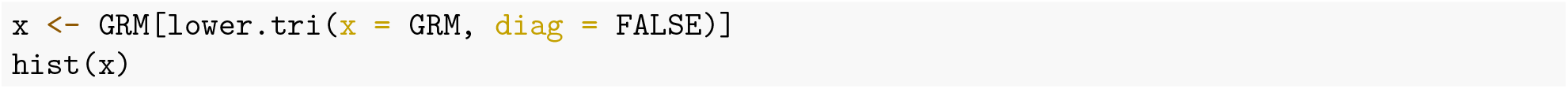

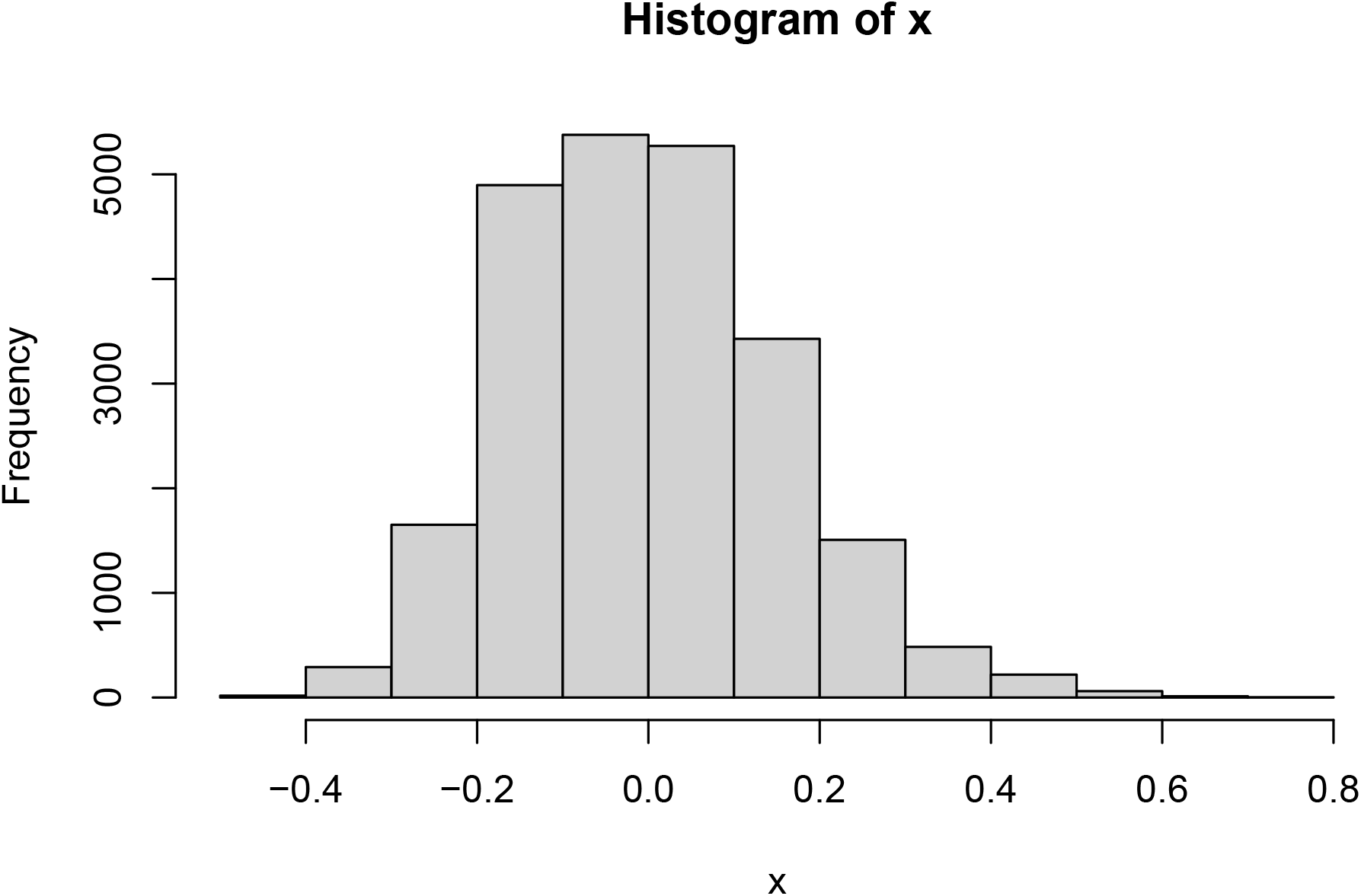

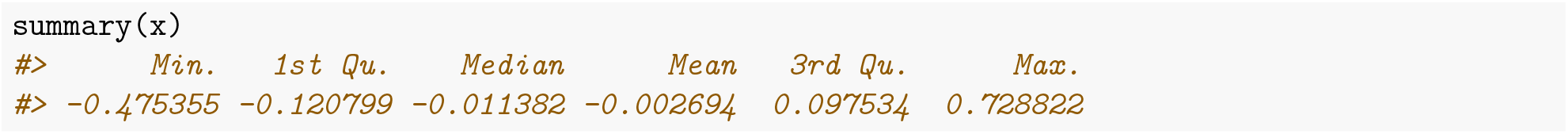

We can also inspect GRM elements between specific caste members:

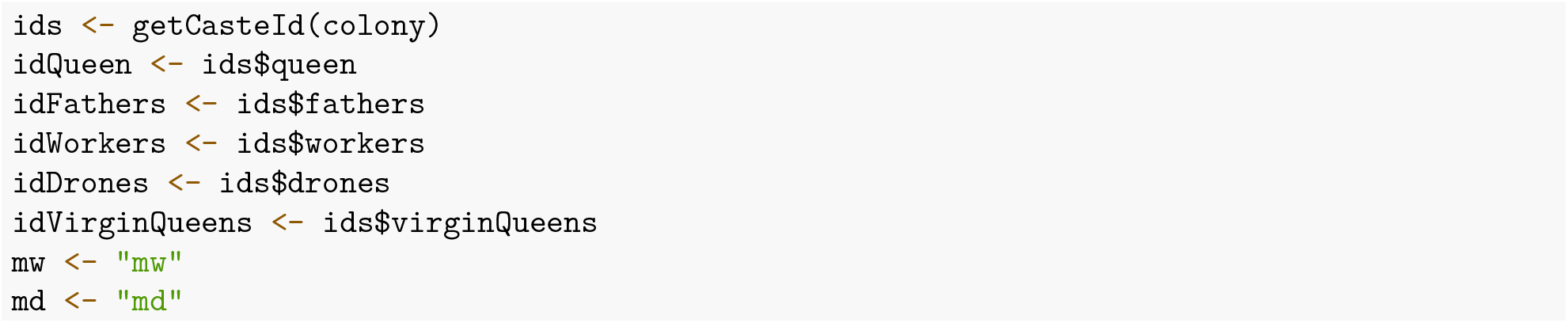

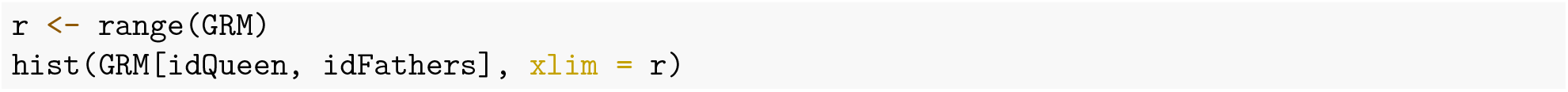

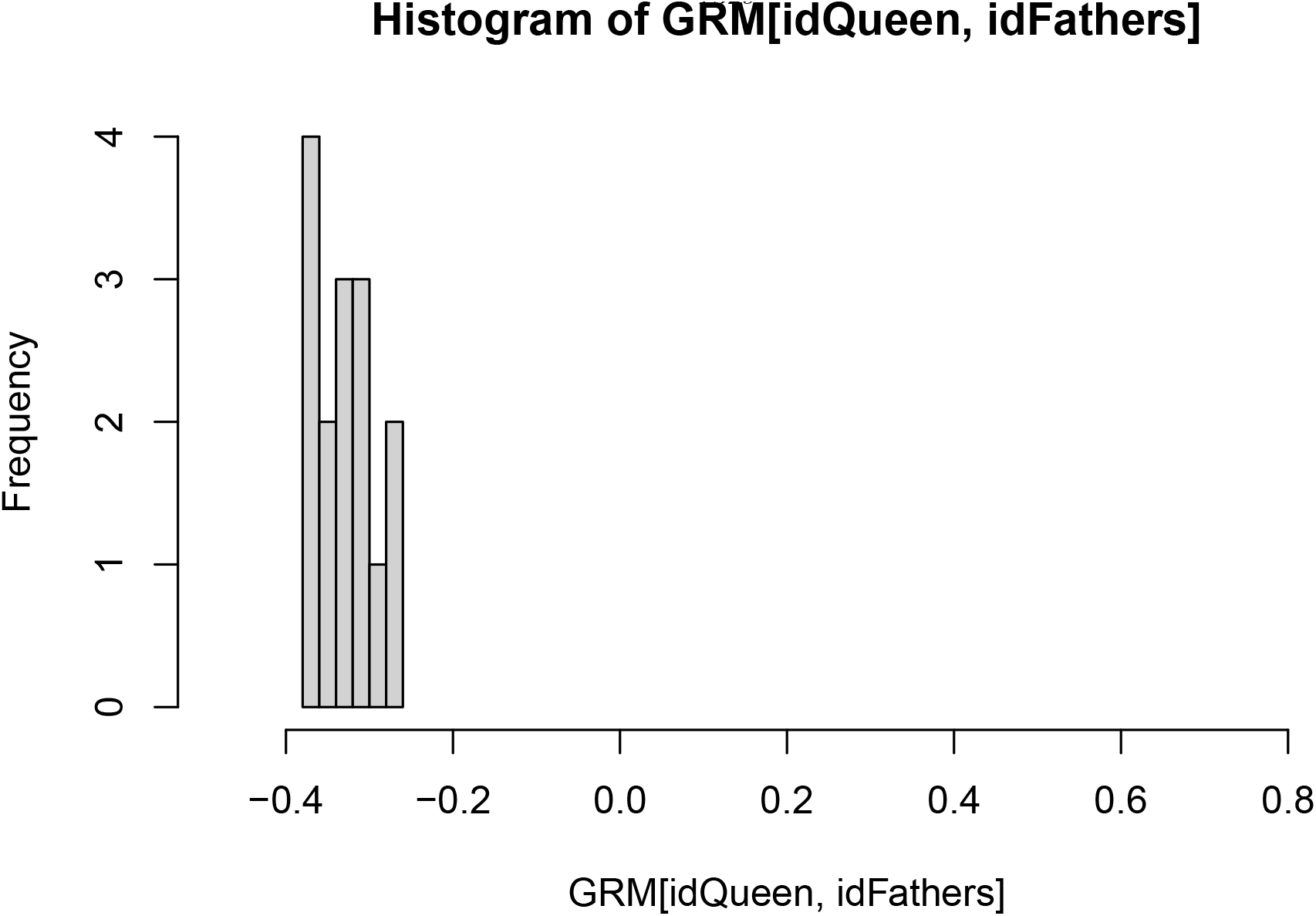

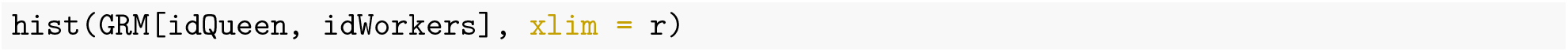

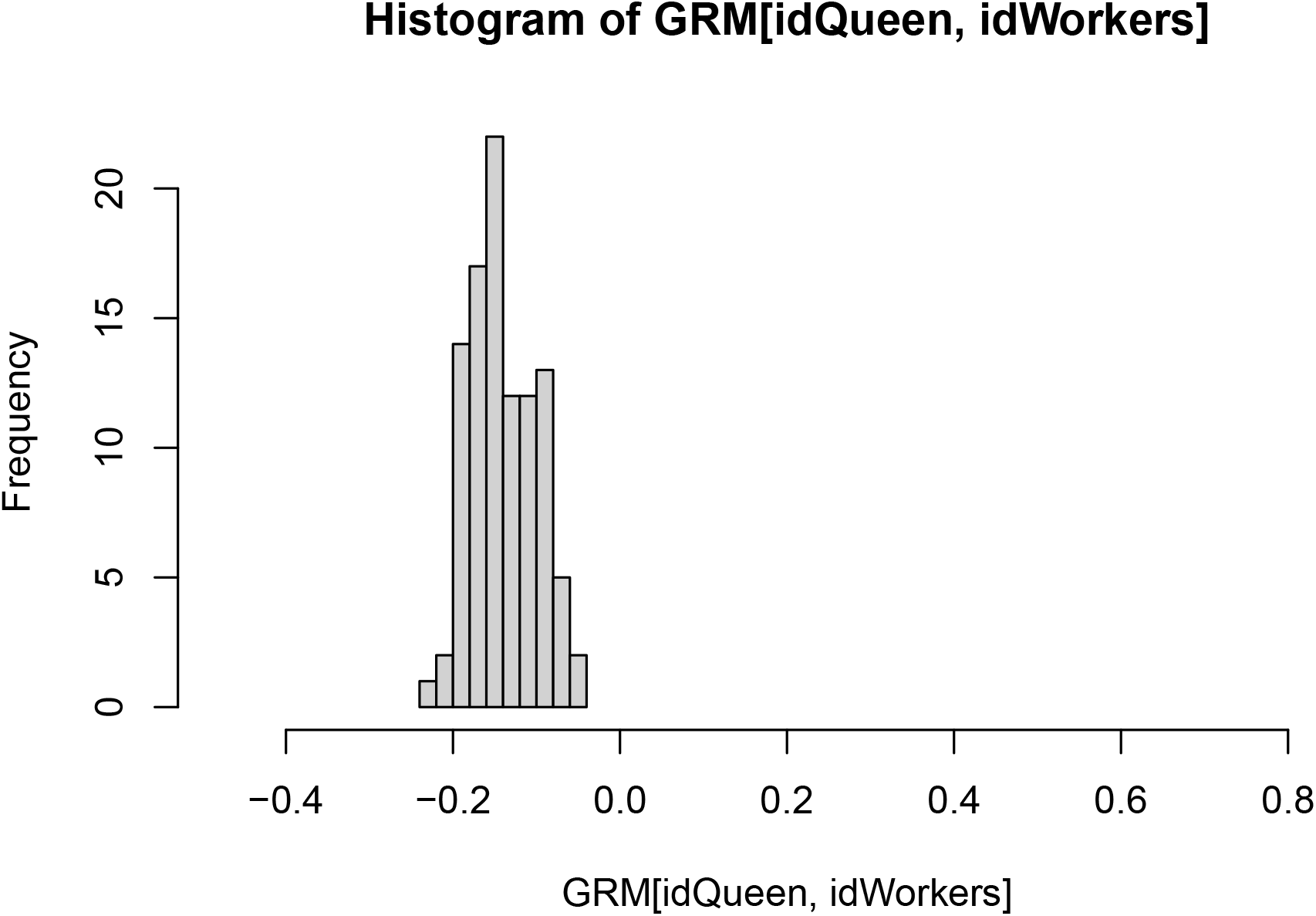

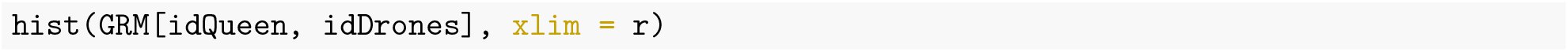

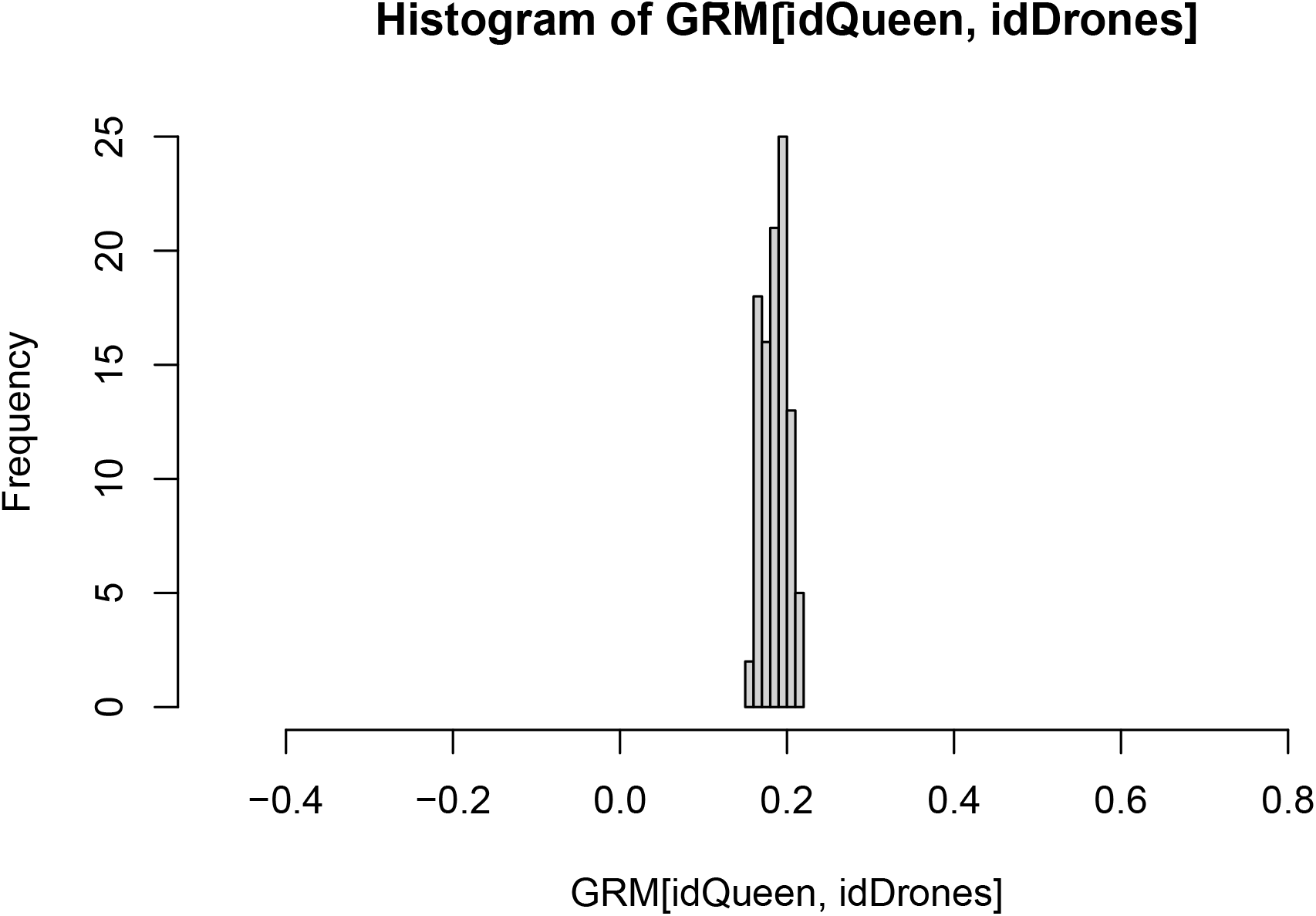

`calcBeeGRMIbs()` uses the `calcBeeAlleleFreq()` function to calculate allele frequencies for centering the honeybee genotypes. You can also use it in some other cases:

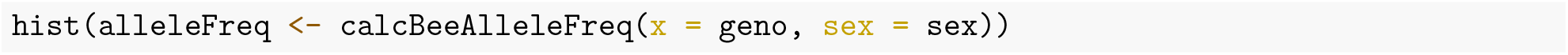

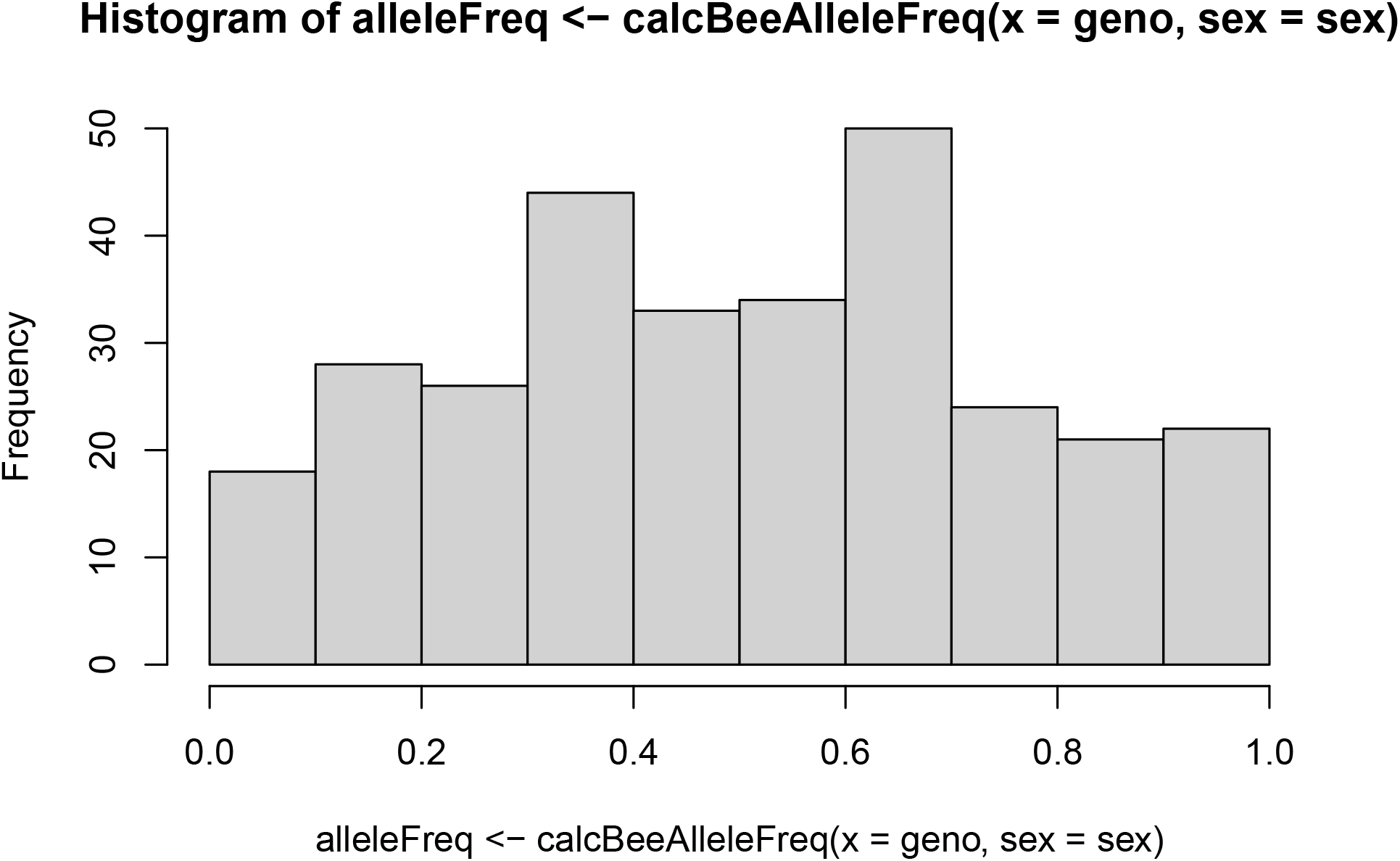

Now lets look at `calcBeeGRMIbd()`. This function creates Genomic Relatedness Matrix (GRM) for honeybees based on Identical-By-Descent (IBD) information. It returns a list with a matrix of gametic relatedness coefficients (between genomes) and a matrix of individual relatedness coefficients (between individuals). Please refer to Grossman and Eisen (1989), Fernando and Grossman (1989), Fernando and Grossman (1990), Van Arendonk, Tier, and Kinghorn (1994), and Hill and Weir (2011) for the background on this function.

Now obtain the IBD haplotypes and compute IBD GRM.

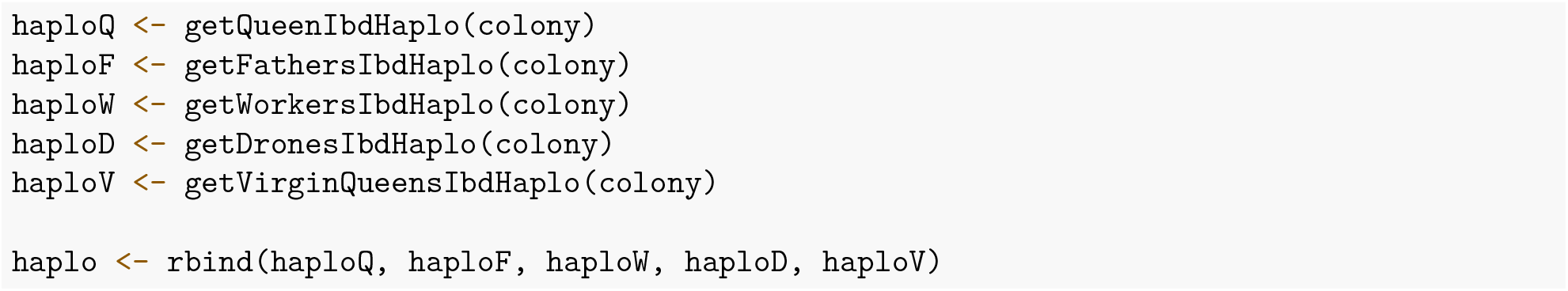

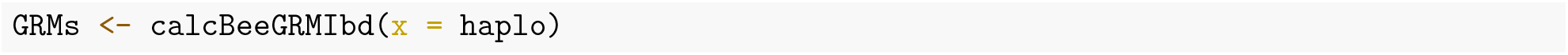

Let’s view this matrix:

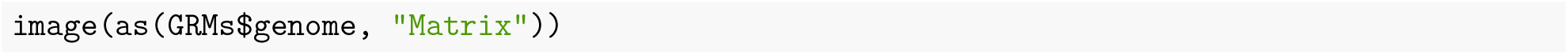

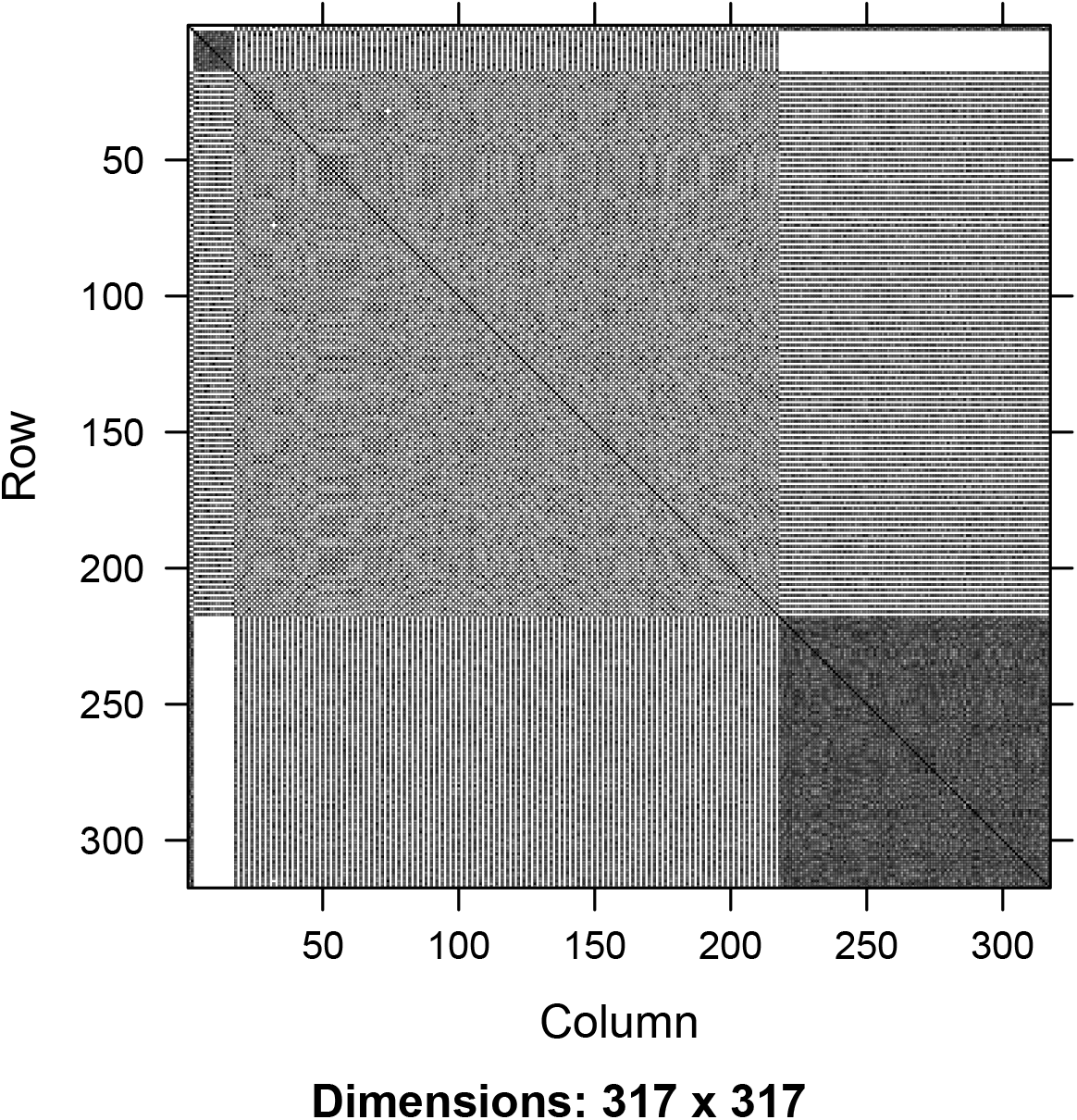

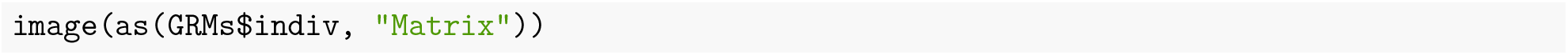

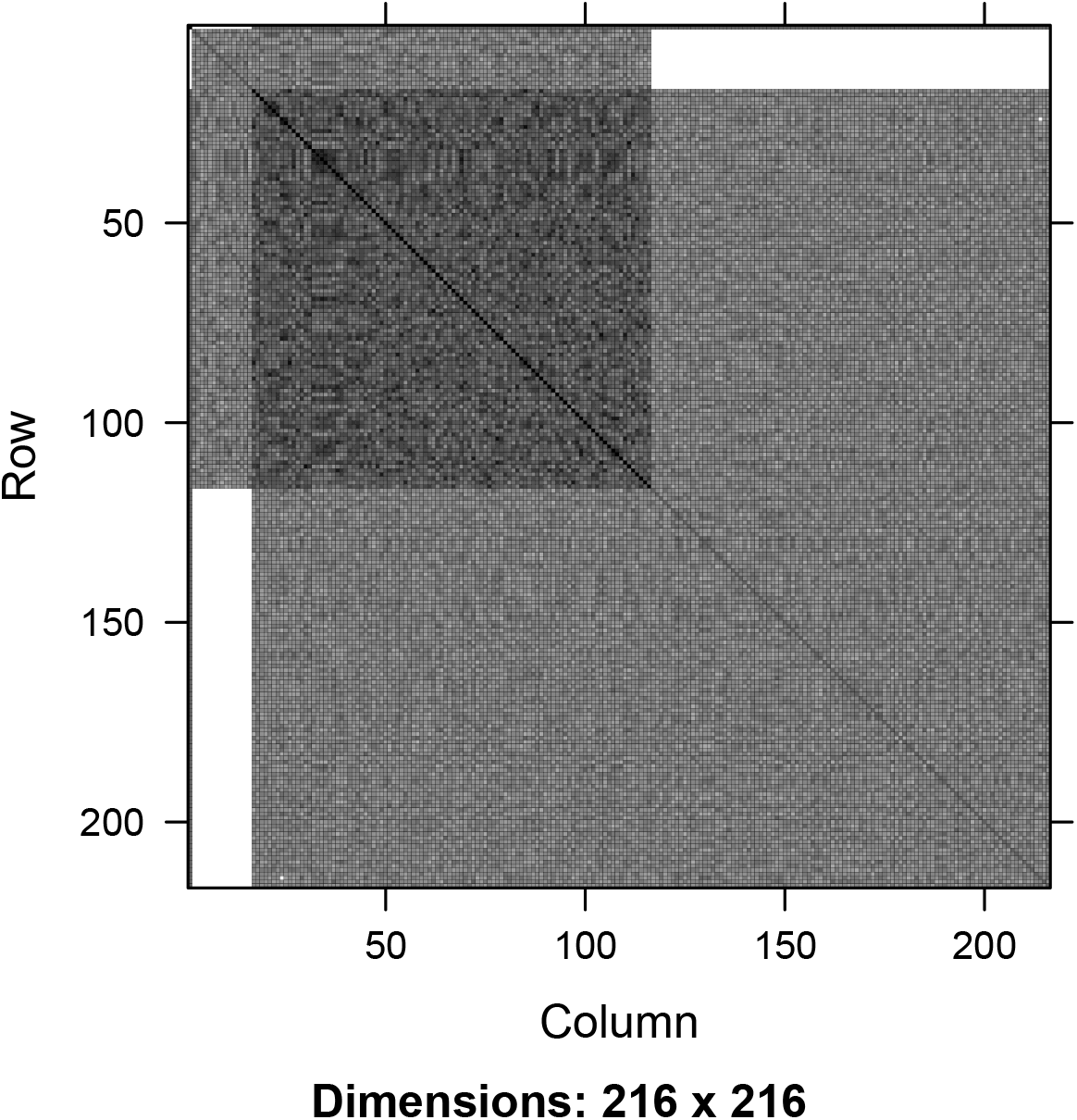

Now we can look at the diagonal of the obtained matrices that represent 1 for a genomes and 1 + inbreeding coefficient individuals.

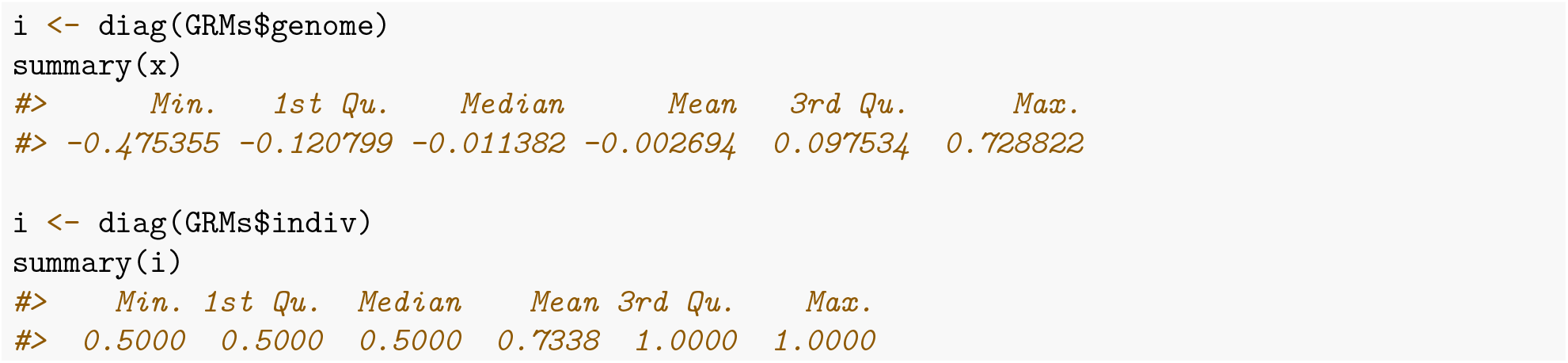

And now the non-diagonals that represent the coefficients of relationship between genomes or between individuals.

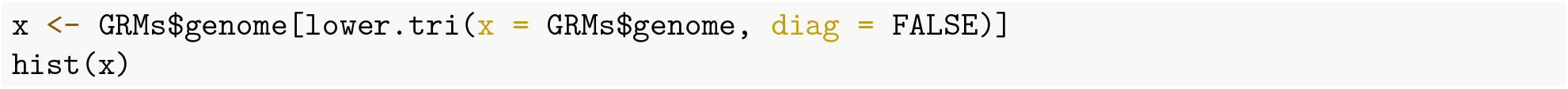

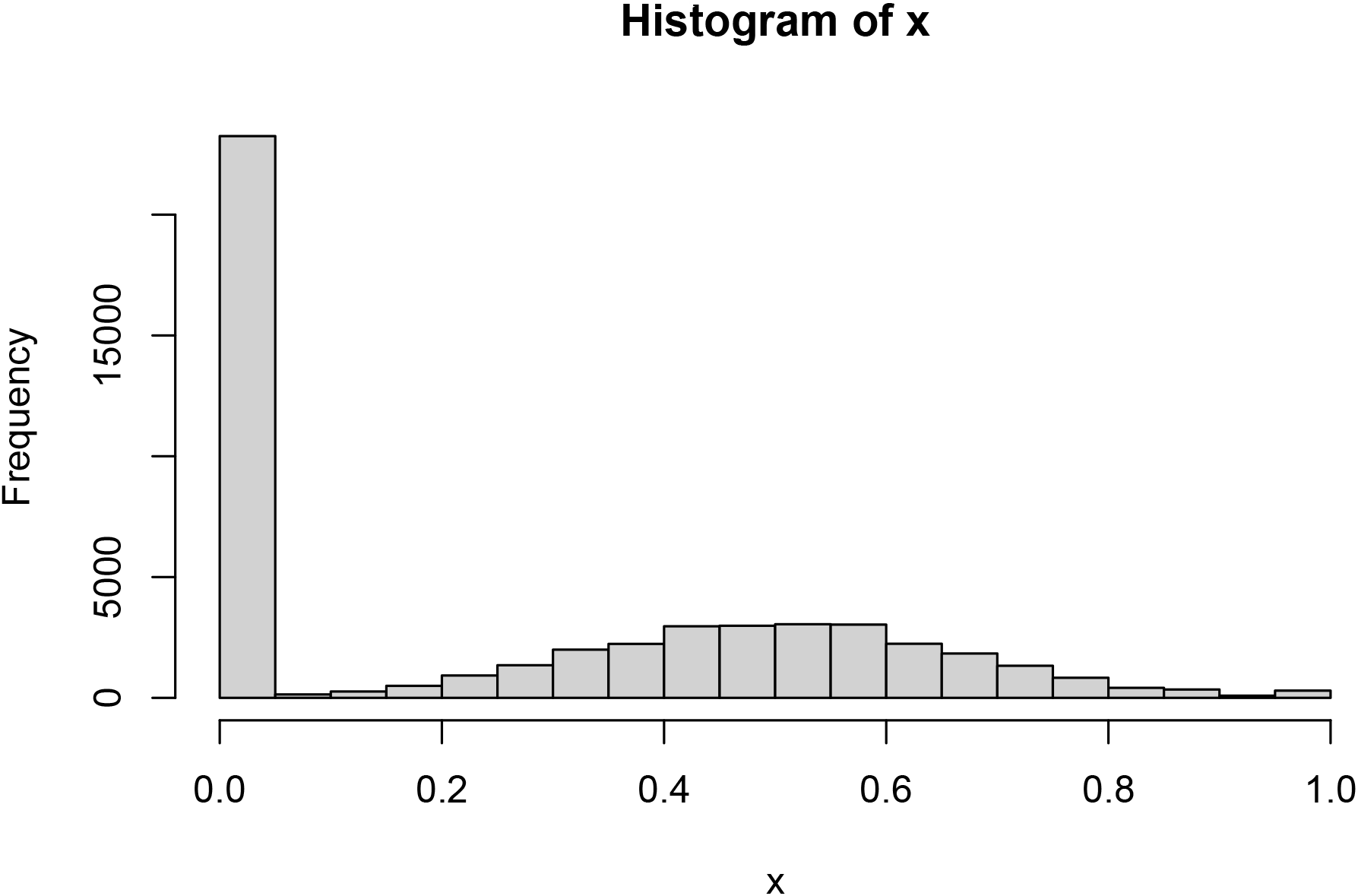

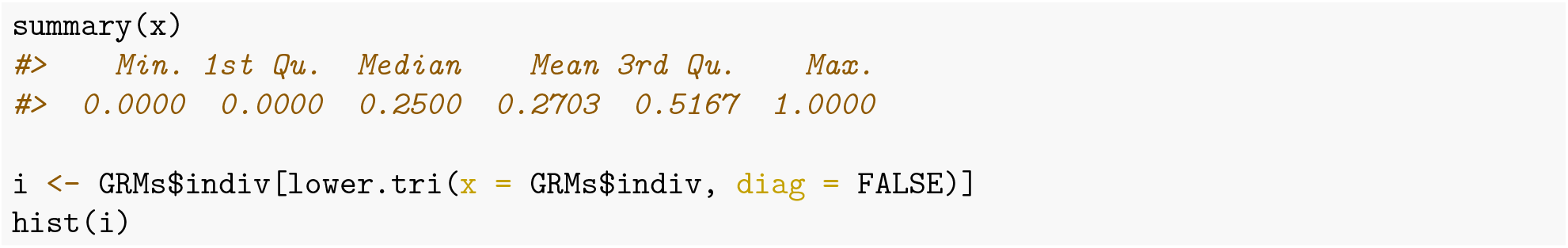

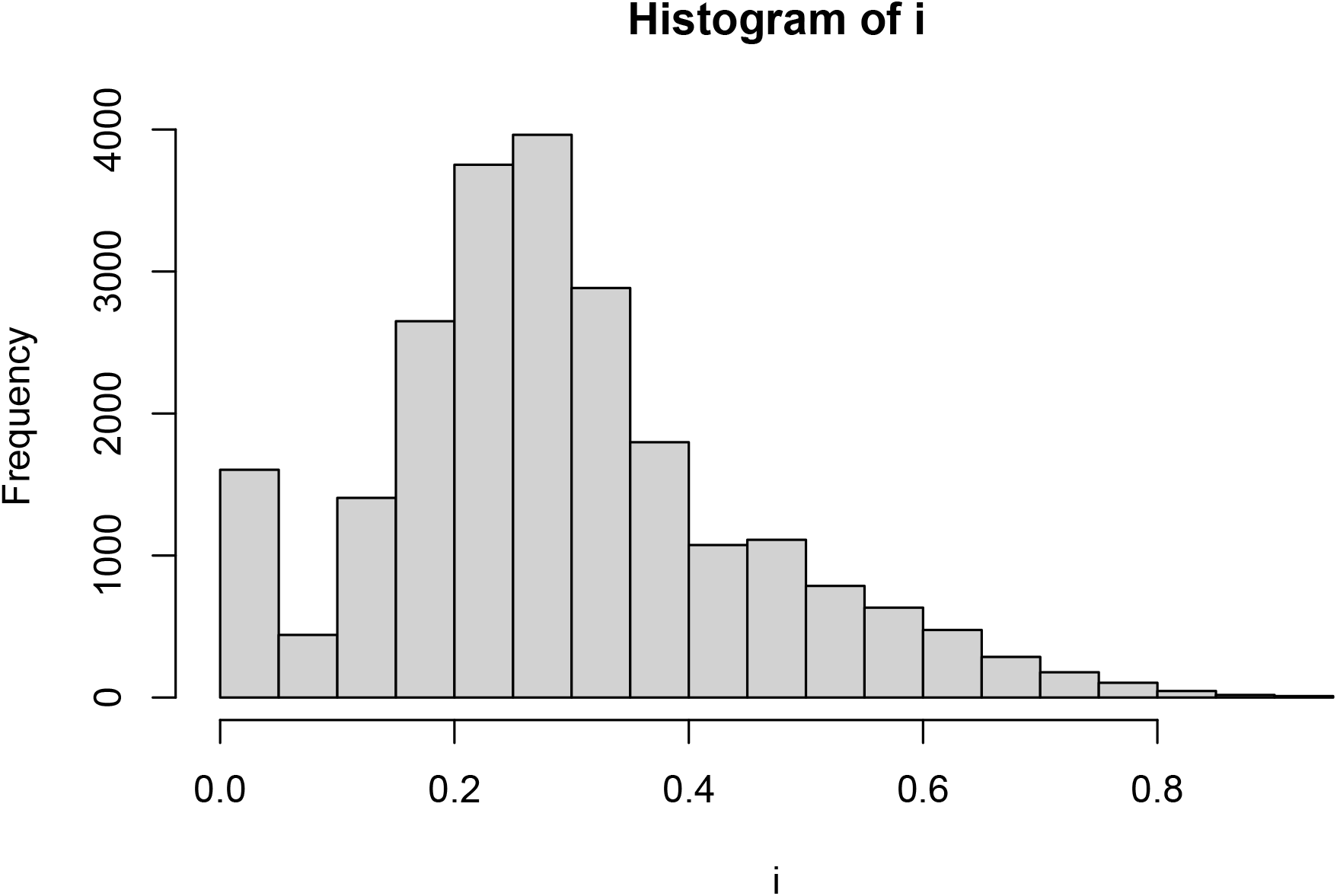

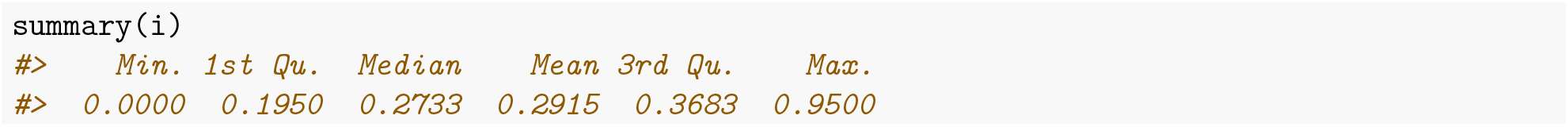

Let’s now compare compare relationships between caste members within a colony.

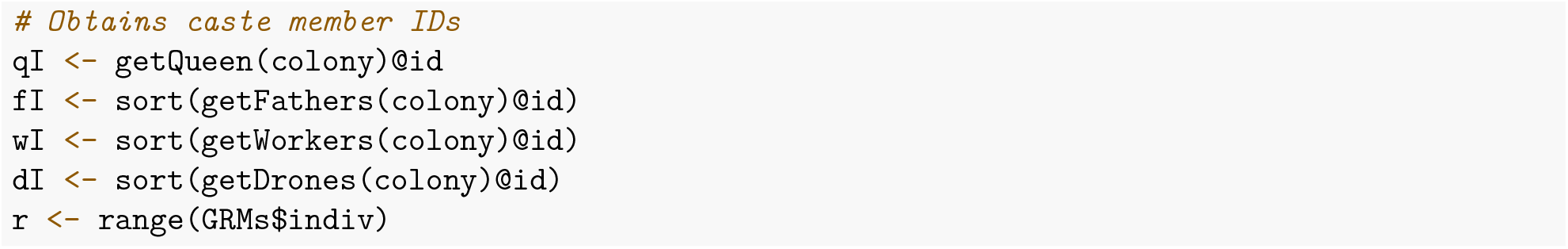

Compare queen and fathers:

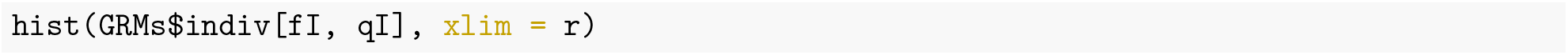

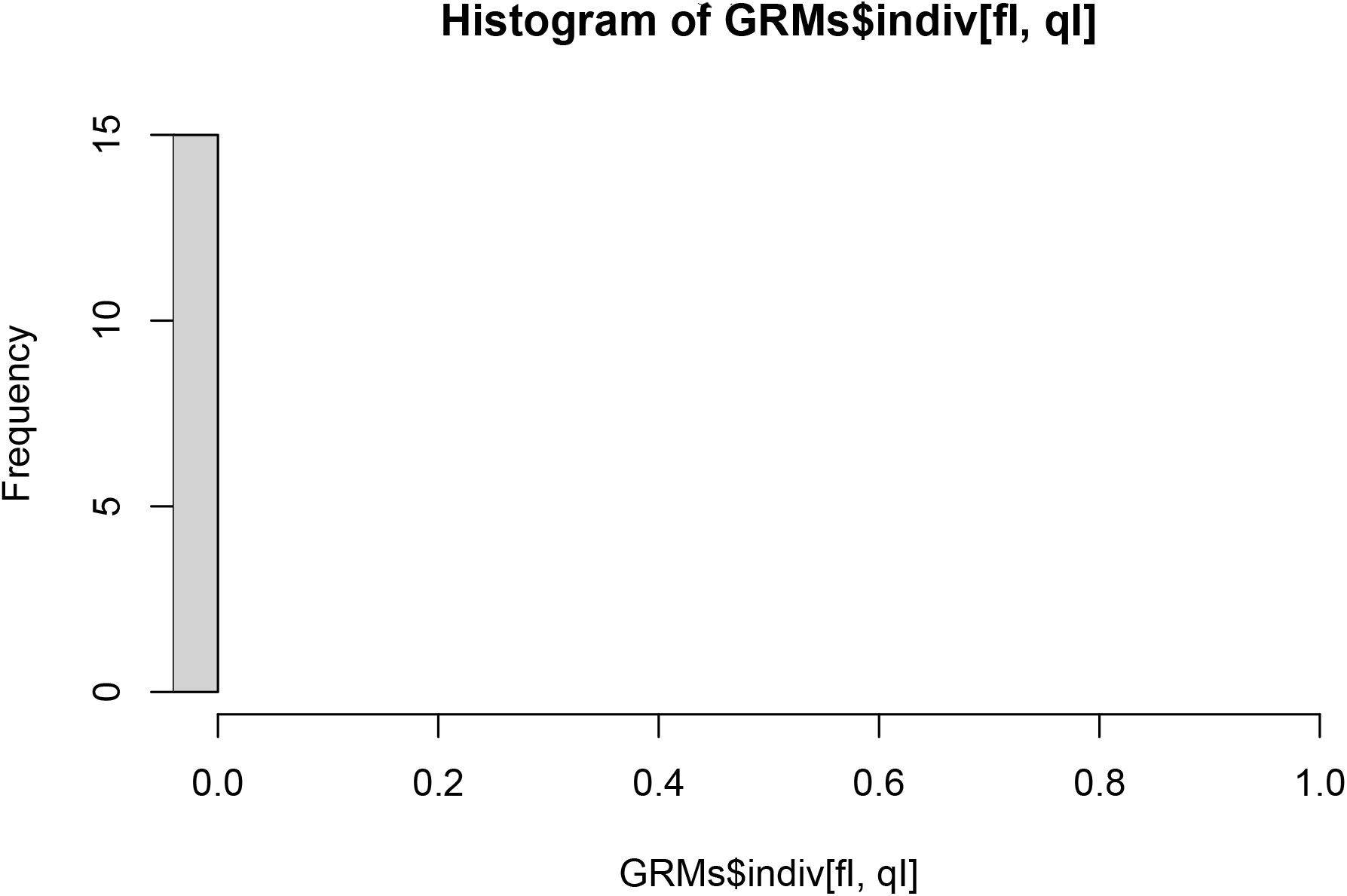

Queen and workers:

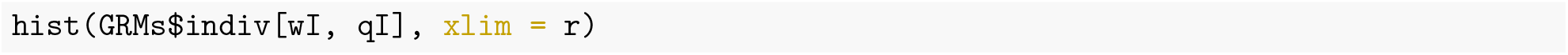

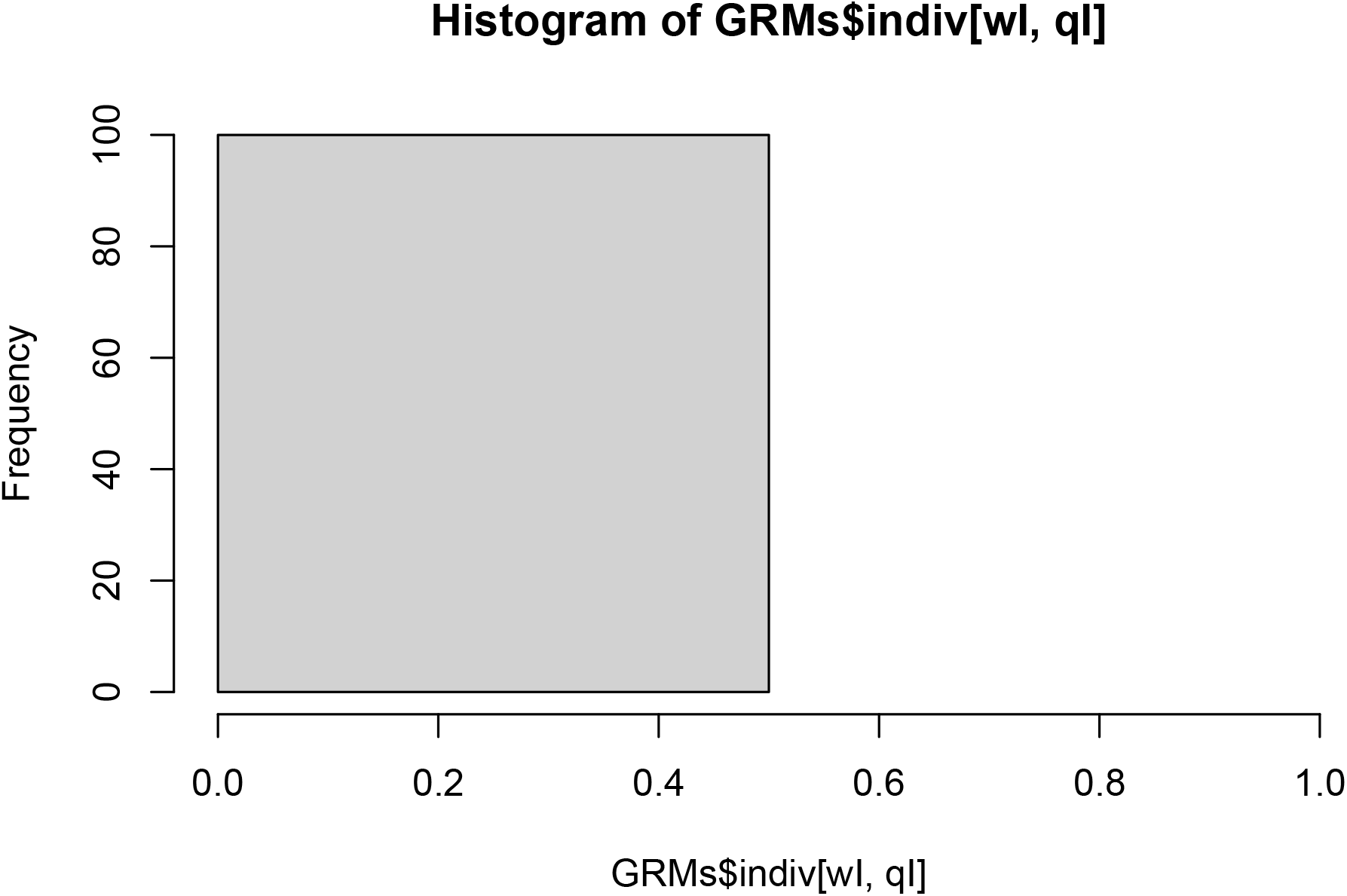

Queen and drones:

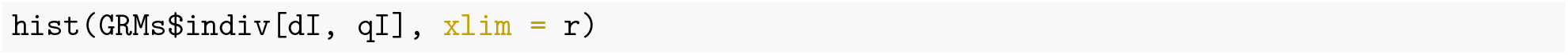

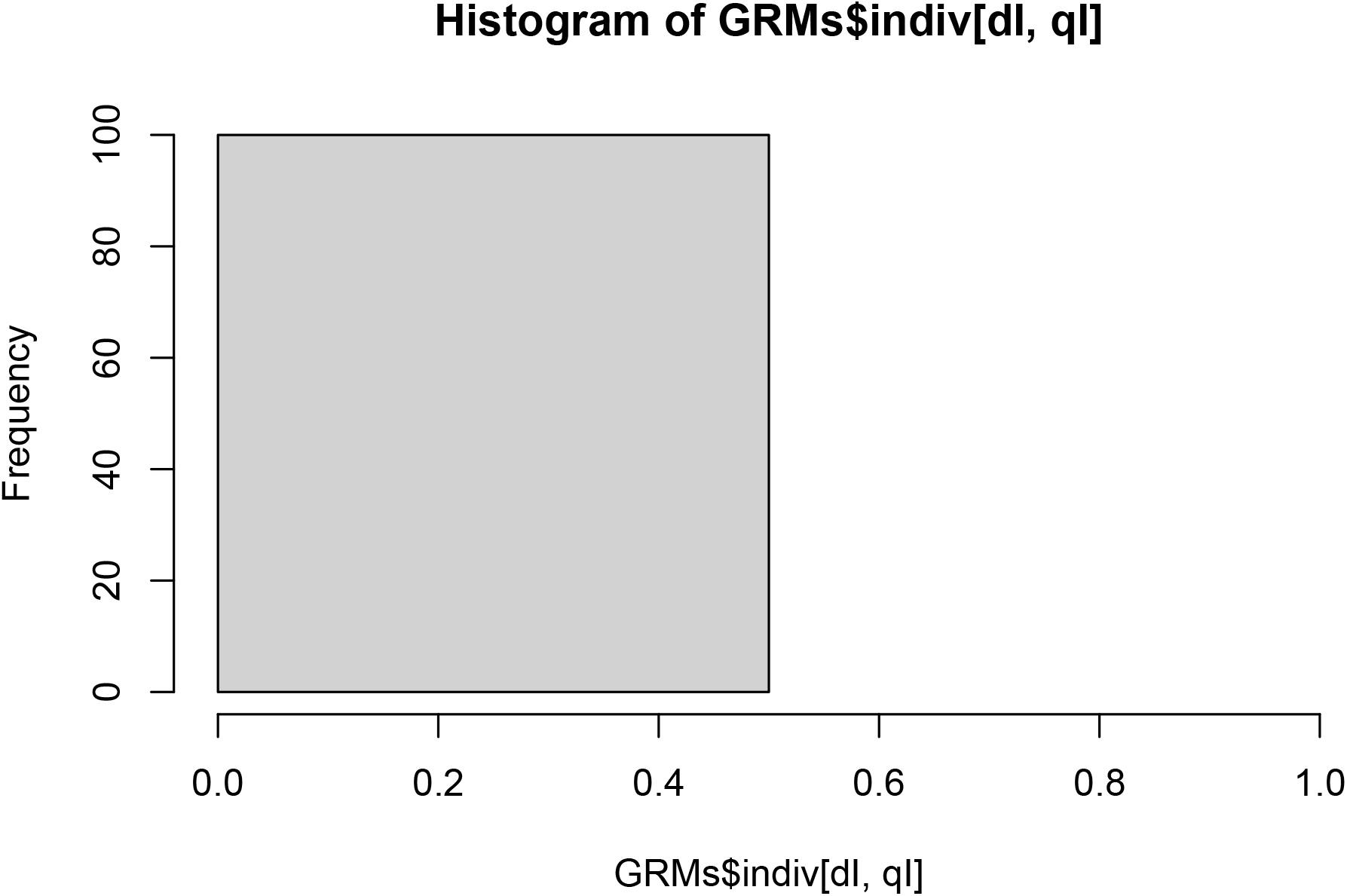

## Additional file 6 – Quantitative genetics vignette

### Introduction

This vignette describes and demonstrates how SIMplyBee implements quantitative genetics principles for honeybees. Specifically, it describes three different examples where we simulate:

1. Honey yield – a single colony trait,
2. Honey yield and Calmness – two colony traits, and
3. Colony strength and Honey yield – two colony traits where one trait impacts the other one via the number of workers.

We start by loading SIMplyBee and quickly simulating genomes for some founder honeybees. Specifically, we will simulate genomes for 20 individuals with 16 chromosomes and 1000 segregating sites per chromosome.

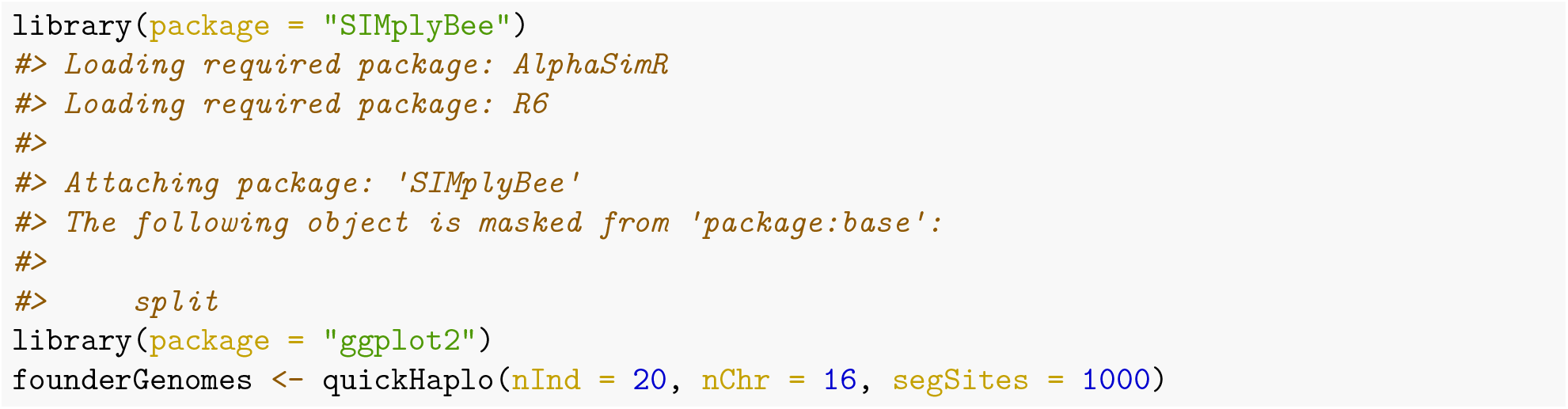

### Honey yield

This section shows how to simulate one colony trait, honey yield, that is influenced by the queen and workers as well as the environment. We will achieve this by:

a. setting base population quantitative genetic parameters,
b. inspecting individual values in the base population,
c. inspecting individual values in a colony,
d. calculating colony value,
e. calculating multi-colony values, and
f. selecting on colony values.

### Base population quantitative genetic parameters

AlphaSimR, and hence SIMplyBee, simulates each individual with its corresponding genome, and quantitative genetic and phenotypic values. To enable this simulation, we must set base population quantitative genetic parameters for the traits of interest in the global simulation parameters via `SimParamBee`. We must set:

1. the number of traits,
2. the number of quantitative trait loci (QTL) that affect the traits,
3. the distribution of QTL effects,
4. trait means, and
5. trait genetic and environmental variances – if we simulate multiple traits, we must also specify genetic and environmental covariances between the traits.

In honeybees, the majority of traits are influenced by the queen and workers. There are many biological mechanisms for these queen and workers effects. Depending on which caste is the main driver of the trait (the queen or workers), we also talk about direct and indirect effects. For example, for honey yield, workers directly affect honey yield by foraging, while the queen indirectly affects honey yield by stimulating workers via pheromone production. The queen and workers effects for a trait can be genetically and environmentally independent or correlated (usually negatively).

Here, we will simulate two traits to represent the queen and workers effects on honey yield. From this point onward we will use the terms the queen effect and queen trait interchangeably. The same applies to workers effect and workers trait. These two effects (=traits) will give rise to honey yield trait. We will assume that colony honey yield is approximately normally distributed with the mean of 20 kg and variance of 4 *kg*^2^, which implies that most colonies will have honey yield between 14 kg and 26 kg (see `hist(rnorm(n = 1000, mean = 20, sd = sqrt(4)))`). Traits like honey yield have a complex polygenic genetic architecture, so we will assume that this trait is influenced by 100 QTL per chromosome (with 16 chromosomes, this gives us 1600 QTL in total).

We will first initiate global simulation parameters and set the mean of queen effects to 10 kg with genetic variance of 1 *kg*^2^, while we will set the mean of workers effects to 10 kg with genetic variance of 1 *kg*^2^. The mean and the variance for the worker effect are proportionally scaled by the expected number of workers in a colony. The mean and variance for the queen effect is assumed larger than for the workers effect, because there is one queen and many workers in colony and we assume that workers effects “accumulate”. Deciding how to split the colony mean between queen and workers effects will depend on the individual to colony mapping function, which we will describe in the Colony value sub-section.

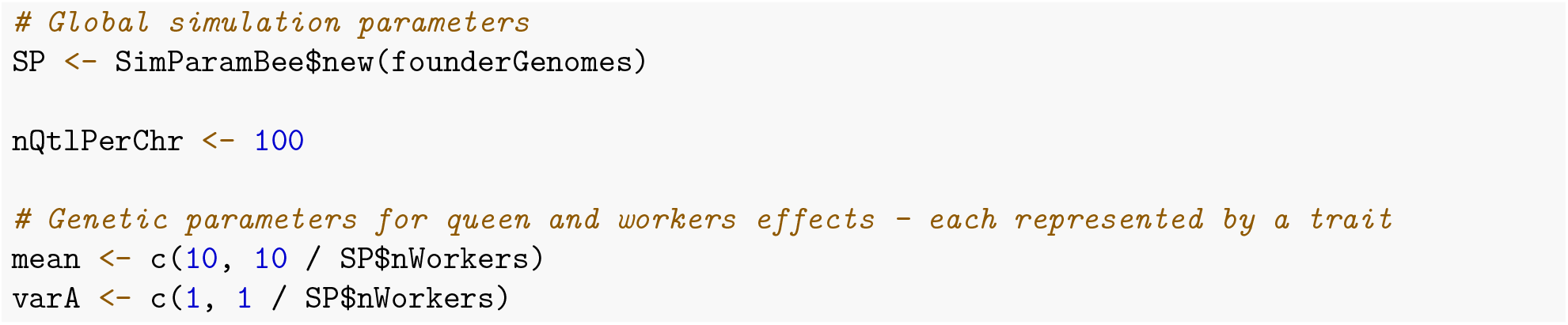

We next set genetic correlation between the queen and workers effects to – 0.5 to reflect the commonly observed antagonistic relationship between these effects. With all the quantitative genetic parameters defined, we now add two additive traits to global simulation parameters and name them `queenTrait` and `workerTrait`. These parameters drive the simulation of QTL effects. Read about all the other trait simulation options in AlphaSimR via: `vignette(topic = “traits”, package=“AlphaSimR”)`.

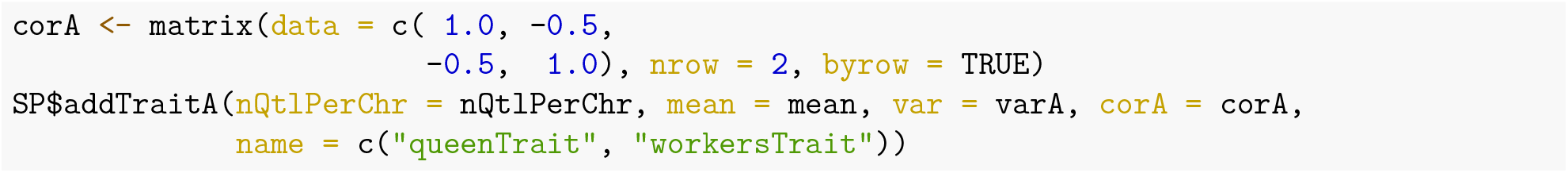

Finally, we set the environmental variance of the queen and workers effects to 3 *kg*^2^ and we again scale the worker variance by the expected number of workers. Contrary to the negative genetic correlation, we here assume that environmental correlation between the queen and workers effects is slightly positive, 0.3. This is just an example! These parameters should be based on literature or simulation scenarios of interest.

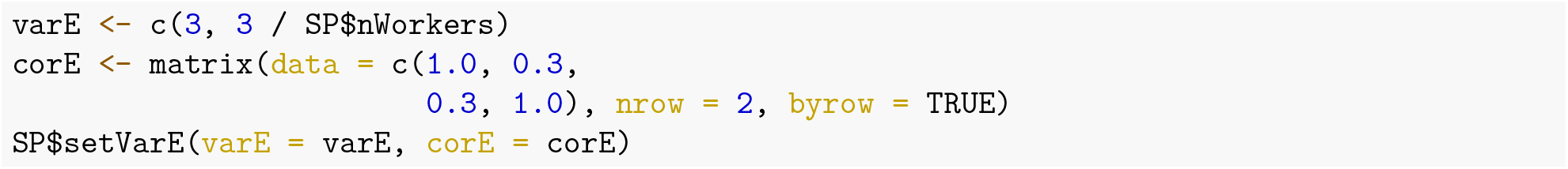

### Individual values in the base population

Now we create a base population of virgin queens. Since we defined two traits, all honeybees in the simulation will have genetic and phenotypic values for both traits. The genetic values are stored in the `gv` slot of each `Pop` object, while phenotypic values are stored in the `pheno` slot.

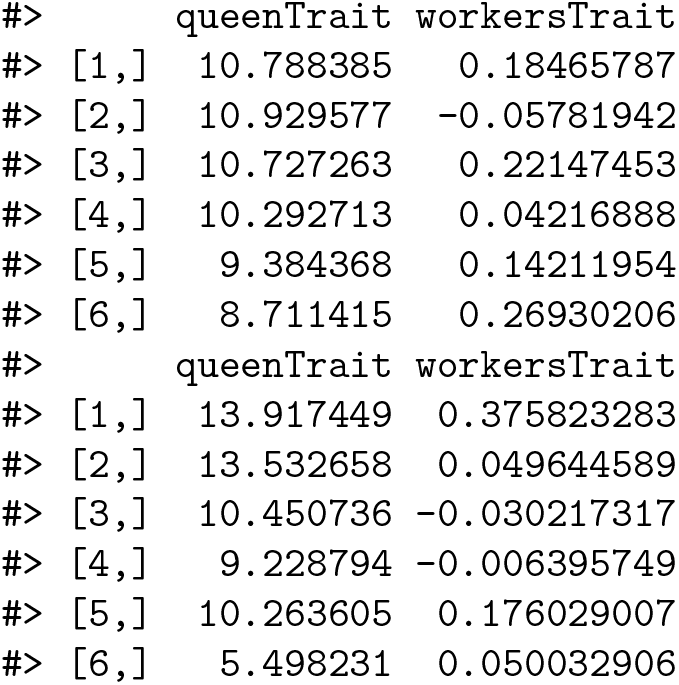

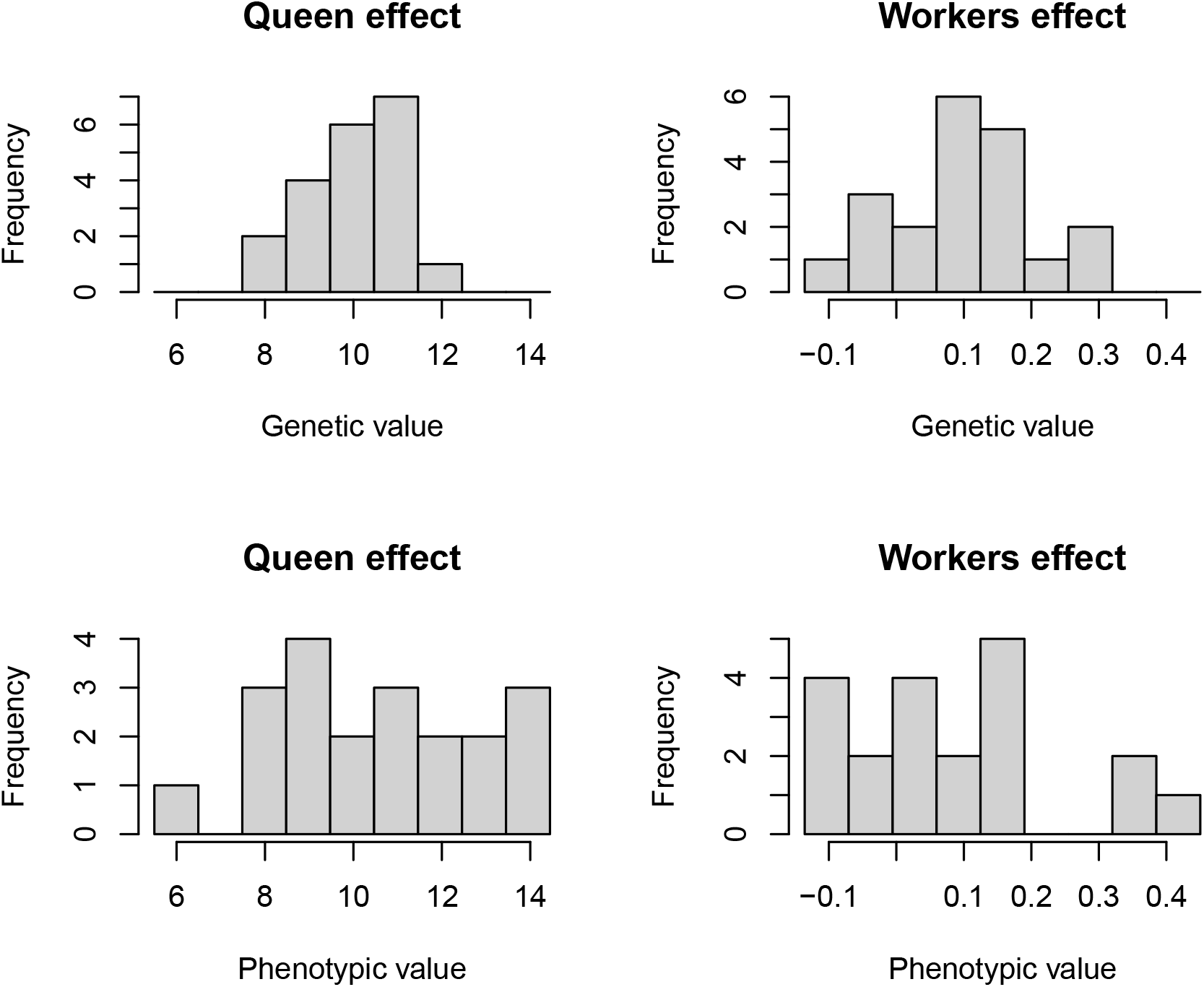

Note that these are virgin queens, yet we obtained queen and workers effect values for them! Is this wrong? No! Virgin queens carry DNA with genes that are differentially expressed in different castes, which would be only showed in their phenotype. Hence, virgin queens have genetic values for the queen and worker effects, but they might never actually express these effects. In this simulation virgin queens also obtained phenotypic values for both of the effects. This is technically incorrect because virgin queens don’t express genes for the worker effect at all, and they also do not express the queen effect, not until they become the queen of a colony. We can treat these phenotypic values for virgin queens as values that we could see if these virgin queens would express these traits. We will show later in the Colony value sub-section how we use these traits from different castes. If existence of these phenotypic values for certain castes is a hindrance, we can always remove them for population or colony objects by modifying the corresponding slots as required.

As with the virgin queens, drones also carry DNA with genes that are expressed in different castes. Therefore, drones will also have the queen and workers effect genetic (and phenotypic values) for honey yield even though they do not contribute to this trait in a colony.

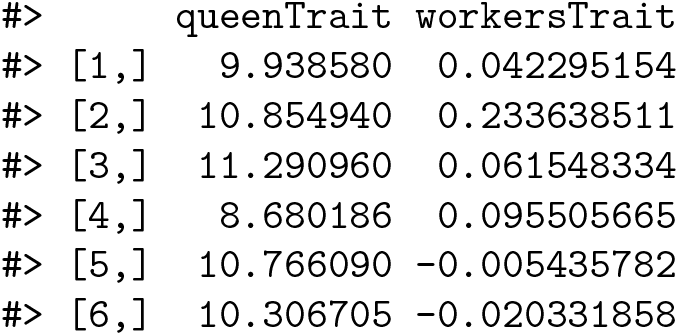

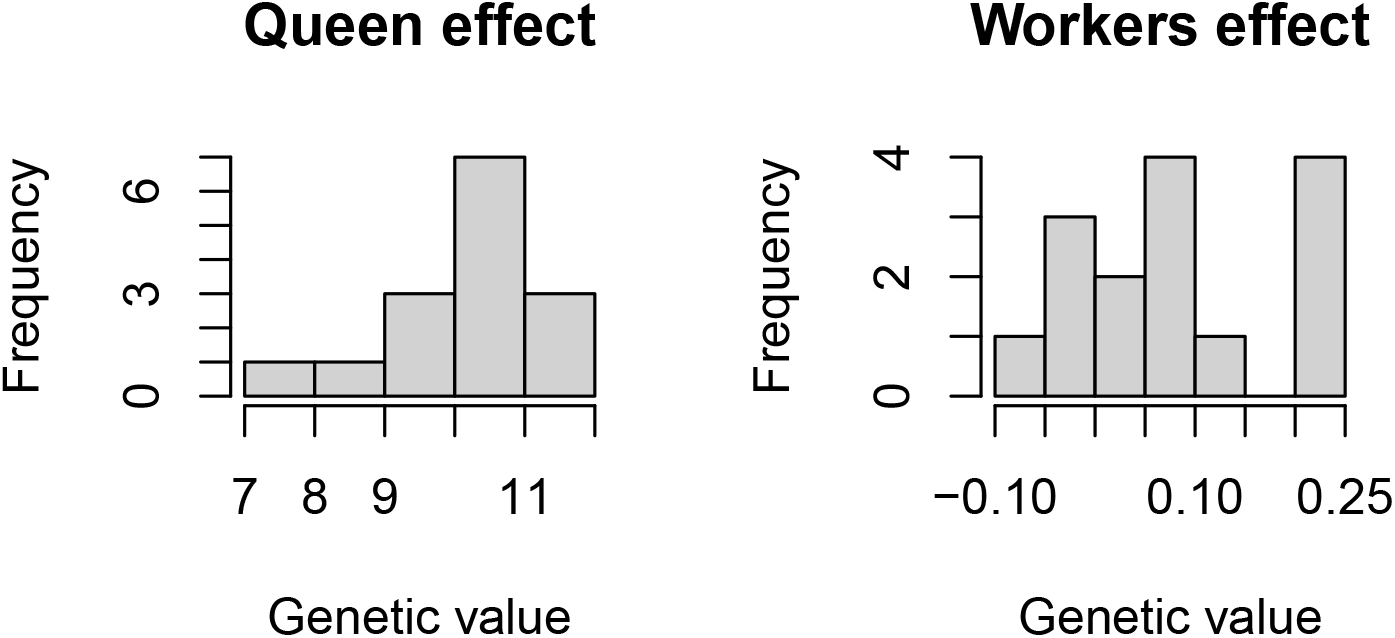

### Individual values in a colony

We continue by creating a colony from one base population virgin queen, crossing it, and adding some workers.

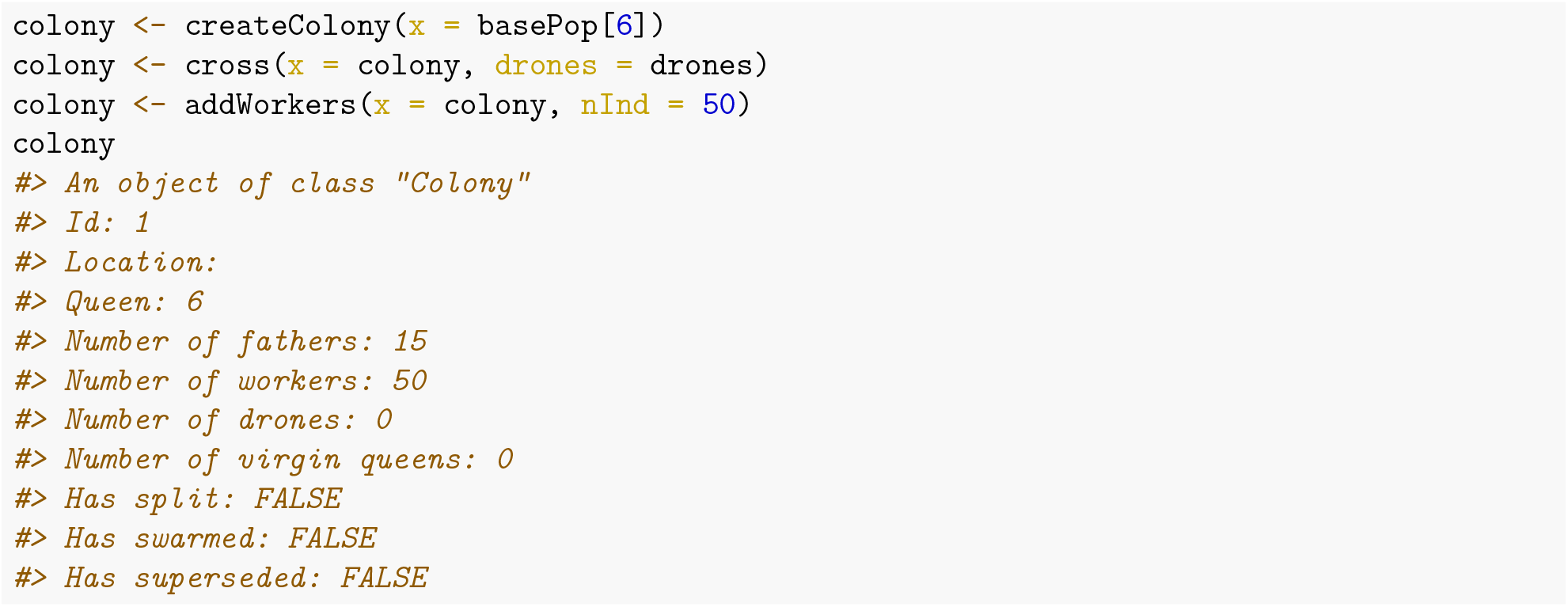

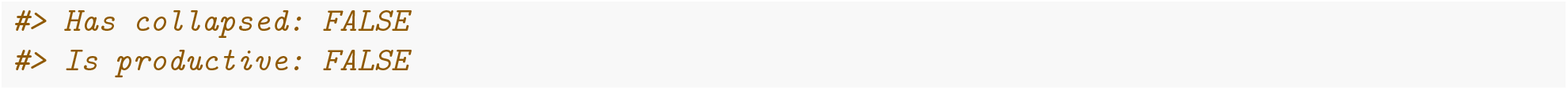

We can access the genetic and phenotypic values of colony members with functions `getGv()` and `getPheno()`, both of which have the `caste` argument (see more via `help(getGv)`).

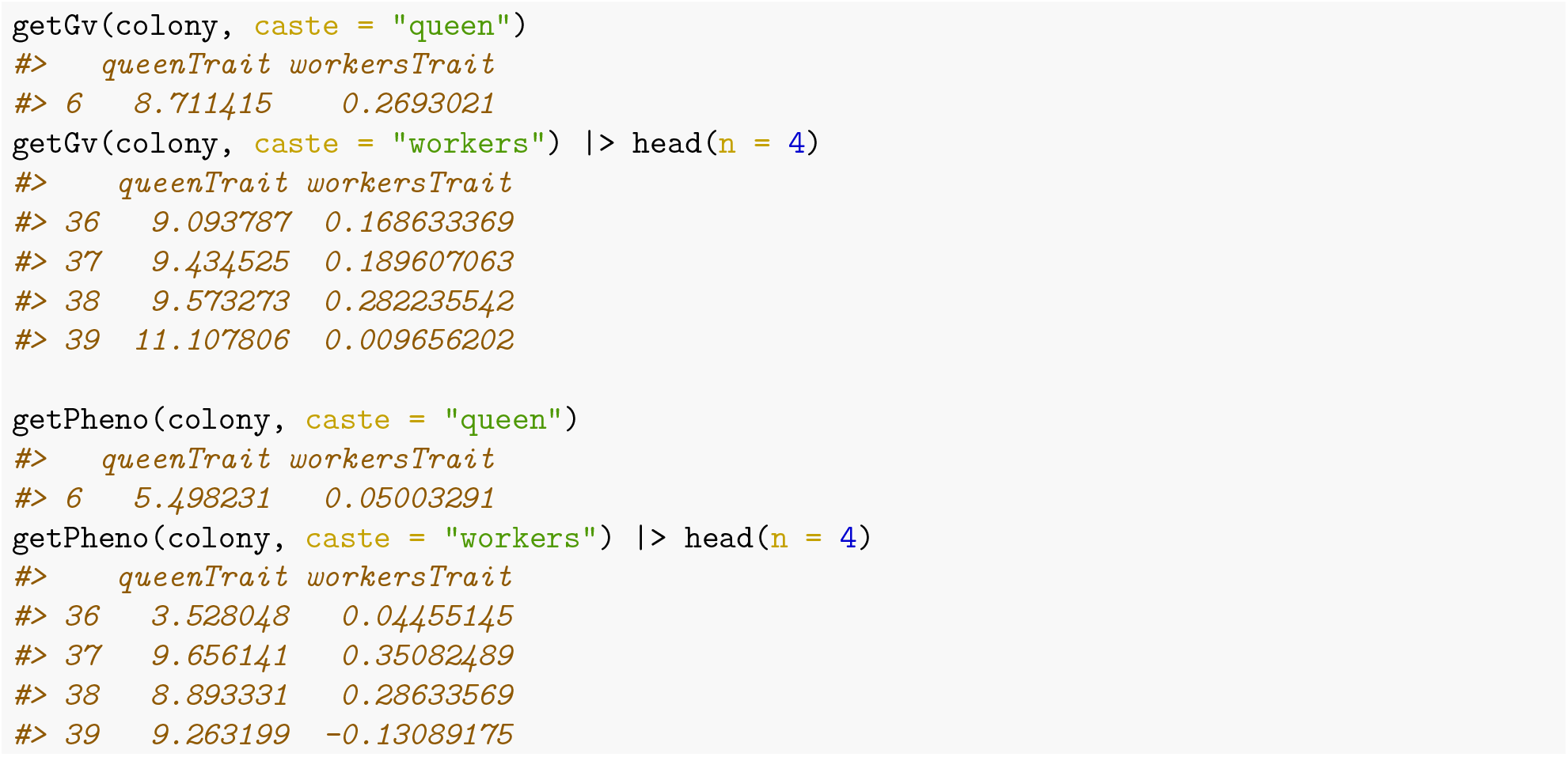

For convenience, there are also alias functions for accessing the genetic and phenotypic values of each caste directly.

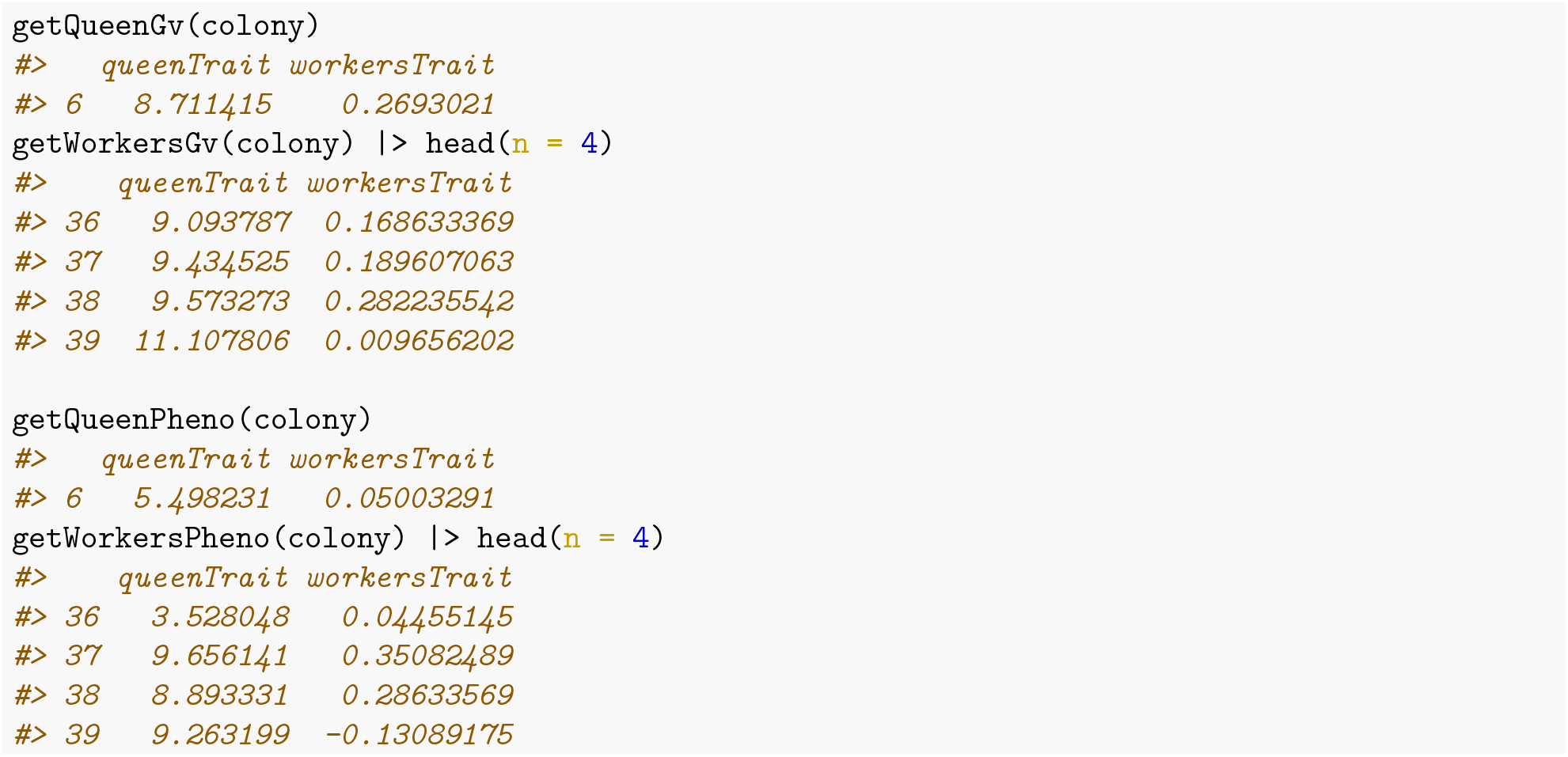

Some phenotypes, such as honey yield, are only expressed if colony is at full size. This is achieved by the `buildUp()` colony event function that adds worker and drones and hence turns on the `production` status of the colony (to `TRUE`). SIMplyBee includes a function `ìsProductive()` to check the production status of a colony.

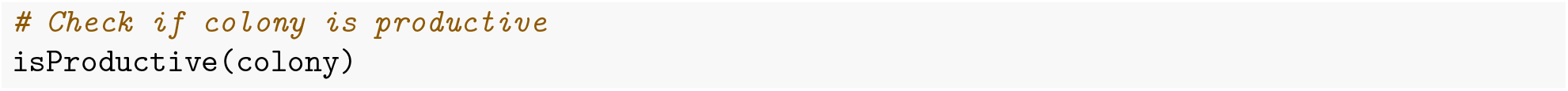

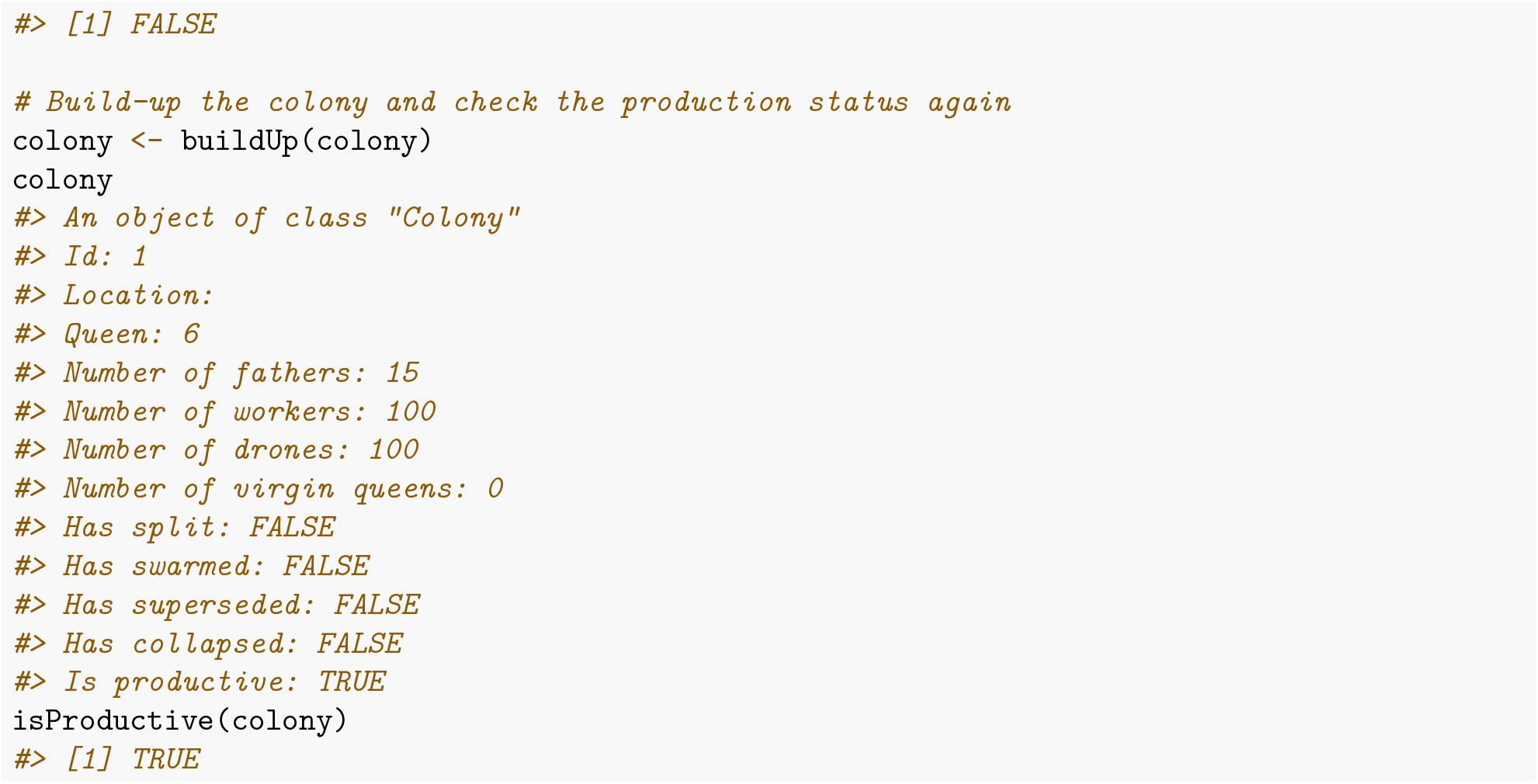

For the ease of further demonstration, we now combine workers’ values into a single data.frame.

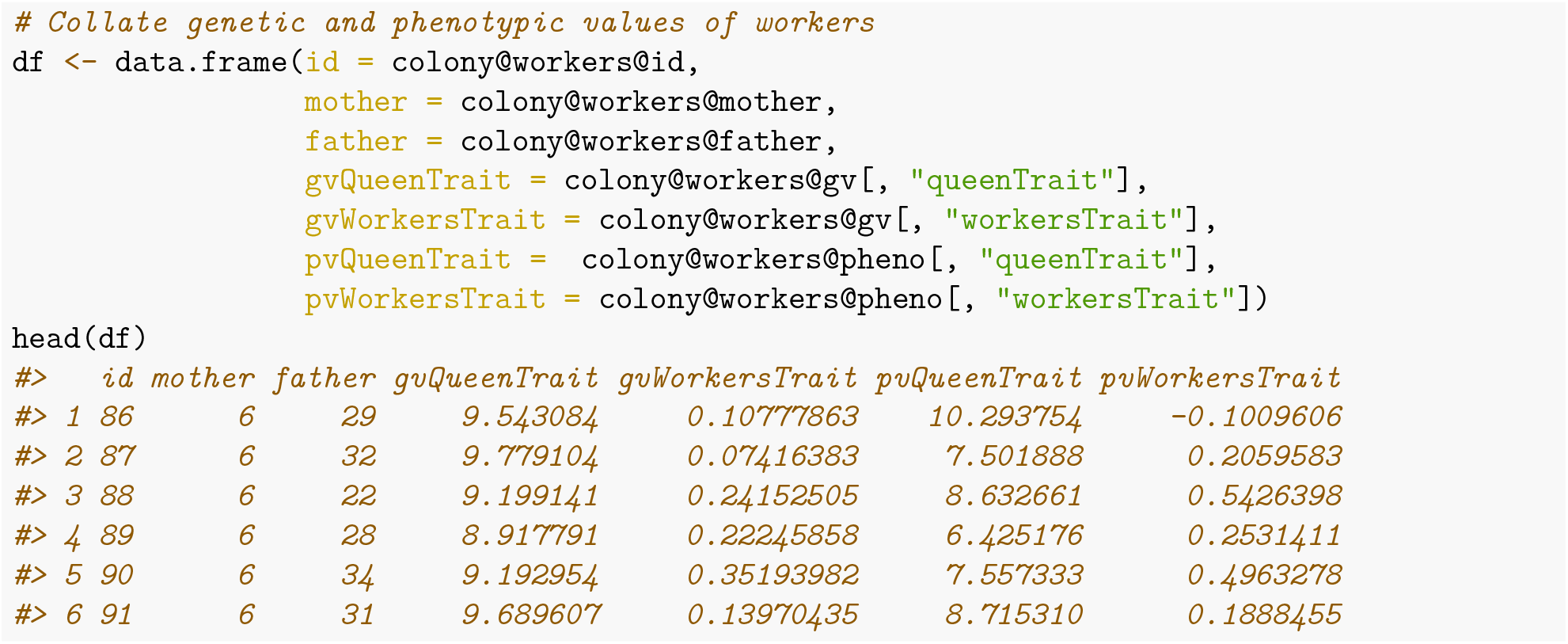

To visualise correlation between queen and workers effects in workers, we plot these effect values against each other.

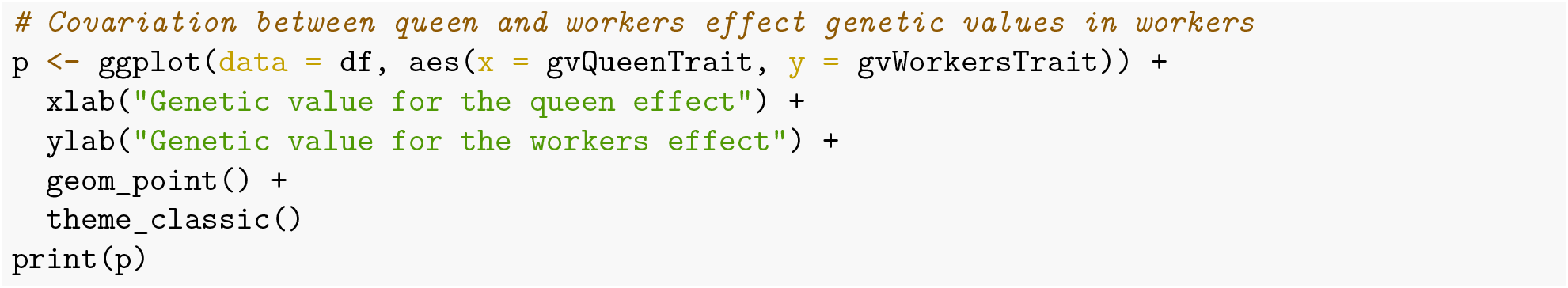

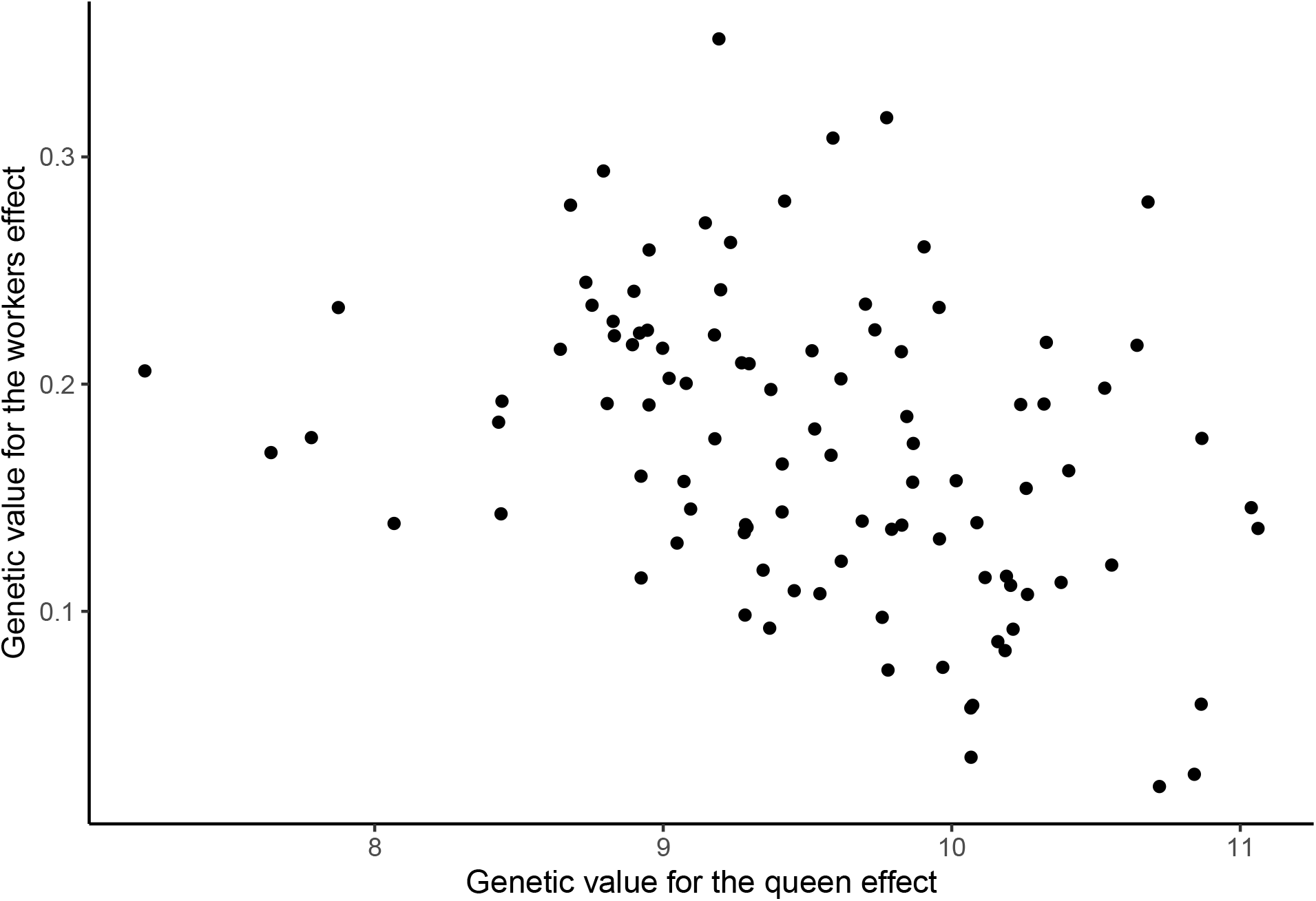

In SIMplyBee, we know genetic values of all individuals, including drones that the queen mated with (=fathers in a colony)!

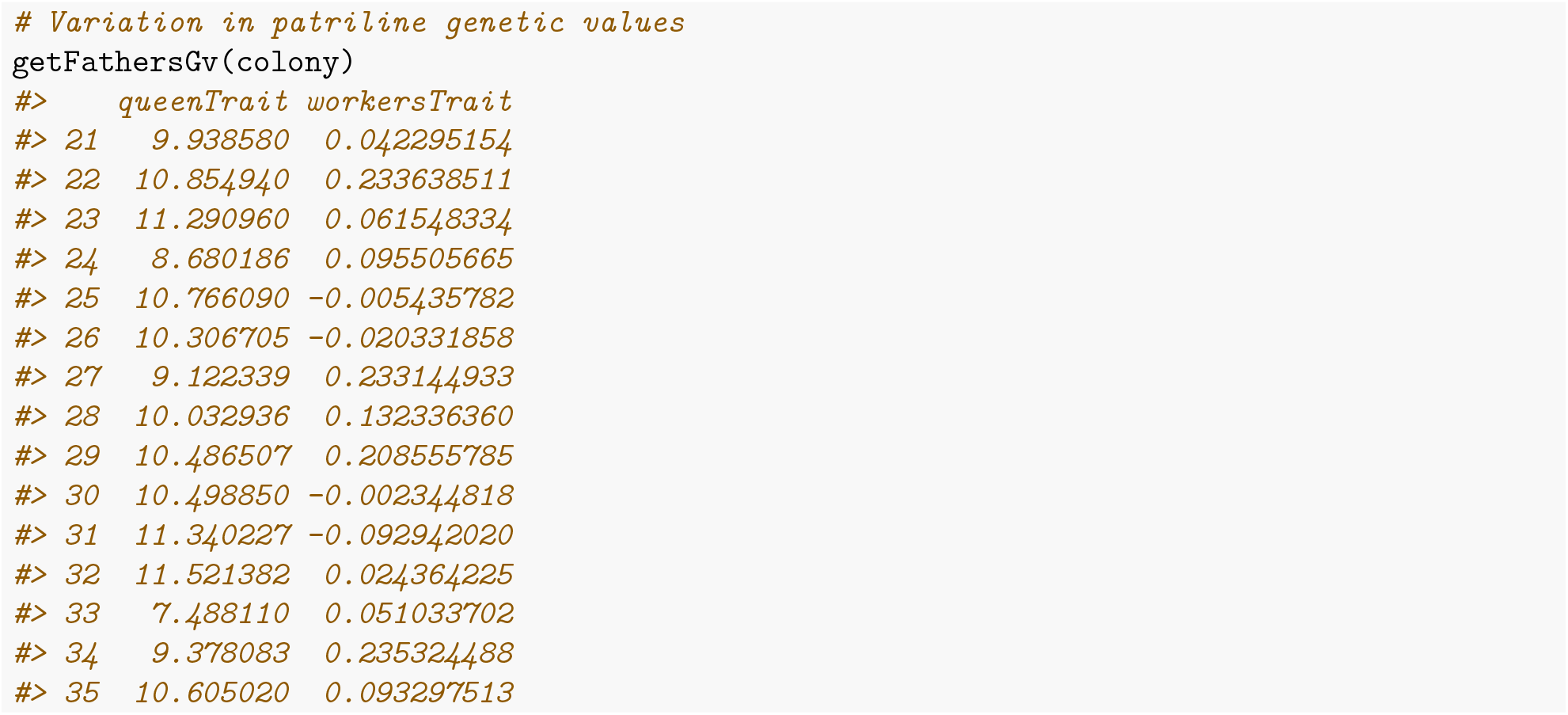

Knowing the father of each worker, we inspect variation in the distribution of genetic values of worker by the patriline (workers from a single father drone) for the workers effect.

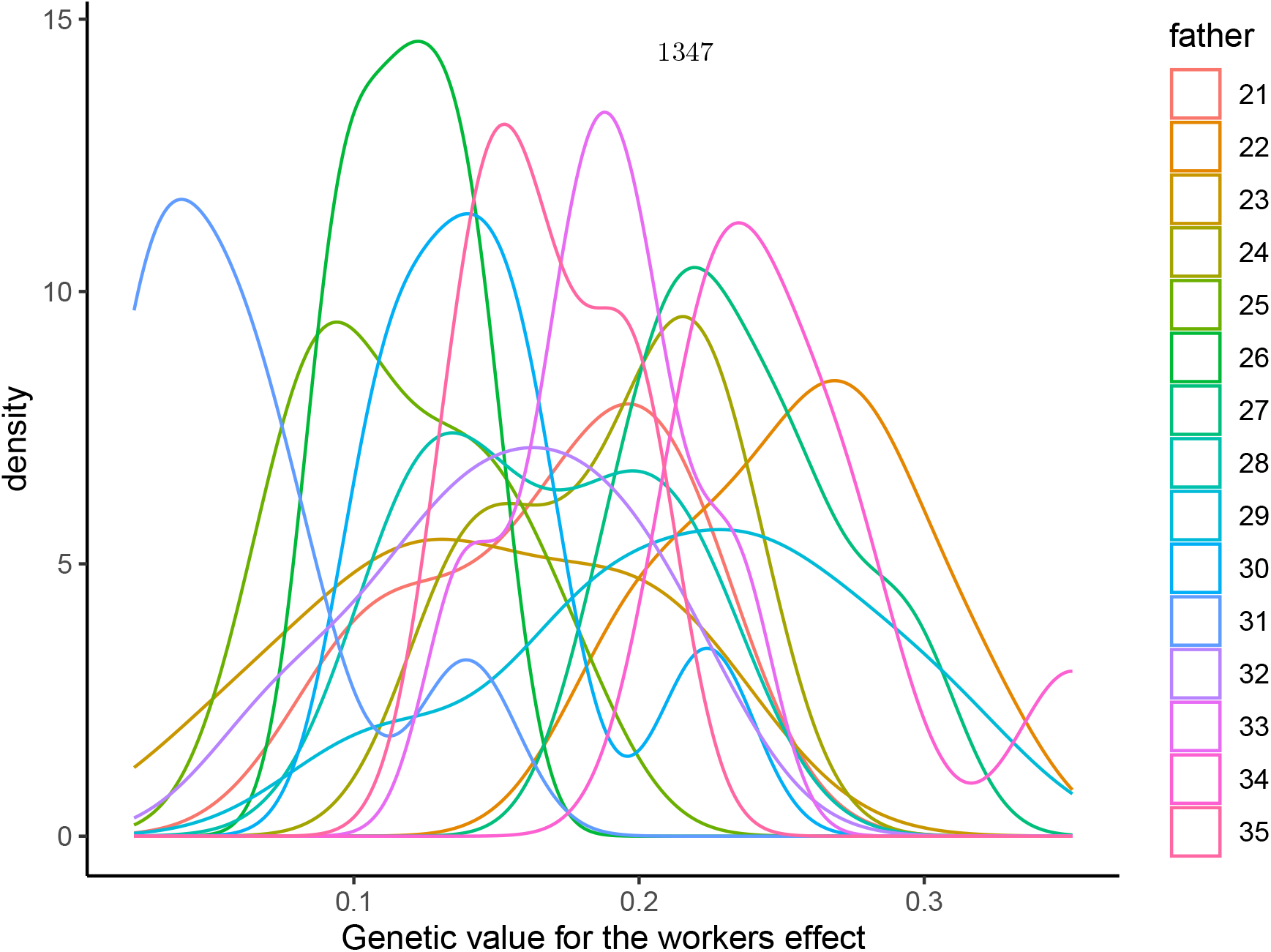

### Colony value

However, in honeybees we usually don’t observe values on individuals, but on a colony. SIMplyBee provides functions for mapping individual values to a colony value. The general function for this is `calcColonyValue()`, which can combine any value and trait from any caste. There are also aliases `calcColonyGv()` and `calcColonyPheno()`. These functions require users to specify the so-called mapping function (via the FUN argument). The mapping function specifies queen and workers traits (potentially also drone traits) and what function we want to apply to each of them before mapping them to the colony value(s). We can also specify whether the colony value(s) depend on the production status. For example, if a colony is not productive, its honey yield would be 0 or unobserved. SIMplyBee provides a general mapping function `mapCasteToColonyValue()` and aliases `mapCasteToColonyGv()` and `mapCasteToColonyPheno()`. These functions have arguments to cater for various situations. By default, they first calculate caste values: leave the queen’s value as it is, sum workers’ values, potentially sum drones’ values, and lastly sum all these caste values together into a colony value. Users can provide their own mapping function(s) too!

We now calculate honey yield for our colony – a single value for the colony.

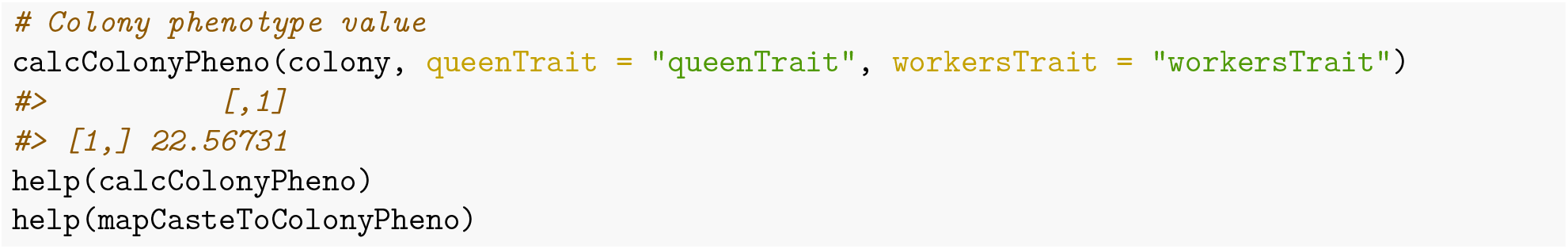

These colony values are not stored in a colony, because they change as colony changes due to various events. For example, reducing the number of workers will reduce the colony honey yield.

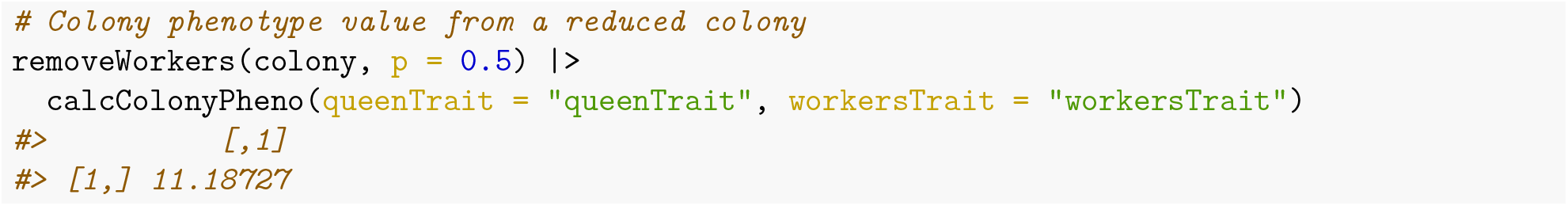

Please note that we assumed that the queen contributes half to colony honey yield and workers contribute the other half. This means that removing workers will still give a non-zero honey yield! This shows that we have to design the mapping between individual, caste, and colony values with care!

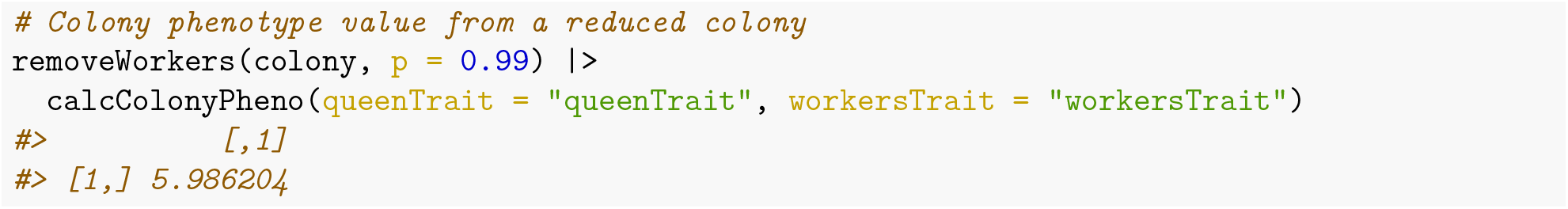

Finally, note that SIMplyBee currently does not provide functionality for breeding values, dominance deviations, and epistatic deviations at caste and colony levels, despite the availabiliy of AlphaSimR `bv()`, `dd()`, and `aa()` functions. This is because we have to check or develop theory on how to calculate these values across active colonies and hence we currently advise against the use of AlphaSimR `bv()`, `dd()`, and `aa()` functions with SIMplyBee as the output of these functions could be easily misinterpreted.

### MultiColony values

The same functions can be used on a `MultiColony` class object. Let’s create an apiary.

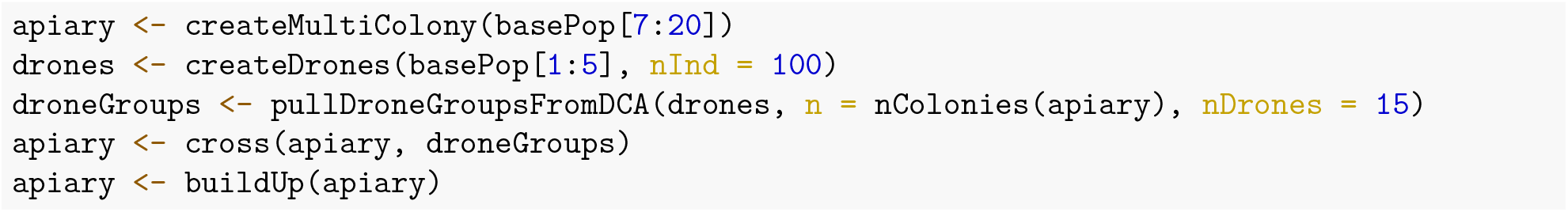

We can extract the genetic and phenotypic values from multiple colonies in the same manner as from a single colony, by using `get*Gv()` and `get*Pheno()` functions. The output of these function is a named list with values for each colony or a single matrix if we set the `collapse` argument to `TRUE`.

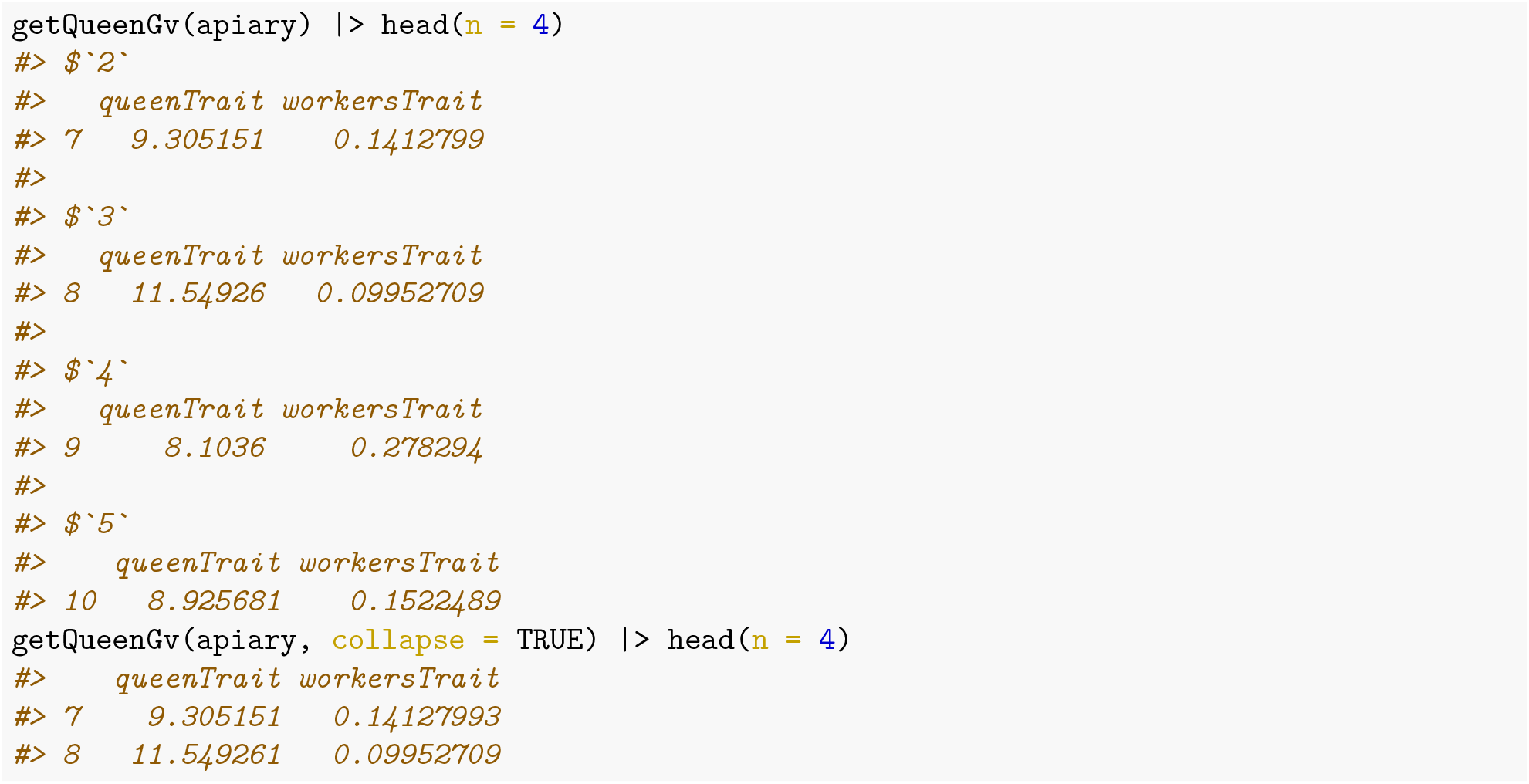

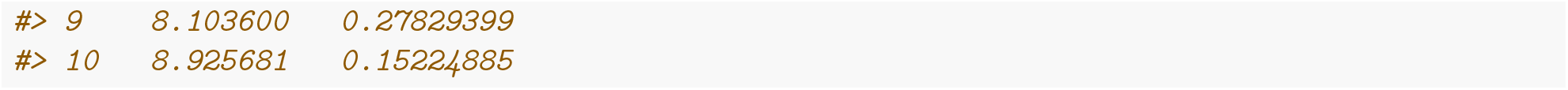

In a similar manner, we can calculate colony value for all the colonies in our apiary, where the row names of the output represent colony IDs.

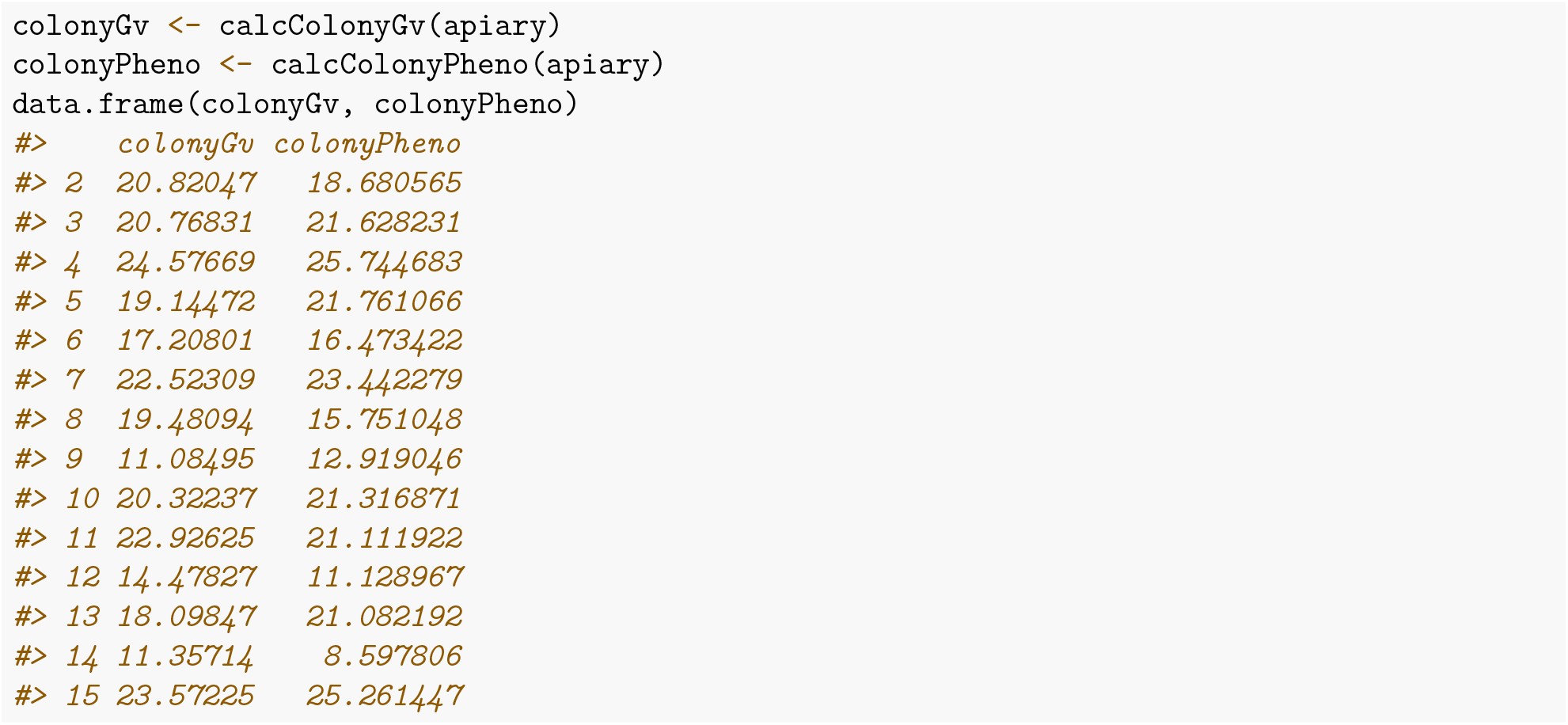

### Selection on colony values

Since the aim of selection is to select the best individuals or colonies for the reproduction, we could select the best colony in our apiary based on either genetic or phenotypic value for grafting the new generation of virgin queens. We can use the function `selectColonies()` that takes a matrix of colony values (the output of `calcColonyValue()` function). The default behavior is to select the colonies with the highest value (argument `selectTop` set to `TRUE`), but you can also select the colonies with the lowest values (argument `selectTop` set to `FALSE`).

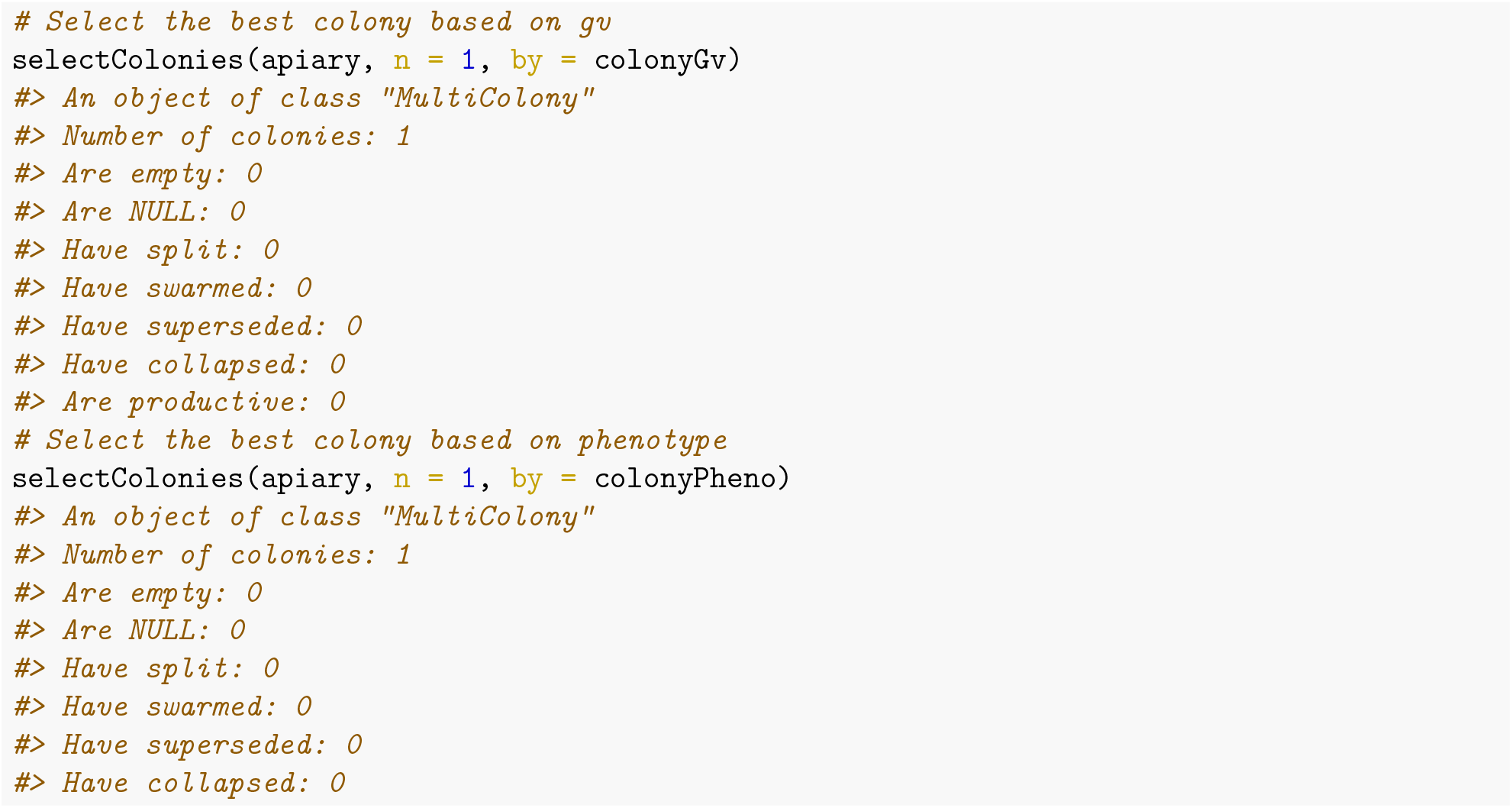

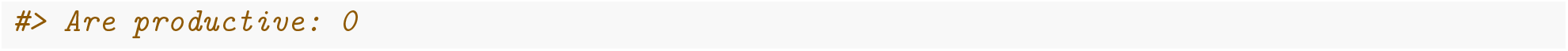

The same functionality is implemented in `pullColonies()` and `removeColonies()`.

### Honey yield and Calmness

In this section we expand simulation to two uncorrelated colony traits with queen and workers effects, honey yield and calmness. We follow the same recipe as in the previous section where we simulated only one colony trait.

We first reinitialize the global simulation parameters because we will define new traits. For honey yield we will use the same parameters as before, while for calmness trait we will assume that the trait is scored continuously in such a way that negative values are undesirable and positive values are desirable with zero being population mean. We will further assume the same variances for calmness as for honey yield, and a genetic (and environmental) correlation between the queen and workers effects of −0.4 (and 0.2) for calmness. We assume no genetic or environmental correlation between honey yield and calmness. Beware, this is just an example to show you how to simulate multiple colony traits – we have made up these parameters – please use literature estimates in your simulations!

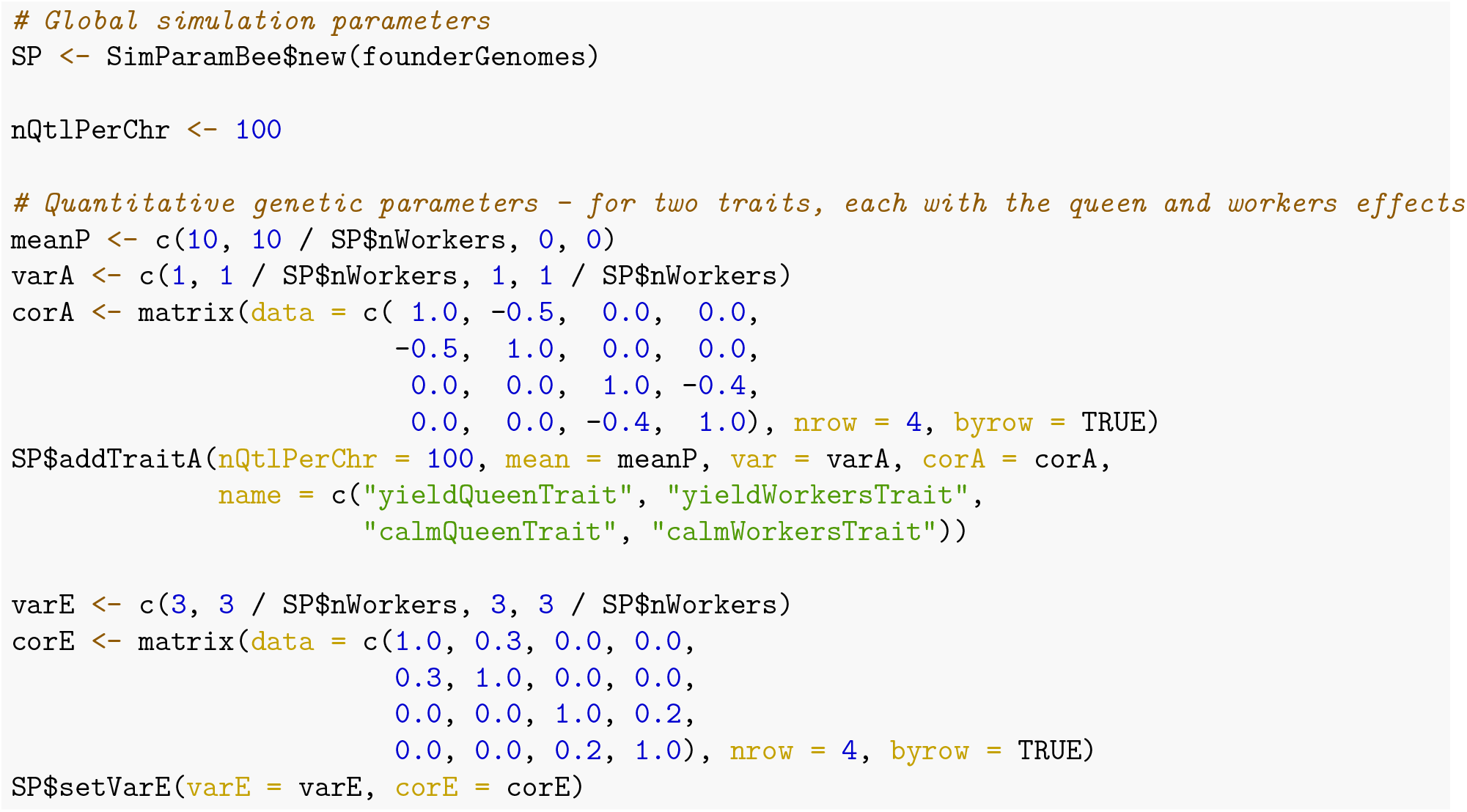

We continue by creating a base population of virgin queens and from them an apiary with 10 full-sized colonies.

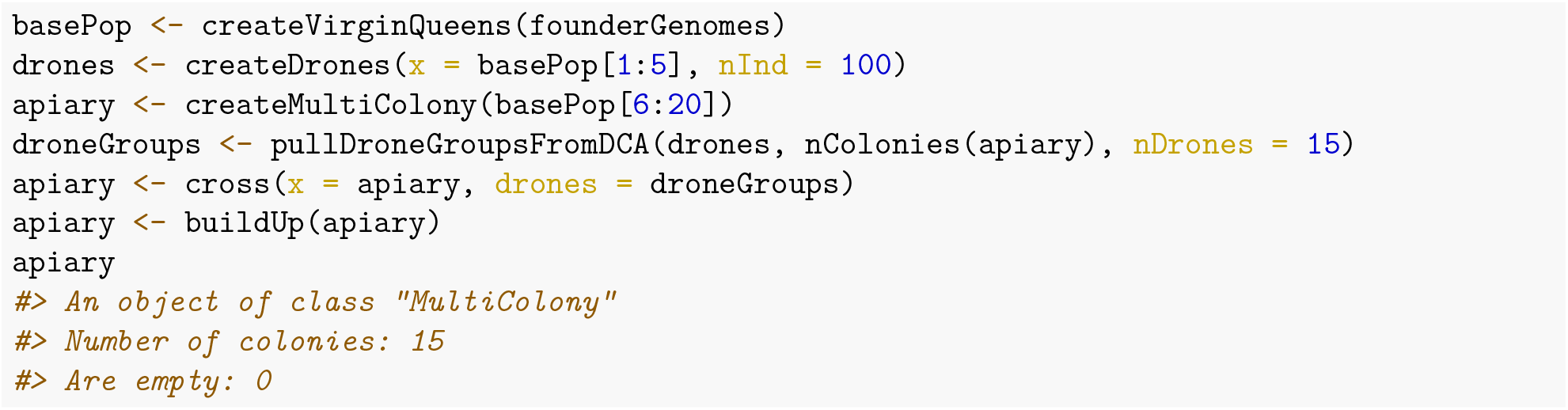

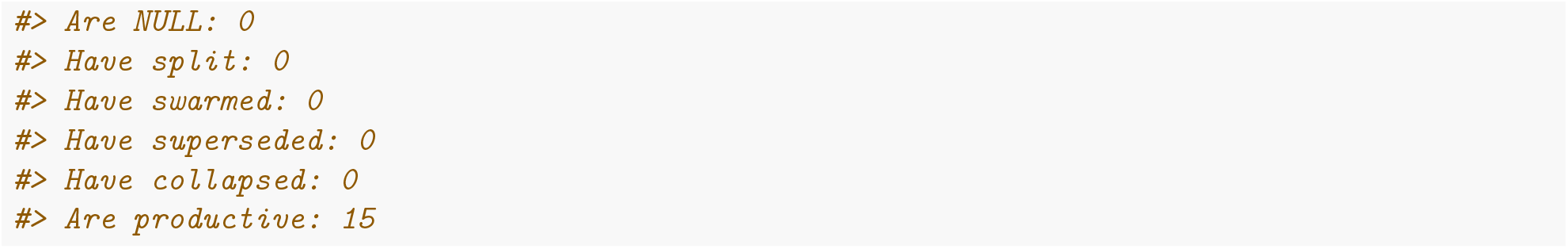

We can again inspect the genetic (and phenotypic) values of all individuals in each colony and whole apiary with `get*Gv()` and `get*Pheno()` functions. Now, the output contains four traits representing the queen and workers effect for honey yield and calmness. These functions also take an `nInd` argument to sample a number of individuals along with their values.

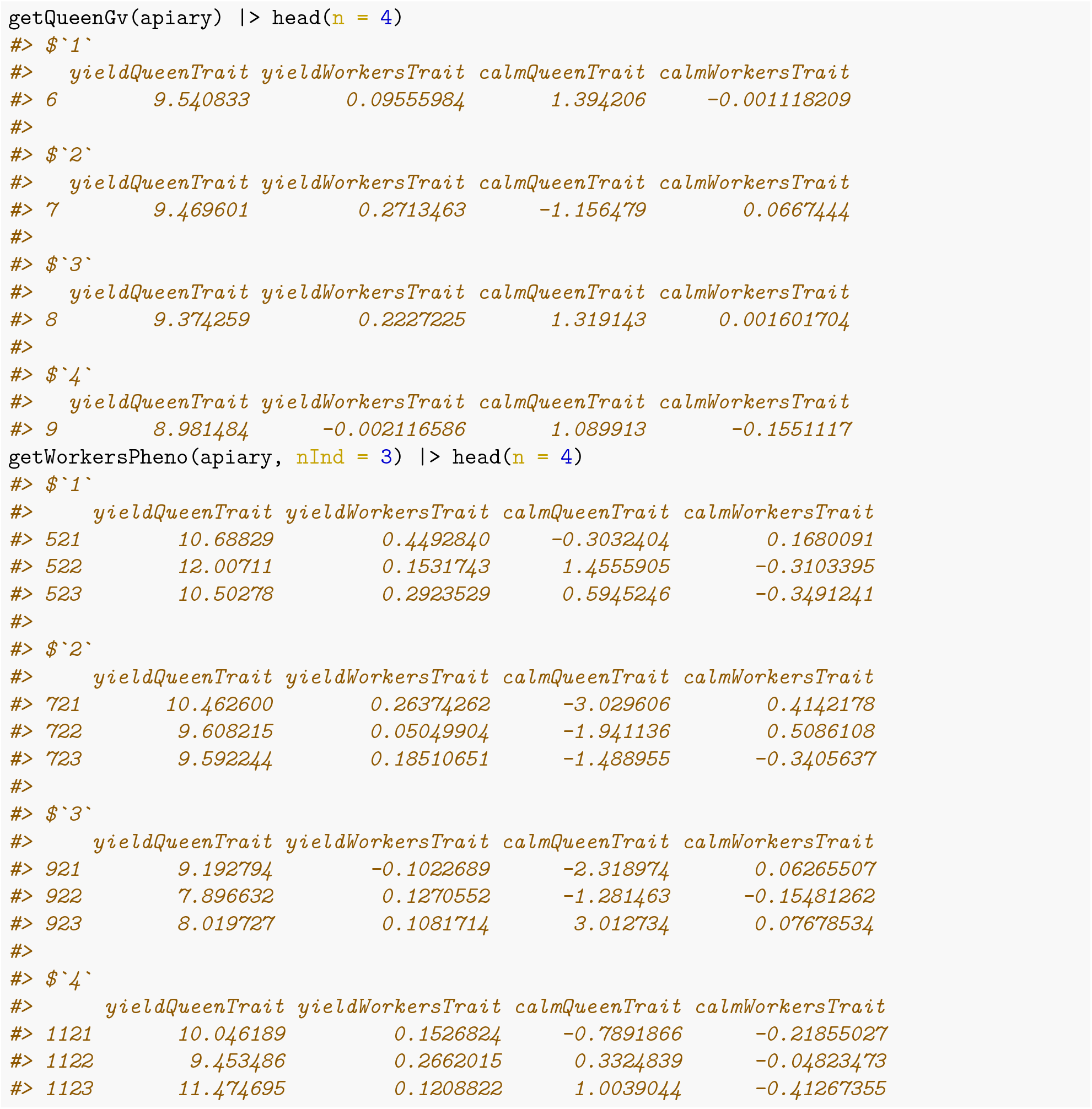

Now, we calculate colony genetic and phenotypic values for all colonies in the apiary. Since we are simulating two traits, honey yield and calmness, we have two ways to calculate corresponding colony values. The first way is to use the default `mapCasteToColony*()` function in `calcColony*()` and only define additional arguments as shown here:

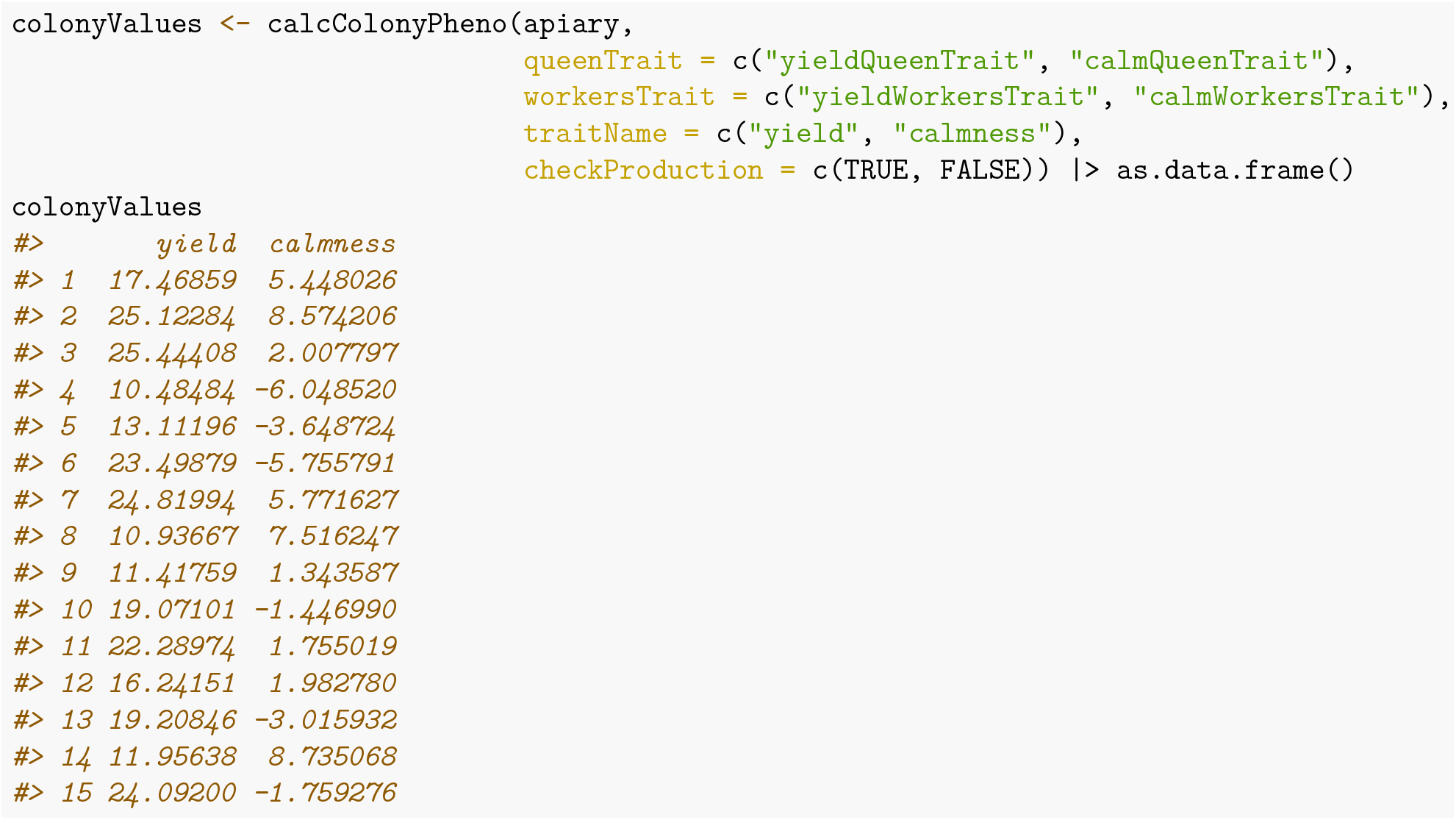

The second way is to create our own mapping function. An equivalent outcome to the above is shown below just to demonstrate use of your own function, but we are simply just reusing `mapCasteToColonyPheno()` twice;)

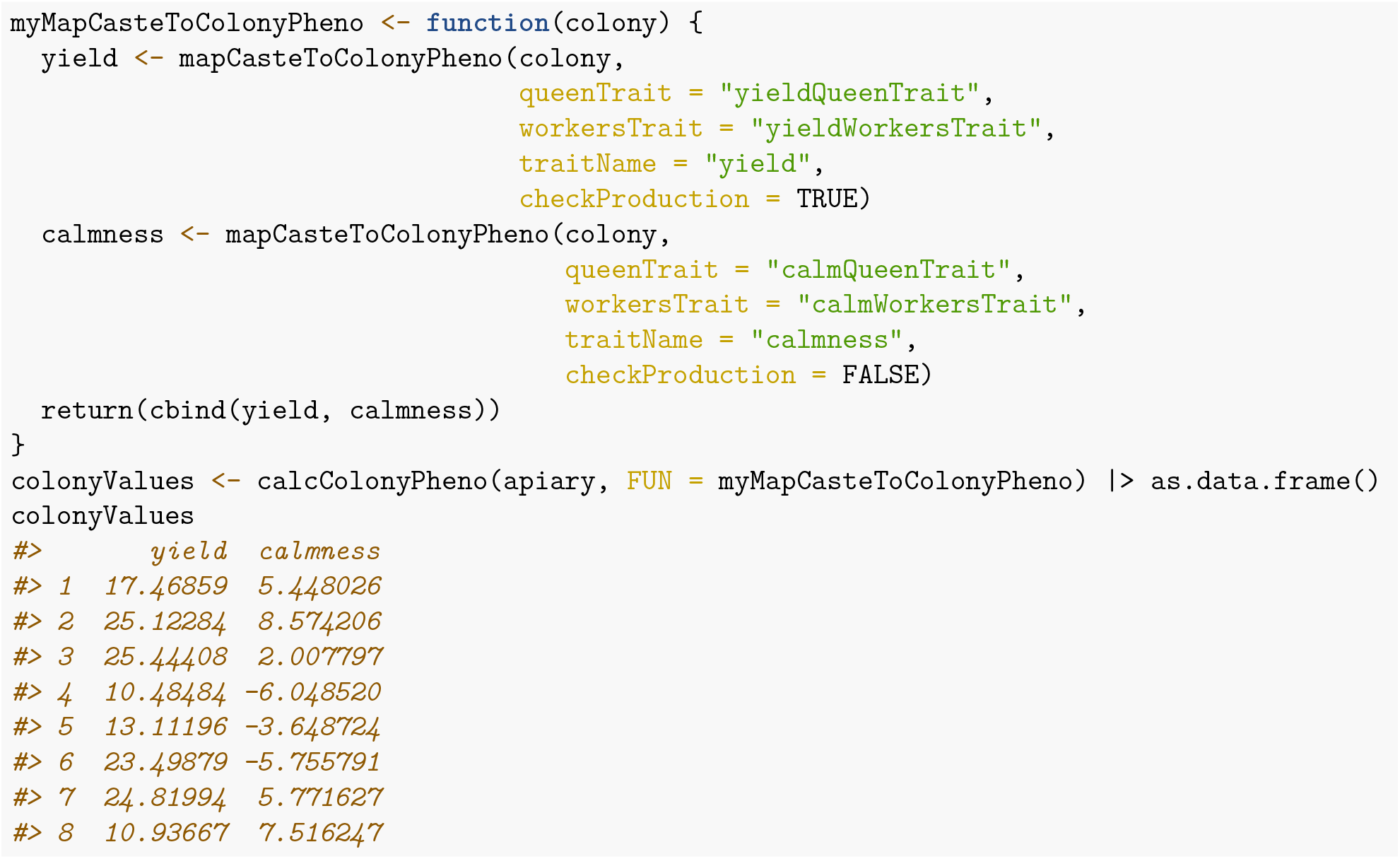

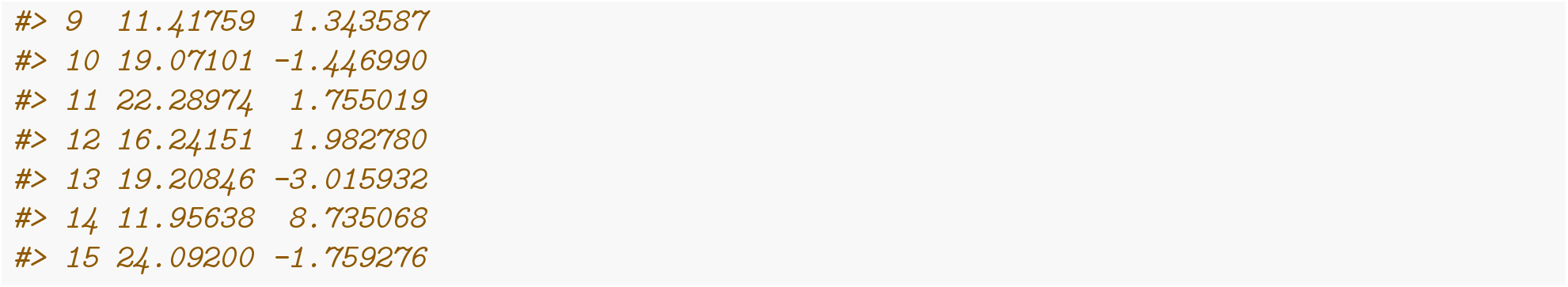

Again, we can now select the best colony based on the best phenotypic value for either yield, calmness, or an index of both. Let’s say that both traits are equally important so we select on a weighted sum of both of them – we will use the AlphaSimR `selIndex()` function that enables this calculation along with scaling. We will represent the index such that it has a mean of 100 and standard deviation of 10 units.

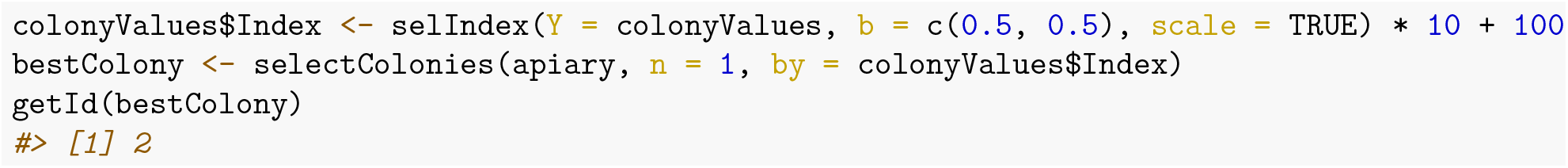

We see that we selected colony with ID “4”, but we would be selecting a different colony based on different selection criteria (yield, calmness, or index).

### Strength and honey yield

In this section we change simulation to two traits where the phenotype realisation of the first trait affects the phenotype realisation of the second trait. Specifically, we will assume that queen’s fecundity, and hence the number of workers, is under the genetic affect of the queen and her environment. Furthermore, we will assume as before that colony honey yield is due to the queen effect and workers effect. Since the value of the workers effect depends on then number of workers, we obtain correlation between fecundity and honey yield, even if these traits would be uncorrelated on the queen level. We emphasise that this is just an example and the biology of these traits might be different.

We follow the same logic as before and simulate three traits that will contribute to two colony traits, queen’s fecundity, that is colony strength, and honey yield. We assume that fecundity is only due to the queen (and not the workers), hence we simulate only the queen effect for this trait. For honey yield we again assume that both the queen and workers contribute to the colony value. For speed of simulation we only simulate 100 workers per colony on average and split honey yield mean between the queen and workers. We measure fecundity with the number of workers, which is a count variable and for such variables Poisson distribution is a good model. This distribution has just one parameter (lambda) that represents both the mean and variance of the variable. To this end we set phenotypic variance to 100 and split it into 25 for genetic and 65 for environmental variance. As before we warn that these are just exemplary values to demonstrate the code functionality and do not necessarily reflect published values!

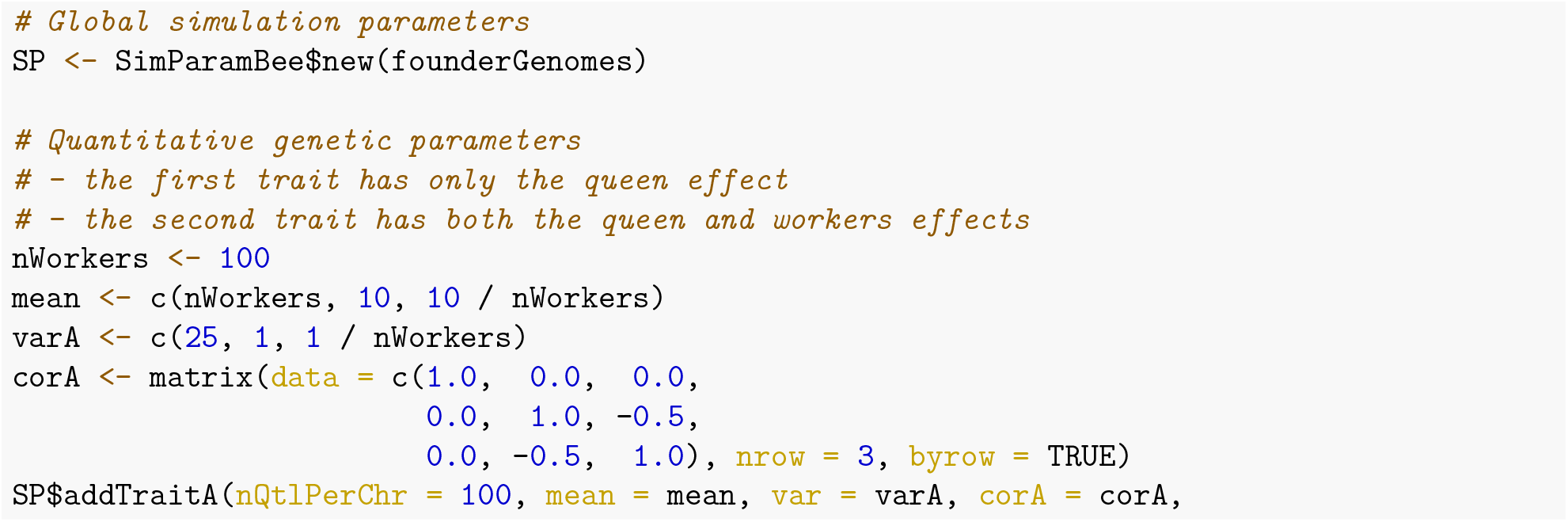

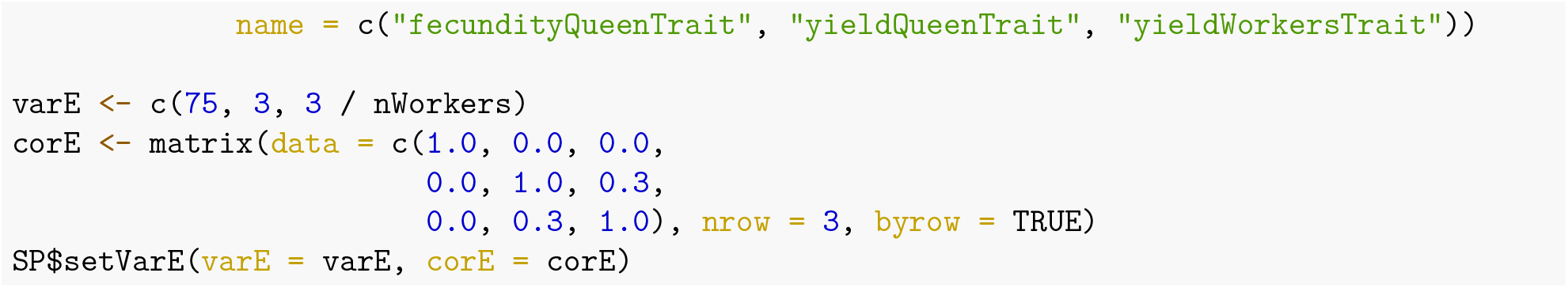

We continue by creating an apiary with 10 colonies.

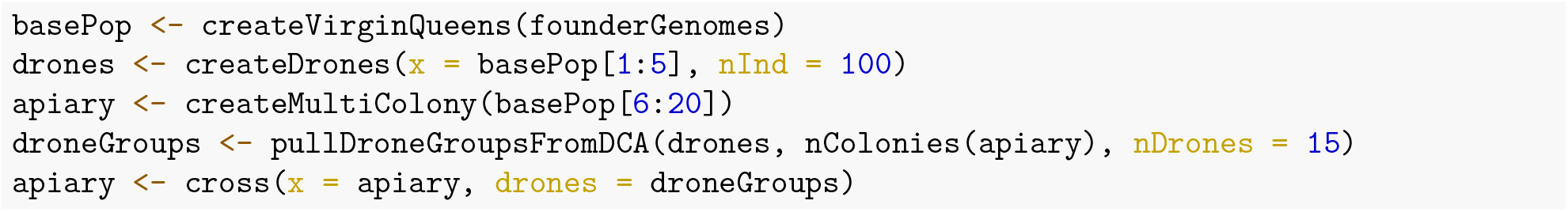

Let’s explore queen’s genetic and phenotypic values for fecundity and honey yield. The below printouts show quite some variation in fecundity between queens at the genetic, but particularly phenotypic level. This is a small example, so we should not put too much into correlations between these three variables. However, if you restart this simulation many times, you will notice zero correlation on average between `fecundityQueenTrait` and the other two traits and negative correlation on average between `yieldQueenTrait` and yieldWorkersTrait. Just like we defined in the global simulation parameters.

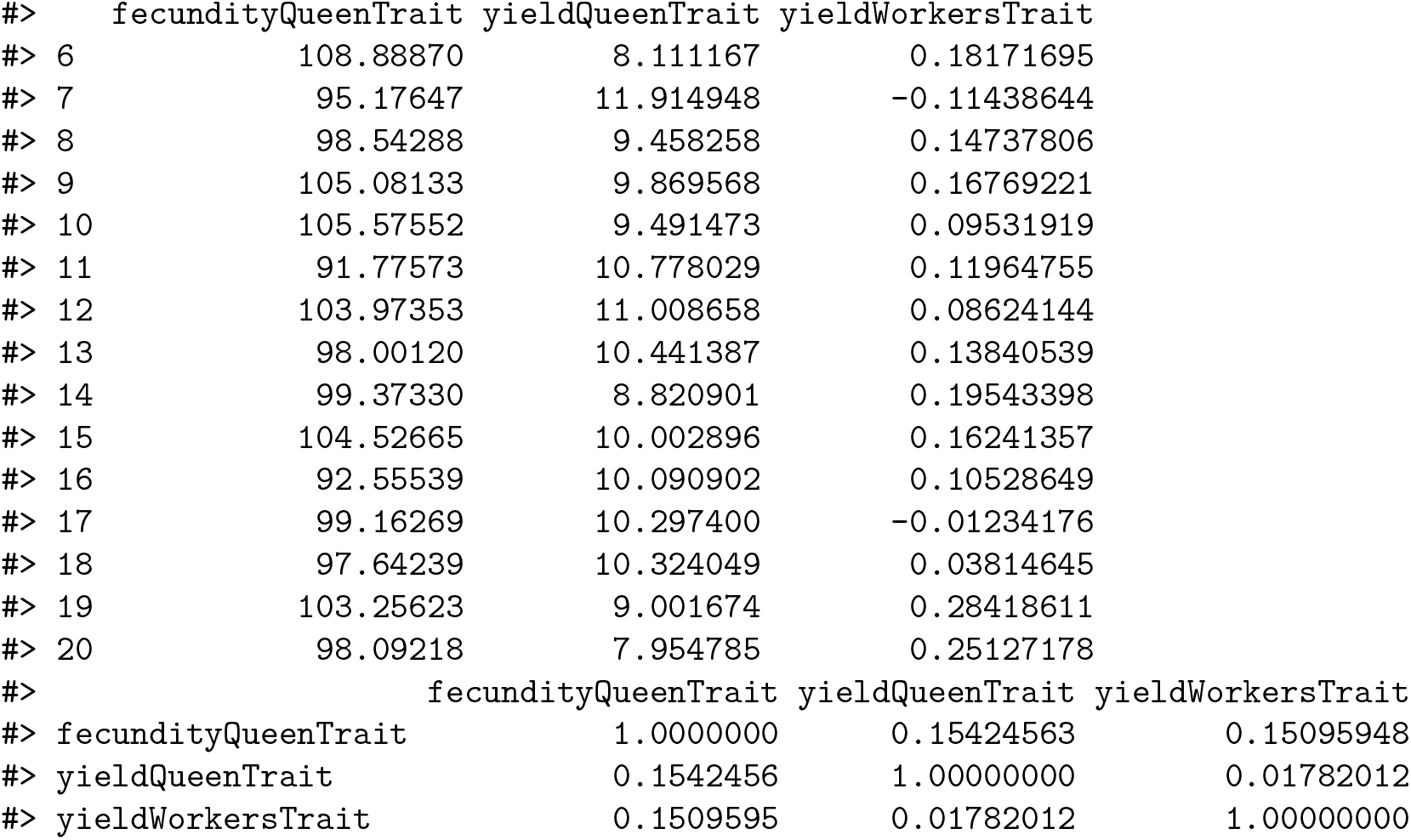

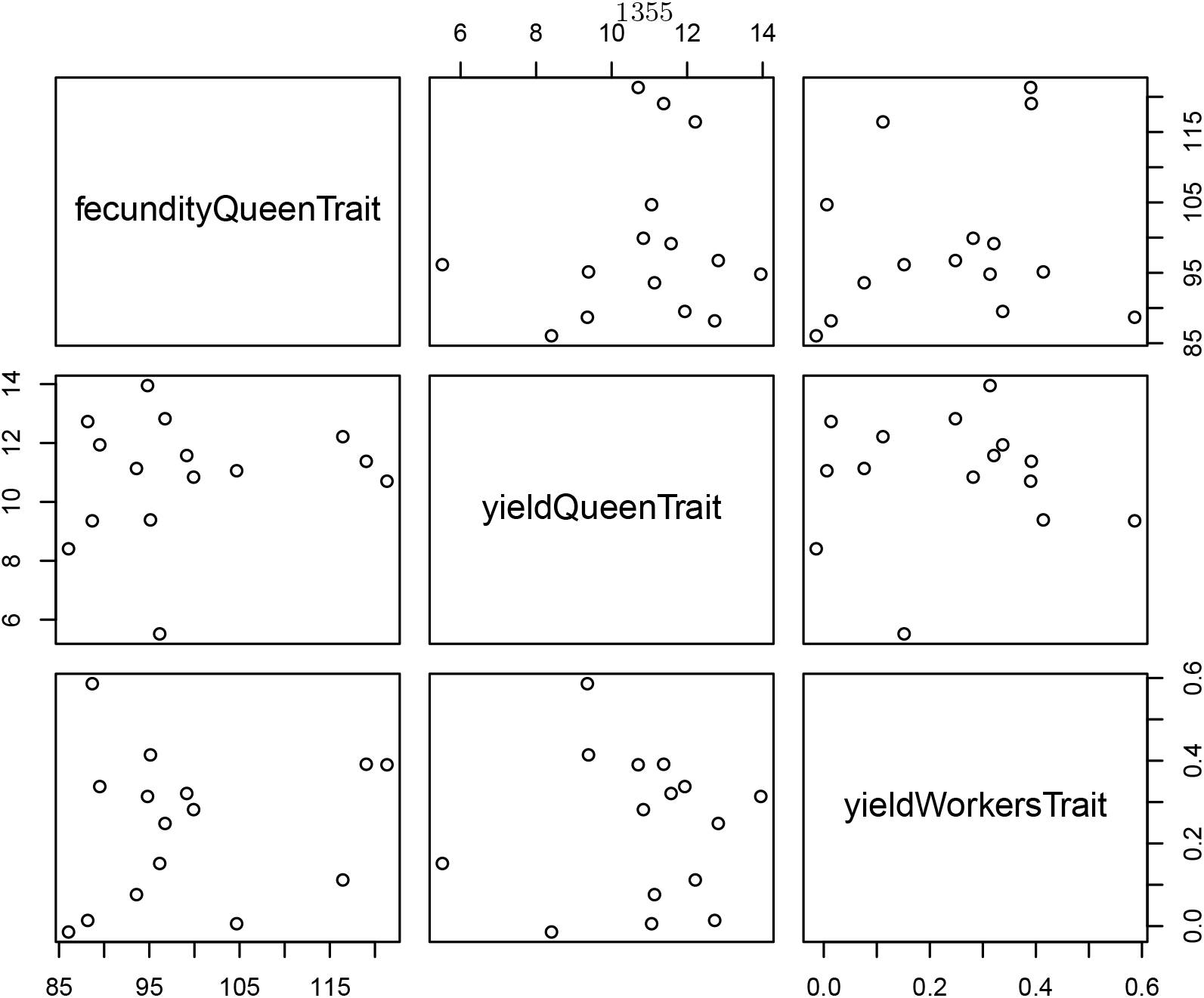

We next build-up colonies in the apiary. But instead of building them all up to the same fixed number of workers, we build them up according to queen’s fecundity. For that we use the sampling function `nWorkersColonyPhenotype()`, that samples the number of workers based on phenotypes of colony members, in our case `fecundityQueenTrait` in queens. Correspondingly, each colony will have a different number of workers. Read more about this function in it’s help page.

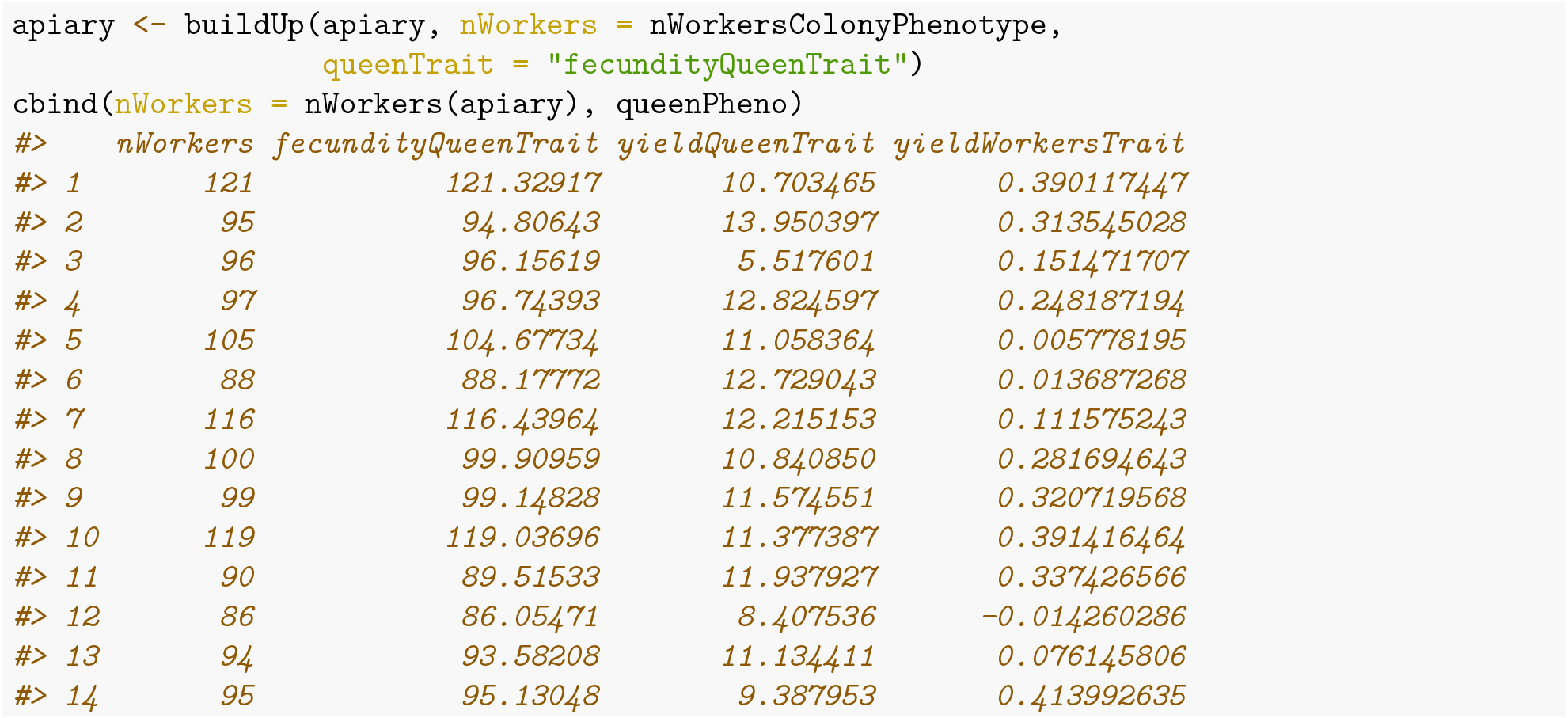

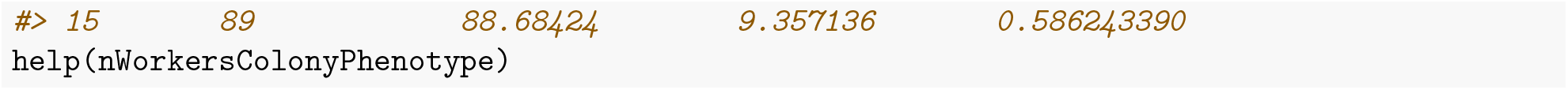

To compute the colony value for honey yield, we again employ the `calcColonyPheno()` function. Correlating the queen and colony values we will now see a positive correlation because our individual to colony mapping function sums workers effect across all workers and the more workers there are the larger the sum.

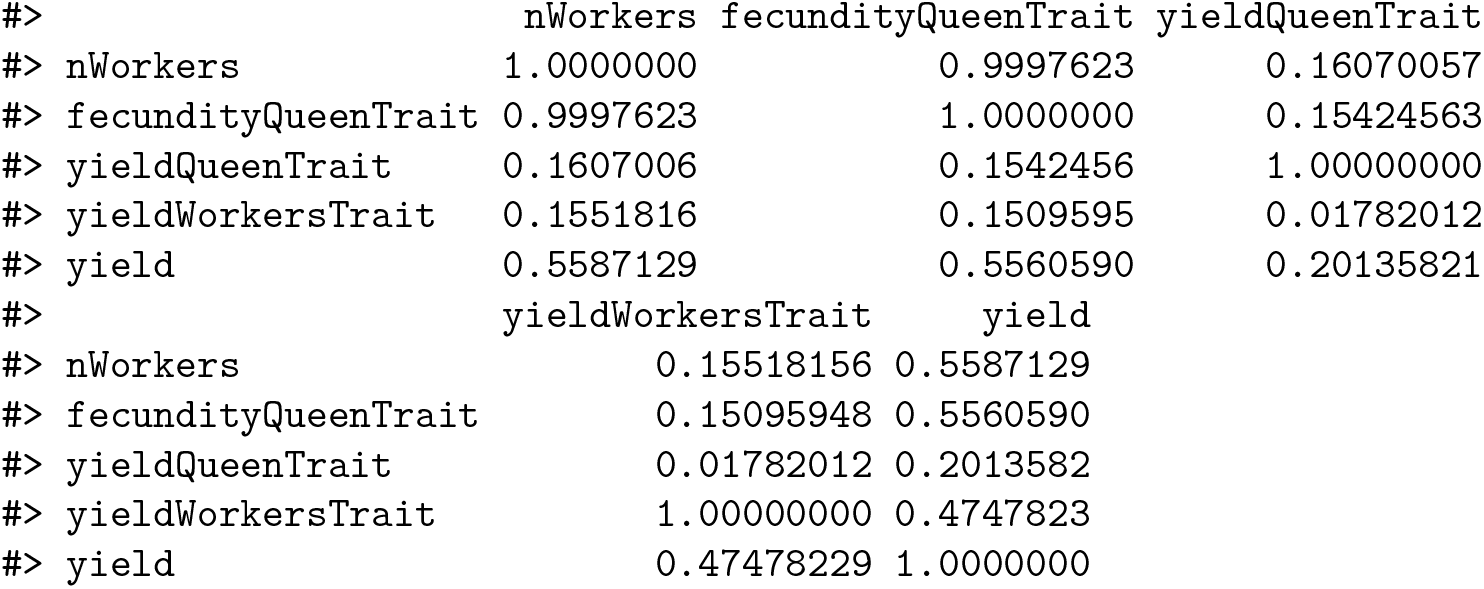

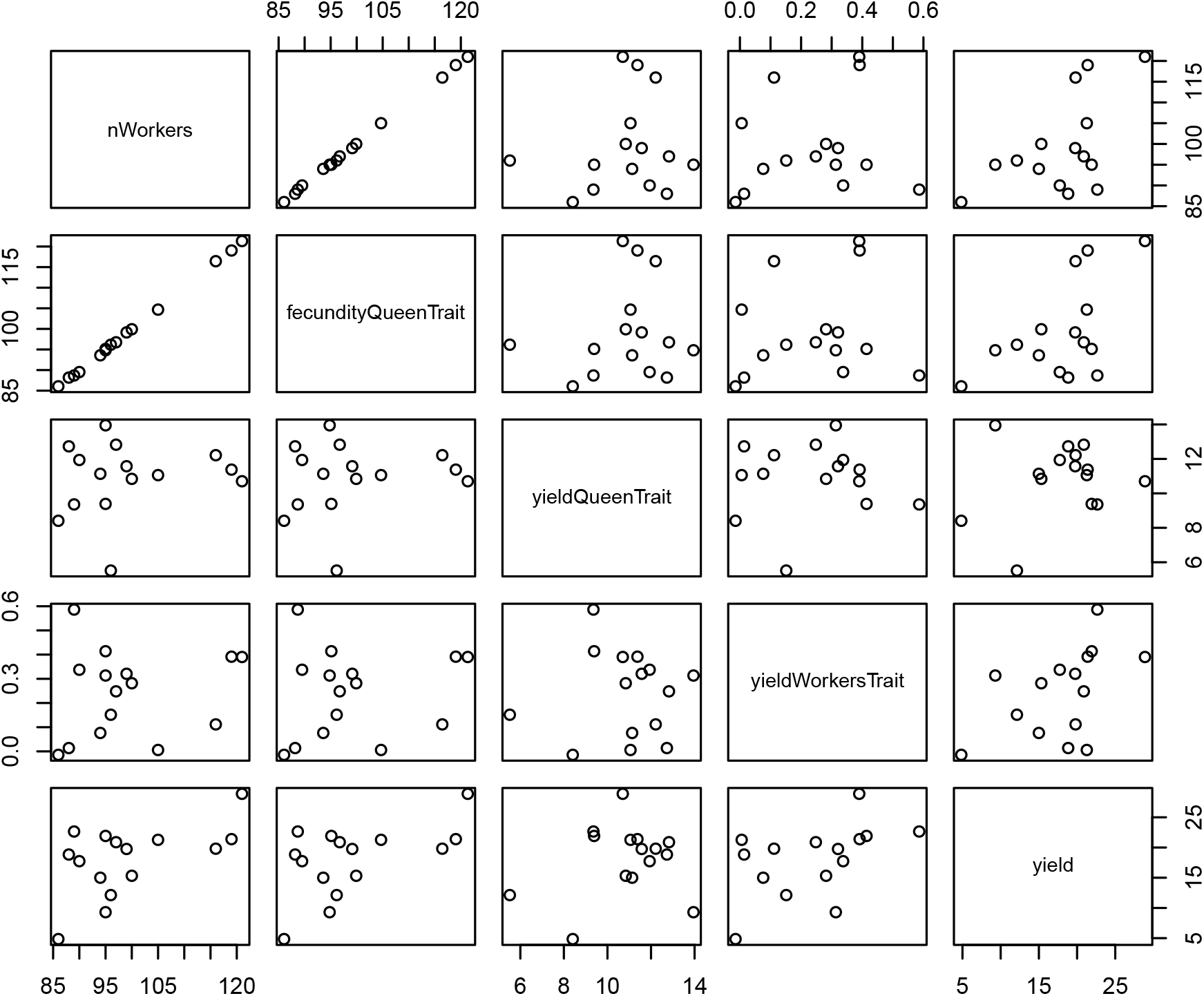

## Additional file 7 – Sampling functions vignette

### Introduction

SIMplyBee includes functions to sample various values that are expected to vary between colonies and events. These functions are used to sample numbers, usually individuals, and proportions. We can use the functions, pass them to other functions, or save them in the `SimParamBee` object so they can be used by default by other functions.

We start by loading the package:

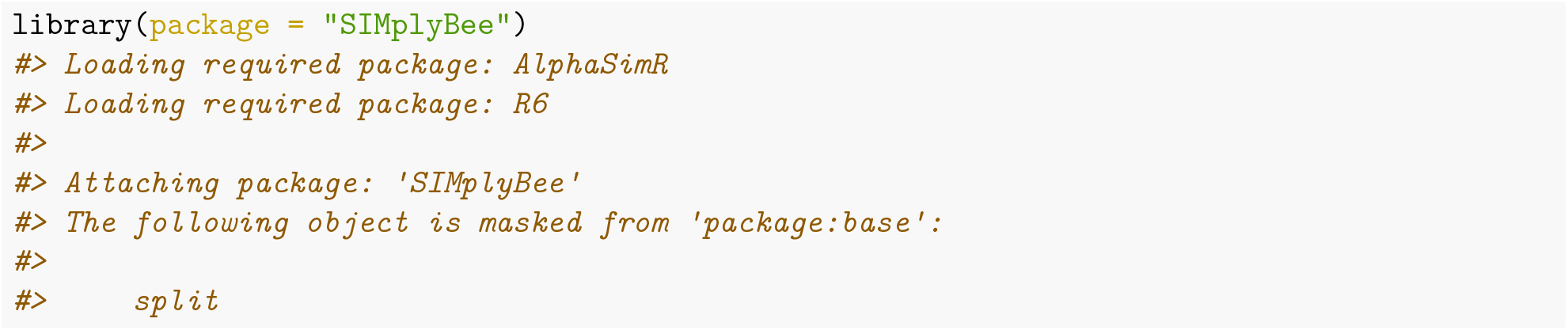

### Functions to sample numbers

First, there are functions to sample the number of caste individuals from either a Poisson or truncated Poisson distribution: `n*Poisson()` and `n*TruncPoisson()`, where `*` is either `Workers`, `Drones`, `VirginQueens`, or `Fathers`. Most SIMplyBee functions that take the number of individuals as an argument can accept these sampling functions as an input, meaning that the output of such function calls will be stochastic. These functions are useful when you want to sample a variable number of individuals around a mean, as for example when mating virgin queens with a variable number of drones.

Let’s start a simulation by creating a DCA and an apiary with 10 virgin colonies:

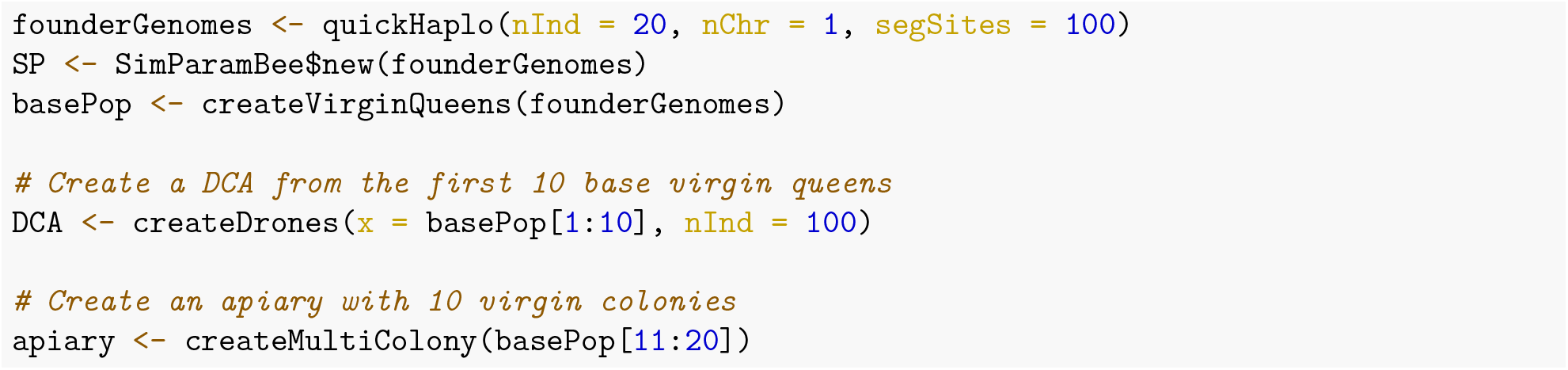

From the literature we know that virgin queens on average mate to 17 drones, but the actual number varies around this mean. Some mate with 10 drones, some with 20, etc. To resemble this variation, we can use the function `nFathersPoisson()` to sample variable number of drones from a DCA. The default average for this function is 15, but you can use any value you want. Let’s use this function to sample 1,000 values and inspect the distribution and the mean.

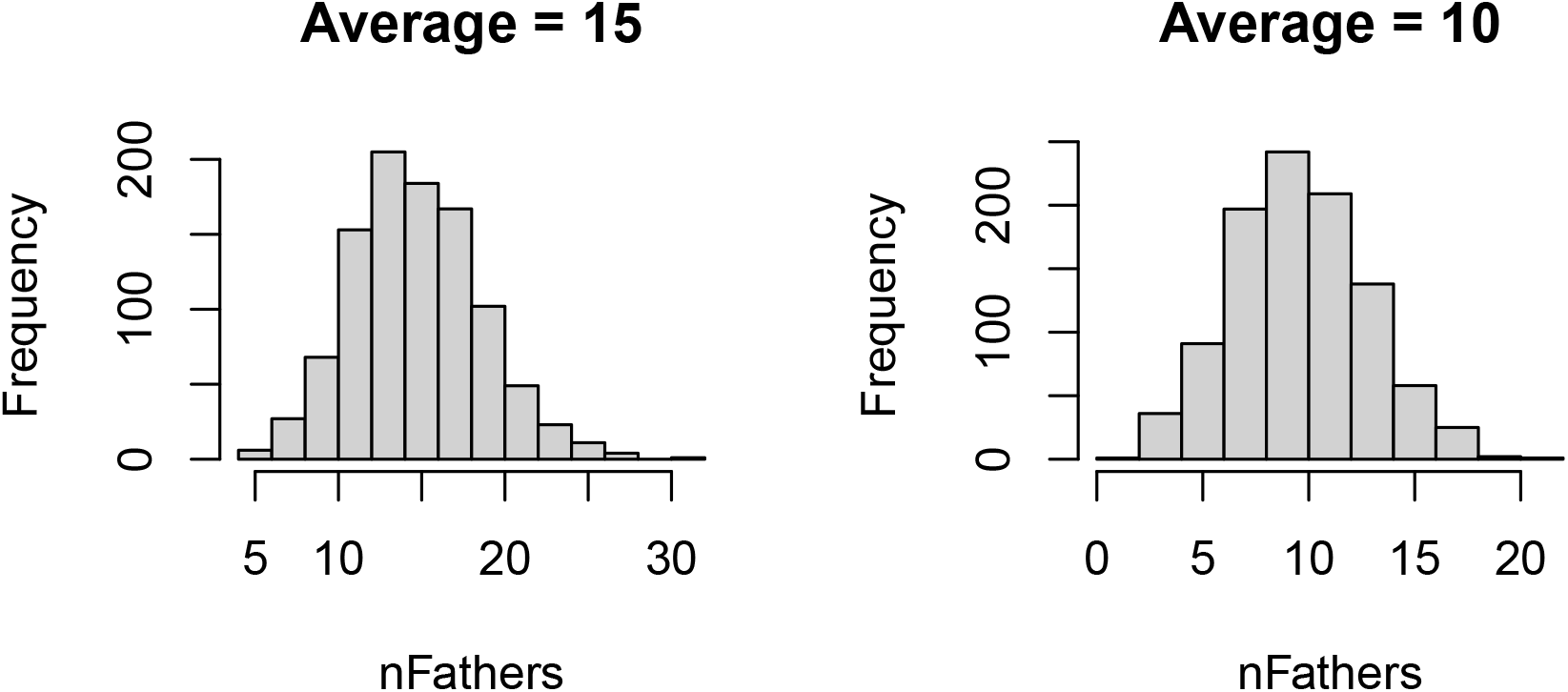

Let’s now use this functionality to sample a variable number of drones from the DCA to mate with each of the 10 virgin queens.

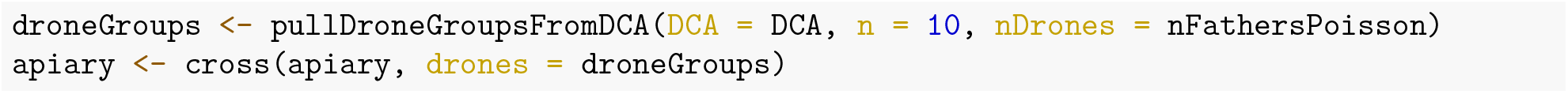

And inspect the number of fathers in each of the colony and their mean.

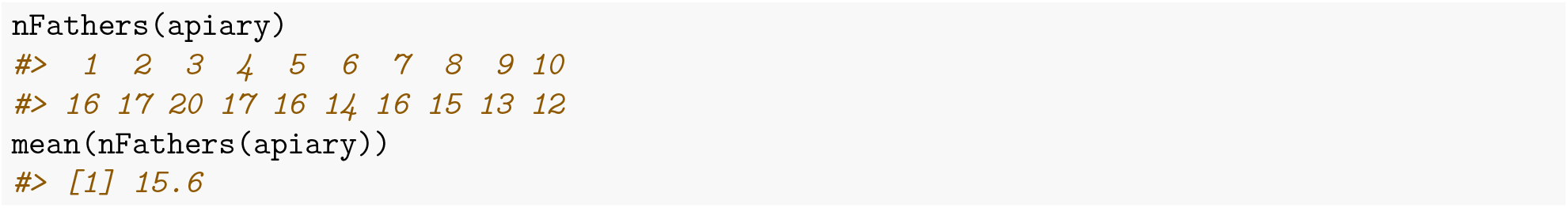

Second, we have a group of functions that will sample the number of individuals according to the colony phenotype, whatever that might be. These functions are named `n*ColonyPhenotype()`, where `*` is either is either `Workers`, `Drones`, or `VirginQueens`. An example of this would be sampling the number of workers and drones according to queen’s fecundity or honey yield. An example of this can be seen in the quantitative genetics vignette in the “Strength and honey yield” example.

### Functions that sample proportions

SIMplyBee also includes functions to sample the proportions of workers that leave or are removed when downsizing, splitting, or swarming a colony from either a uniform distribution or from a beta distribution that accounts for the number of individuals in a colony (colony strength). These functions are named `*PUnif()`, where `*` can be either `swarm`, `split`, or `downsize`. There is an additional function, `splitPColonyStrength()`, that determines the number of workers to be removed in a split according to the colony strength.

Let’s say we want to swarm all the colonies in our apiary with a variable percentage of workers that leave. We want to sample this percentage from an uniform distribution with the mean of 0.6. For this, we use `swarmPUnif()` function that takes a `min` and a `max` values and sample a value between them. By default, the `min` is set to 0.4 and `max` to 0.6. Let’s use this function to sample a 1,000 values between 0.5 and 0.7 and inspect the mean.

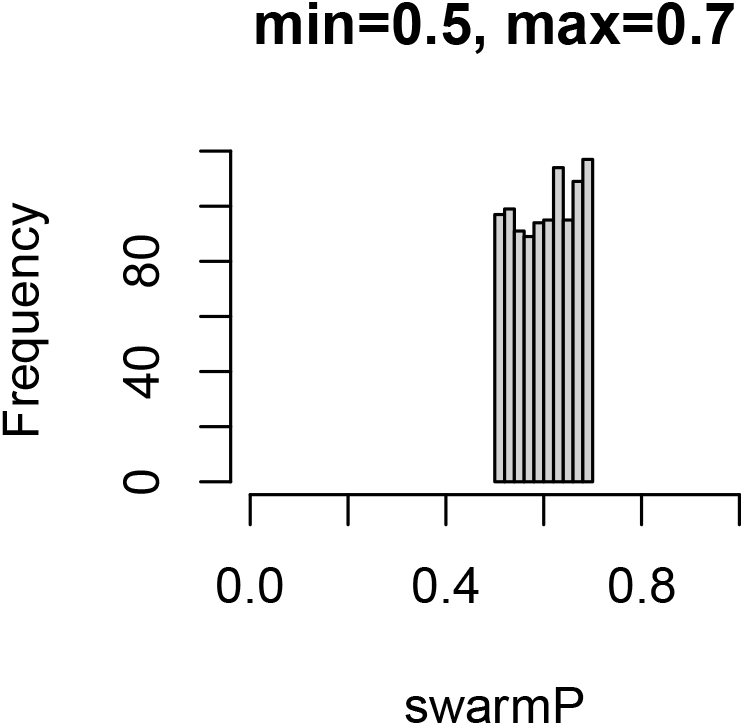

Let’s now swarm all the colonies in our apiary with a variable percentage of workers that leave.

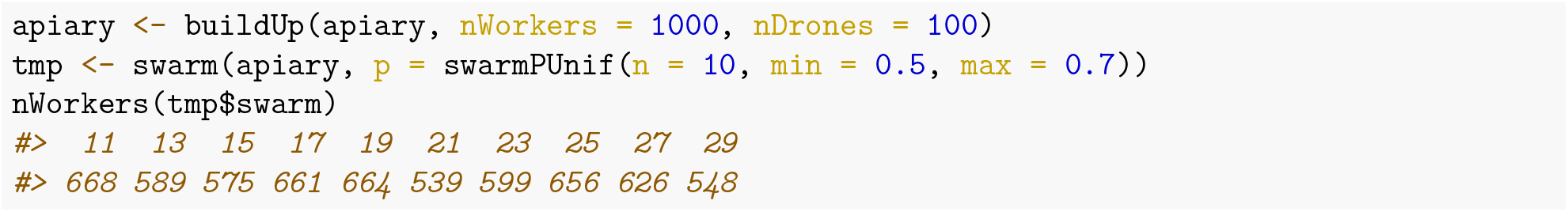

We see that each colony swarmed with a different percentage between 0.5 and 0.7.

## Notes

http://simplybee.info/

https://cran.r-project.org/web/packages/SIMplyBee/index.html

### Competing Interest Statement

The authors have declared no competing interest.

